# Quantitative Modeling of TLR Signaling Reveals Missing Negative Feedback Guiding Identification of TANK-IKKε Checkpoint

**DOI:** 10.64898/2026.08.03.742528

**Authors:** Nathan P. Manes, Fengkai Zhang, Bin Lin, Jing Sun, Sergio A. Hassan, Anthony A. Armstrong, Yuting Shao, Jessica M. Calzola, Pauline R. Kaplan-Stafford, Rachel A. Gottschalk, Matthew J. Marino, Doeun Kim, Ronald N. Germain, Martin Meier-Schellersheim, Iain D. C. Fraser, Aleksandra Nita-Lazar

**Affiliations:** Functional Cellular Networks Section, Laboratory of Immune System Biology, National Institute of Allergy and Infectious Diseases, National Institutes of Health, Bethesda, MD, USA; Computational Biology Section, Laboratory of Immune System Biology, National Institute of Allergy and Infectious Diseases, National Institutes of Health, Bethesda, MD, USA; Signaling Systems Section, Laboratory of Immune System Biology, National Institute of Allergy and Infectious Diseases, National Institutes of Health, Bethesda, MD, USA; Bioinformatics and Computational Biosciences Branch, National Institute of Allergy and Infectious Diseases, National Institutes of Health, Bethesda, MD, USA; Dept. of Immunology, University of Pittsburgh, Pittsburgh, PA, USA; Lymphocyte Biology Section, Laboratory of Immune System Biology, National Institute of Allergy and Infectious Diseases, National Institutes of Health, Bethesda, MD, USA

**Keywords:** Toll-like receptor 4, macrophage, rule-based modeling, pathway deactivation, IKKε, TANK, IRAK1, TRAF6

## Abstract

Sensing of tissue injury or infection by Toll-like receptors (TLRs) must rapidly mobilize host defense, but that activation must also be terminated on an appropriate time scale to avoid sustained, tissue-damaging inflammation. The full network of molecular interactions that governs both the rapid onset and the properly timed shutoff of TLR signaling remains incompletely characterized at a deep mechanistic level. To this end, we built a rule-based model of mouse macrophage TLR4 signaling at the molecular-interaction level, parameterized with measured protein copy numbers, RNA-seq-based abundance estimates, literature- and structure-informed reaction rates, and 979 dynamic experimental constraints, achieving a level of granularity beyond that of prior TLR models. The trained model reproduced much of the TLR4-induced NF-κB and MAP kinase response, but it consistently failed to capture deactivation of MyD88, TRAF6-associated species, and IKKα/β. Rather than treating this as simple model error, we used the recurrent failure as a biological signal that localized missing regulation to the proximalMyD88-IRAK-TRAF6 module and motivated experimental evaluation of IKKε and its scaffold TANK. Loss of IKKε enhanced transcriptional, cytokine, MAP kinase, and NF-κB responses to MyD88-specific TLR ligands, and TANK deficiency produced a similar cellular phenotype while abolishing stimulus-induced IKKε phosphorylation. Deficiency of either protein increased IRAK1and TRAF6 ubiquitination without increasing MyD88 ubiquitination, placing the inhibitory checkpoint at or immediately downstream of the IRAK1-TRAF6 ubiquitin-signaling node. Overlapping but non-identical in vivo phenotypes further supported a shared regulatory axis with additional protein-specific functions. Together, these findings illustrate a model-experiment discovery cycle in which quantitative pathway discordance identifies missing biology, revealing a TANK-dependent IKKε checkpoint that restrains MyD88-driven inflammation.

## Introduction

Toll-like receptors (TLRs) detect microbial and danger-associated molecular patterns and initiate innate host defense [1]. TLR3 signals through TRIF, TLR4 through both TRIF and MyD88, and most other TLRs through MyD88. The MyD88 branch activates MAP kinases and NF-κB through TRAF-linked kinase cascades, rapidly inducing inflammatory mediators. This activation is essential for antimicrobial defense, but if left unchecked it can instead drive chronic inflammation and autoimmune disease [2–4], which makes the mechanisms that terminate signaling as physiologically important as those that initiate it.

Understanding how the signaling termination is achieved first requires understanding how the receptor-proximal machinery is organized. TLR4 signaling spans bimolecular interactions, receptor trafficking, as we have described previously [5], and the assembly and reorganization of supramolecular organizing centers (SMOCs, also called signalosomes) [6, 7]. However, important features of receptor activation and signalosome dynamics remain unresolved [8], which complicates efforts to model the pathway accurately. Mechanistic models offer a way to test whether current pathway knowledge is sufficient to explain observed dynamics, and, where it is not, to generate experimentally verifiable hypotheses about what is missing [9, 10]. Rule-based approaches are well suited to this task in molecular signaling networks [11–13], but building a predictive model also requires reactant abundances, reaction-rate estimates, and time-resolved measurements distributed across the pathway [14, 15]; targeted mass spectrometry provides one route to this quantitative parameterization [16, 17].

Several previous TLR models have addressed complementary aspects of this problem [18–21] . An and Faeder developed an early BioNetGen model of TLR4 signaling and LPS preconditioning that represented A20- and IκB-mediated negative feedback and reproduced dose-dependent TNF responses and endotoxin tolerance; however, it was intended as qualitative dynamic knowledge representation rather than quantitative biochemical parameterization [18]. Subsequent models examined proximal TLR4–MyD88 complex assembly [20], TLR3–TLR7 crosstalk and innate immune memory [21] , and heterogeneous macrophage signaling across TLR4, TNFR, IFNAR, and IL10R inputs [19]. Building on this body of work, we sought a different level of resolution: an extensively data-constrained, molecular-interaction-level model of macrophage TLR4 signaling with biochemical parameterization of both reactant concentrations and reaction rates.

A central challenge in constructing such a model is pathway termination itself. Pattern-recognition receptor networks are highly redundant [5, 22], yet the molecular mechanisms of negative regulation remain far less complete than those governing activation [23–27]. This asymmetry matters functionally, because interacting positive and negative circuits can generate complex signaling dynamics [23], and loss of individual inhibitory proteins can cause spontaneous inflammation or autoimmunity [3, 28–31], while excessive MyD88 activity promotes systemic inflammation and endotoxin sensitivity [32].

Given this incomplete picture of negative regulation, we asked whether reproducible discrepancies between model and experiment could be used to localize missing deactivation biology, rather than being treated merely as fitting error. Our model reproduced much of TLR4 activation but repeatedly failed to reproduce proximal signal termination, and this localized failure led us to test candidate molecules for a role in this missing negative regulation. TANK is a TRAF-associated scaffold and established negative regulator of TLR signaling, although its inhibitory mechanism is incompletely defined [33]. The related kinase IKKε is best known for interferon-dependent antiviral functions [34] and also participates in IL-17 signaling [35]; combined inhibition of IKKε and TBK1 enhances IL-1-driven responses [36], but an inhibitory role for IKKε alone in a functional MyD88-dependent inflammatory context has not been established. Our experiments with IKKε-deficient BMDM and with TANK-deficient BMDM showed that these two molecules each contribute to this inhibitory activity and suggest that TANK acts upstream of IKKε. These findings reveal the value of the detailed modeling we report here, showing how a limitation to that model led to discovery of additional regulatory components critical for avoiding pathologic inflammation.

## Methods

### Outline of the Targeted LC-MS Experiments

To parameterize the model with measured protein abundances, we first generated a targeted LC-MS dataset, described in detail in our Data Descriptor article [37]. Target proteins were selected for absolute quantitation to study the chemotaxis [16] and TLR4 signaling pathways in mouse macrophages. Tryptic target peptides were selected, and peptide standards were purchased and analyzed using data-dependent acquisition (DDA) LC-MS. The resulting spectra were used to make spectral libraries and scheduled parallel reaction monitoring (PRM) LC-MS assays. Heavy isotope-labeled, purified, quantitated peptide standards were then used in these PRM LC-MS assays to measure the absolute abundance (copy/cell) of the target proteins; these values, together with an RNA-seq dataset from our laboratory, were used to estimate proteome-wide protein absolute abundance.

### Protein and Peptide Selection for Targeted LC-MS

Target proteins were selected based on our earlier mechanistic review of PRR signaling pathways [5]. Multiple rounds of assay development were performed, and at least two target peptides were used to quantitate each target protein, if possible. Peptides were manually selected to be both proteotypic (efficiently identified and quantitated by LC-MS), and quantotypic (the peptide quantity is an accurate measurement of the target protein quantity). Each peptide was fully tryptic with no missing cleavages. Internal KP and RP sites and neighboring trypsin cleavage sites (e.g., …AAKRAA…) were avoided. Proteolysis was required to produce exactly one copy of each target peptide from each target protein (Ile/Leu substitution was considered). Preferentially, the peptide length was 5-20 amino acids. Cys and Met (oxidation), Asn and Gln (deamidation), amino-terminal Gln (formation of pyroglutamate), and N/C-terminal peptides (prone to PTMs) were avoided. Other covalent modifications and natural genetic variants annotated in UniProt [38] were avoided including those at the trypsin sites. Preferentially, each peptide was designed so that it would be useful for an assay of the human ortholog (the trypsin sites and Ile/Leu substitution were considered). The peptide was required to be unique to the target protein (or to a small set of closely related homologs; Ile/Leu substitution and RNA-seq splice-isoform abundance data were considered). The target peptide was required to be proteotypic by using LC-MS proteomics data from The Global Proteome Machine [39], from the National Institute of Standards and Technology [40] and from our own laboratory [41–43].

### Peptide Standards for LC-MS

Custom peptide standards were purchased from JPT Peptide Technologies GmbH (Berlin, Germany) and Thermo Fisher Scientific Inc. (Waltham, MA). Purified quantitated heavy-labeled (^13^C_6_ ^15^N_4_ Arg, ^13^C_6_ ^15^N_2_ Lys) internal peptide standards were purchased for quantitative analyses. When possible, each internal peptide standard was flanked by leading and trailing regions (each was three amino acids in length) to mimic the cleavage sites of the target protein. An aliquot (1 nmol) of each quantitated peptide standard was dissolved in 20% v/v acetonitrile (ACN), vortexed for 2.5 min, and bath sonicated for 5 min at room temperature. Peptides were pooled and concentrated in a SpeedVac vacuum concentrator (Thermo Fisher Scientific Inc.) at 40 °C to a final concentration of 4 μM (of each peptide) in 20% v/v ACN. This procedure was performed in duplicate, and the duplicates were then combined.

### Sample Preparation for LC-MS

Because the quantitative assays required primary and immortalized macrophages, we prepared them as follows. Bone marrow-derived macrophages (BMDMs) from C57BL/6 mice were prepared methods that we and others have described previously [37, 44–47]. These animal-based procedures were conducted under the protocol LISB-17E approved by the NIAID Animal Care and Use Committee following the guidelines of the Office of Animal Care and Use (OACU) on 2 January 2021. Briefly, bone marrow progenitors from femurs and tibias were differentiated into BMDMs for six days in complete medium supplemented with 60 ng/ml mouse MCSF (R&D Systems Inc, Minneapolis, MN). RAW264.7 (American Type Culture Collection TIB-71) mouse macrophages and immortalized BMDMs (iBMDMs) [48] were cultured in Dulbecco’s Modified Eagle’s medium (DMEM; 4.5 g/L glucose; 4 mM L-glutamine), 20 mM HEPES, 1 mM sodium pyruvate, and 10% v/v fetal bovine serum (not heat inactivated). Cells were counted manually or using a Cellometer Auto T4 (Nexcelom Bioscience LLC, Lawrence, MA; cell diameters were also measured). Trypan blue staining was used to measure viability.

Sample preparation followed our published protocol [49]. Briefly, cells were lysed and homogenized using a Bioruptor Plus (Diagenode Inc., Denville, NJ) in the following lysis buffer: 8 M urea, 100 mM HEPES·NaOH, pH 8, 10 μM bestatin, 10 μM pepstatin A, 1x Halt phosphatase inhibitor cocktail (provided as a 100x solution; Thermo Fisher Scientific Inc.). Cell lysate protein concentrations were measured using a bicinchoninic acid (BCA) assay kit (Thermo Fisher Scientific Inc.). Either 0 μg or 50 μg (protein mass) of BMDM homogenate was mixed with 0, 50, 500, or 5000 fmol (of each peptide) of the quantitated heavy labeled peptide standards mixture. Dithiothreitol (DTT) was added to reduce protein cystines, and iodoacetamide was added to alkylate protein cystines. The samples were diluted with 100 mM HEPES·NaOH pH 8 such that the final urea concentration was ≤1 M, and they were digested with sequencing grade modified trypsin (Promega Corp., Madison, WI) at 37°C. To halt the reactions, 1% v/v formic acid (FA; final concentration) was added. The samples were microcentrifuged (10,000 × g for 20 min at room temperature) and underwent solid-phase extraction (SPE) using Sep-Pak C-18 SPE columns (1 ml, 100 mg C-18 media, Waters Corp., Milford, MA; Mobile Phase A = 0.1% v/v FA, Mobile Phase B = 0.1% v/v FA, 80% v/v ACN).

### Targeted LC-MS

PRM LC-MS was performed using an UltiMate 3000 nanoLC coupled to a Q Exactive HF mass spectrometer (Thermo Fisher Scientific Inc.). The samples were trapped onto a trap column (Acclaim PepMap 100, C-18, 75 μm i.d., 2 cm length, Thermo Fisher Scientific Inc.) and resolved using an EASY-Spray column-ESI-tip cartridge (PepMap RSLC C18, 75 μm i.d., 50 cm length, 2 μm bead diameter with 100 A pores, Thermo Fisher Scientific Inc.). Analytes were separated using a 60 min linear gradient (2-30% Mobile Phase B; Mobile Phase A = 0.1% v/v FA; Mobile Phase B = 0.1% v/v FA in ACN; flow rate = 200 nl/min) and electrosprayed at 1.8 kV into the MS. Each MS1 scan (200-2000 m/z, Resolution = 60k) was followed by fourteen PRM scans (Resolution = 60k, maximum isolation time = 110 ms, normalized collision energy = 27), with LC-MS scheduling performed using Dynamic Retention Time.

### LC-MS data analysis

DDA LC-MS spectra were analyzed using Proteome Discoverer (v. 2.2.0.388, Thermo Fisher Scientific Inc.) using Sequest HT and Mascot (v. 2.6.2, Matrix Science, Boston, MA) to perform database searching against a FASTA of the target peptides. The static modifications were carbamidomethylation (C) and the heavy-isotope labels (K, R). The dynamic modifications were oxidation (H, M, W), acetylation (peptide N-term), deamidation (N, Q), Gln conversion to pyro-Glu (peptide N-term Q), and carbamidomethylation (peptide N-term). The results were imported into Skyline (64-bit, v. 4.2.0.19072) [50] to create spectral libraries for LC-PRM assays. The LC-PRM spectra were imported into Skyline (64-bit, v. 19.1.0.193), and target protein abundance values in units of copies/cell were calculated.

### Protein Abundance Estimation using RNA-seq

To extend these measurements across the full proteome, BMDMs were also analyzed using RNA-seq as described [46] (NCBI GEO accession GSE70510). For each sample and gene, the abundance in units of transcripts per million (TPM) was calculated [51]. A linear regression was performed using the Log_10_-transformed TPM values and the LC-PRM Log_10_-transformed copy/cell values. The resulting equation was used to estimate protein copies/cell values for the entire mouse BMDM proteome.

### Pathway Modeling Overview

With reactant concentrations in hand, we next built the reaction network. The mouse macrophage TLR4 signaling pathway was modeled at the molecular reaction level using the Simmune software suite v. 2.4.3-20260211 (build 7531) [52, 53] running on a Linux workstation installed with Ubuntu v. 22.04. Simmune was developed for rule-based biological pathway modeling. SimModeler was used to construct the reaction network, SimAnalyzer was used to perform pathway simulations to train the model, Simmune Network Viewer was used to visualize the reaction network, and Simmune Python Analysis Module (version compiled 2026-Feb-12) to analyze and visualize the simulation results. Simmune requires only a biochemical description of the pathway (the network of reactions, reaction rate constants, and reactant concentrations) and models molecular association, dissociation, and transformation reactions using law of mass action kinetics, along with production and degradation reactions. The resulting network contained 445 unknown variables (57 molecule concentrations and 388 reaction rate constants) and 803 ODEs, averaging 7.26 terms each (minimum = 3, maximum = 136, including the derivative term), with no reaction dead-ends. SimAnalyzer was used to explore this high-dimensional parameter space and score parameter sets against 979 experimental time-course ratios (2.2 ratios per unknown variable), training all model parameters simultaneously. Simmune modeled the extracellular and intracellular environment and automatically adjusted kinetics equations for reactants that switched between freely diffusing and membrane-associated states. Simmune terminology is capitalized and italicized below for clarity.

### Reaction Network Modeling

We constructed the detailed mechanistic network of the TLR4 signaling pathway based on our earlier description [5]. SimModeler was used to build the TLR4 signaling pathway reaction network visualized by the Simmune Network Viewer (Supplemental Figure S4A). Each Simmune *Molecule* was composed of one or more *Molecular Components* (e.g., intracellular and extracellular domains; the *Molecules* are described in Supplemental Table S1). Each *Molecular Component* optionally contained *Binding Sites* (which could be *Bound* or *Unbound*) and *Features* (which could be *On* or *Off*, e.g., a phosphorylation site). Each Simmune *Complex* contained one or more *Molecules* bound together. A Simmune *Complex* can be *Well-Defined* or in a *Pattern*. Each *Well-Defined Complex* had all its *Features fully defined (on or off)* and *Binding Sites Unbound*. A *Pattern Complex* is a not *Well-Defined Complex*, which that could potentially match multiple *Well-Defined Complexes*. Each *Pattern Complex* could have one or more ambiguous *Feature* states (set to *Don’t Care*, meaning compliance with both *On* and *Off*) and/or have one or more ambiguous binding sites (either set to *Don’t Care*, or set to *Bound* but the binding partner was not specified). For example, pp-ERK1 having any binding state was one of the *Pattern Complexes*. Simmune required complexes of interest (e.g., pp-ERK1) be defined prior to performing simulations and model training.

Using this formalism, SimModeler was used to design the TLR4 signaling pathway reactions (Supplemental Table S2). An *Association Reaction* was used to model two *Complexes* undergoing association to become one *Complex*, with *Features* optionally changing their state: *Association Rate* (in units of M/s) = k_on_ (1/(M×s)) × *Complex A Concentration* (M) × *Complex B Concentration* (M). All the *Association Reactions* were defined as three-dimensional reactions (Simmune automatically converted these into two-dimensional reactions if appropriate). Because of the way that the *Association Reactions* were defined, the maximum possible number of *Molecules* in a *Complex* was set to twelve. A *Dissociation Reaction* was used to model one *Complex* undergoing dissociation to become two *Complexes*, with *Features* optionally changing their state: *Dissociation Rate* (M/s) = k_off_ (1/s) × *Complex Concentration* (M). All the *Dissociation Reactions* were “normal” (i.e., none were restricted to being intra-complex). All the *Association* and *Dissociation Reactions* had “no orientation” (“cis-binding” and “trans-binding” orientations were not used; they are used for modeling cell-cell interactions).

A *Transformation Reaction* was used to model one *Complex* undergoing alteration of the state of its *Features*: *Transformation Rate* (M/s) = k_trans_ (1/s) × *Complex Concentration* (M). A *Dissociation Reaction* was used to model simultaneous transformation and dissociation (e.g., Michaelis-Menten kinetics) with potential simultaneous transformation, and k_cat_ was used to represent these rate constants. Enzymatic reactions were accordingly modeled as unidirectional two-step reactions: an *Association Reaction*, and a *Dissociation Reaction* which included a simultaneous catalysis event (e.g., phosphorylation). Protein phosphatases were not explicitly modeled because TLR-mediated upregulation of dephosphorylation remains poorly understood [25, 26]; instead, each dephosphorylation reaction was modeled using a *Transformation Reaction* attributable to unknown phosphatases.

Two additional reaction types captured gene expression. A *Production Reaction* was used to model a *Precursor Complex* (e.g., IκBα mRNA) generating an increase in the concentration of a *Product Complex* (e.g., IκBα protein): *Production Rate* (M/s) = k_prod_ (1/s) × *Precursor Complex Concentration* (M). A *Degradation Reaction* was used to model a *Complex* undergoing a reduction in its concentration: *Degradation Rate* (M/s) = k_deg_ (1/s) × *Complex Concentration* (M). Finally, *Rate Constraints* were used to ensure that two related reaction rate parameters were constrained to be equal (e.g., p50:p50 k_on_ were required to be equal in the cytosol and nucleus). Details of homomer modeling, homolog modeling, polyubiquitination modeling, macrophage modeling and model simulation protocols are described in the Supplemental Methods.

### Initial Model Reactant Concentration Values

Having defined the reaction rules, we next assigned starting concentrations for every reactant (Supplemental Table S1). PRM LC-MS results were used to calculate initial protein concentration values; where no LC-MS measurement was available, RNA-seq-based abundance estimates were used instead. The model also contained four non-protein reactants (LPS, mRNA_IkBa, mRNA_IkBb, and PI45P2_x4CG): LPS concentration was set using the modeling protocol, PI(4,5)P2 concentration was based on experimental measurements (Supplemental Table S1), andIκBα/β mRNA concentrations were set equal to the corresponding protein concentrations (discussed below).

### Experimental Reaction Rates and Rate Constraints for Modeling

Reactant concentrations alone are insufficient without corresponding reaction rates, so we performed a literature review to identify experimentally measured rate constants for the TLR4signaling pathway. This yielded 63 k_on_, 65 k_off_, 28 k_trans_, 114 K_D_ (dissociation constant), 112 K_M_ (Michaelis constant), and 112 k_cat_ measurements were found (Supplemental Table S3). Here, k_cat_ is a rate constant for a simultaneous transformation and dissociation reaction (e.g., Michaelis-Menten kinetics). Some of these measurements were for reactions that were ultimately excluded from the model, and replicate measurements were averaged as geometric means. This process resulted in 19 k_on_, 24 k_off_, 22 k_trans_, 48 K_D_, 32 K_M_, and 33 k_cat_ values used for the model (Supplemental Table S4).

The k_on_, k_off_, k_trans_, and k_cat_ values served as initial values of the reaction rate parameters. The dissociation constant K_D_ of a binding reaction is equal to k_off_ / k_on_. The Michaelis constant K_M_ of a Michaelis-Menten enzyme is equal to (k_off1_ + k_cat_) / k_on1_, where k_on1_ and k_off1_ are the enzyme-substrate association and dissociation rate constants, respectively. The K_M_ and K_D_ values could not be used to directly parameterize the Simmune model. Instead, each could be used as a *Rate Constraint* to solve for an unknown parameter (e.g., k_on_ could be calculated using K_D_ and k_off_). As described above, *Rate Constraints* were sometimes needed to ensure that two related reaction rate parameters were equal (e.g., p50:p50 k_on_ in the cytosol and nucleus). Note that during model training (described below), the genetic algorithm implemented in SimAnalyzer ignores all *Rate Constraints*.

### Protein Complex Structure Modeling

For rate constants that could not be found in the literature, we turned to structural modeling. Protein complex structures were each modeled using up to three levels of complexity: as a simple two subunit complex, as medium-sized complex (maximum of six subunits), and as a large-sized complex (maximum of eight subunits). For example, the activated form of TRAF6 was modeled as TRAF6:Ubc13, [TRAF6]_2_:Ubc13:Uev1a, and [TRAF6]_6_:Ubc13:Uev1a. None of the protein complex models included a known transmembrane region (complexes were designed to be exclusively intracellular or extracellular; only the intracellular or the extracellular region of each transmembrane protein was included). If a protein was reported to have two structured domains tethered by a structurally disordered region (e.g., MyD88), and only one of the domains was known to directly bind to a complex (e.g., to MAL), then only the relevant domain(s) were modeled.

AlphaFold-Multimer v. 2.3.2 [54] was used to predict protein complex structures. AlphaPickle v. 1.5.4 (https://github.com/mattarnoldbio/alphapickle) was used to extract the AlphaFold results. There were 630 AlphaFold analyses performed (25 models/analysis). For each analysis, the models were ranked using the Model Confidence score (this is the default method) which was defined as (0.8 × ipTM) + (0.2 × pTM), where pTM is the predicted template modeling score (of the whole protein complex) and ipTM is the interface pTM (the pTM but only considering the PPI interface residues) (AlphaFold-Multimer produced one pTM score and one ipTM score per model). Of the 630 analyses, three were designed incorrectly and were discarded, and 16 failed (timed-out or memory limit exceeded).

### Protein Complex Structure Model Refinement

In this section, a chain refers to an individual protein molecule. Each model was refined by removing low-confidence regions according to the criteria described below (see SI for details). The main goals were twofold: to preserve high-quality interchain interfaces for rigid-body simulations, as required here for Brownian dynamics simulations used to estimate association rates, and to maintain the structural integrity of each chain if molecular dynamics simulations are needed (although not performed in this work, this criterion is included in the pipeline for completeness).

First, each residue was classified based on pLDDT as confident (pLDDT > 60) or low-confidence (pLDDT ≤ 60), defining the initial set of exclusively confident or low-confidence sequence segments, {c/u}_1_. Then, short low-confidence segments (<9 residues), which may represent genuine loops, linkers, or flexible regions, were reclassified as confident to generate the refined set {c/u}_2_. Subsequently, low-confidence segments near interchain interfaces were reevaluated. Specifically, residues within low-confidence segments were reclassified as confident if the distance between their Cα atoms and those of any residue in a confident segment belonging to a different chain was ≤7 Å. If this resulted in more than 60 % of residues in a segment being reclassified, the entire segment was retained as confident. This generated {c/u}_3_. Next, terminal confident segments were reevaluated and reclassified as low-confidence if they flanked low-confidence segments and lacked contacts (≤7 Å between Cα atoms) with any confident segment in {c/u}_3_. This step removed terminal domains connected to the core structure through flexible or disordered linkers, which may behave as loosely tethered regions and are unlikely to contribute to *k*_on_ prediction.

The resulting structures can be used directly in rigid-body simulations (BD or MC) without further processing, as relevant interchain interfaces are preserved and most extraneous domains that may interfere with association/dissociation processes are removed. The criteria also aim to maximize the likelihood of retaining intact chains suitable for MD simulations; however, fragmented chains may still be present, and visual inspection is recommended, as they could disintegrate during dynamics. Depending on the application, such fragmentation may not be a concern.

### Initial Association Rate Values from Molecular Simulation

The refined and unrefined models generated by AF were used to perform molecular simulations to predict association rate constants. Both the unrefined and refined models were analyzed using molecular simulations. The models were split into two complexes (substrates A and B). There were seven AF structures, each corresponding to two association reactions (for each, the protein complex was split in two ways). Consequently, there were 618 complexes and 1,235 molecular simulation analyses. In 570 of the 618 complexes, there was no ambiguity when identifying substrates A and B. In the remaining 48 complexes, there were multiple copies of the same protein that were partitioned between two different substrates, so there was some ambiguity when isolating substrates A and B. In 19 of the 48 complexes, manual inspection of the structure was sufficient to achieve a highly confident identification of substrates A and B. There were 29 instances where manual isolation of substrates A and B was less confident.

Brownian dynamics simulations are a powerful method for predicting association rate constants [55]. PDB2PQR v. 2.1.1 [56] (http://www.poissonboltzmann.org/) and PROPKA v. 3.0 revision 182 [57] were used to determine atomic charge states, whereas APBS v. 1.1.0 [56] (http://www.poissonboltzmann.org/) was used to generate electrostatic potential grids. TransComp v. 1.0 [58] was used to predict protein-protein k_on_ values. Standard physiological conditions were used (pH 7.4, 37 °C, ionic strength = 150 mM), except for TransComp, which is optimized and hardwired for 298 K (24.85 °C).

There were 1,235 TransComp analyses of TLR4 pathway PPIs (10,000,000 trajectories per TransComp analysis), resulting in 276 k_on_ predictions TransComp is programmed to avoid producing low confidence k_on_ predictions, and most TransComp analyses did not produce a k_on_ prediction. Predicted k_on_ values less than 10^4^/(M×s) were discarded. One predicted k_on_ was 2.15×10^10^/(M×s) and was replaced with the upper cutoff value of 10^10^/(M×s). In the original TransComp publication [58], the software was validated using an experimental dataset with measured k_on_ values ranging from 2.1×10^4^/(M×s) to 1.3×10^9^/(M×s). Additionally, the full theoretical range of PPI k_on_ values is approximately 1-10^10^ /(M×s) [58–62]. After filtering the data and removing redundant predictions, there were 122 non-redundant k_on_ predictions (Supplemental Table S5).

### Production and Degradation Reaction Modeling

Given the targeted proteolysis and recovery of IκBα and IκBβ during TLR4 signaling [46], *Production* and *Degradation Reactions* were defined for these two proteins. The protein *Production Rate* (in units of M/s) was equal to k_prod_ (1/s) × *mRNA concentration* (M). The protein *Degradation Rate* (M/s) was equal to k_deg_ (1/s) × *Protein Concentration* (M). The mRNA reactants were only used to produce the corresponding proteins, so the initial concentration value of each mRNA was set to equal the concentration of each corresponding protein for simplicity (222 nM IκBα, 1629 nM IκBβ). The maximum IκBα degradation half-life in mouse BMDMs was estimated to be t_1/2_ = 20 min (thus, initial k_deg_ = ln(2)/t_1/2_ = 0.000578/s) based on [46]. The same initial value was used for IκBβ k_deg_. Unbound IκBα/β and K48 polyubiquitinated IκBα/β were both targeted for degradation. For each IκB, the initial value of k_prod_ was set to be equal to k_deg_.

Because IRAK1 and IRAK4 have also been reported to multiply phosphorylate MAL, resulting in its rapid degradation [63–65], the initial p-MAL k_deg_ parameter was set to 0.05/s.

### Other Initial Model Reaction Rate Values

Many of the rate parameters (k_on_, k_off_, k_trans_, and k_cat_) of the core TLR4 pathway model had no experimental measurement or simulation-based estimate available by the methods above. For each parameter type, we therefore calculated the empirical cumulative distribution function (eCDF) of the experimentally measured values (Supplemental Table S4; Supplemental Figure S2). reasoning that, absent hidden biases, this eCDF should approximate the corresponding distribution across the entire core pathway. From each eCDF, a default initial value was determined ad hoc: 10^6^/(M×s) for k_on_, 0.02/s for k_off_, 0.05/s for k_trans_, and 0.2/s for k_cat_ (Supplemental Figure S2).

A related issue arose for the seven transcription factors that undergo nuclear import and export in the model (ELF4, FOS, p50, p52, RelA, RelB, and cRel). Because the model does not represent transcription factor sequestration by chromatin, the nuclear export k_trans_ rate needed to implicitly account for nuclear sequestration as well as true export. Earlier studies of RelA and cRel in LPS stimulated mouse BMDMs found that the maximum rate of nuclear export is significantly slower than the maximum rate of nuclear import [66, 67]. Thus, the initial value of the nuclear export k_trans_ for these seven transcription factors was set to 0.005/s (10% of the nuclear import k_trans_ initial value).

### Experimental Time-Course Data for Model Training

With initial concentrations and rates established, we assembled the dynamic data used to train the model. Simmune model training used 979 protein quantitation ratio values from experiments in LPS-treated mouse and human macrophages, organized into two tables: quantitative LC-MS phosphoproteomics data (Supplemental Table S9) [43, 68–71] and other time-course data (Supplemental Table S10) [45, 46, 67, 72–75]. The key regulatory phosphorylation sites and upstream protein kinases are listed in Supplemental Table S8. LPS concentrations were converted from ng/ml to nM using the approximate average molecular mass of rough LPS, 3.5 kDa [76]. Most protein quantitation values were relative abundance values (e.g., western blot data); the remainder were reported as abundance ratios, such as the ratio of nuclear to total RelA in mouse BMDMs measured by confocal microscopy [67]. For each experiment (time-course or dose response curve), the protein quantitation values were normalized using the maximum value, yielding fold-change values ranging from zero to one; to avoid issues from values equaling exactly zero or one, these were slightly winsorized to a range of 1% to 99%.

### Mice

*Tank*-/- mice were generated by the Akira laboratory [33]. *Ikbke-/-* mice were purchased from The Jackson Laboratory (strain 006908). Wild-type C57BL/6N mice were obtained from Taconic, and mice were maintained under specific-pathogen-free conditions. *Ikbke-/-* and *Tank-/-* mice were each backcrossed onto a C57BL/6 background for at least 10 generations before use, and double-knockout *(Ikbke-/- Tank-/-)* mice were generated by intercrossing the single-knockout strains. Male and female mice 8-12 weeks of age were used for in vitro assays, and mice were aged to 6months for autoimmunity experiments. All procedures were approved by the NIAID Animal Care and Use Committee under protocol LISB-3E (National Institutes of Health, Bethesda, MD).

### Bone marrow–derived macrophage (BMDM) culture

Bone marrow cells were flushed from femurs and tibias using sterile cDMEM and passed through a 70-µm cell strainer. Red blood cells were lysed using ammonium-chloride-potassium (ACK) buffer. Cells were cultured in complete DMEM (DMEM with 4.5 g/L glucose without L-glutamine (Lonza, Walkersville, MD, cat# 12-614F/12), 10% FBS (Gemini Bio-Products, West Sacramento, CA), 2 mM Glutamine (Lonza) and 20 mM HEPES (Lonza)) containing 60 ng/mL recombinant mouse M-CSF (R&D, Cat#: 416-ML-050) for 6 days. Adherent macrophages were harvested on day 7 and used for stimulation assays.

### Stimulation assays

BMDMs were plated at 0.22 x 10^6^ cells/mL in microwell plates and rested overnight before stimulation. Cells were stimulated with 25ng/ml Resiquimod (R848; Invivogen, CAS#:144875-48-9) for ELISA and mRNA harvesting, and 100 ng/mL for western blot; or 100ng/ml Pam3CSK4 (P3C, Invivogen, Cat#: **tlrl-pms**) for the indicated times. Vehicle-treated controls were included in all experiments.

### Quantitative real-time PCR

BMDM were plated at a density of 0.22 x 10^6^ cells/ml in a 24-well plate (BD Falcon) in a total volume of 0.9 mL. 24 hr after plating, the cells were treated with TLR ligands diluted in 0.1 mL of complete DMEM, for time periods as indicated. Cells from 4 wells were pooled for total RNA extraction with RNeasy Mini Kit (Qiagen, Cat. No. 74106), then reverse transcribed with iScript cDNA synthesize kit (Biorad, cat# 1708891). Quantitative PCR was performed using Roche Light Cycler probe master mix (cat# 04707494001) on a Quantstudio 6 Flex instrument. Primer sequences for *Il1a, Il1b, Il6, Il10, Cxcl1,* and *Nfkbiz* were validated for efficiency (90–110%) and are listed in Supplemental Table S11. Transcript levels were normalized to *Hprt1* expression and analyzed using the ΔΔCt method. Each condition was performed in triplicate, and experiments were repeated at least three times independently.

### Cytokine ELISA

BMDMs were plated at 2.2 × 10 cells/mL in 96-well plates (BD Falcon) in a total volume of 90 µL. After 24 h, cells were treated with TLR ligands diluted in 10 µL of complete DMEM for the indicated times. Supernatants were collected after 24 h of stimulation. TNFα, IL-6, and IL-12p40 were measured by ELISA (R&D Systems DuoSet; DY410 for mouse TNFα, DY406 for IL-6, and DY499 for IL-12p40). Cytokine concentrations were determined from vendor-provided protein standard curves.

### Immunoblotting and phospho-protein analysis

Cells were lysed in RIPA buffer (Cell Signaling Technology, Danvers, Mass; catalog no. 9086) containing protease inhibitors (Roche Applied Science, Penzberg, Germany; catalog no. 11836170001), phosphatase inhibitors (Roche Applied Science, catalog no. 4906837001) and 100 mM of N-ethylmaleimide (Sigma, St Louis, Mo; catalog no. E3876). Protein concentrations were normalized by BCA assay. Equal amounts of protein (20–30 µg) were separated on SDS– PAGE gels, transferred to PVDF membranes, and probed with phospho-specific antibodies to p-Erk1/2 (Thr202/Tyr204, Cell Signaling Technology, Danvers, Mass; catalog no. 4370s), p-p38 (Thr180/Tyr182, Cell Signaling Technology, Danvers, Mass; catalog no. 4511s), p-JNK (Thr183/Tyr185, Cell Signaling Technology, Danvers, Mass; catalog no. 4668s), p-IKKα/β (Ser176/180), p-TBK1 (Ser172, Cell Signaling Technology, Danvers, Mass; catalog no. 5483s), and p-IKKε (Ser172). Membranes were stripped and reprobed with total protein antibodies or RhoGDI (Sigma, Cat# R3025) as a loading control. Blots were developed by SuperSignal West Dura Extended Duration Substrate (Thermo-Fisher Scientific, catalog no. 34076), captured with Bio-Rad ChemiDoc and quantified using Bio-Rad Image Lab software.

### Ubiquitination assays (Halo-TUBEs)

To enrich for ubiquitinated proteins, we prepared Halo-TUBEs according to the manufacturer’s instructions [77, 78]. HaloLink Resin and the pFN18A HaloTag T7 Flexi Vector were purchased from Promega. TUBES expression vectors were purchased from Univeristy of Dundee, United Kingdom. BMDMs were stimulated with R848 for 20 minutes, lysed, and incubated with resin overnight. Precipitated proteins were analyzed by immunoblotting with antibodies to TRAF6 (Abcam, Cat#: ab33915), MyD88 (R&D, Cat#: AF3109) or IRAK1 (CST, Cat#: 4504S). Input lysates were run in parallel as loading controls. Relative enrichment of ubiquitylated proteins across genotypes were quantified using Image J.

### Autoimmunity assays

To evaluate longer-term consequences of these pathway defects, mice were maintained for 6 months, at which point spleens and lymph nodes were harvested and fixed in 10% neutral-buffered formalin. Organ size was measured using image analysis software Image J, and area indices were calculated relative to wild-type controls. Serum was collected by retro-orbital bleed, and anti-dsDNA antibody titers were determined by ELISA (Alpha Diagnostic International, Cat#: 5120).).

### Endotoxin-induced shock model

Mice were injected intraperitoneally with lipopolysaccharide from *Salmonella enterica* serotype Minnesota (LPS) (Sigma, Cat#:L6261) at 5 mg/kg body weight. Animals were monitored at 6–12 hour intervals for 10 days and euthanized when humane endpoints were reached. Survival was plotted by Kaplan–Meier method, and differences were assessed by log-rank test.

### Statistical analysis

All experiments were performed at least three times unless otherwise indicated. Data are presented as mean ± SD. Statistical significance was assessed using unpaired two-tailed Student’s *t*-tests for pairwise comparisons or ANOVA with post-hoc Tukey correction for multiple comparisons. For survival analyses, the log-rank test was used. *P* values less than 0.05 were considered significant. Significance levels are indicated as *p* < 0.05 (\*\**), p < 0.01 (**), p < 0.001 (\*\*\**), and p < 0.0001 (****).

## Data Availability

The targeted LC-MS proteomics results are described in our Data Descriptor article [37], and all raw data and analyzed results have been deposited in the ProteomeXchange [79] (ID: PXD031697) and Panorama Public [80] (https://doi.org/10.6069/44s8-9f68) proteomics public data repositories.

The untrained and trained pathway models are provided as text files of the ODEs, as Systems Biology Markup Language (SBML) XML files (four versions) [81], and as Simmune files (Supplemental Dataset 1). The Simmune files also include the model training data and parameters and the pre- and post-training reactant concentrations.

## Results

### Quantitative parameterization of a molecular TLR4 signaling model

To construct a molecular-interaction-level model of TLR4 signaling in mouse bone marrow-derived macrophages (BMDMs), we integrated targeted LC-MS, RNA-seq, published protein-interaction and enzyme-kinetic measurements, phosphoproteomics, flow cytometry, and microscopy with structural modeling, molecular simulation, and rule-based pathway modeling (Figure 1A, B). The targeted LC-MS workflow and dataset are described elsewhere [37]; briefly, LC-PRM measurements ranged from 1,332 copies/cell for Elk1 to 227 million copies/cell for actin, and RNA-seq-based estimates extended parameterization across the BMDM proteome (Figure 1C). As previously reported [37], the median deviation between measured and estimated protein abundance was threefold, giving us a sense of the uncertainty carried into the model by proteins without direct LC-MS measurements.

**Figure 1.**
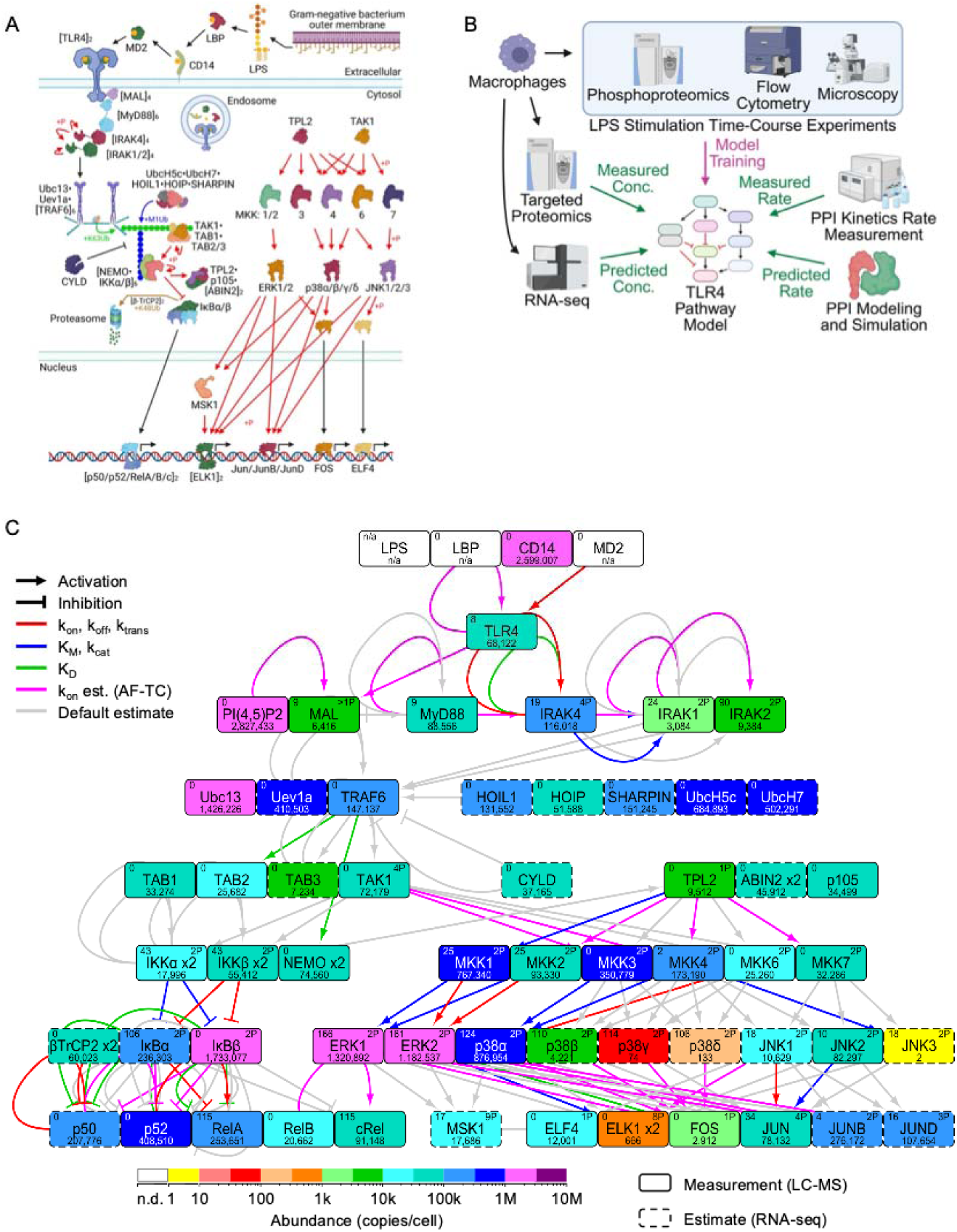
TLR4 Signaling Pathway Model, Workflow and Model Parameterization. The MyD88-dependent TLR4 signaling pathway in mouse BMDMs was modeled. Red, green, blue, and brown arrows depict phosphorylation and K63, M1, and K48 polyubiquitination, respectively (A). Workflow of the experiments and computational analyses (B): mouse BMDMs were analyzed using targeted LC-MS and RNA-seq to produce measurements and estimates, respectively, of protein concentrations for initial model parameterization. Kinetics experiments (from publications from other laboratories, e.g., surface plasmon resonance and enzyme assays) and molecular simulations were used to produce measurements and estimates, respectively, of reaction rate constants for initial model parameterization. Experimental time-course data (*FC Ratios* from publications from our laboratory and other laboratories) were used to train the TLR4 signaling pathway model. The modeled TLR4 signaling pathway (C). Each node indicates the initial protein (homodimer, if indicated) copy number (bottom), number of experimental *FC Ratios* (upper left), and number of phosphorylation sites (upper right). Each edge summarizes the source of the initial reaction rate parameter(s): experimental measurement, estimate from AlphaFold and TransComp simulations, or neither.

The reaction network was based on our mechanistic description of TLR4 signaling [5] (Figure 1A; Supplemental Tables S1 and S2). Since explicit representation of every subunit in large homomeric assemblies was computationally prohibitive, homomers were coarse-grained. The final model contained 445 unknown variables (57 molecular concentrations and 388 reaction-rate constants) and 803 ordinary differential equations. Initial concentrations came from LC-MS measurements or RNA-seq-based estimates. Initial rate values came from published measurements or from 630 AlphaFold analyses and 1,235 TransComp analyses, which yielded 122 non-redundant association-rate predictions. Applied to 27 standards, the same prediction workflow had a median deviation of 4.3-fold (Supplemental Figure S1; Supplemental Tables S3-S6).

Most measured rate constants, and all TransComp simulations, represent buffered *in vitro* conditions that can differ from the intracellular environment due to crowding and other effects [55, 82, 83]. Although some protein-association rates measured in vitro and in living cells differ by only about twofold, with uncertain generality [84], we treated measured and predicted values as informed starting points rather than fixed truths. Better-supported parameters were therefore varied only locally during model training, whereas unmeasured parameters were sampled more broadly (Supplemental Tables S4 and S7; Supplemental Figures S2 and S3).

Model training used 979 normalized protein-abundance ratios from LPS-stimulated mouse and human macrophages (Supplemental Tables S8-S10). Because these measurements, although numerically exceeding the unknown variables, were unevenly distributed and often correlated, their number alone does not establish parameter identifiability or eliminate overfitting. Consistent with this caveat, direct parameter sampling produced poor fits even after millions of simulations. However, genetic-algorithm optimization [85] performed markedly better, reducing discordance below one after 23 iterations and to 22.2% after 85 iterations (Supplemental Figure S4B), over a total of 5,175,240 non-redundant simulations (the top 20 models and corresponding experimental data are provided in Supplemental Dataset S2).

### The model reproduces TLR pathway activation but exposes a proximal deactivation deficit

Having trained the model, we next asked how well it reproduced the known biology of TLR4 activation and deactivation, starting at the receptor. The model represented transfer of LPS through LBP and CD14 to MD2-TLR4 and subsequent receptor dimerization, following the established pathway architecture [5] and LPS biology [86] while acknowledging unresolved features of TLR4 activation [8]. Consistent with this residual uncertainty, simulated cell-surface TLR4 agreed moderately with measurements at 0.03, 30, and 300 nM LPS but was inaccurate at 0.3 and 3 nM (Figure 2B; Supplemental Dataset S2). Given that TLR4 endocytosis can attenuate MyD88 signaling [87, 88] and has been linked to TRIF pathway engagement [88, 89], and because recent evidence distinguishes receptor endocytosis from endosomal signaling [90], this partial agreement points to receptor trafficking as a remaining source of uncertainty.

**Figure 2.**
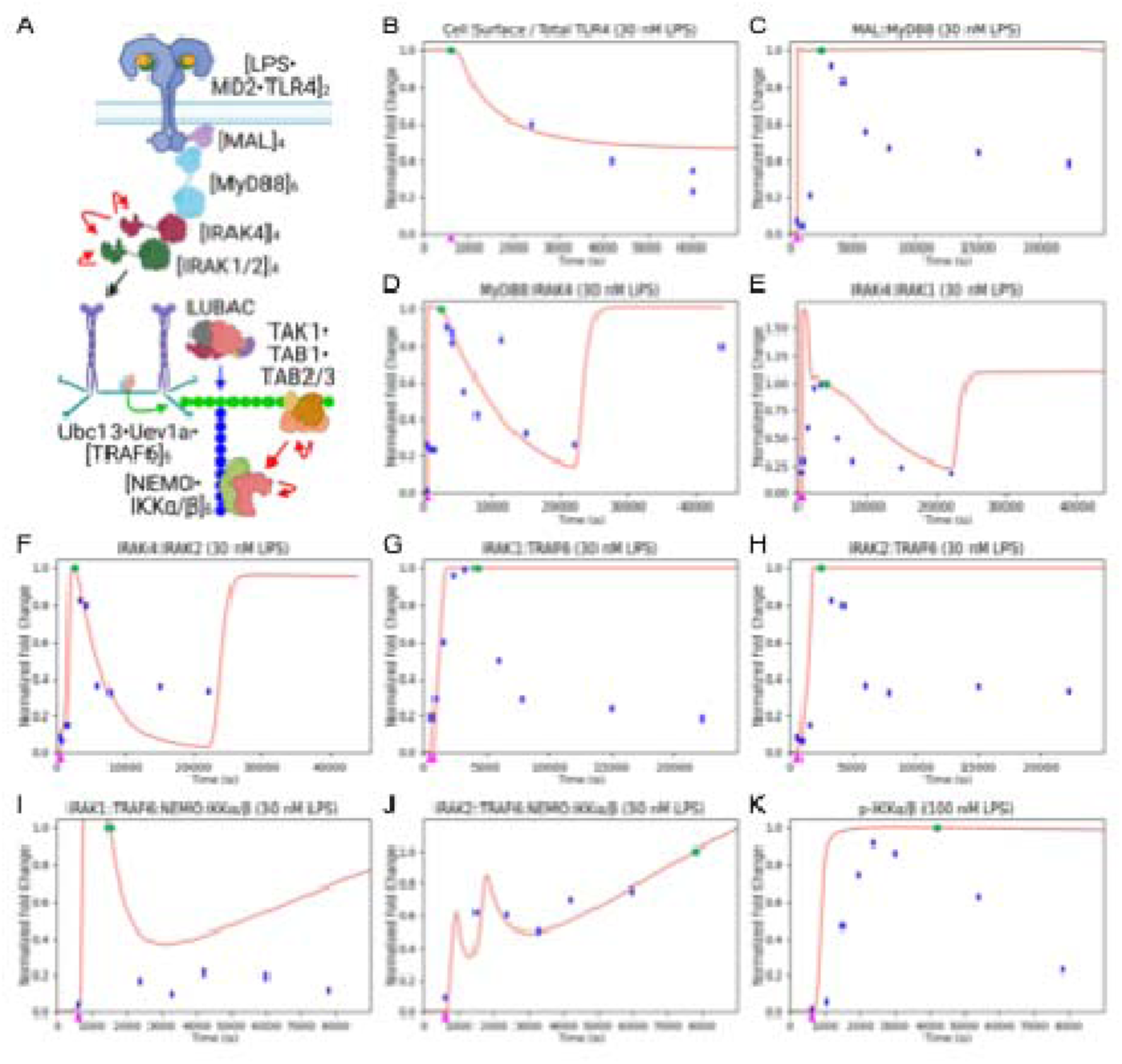
Quantitative Accuracy of Proximal TLR4 Signaling Dynamics. Model simulations (red lines) were compared with experimentally measured normalized abundances (blue points and green *FC Reference Values*). Magenta triangles indicate the addition of LPS at the indicated concentration. Panels depict proximal TLR4 signaling (A) and dynamics of receptor endocytosis (B), myddosome assembly (C-F), TRAF6 signaling (G, H), IKK complex recruitment (I, J), and IKK activation (K).

Moving downstream from the receptor to the early components of the signaling pathway, a clearer and more consistent pattern emerged: the model activated the pathway appropriately but failed to turn it off. Experimentally measured MyD88 activation peaked rapidly and then declined, whereas the model captured activation but not the subsequent decrease [75] (Figure 2C). IRAK4 and the early IRAK1/2 responses were reproduced more accurately, although the model predicted late IRAK reactivation that has not been observed experimentally [75] (Figure 2D-F). In the modeled mechanism, myddosome assembly [91], IRAK phosphorylation and trans-autophosphorylation [33, 92–95], and IRAK dissociation recruit and activate TRAF6 [96]; here, only TRAF6 auto-polyubiquitination was represented among the several known ubiquitination events [96]. As with MyD88, IRAK1- and IRAK2-associated TRAF6 activation was reproduced, but their deactivation was not [75, 97] (Figure 2G, H). Further downstream the model captured formation of IRAK-TRAF6-NEMO-IKK complexes but activated IKKα/β too rapidly and failed to reproduce its decline (Figure 2I-K).

By contrast, performance of the model was considerably stronger when examining later (downstream) aspects of the signaling pathway, suggesting that the deactivation defect was specifically localized to the proximal portion of the pathway rather than reflecting a general limitation of the model. The simulations captured the dose-dependent loss of IκBα/β, although not its partial recovery, and reproduced RelA and cRel nuclear localization with high accuracy (Figure 3; Supplemental Figures S5 and S7). The model also reproduced p38 activation and deactivation, mid-to-high-dose ERK responses, early JNK responses, and Jun-family phosphorylation, while missing MKK1/2 deactivation and exaggerating some low-dose MAPK responses (Figure 4; Supplemental Figure S6). Taken together, a model that reproduced much of the downstream NF-κB and MAPK output nevertheless showed a recurrent proximal defect, with MyD88, TRAF6-associated species, and IKKα/β remaining active longer than observed. Because this pattern persisted after broad parameter exploration, we interpreted it as evidence for missing regulation localized to the MyD88-IRAK-TRAF6/IKK region, rather thanas a uniform failure of the network.

**Figure 3.**
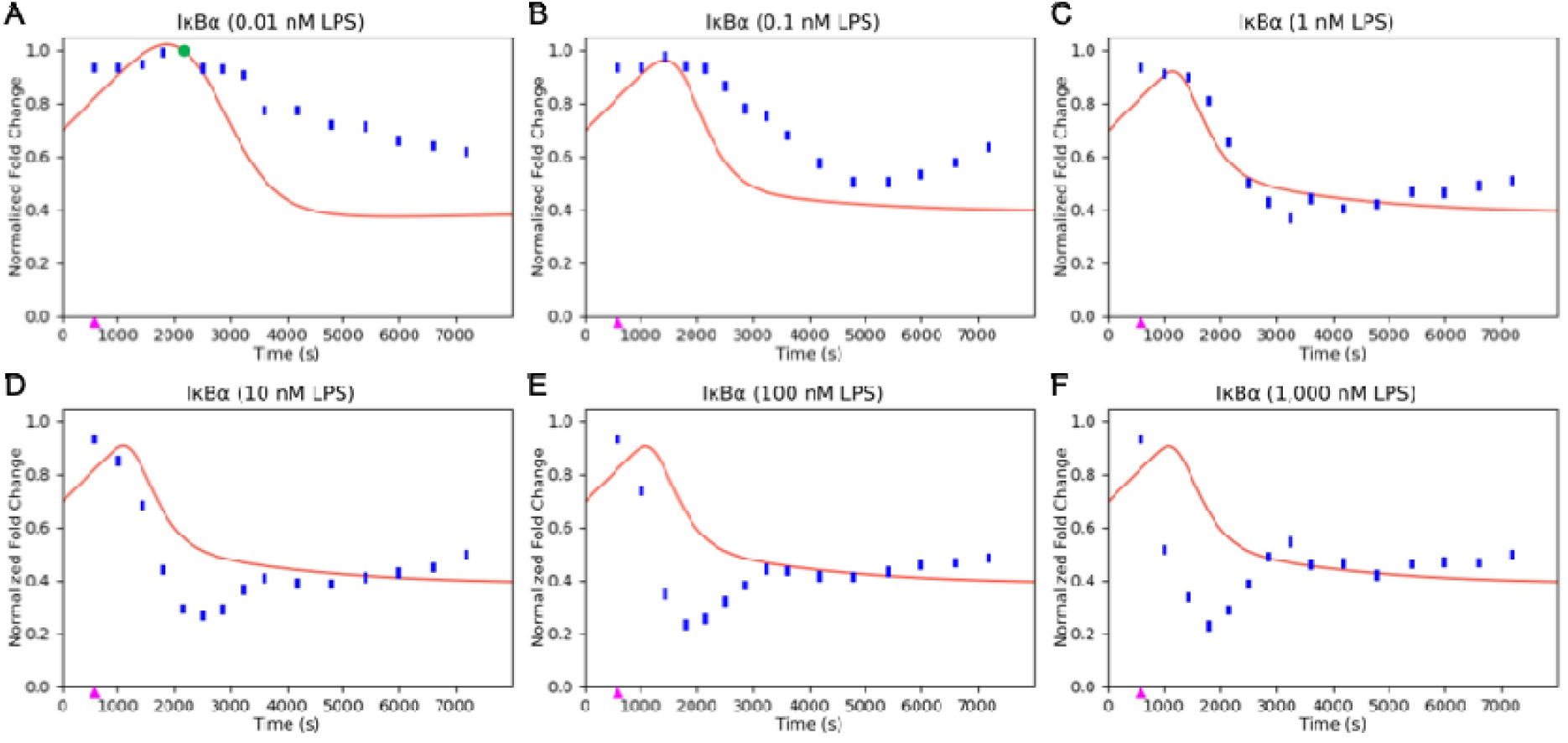
IκBα/β Model Accuracy. Model simulations (red lines) were compared with experimentally measured normalized abundances (blue points and green *FC Reference Values*). Magenta triangles indicate the addition of LPS at the indicated concentration.

**Figure 4.**
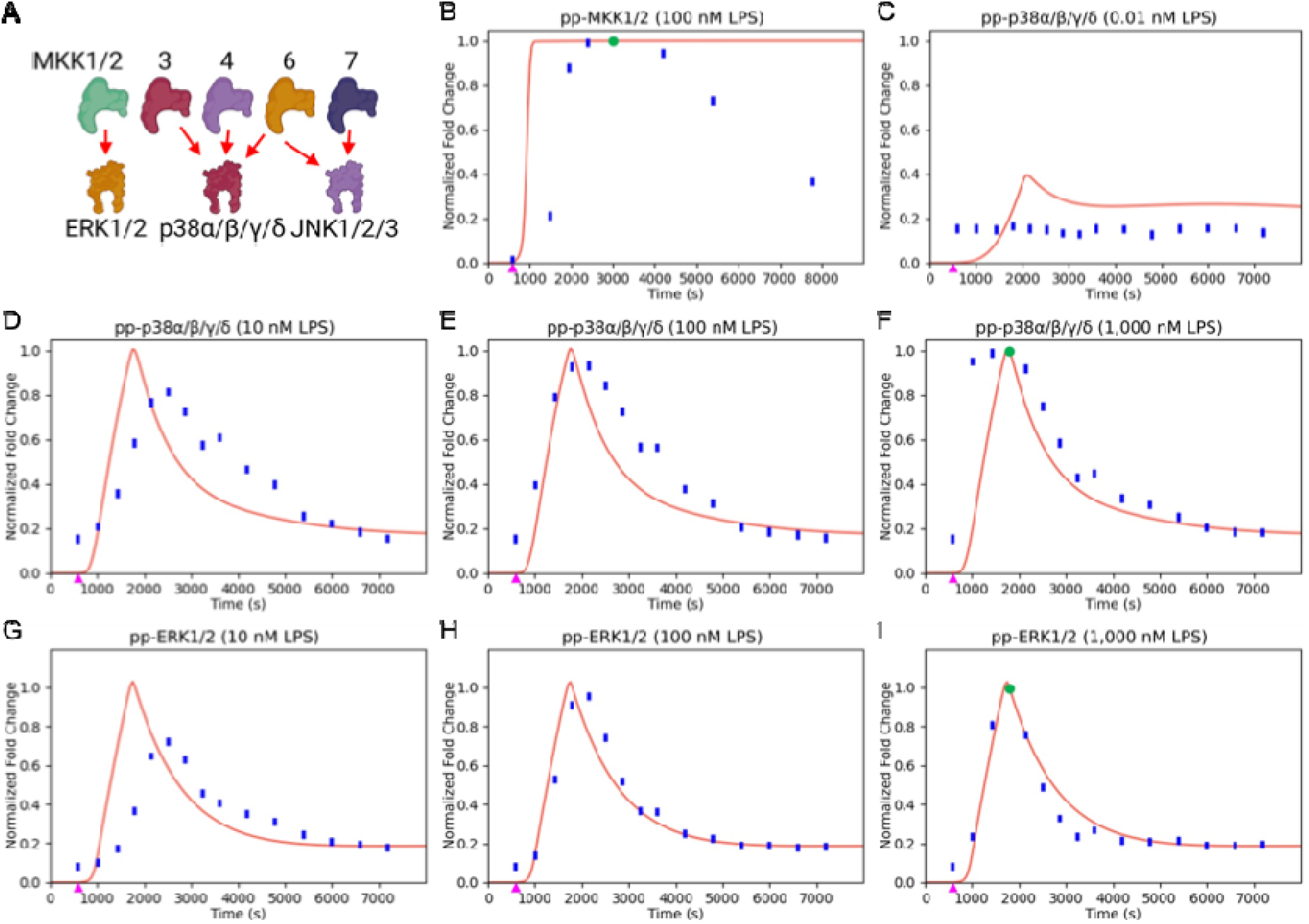
Model Accuracy of MAP Kinase Signaling. Model simulations (red lines) were compared with experimentally measured normalized abundances (blue points and green *FC Reference Values*). Magenta triangles indicate the addition of LPS at the indicated concentration. Panels depict MKK–MAPK signaling (A), pp-MKK1/2 (B), pp-p38α/β/γ/δ (C-F), pp-ERK1/2 (G-I).

### Model failure guides testing of IKK**ε** as a proximal inhibitory regulator

This localized discrepancy gave us a specific, testable prediction that an additional negative regulator should act between the myddosome and TRAF6-dependent activation. IKKε has been associated primarily with TRIF-dependent TLR signaling [98], is recruited by the TRAF-associated scaffold TANK [36], and TANK is an established negative regulator of TLR responses [33], making this pathway a natural candidate, even though the model did not uniquely predict IKKε or TANK as the involved molecules but rather identified the proximal region in which an additional negative regulator module was required.

To test whether IKKε had the postulated negative regulatory role and to do this test in a system with MyD88-dependent signaling separated from TLR4-specific TRIF contributions, we began with stimulated wild-type and *Ikbke*-/- BMDMs with R848 (TLR7) or Pam3CSK4 (TLR1/2), rather than with LPS. *Ikbke*-/- macrophages showed greater induction of *Il1a*, *Il1b*, *Il6*, *Il10*, *Cxcl1*, and *Nfkbiz*, together with increased TNFα, IL-6, and IL-12p40 secretion (Figure 5A, B). These data genetically assign IKKε an inhibitory role in MyD88-dependent macrophage responses.

**Figure 5.**
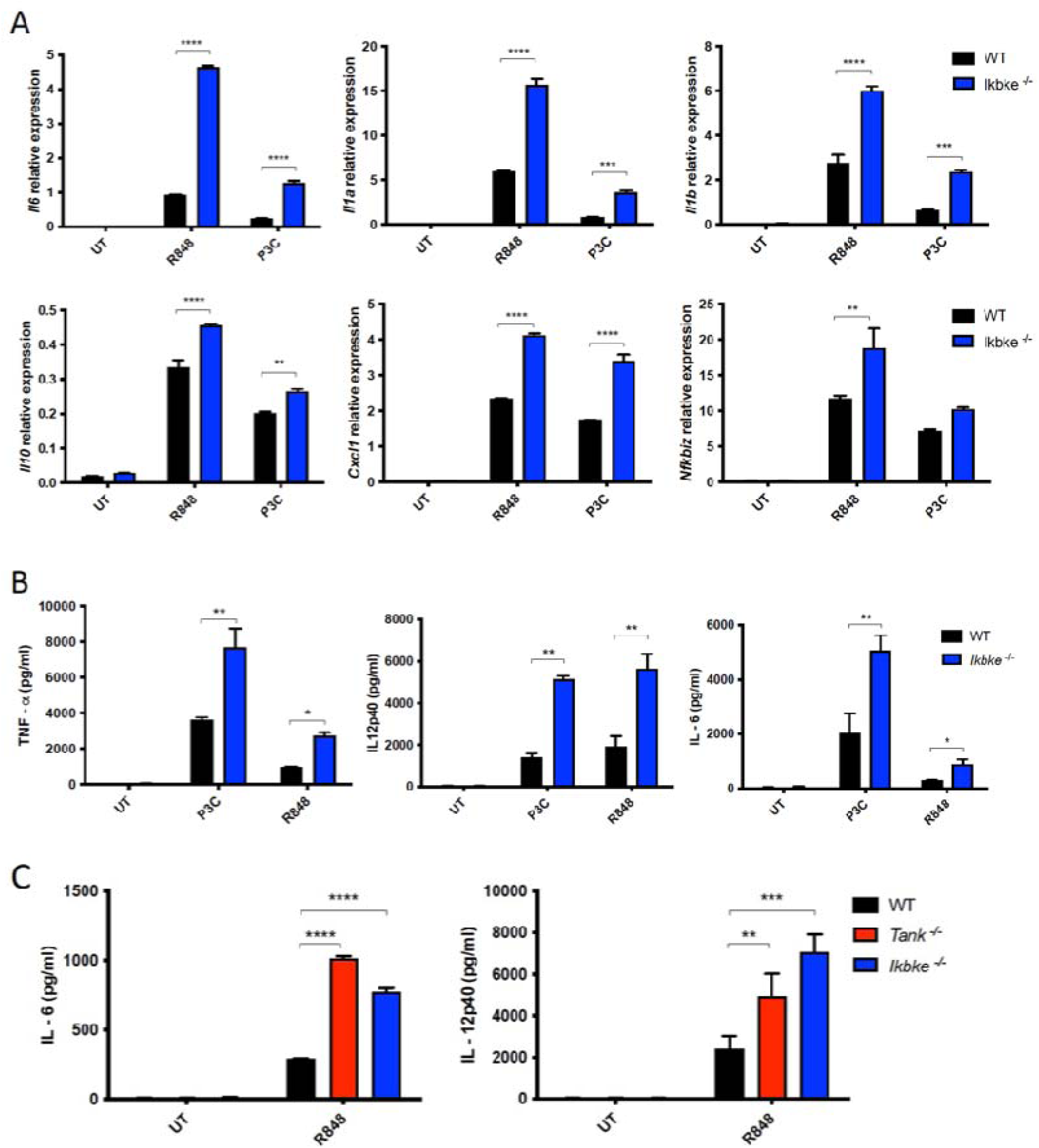
IKKε deficiency enhances MyD88-dependent macrophage responses. (A) Gene-expression analysis of BMDMs from wild-type (WT) and *Ikbke*-/- mice stimulated with 100 ng/mL Pam3CSK4 (P3C; TLR1/2 agonist) or 25 ng/mL R848 (TLR7 agonist) for 24 h. Quantitative RT-PCR was performed for *Il1a*, *Il1b*, *Il6*, *Il10*, *Cxcl1*, and *Nfkbiz*. Expression was normalized to unstimulated controls; *Ikbke*-/- macrophages showed greater induction than WT cells. (B) Cytokine secretion from WT and *Ikbke*-/- BMDMs after 24 h of P3C or R848 stimulation. Supernatants were analyzed by ELISA for TNFα, IL-6, and IL-12p40. Cytokine release was increased in *Ikbke*-/- macrophages across both stimuli. Data are representative of at least three independent experiments. Statistical significance was determined using Student’s t-test; p < 0.05 (*), p < 0.01 (**), p < 0.001 (***), and p < 0.0001 (****). (C) Cytokine secretion by WT, *Ikbke*-/-, and *Tank*-/- macrophages stimulated with 25 ng/mL R848. IL-6 and IL-12p40 secretion was increased in both knockout strains. Data are representative of three independent experiments; statistical testing was as in panel B.

### TANK is required for stimulus-induced IKK**ε** phosphorylation and shares its inhibitory phenotype

Having established an inhibitory role for IKKε, we next asked whether its scaffold TANK was required for this function. Following R848 stimulation, *Ikbke*-/- BMDMs showed enhanced phosphorylation of ERK, p38, JNK, IKKα/β, and TBK1 (Supplemental Figure S8A). *Tank*-/- BMDMs displayed a similar pattern of MAPK and canonical IKK hyperactivation. Critically, R848-induced IKKε phosphorylation was abolished in *Tank*-/- cells, whereas TBK1 phosphorylation was retained (Supplemental Figure S8B), demonstrating that TANK is required for stimulus-induced IKKε phosphorylation under these conditions. Consistent with this shared requirement, R848-induced IL-6 and IL-12p40 secretion was comparably increased in *Ikbke*-/- and *Tank*-/- macrophages (Figure 5C). The convergence of signaling and cytokine phenotypes, together with TANK-dependent IKKε phosphorylation, is consistent with a shared inhibitory axis, although it does not imply that the two proteins have identical functions.

### IKK**ε** and TANK constrain IRAK1 and TRAF6 ubiquitination

As IKKε _and TANK act as a shared inhibitory axis, we next sought to position this checkpoint more precisely within the proximal signaling module identified by the model. To do so, we enriched ubiquitinated proteins after 20 min of R848 stimulation. TRAF6 ubiquitination, a marker of TRAF6 activation [92, 99], was increased in both *Ikbke*-/- and *Tank*-/- BMDMs (Figure 6A, B). MyD88 ubiquitination was weaker and was not increased in either knockout, consistent with the requirement for unmodified MyD88 in efficient myddosome assembly [91] (Figure 6C, D). IRAK1 ubiquitination, however, was significantly increased in both knockouts (Figure 6E, F). These results place the TANK-IKKε checkpoint at or immediately downstream of the IRAK1-TRAF6 ubiquitin-signaling node, although they do not identify the direct IKKε substrate.

**Figure 6.**
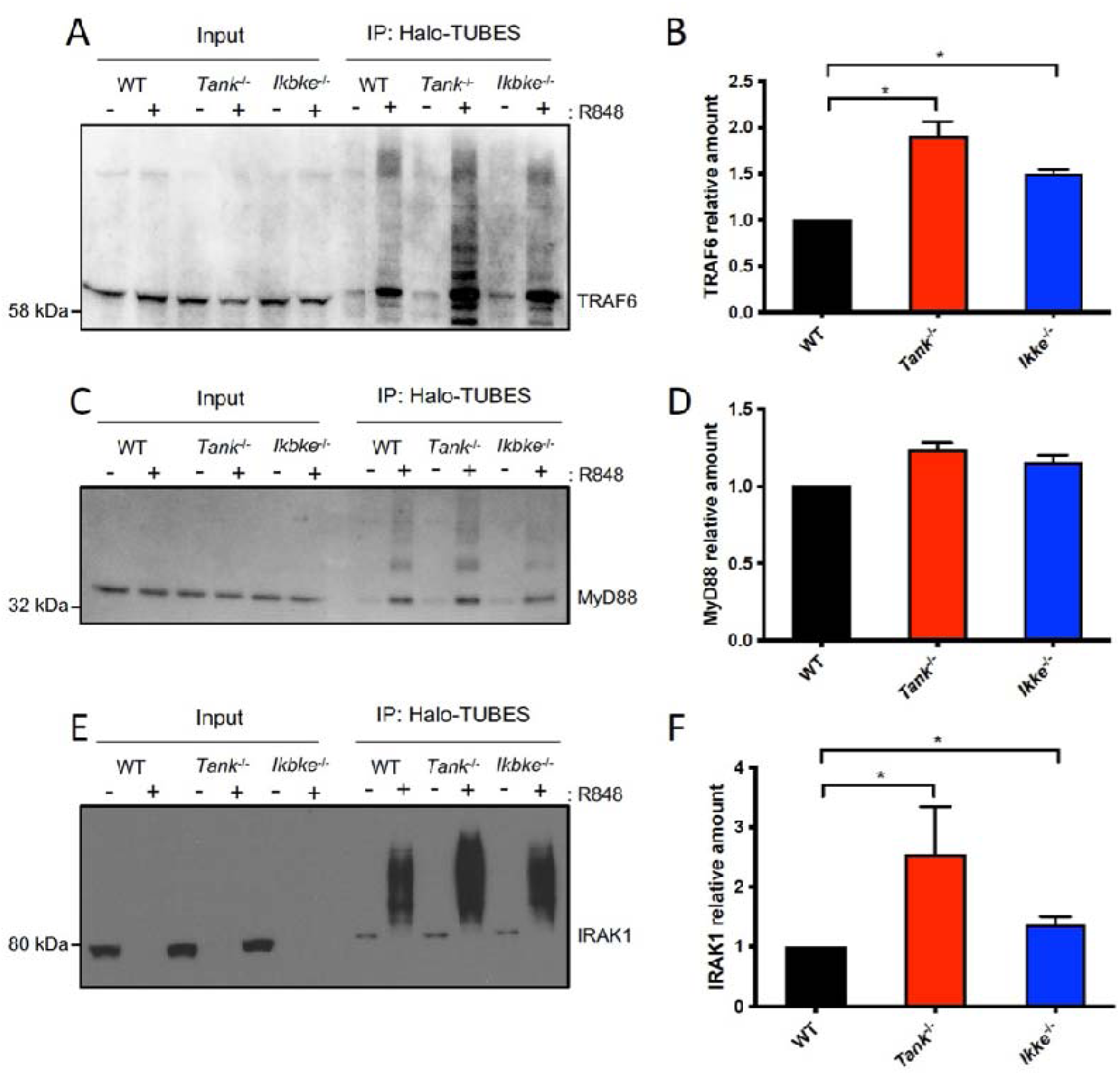
IKKε and TANK constrain IRAK1 and TRAF6 ubiquitination. (A, B) TRAF6 ubiquitination was measured in WT, *Ikbke*-/-, and *Tank*-/- BMDMs using Halo- TUBES precipitation after 100 ng/mL R848 stimulation. Both knockout strains showed increased TRAF6 ubiquitination relative to WT. (C, D) MyD88 ubiquitination was detectable after R848 stimulation but was not increased in *Ikbke*-/- or *Tank*-/- cells, indicating that the inhibitory effect is positioned downstream of this step. (E, F) IRAK1 ubiquitination was induced by R848 in WT cells and further increased in both *Ikbke*-/- and *Tank*-/- macrophages. Together with the TRAF6 results, these data place the TANK- IKKε checkpoint at or immediately downstream of the IRAK1-TRAF6 node without identifying a direct substrate. Immunoblots are representative of three independent experiments; densitometry is shown as mean ± SD. Statistical significance was determined using Student’s t- test; p < 0.05 (*).

### IKK**ε** and TANK have overlapping but non-identical functions *in vivo*

Finally, we asked whether these biochemical phenotypes translate into overlapping physiological consequences *in vivo.* At six months of age, both *Ikbke*-/- and *Tank*-/- mice developed splenomegaly and lymphadenopathy, with a stronger phenotype in *Tank*-/- animals (Figure 7A- D). *Ikbke*-/- *Tank*-/- double-knockout mice did not show additive organ enlargement, consistent with substantial pathway overlap. However, the two genotypes diverged in other respects: anti-dsDNA antibodies were significantly increased in *Tank*-/- but not *Ikbke*-/- mice; the increase in double-knockout mice did not reach statistical significance (Figure 7E), suggesting that TANK has additional functions in autoantibody control. Similarly, after intraperitoneal LPS challenge (5 mg/kg), *Ikbke*-/- mice showed significantly reduced survival, whereas *Tank*-/- mice had only a modest increase in susceptibility (Figure 7F). Together, the organ, serological, and endotoxin- challenge phenotypes support overlapping roles for TANK and IKKε in inflammatory restraint while also revealing protein-specific functions.

**Figure 7.**
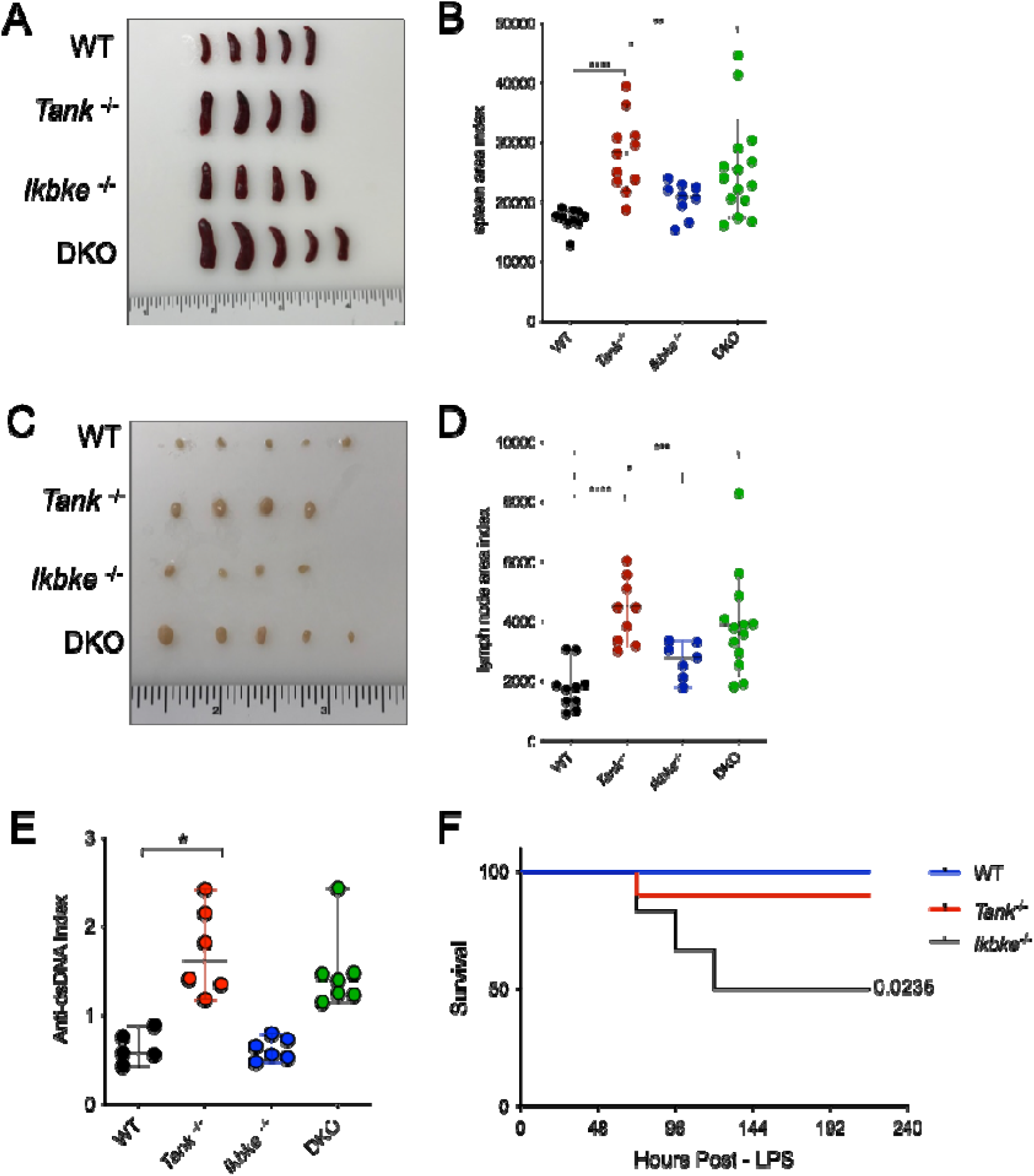
IKKε- and TANK-deficient mice display overlapping and distinct inflammatory phenotypes. (A-D) Six-month-old WT, *Ikbke*-/-, *Tank*-/-, and *Ikbke*-/- *Tank*-/- double-knockout (DKO) mice were assessed for lymphoid-organ enlargement. Spleen and lymph-node areas were increased in both single-knockout strains, with no additive enlargement in DKO mice. (E) Serum anti-dsDNA antibodies were measured by ELISA. Titers were significantly increased in *Tank*-/- mice but not in *Ikbke*-/- mice; the increase in DKO animals did not reach statistical significance. (F) Survival after intraperitoneal LPS injection (5 mg/kg). *Ikbke*-/- mice showed significantly reduced survival, whereas *Tank*-/- mice showed a milder increase in susceptibility. Data are representative of at least three independent experiments. Statistical significance was determined using Student’s t-test for panels B, D, and E; p < 0.05 (*), p < 0.01 (**), p < 0.001 (***), and p < 0.0001 (****), or by Kaplan-Meier analysis with a log-rank test for panel F.

## Discussion

Here we report a an enhanced, highly parametrized model for TLR signaling and show the value of building such a detailed model in for biological discovery. Our detailed model reproduced much of what is known and we could measure with respect to TLR4 activation and downstream signaling but repeatedly failed despite broad parameter testing to reproduce signal termination. The data localized the missing biology to the proximal MyD88/IRAK/TRAF6/IKK region of the pathway. Targeted experiments then identified a TANK-dependent IKKε checkpoint that restrains IRAK1 and TRAF6 ubiquitination. Since the model did not uniquely nominate IKKε or TANK, it is most correct to state that modeling guided the identification of these missing factors by defining where additional regulation was required. This use of informative model failure illustrates the value of mechanistic approaches for resolving incompletely characterized signaling networks [1, 2, 13].

The recurrent deactivation failures are especially biologically informative precisely because TLR activation is much better characterized than its termination [25], and the dominant negative regulatory mechanisms remain unresolved [24]. LPS can induce rapid dephosphorylation through incompletely understood processes [26]; we recently observed hundreds of phosphosites dephosphorylated within 5 min [69], too rapidly to be explained solely by newly expressed dual-specificity phosphatases [27]. Pre-existing phosphatases, deubiquitinases, trafficking pathways, and inhibitory scaffolds are therefore plausible missing components. Independently, RNAi screening identified additional regulators, including Arf6, CYLD, and A20 [103]. Alternative receptor-proximal mechanisms, such as transient MAL dependence followed by direct TLR4– MyD88 signaling [20, 64] or IRAK3-mediated inhibition [104], could also contribute to the observed mismatch. These possibilities reinforce our interpretation that TANK–IKKε represents one component of a redundant deactivation system rather than the unique mechanism responsible for inflammatory shutdown [24].

Studies in other biological systems provide useful context for this strategy of gaining biological insight from model failure. Aldridge et al. used a fuzzy-logic model of TNF-, EGF-, and insulin-induced signaling to infer previously unrecognized pathway crosstalk, including unexpected inhibition of IKK after EGF treatment that was proposed to arise through receptor downregulation and disruption of an autocrine signaling circuit [100]. Their work demonstrated that model structure and model–data discrepancies can reveal inhibitory relationships that are not apparent from inspection of the data alone. In a different context, von Dassow et al. showed that a model of the Drosophila segment-polarity network generated the appropriate developmental pattern across broad ranges of kinetic parameters, establishing that robust behavior can arise from network architecture without precise tuning of every reaction constant [101]. Together, these studies illustrate the significance of our findings: Aldridge et al. illustrate how modeling can expose hidden inhibitory regulation, whereas von Dassow et al. illustrate how a sufficiently complete network can maintain function despite kinetic uncertainty. In our model, the persistence of deactivation errors during broad parameter exploration argued that the modeled TLR4 pathway description lacked one or more regulatory reactions, rather than merely containing inaccurate individual rate constants.

It is useful to contrast the present model with those reported earlier [18–21]. An and Faeder previously developed a BioNetGen model of TLR4 signaling and preconditioning that represented pathway interactions and A20- and IκB-mediated negative feedback in rule-based form [18], reproducing dose-dependent TNF production and endotoxin tolerance and showing that the two inhibitory mechanisms made distinct contributions to attenuation of the initial response and maintenance of the tolerant state. That model, however, was intentionally designed as a detailed qualitative knowledge representation, with protein abundances and reaction coefficients assigned approximate or default values rather than constrained by molecular measurements. Our work extends this earlier rule-based approach through measured or RNA-seq-estimated protein concentrations, literature- and structure-informed reaction-rate estimates, extensive dynamic training data, and systematic use of persistent model–data discordance to localize missing deactivation biology. Li et al. focused on proximal TLR4–MyD88 complex assembly [20], Liu et al. addressed TLR3–TLR7 crosstalk and innate immune memory [21], and Guo et al. emphasized multi-receptor integration, single-cell heterogeneity, and information transmission [19]. The present model instead emphasizes biochemical and molecular-interaction detail across the receptor-to-transcription-factor pathway. Its measured protein copy numbers, literature- and structure-informed rate estimates, broad dynamic training set, and millions of simulations place it among the most extensively data-constrained molecular models of macrophage TLR signaling.

This parameterization strategy also defines important limitations. Our earlier macrophage S1P model used targeted proteomics and microscopy to constrain a pathway operating on a timescale of seconds [16], whereas TLR4 responses unfold over minutes and hours [45, 46], which permits a correspondingly wider range of plausible parameters here. *In vivo* approaches such as fluorescence cross-correlation spectroscopy may improve direct measurement of reaction rates [102], but structural modeling and molecular simulation will remain useful wherever such measurements are unavailable. Moreover, the 979 experimental ratios do not represent 979 independent constraints: time points, doses, and pathway readouts are correlated and unevenly distributed; the resulting model should therefore be viewed as a constrained foundation for hypothesis generation rather than evidence that every fitted parameter is uniquely identifiable. Network redundancy and population-level averaging further limit inference of unmeasured intermediates [22]. Model structure may contribute to the aditional discrepancies between the model output and experimentally measured values. Cooperative higher-order assembly can generate switch-like signaling behavior [6, 7], whereas the present model coarse-grained large homomeric complexes to maintain computational tractability. Given that myddosome number and size influence downstream signaling in individual cells [73, 105], population averages can convert digital signalosome-assembly events into apparently graded responses. More explicit representations of signalosome assembly, spatial organization, and single-cell heterogeneity may therefore help reduce the predicted late IRAK reactivation and improve the accuracy of proximal signaling dynamics in future iterations of the model.

The biological advance achieved by identification of TANK and IKKε as central to the model’s failure to show appropriate deactivation is relevant in this context. TANK is known to suppress TLR signaling [33], IKKε is established in antiviral and interferon responses [34] and in IL-17 signaling [35], and combined inhibition of IKKε and TBK1 can enhance IL-1-driven signaling [36]. Building on this body of literature on TLR and innate immune pathway organization [4, 5], our data add genetic evidence that IKKε alone restrains MyD88-specific TLR responses, demonstrate that TANK is required for stimulus-induced IKKε phosphorylation, and localize their shared inhibitory effect to IRAK1 and TRAF6 ubiquitination rather than MyD88 ubiquitination. The resulting increases in MAPK and NF-κB activation, cytokine production, lymphoid-organ enlargement, and sensitivity to endotoxin establish the physiological relevance of this regulatory axis. At the same time, several observations prevent TANK and IKKε from being treated as functionally identical. Their similar cellular phenotypes and the absence of additive lymphoid-organ enlargement in double-knockout mice support substantial pathway overlap, but their autoantibody and endotoxin-shock phenotypes differ, and TANK also participates in TRAF-dependent pathways beyond IRAK signaling [20, 33]. Regulation at TRAF6, with IRAK1 hyperubiquitination as a secondary effect, may therefore be more plausible, and the differential knockout phenotypes suggest that TANK has partners beyond IKKε.

Considering that dysregulated TLR signaling contributes to autoimmunity and hyperinflammatory states such as sepsis, the IKKε–TANK axis may ultimately offer a therapeutic entry point. Small-molecule IKKε inhibitors have been explored in cancer and viral infection, but interventions must preserve protective immunity, and the effects of IKKε modulation may differ by context. Overall, our results define a regulatory pathway in which IKKε and TANK limit MyD88-dependent signaling at the IRAK1–TRAF6 ubiquitination node. Future model refinement should incorporate this axis, rapid phosphatase activation, and signalosome dynamics to improve prediction and to guide experiments and therapies for inflammatory disease.

## Acknowledgements

The authors are grateful for the assistance provided by Clinton J. Bradfield, Juan Brandi, Danielle A. Castro, Casey M. Daniels, Michael G. Dorrington, Orna R. Ernst, Joseph G. Gillen, David R. Goodlett, Thomas A. Johnson, Mohd M. Khan, David Le, Bernadette Marrero, Andrew J. Martins, Arthur G. Nuccio, Matthew W. Scandura, Jian Song, James Wilson, and Sung Hwan Yoon.

This research was supported by the Intramural Research Program of the National Institutes of Health (NIH). The contributions of the NIH authors are considered Works of the United States Government. The findings and conclusions presented in this paper are those of the authors and do not necessarily reflect the views of the NIH or the U.S. Department of Health and Human Services. This study utilized high-performance computing (HPC) resources from the Office of Cyber Infrastructure and Computational Biology (OCICB) at the National Institute of Allergy and Infectious Diseases (NIAID) and the Biowulf Linux cluster at the National Institutes of

Health, Bethesda, MD (http://biowulf.nih.gov). This project has been funded in part with Federal funds from the National Institute of Allergy and Infectious Diseases (NIAID), National Institutes of Health, Department of Health and Human Services under BCBB Support Services Contract HHSN316201300006W/75N93022F00001 to MEDICAL SCIENCE & COMPUTING.

## Author Contributions

Conceptualization: N.P.M., I.D.C.F., R.N.G., M.M.S., A.N.L. Investigation: N.P.M., J.M.C., P.R.K., R.A.G., B. L., J. S. Formal analysis: N.P.M., F.Z., S.A.H., A.A.A., Y.S., J.M.C., P.R.K., R.A.G., M.J.M., D.K., M.M.S. Writing - Original Draft: N.P.M. Writing - Review & Editing: all authors. Project administration: A.N.L., I.D.C.F., M.M.S. N.P.M., J.M.C., P.R.K., and D.K. performed LC-MS proteomics experiments and data analysis. N.P.M., S.A.H., A.A.A., and M.J.M. performed protein complex structure modeling and association rate constant prediction. N.P.M., F.Z., Y.S., and M.M.S. performed TLR4 pathway modeling. B. L., J. S., and R.A.G. performed PCRs, ELISAs, pull-downs, immunoblotting, autoimmunity assays, and endotoxic shock assays (experiments and data analysis).

**Supplemental Figure S1. Accuracy of the Predicted k_on_ Rates**

AlphaFold and TransComp were used to predict TLR4 signaling pathway PPI k_on_ rate values. To estimate the accuracy of these predictions, 27 k_on_ standards were also analyzed. These diverse PPIs are unrelated to TLR4 signaling, and their k_on_ values were previously measured *in vitro* and predicted using AlphaFold and TransComp (Supplemental Table S6). The mean, geometric mean, and median fold-deviation were 19.5, 7.3, and 4.3, respectively (i.e., half of the predictions were accurate to within give-or-take 4.3-fold). Linear regression (blue line) and 45° line (green line).

**Supplemental Figure S2. eCDFs and Log-Normal CDFs used for Model Training**

The experimentally measured reaction rate constant values (Supplemental Table S4) were summarized and used to produce eCDF curves (red lines). Based on each eCDF, a default initial value (green line) and a Log-Normal distribution (black line) were produced.

**Supplemental Figure S3. Log-Normal Distribution CDFs used for Model Training**

All the distributions used for model training are depicted. The initial parameter values with the highest estimated accuracy were multiplied by a factor randomly sampled from the narrowest distribution (Q3 = 2.5). Likewise, the initial parameter values with the lowest estimated accuracy were multiplied by a factor randomly sampled from the widest distribution (Q3 = 15). The criteria are listed in Supplemental Table S7.

**Supplemental Figure S4. A. The Simmune Reaction Network.** Each node is a *Complex Species*, and each edge is a reaction. LPS is at the top level, and each level downward is an additional degree of separation from LPS. Visualized in Simmune Network Viewer. **B. Model Training using the GA.** Initially, 200,000 *Analyses* (2,800,000 simulations) were performed (GA Iteration 0). Subsequently, 85 iterations of the GA were performed (170,000 *Analyses*, 2,375,240 non-redundant simulations). For each GA iteration, the minimum *Discordance* is charted (i.e., the value from the model with the smallest *Discordance*). The best model had *Discordance* = 22.2%.

**Supplemental Figure S5. Model** Accuracy of RelA and cRel Nuclear Localization

Model simulations (red lines) were compared with experimentally measured normalized abundances (blue points and green *FC Reference Values*). Magenta triangles indicate the addition of LPS at the indicated concentration.

**Supplemental Figure S6. Model Accuracy of JNK and Jun Signaling**

Model simulations (red lines) were compared with experimentally measured normalized abundances (blue points and green *FC Reference Values*). Magenta triangles indicate the addition of LPS at the indicated concentration. Panels pp-JNK1/3 (A-B), pp-JNK2 (C), p-Jun (D), p-JunB (E), p-Jun/p-JunD (F).

**Supplemental Figure S7. Precision of RelA Nuclear Localization Modeling**

Simulation results were produced using the top 20 ranked models. Magenta triangles indicate the addition of 30 nM LPS. Concentration (A), nuclear / total concentration ratio (B), and nuclear / total concentration ratio normalized using the maximum value (C).

**Supplemental Figure S8. TANK and IKKε deficiencies produce similar hyperactivation of MAPK and NF-κB signaling.**

(A) Immunoblot analysis of WT and *Ikbke-/-* BMDMs stimulated with 100 ng/ml R848 for the indicated times. Phosphorylation of Erk, p38, JNK, IKKα/β, and TBK1 was elevated in *Ikbke-/-* macrophages compared with WT. As expected, IKKε phosphorylation was absent in knockout cells. RhoGDI served as a loading control.

(B) Immunoblot analysis of WT and *Tank*-/- BMDMs stimulated with 100 ng/mL R848. *Tank*-/- macrophages displayed enhanced phosphorylation of MAPK and NF-κB signaling proteins, and IKKε phosphorylation was absent, consistent with TANK-dependent IKKε phosphorylation.

