## Supplemental Datasets for "Quantitative Modeling of TLR Signaling Reveals Missing Negative Feedback Guiding Identification of TANK-IKKε Checkpoint": DatasetS2.pdf

### Data-Driven Modeling of the Mouse Macrophage Toll-like Receptor Signaling Pathway

Manes et al, 2026

Supplemental Material

Figure S6. All of the experimental data (normalized fold change ratios) used to train the model are plotted (blue bars; height = 0.02). The corresponding simulation results from the top 20 ranked models are also plotted (red lines). The vertical magenta line indicates the time at which LPS was added (t = 600 s).

Experimental Measurement (*FC Ratio*)

Simulation Results (top 20 ranked models)

LPS Added (t = 600 s)

### TLR4

### Experiment vs. Simulation: 30 nM LPS, Cell Surface / Total TLR4

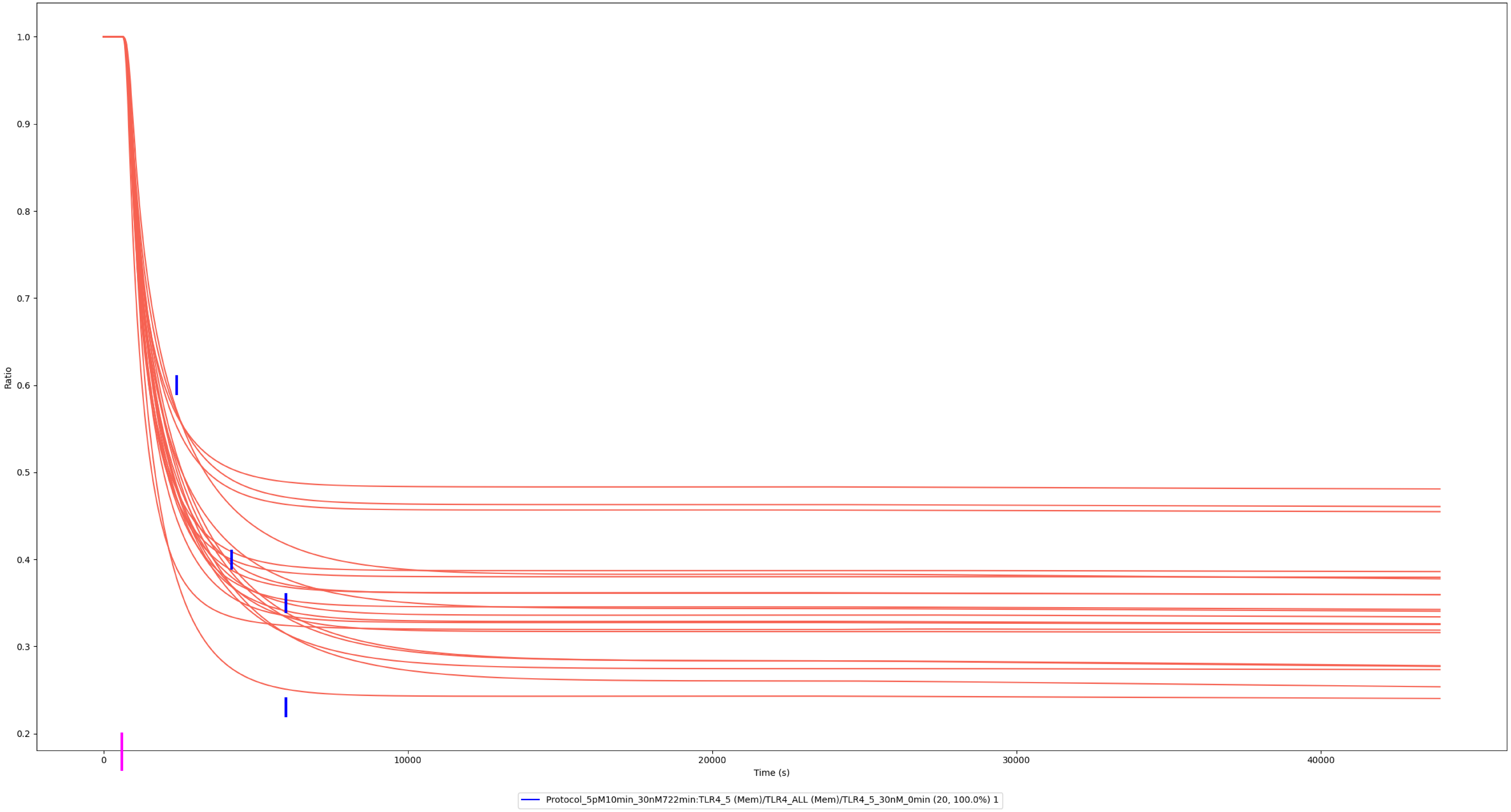

### Experiment vs. Simulation: 0.03 nM LPS, Cell Surface / Total TLR4

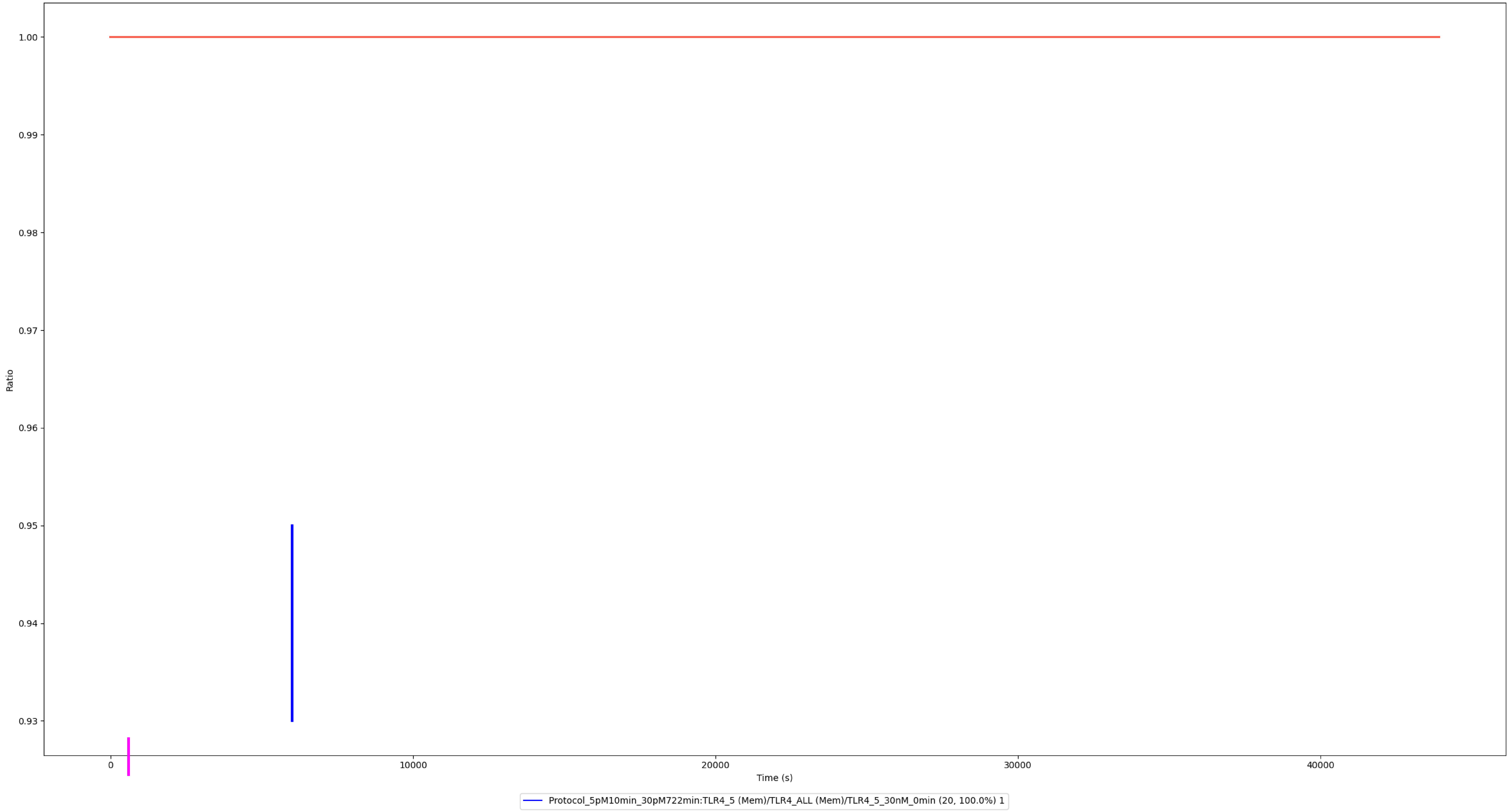

### Experiment vs. Simulation: 0.3 nM LPS, Cell Surface / Total TLR4

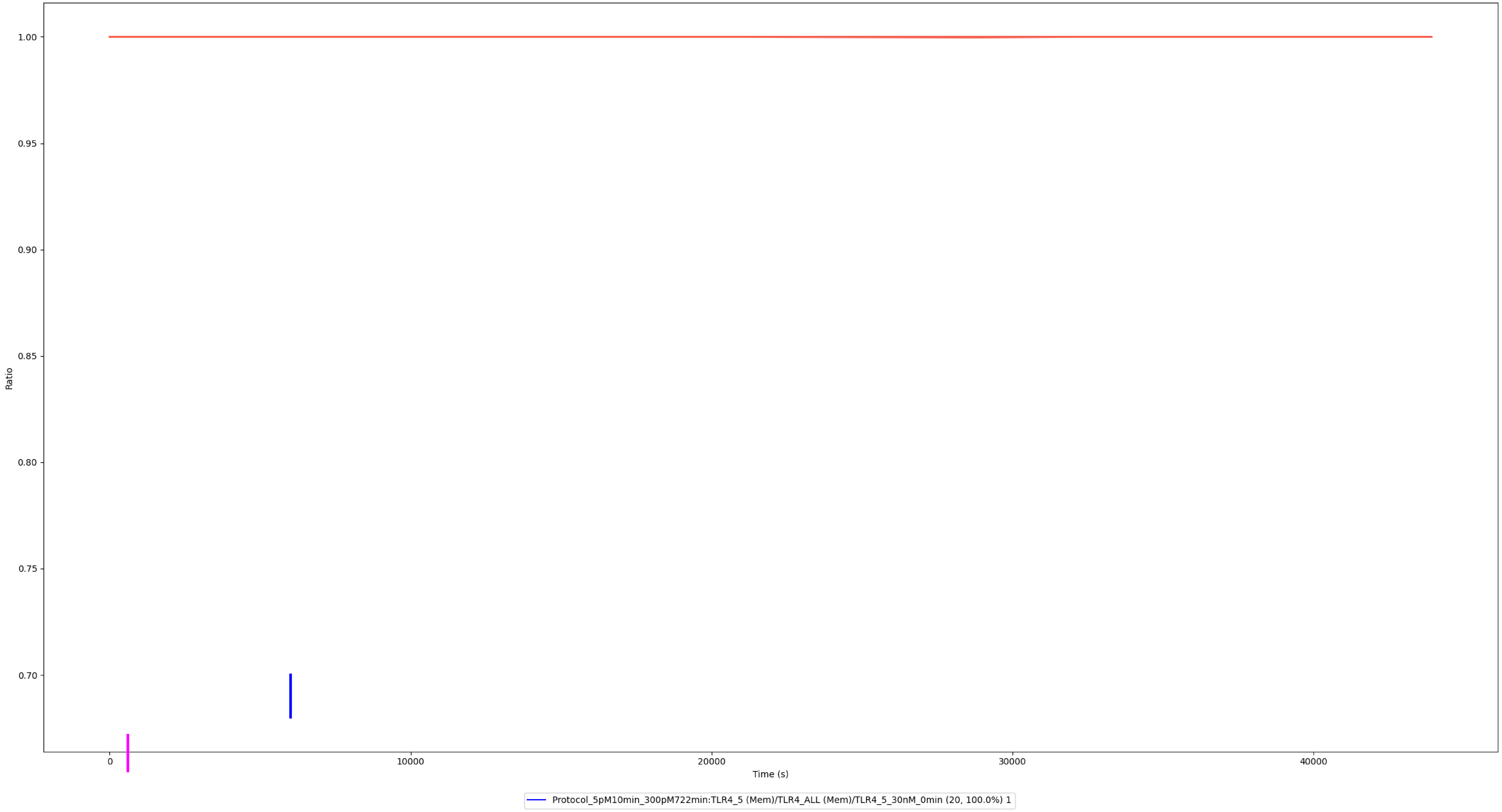

### Experiment vs. Simulation: 3 nM LPS, Cell Surface / Total TLR4

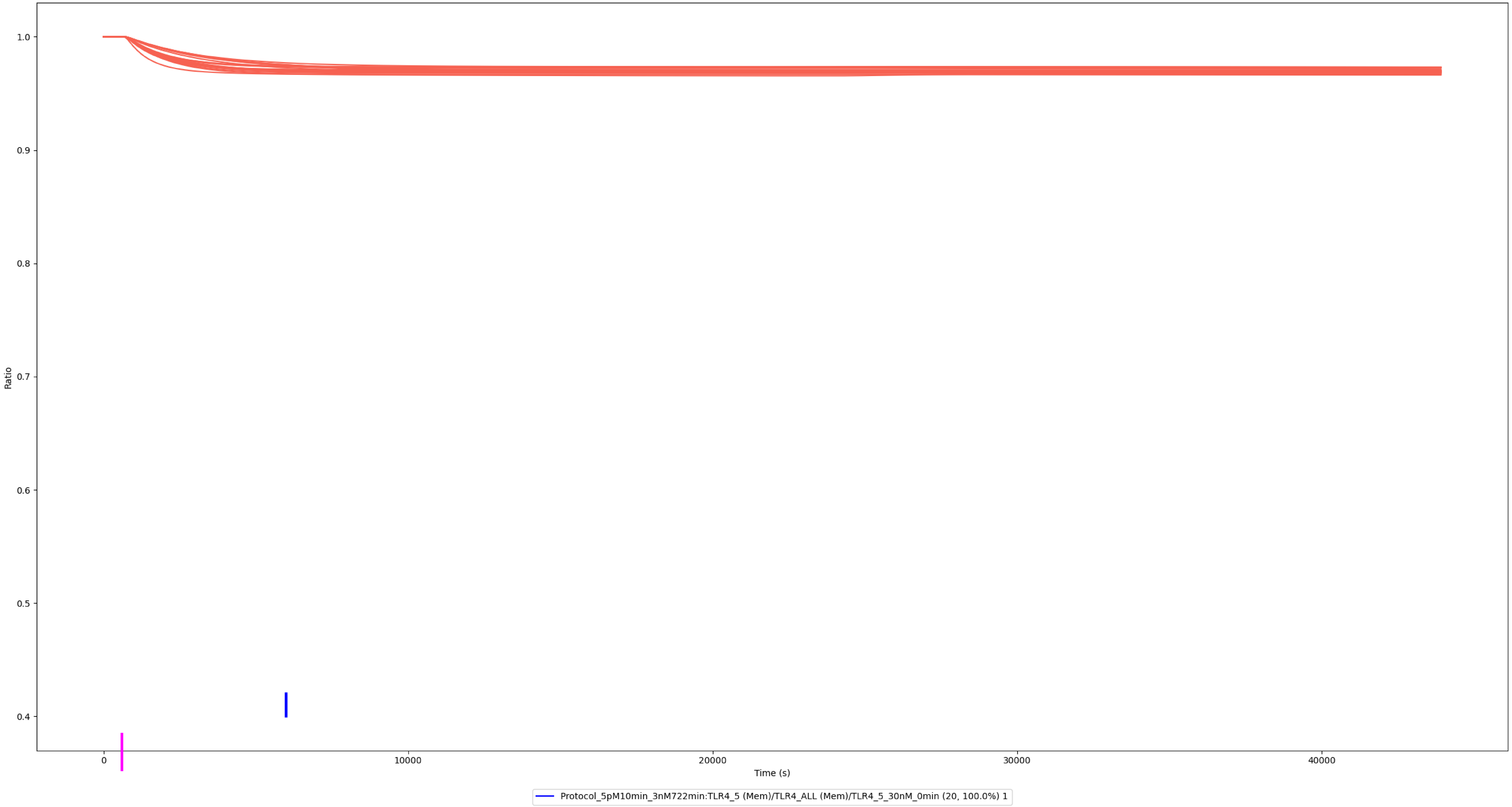

### Experiment vs. Simulation: 300 nM LPS, Cell Surface / Total TLR4

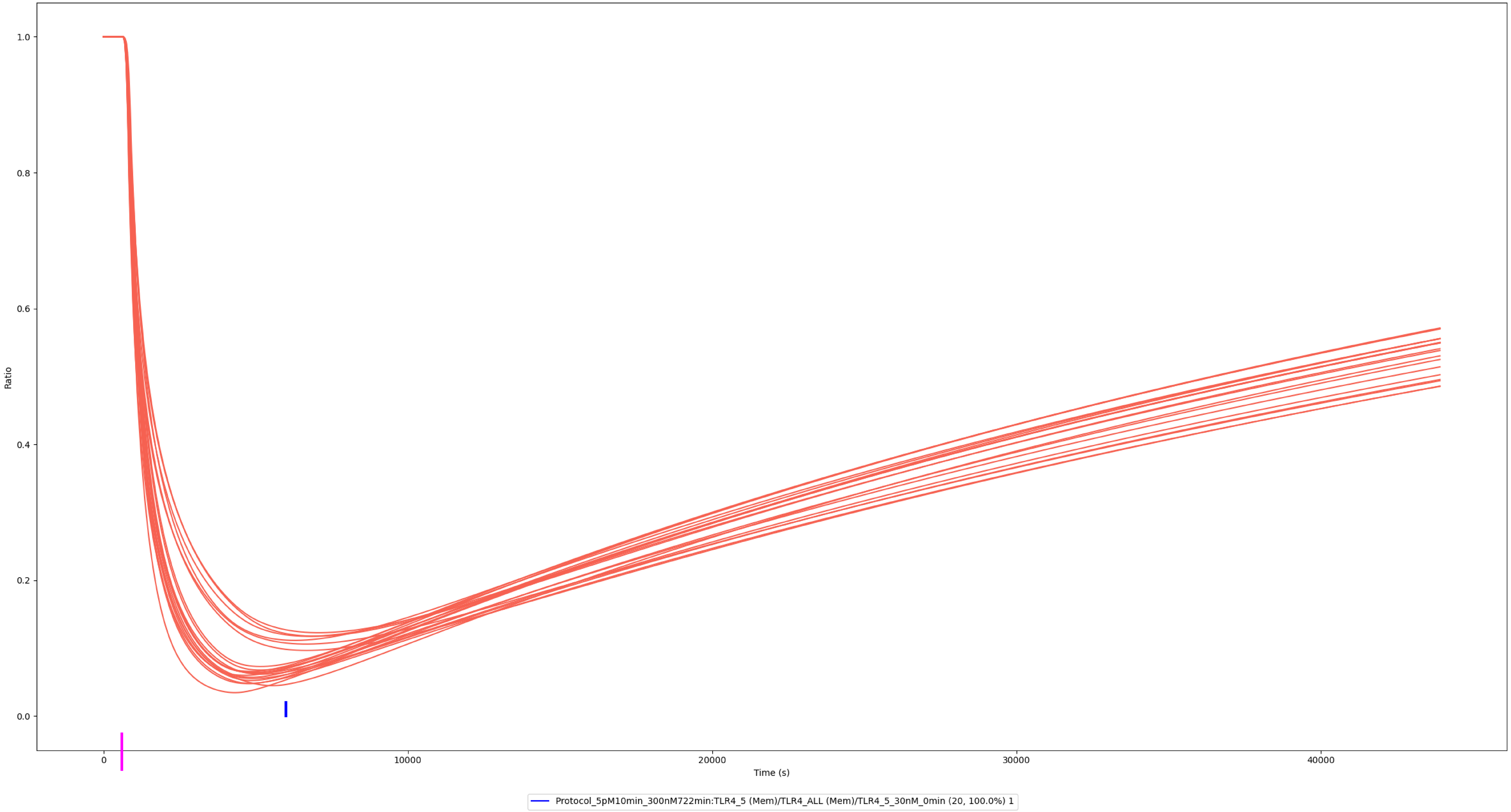

### MyD88

### Experiment vs. Simulation: 30 nM LPS, MAL:MyD88

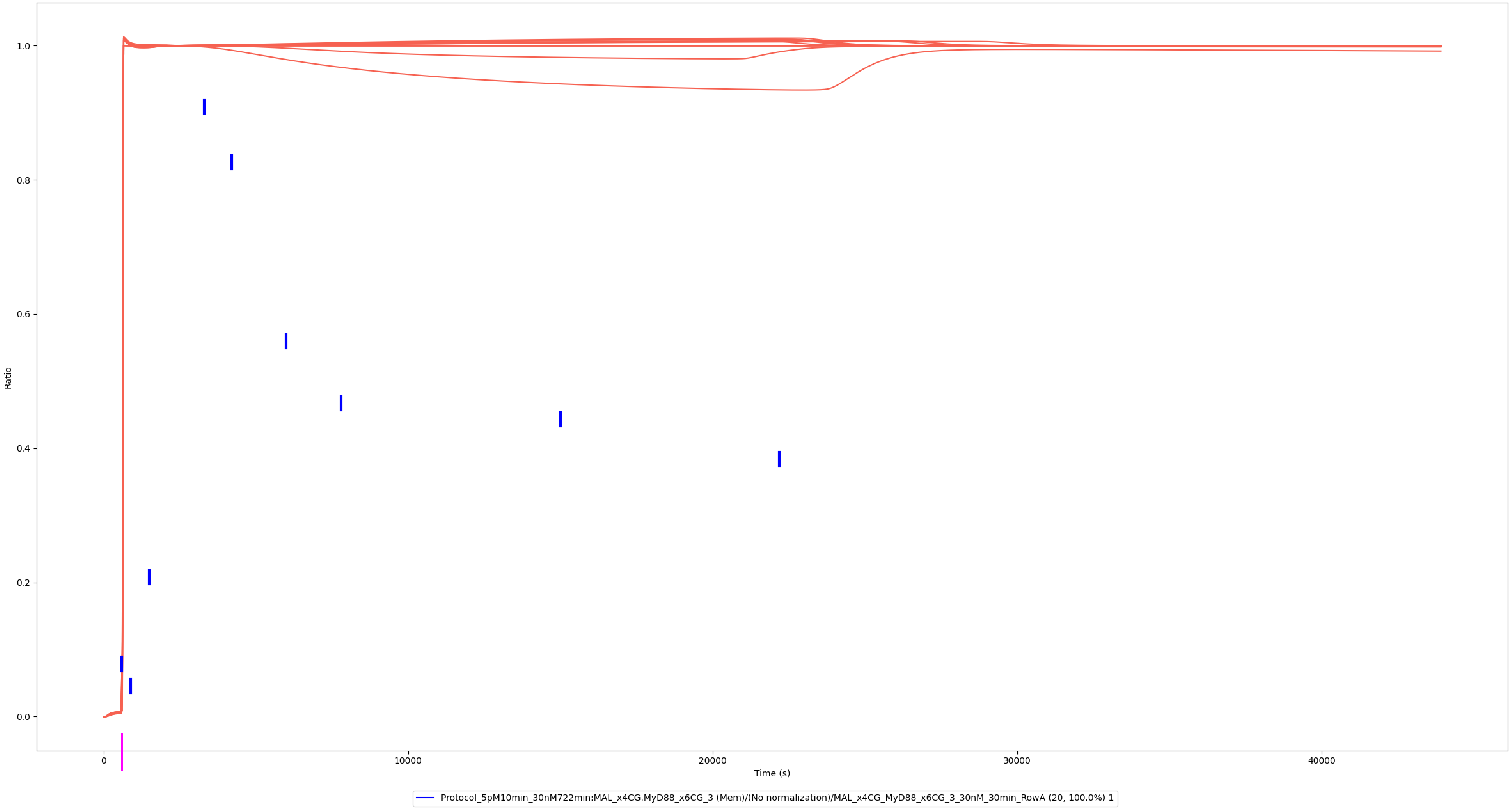

### Experiment vs. Simulation: 30 nM LPS, MAL:MyD88

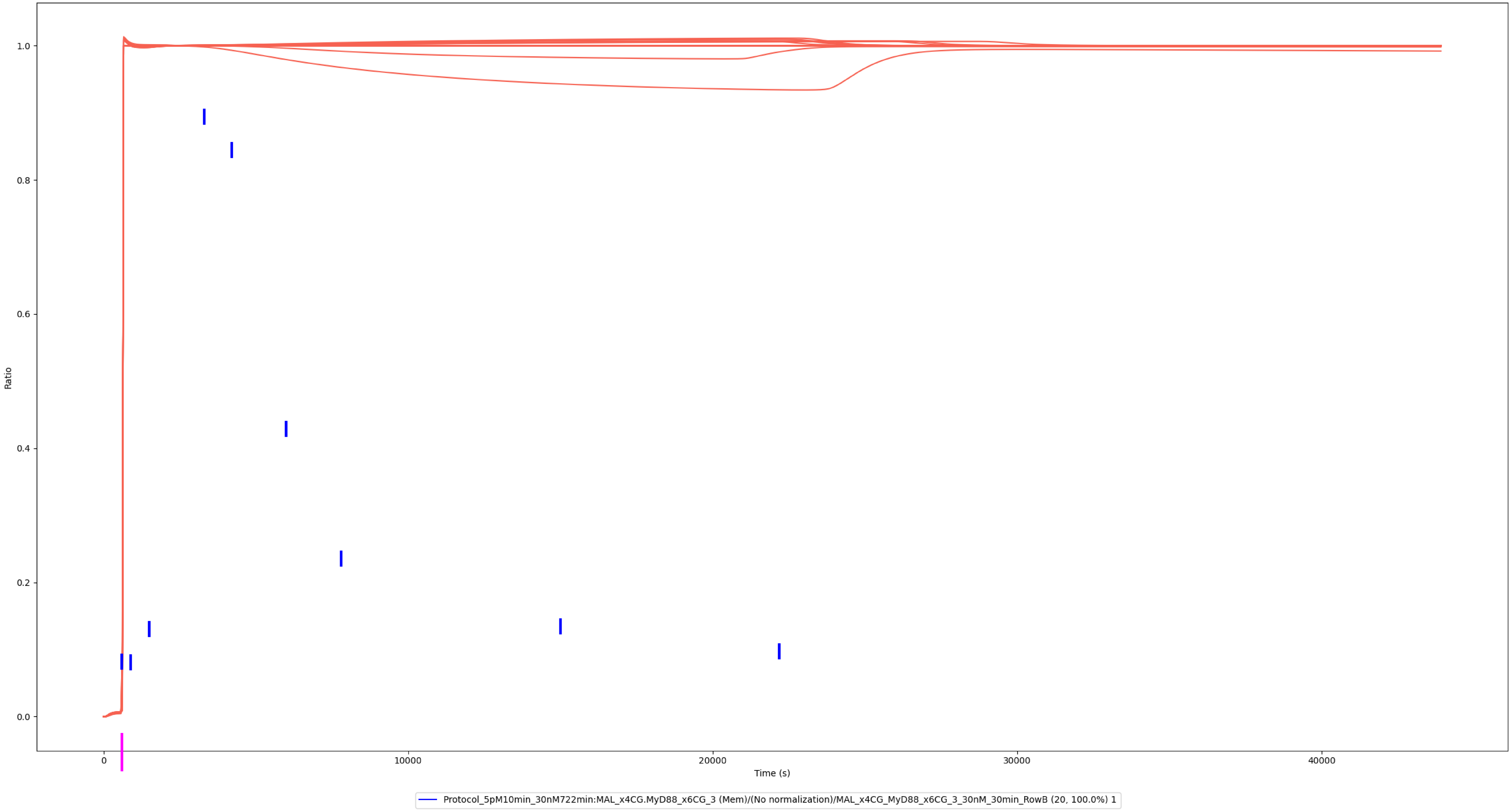

### IRAK4

### Experiment vs. Simulation: 30 nM LPS, MyD88:IRAK4

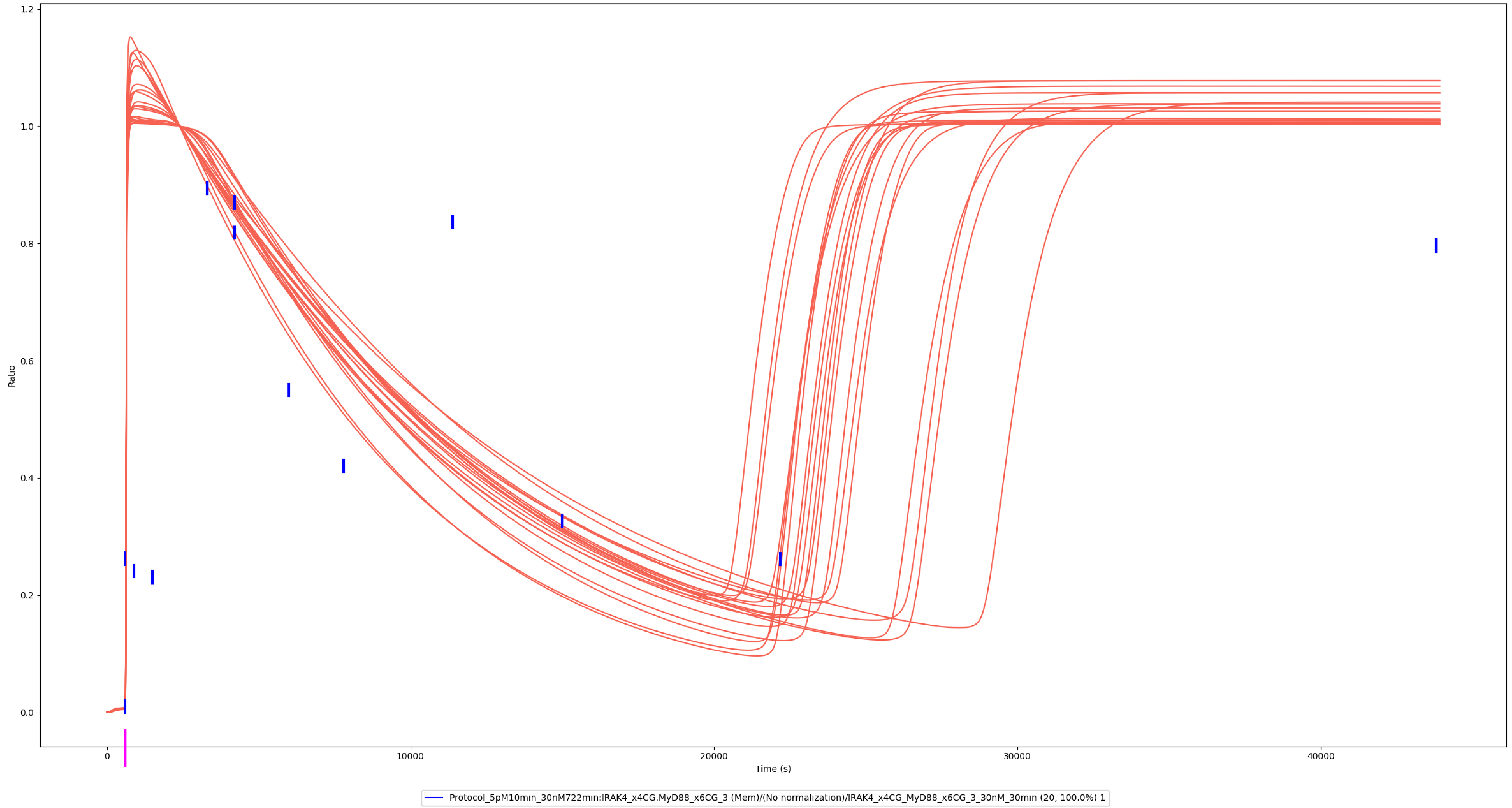

### Experiment vs. Simulation: 30 nM LPS, MyD88:IRAK4

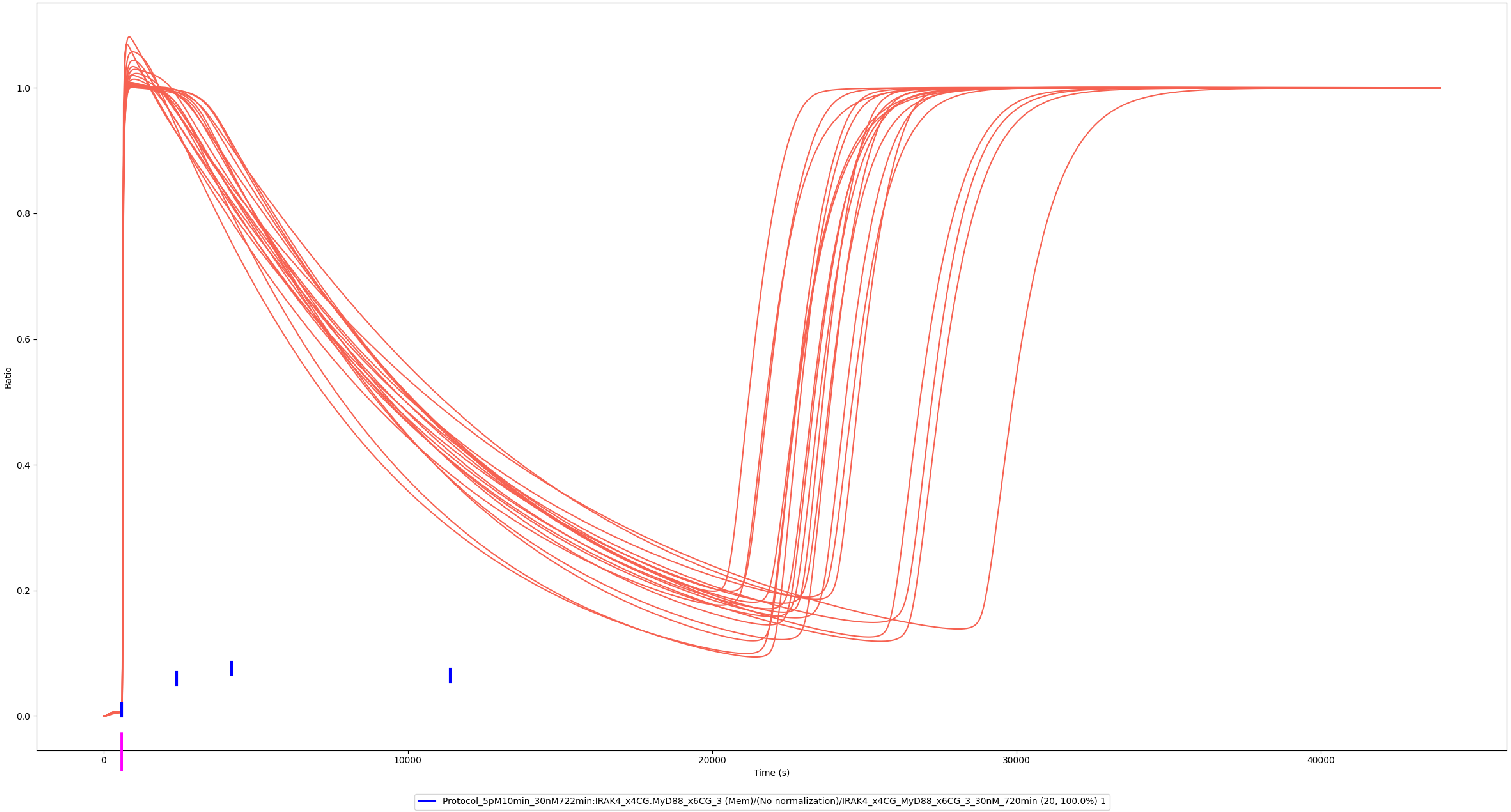

### Experiment vs. Simulation: 30 nM LPS, pppp-IRAK4

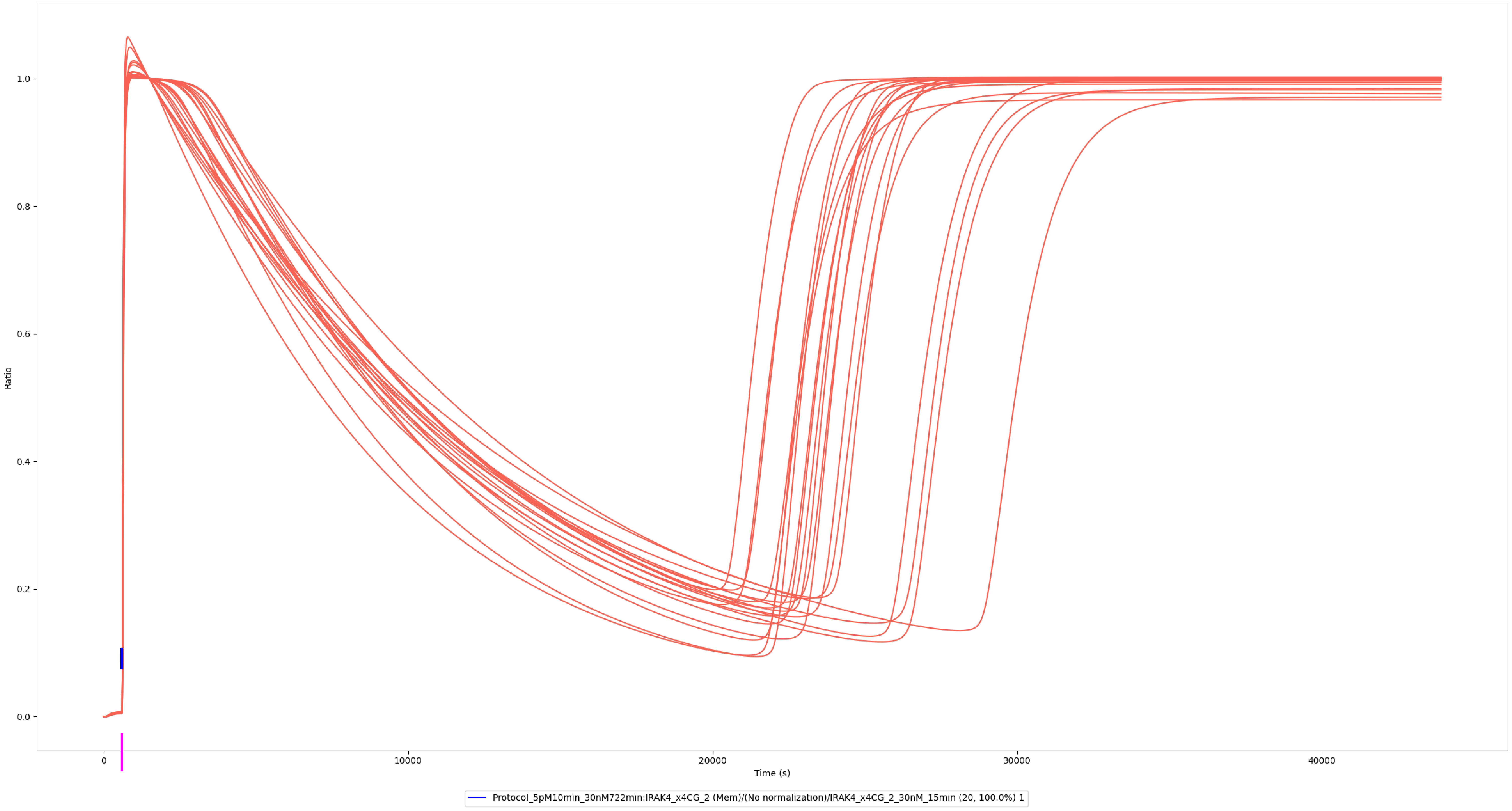

### IRAK1

### Experiment vs. Simulation: 30 nM LPS, IRAK4:IRAK1

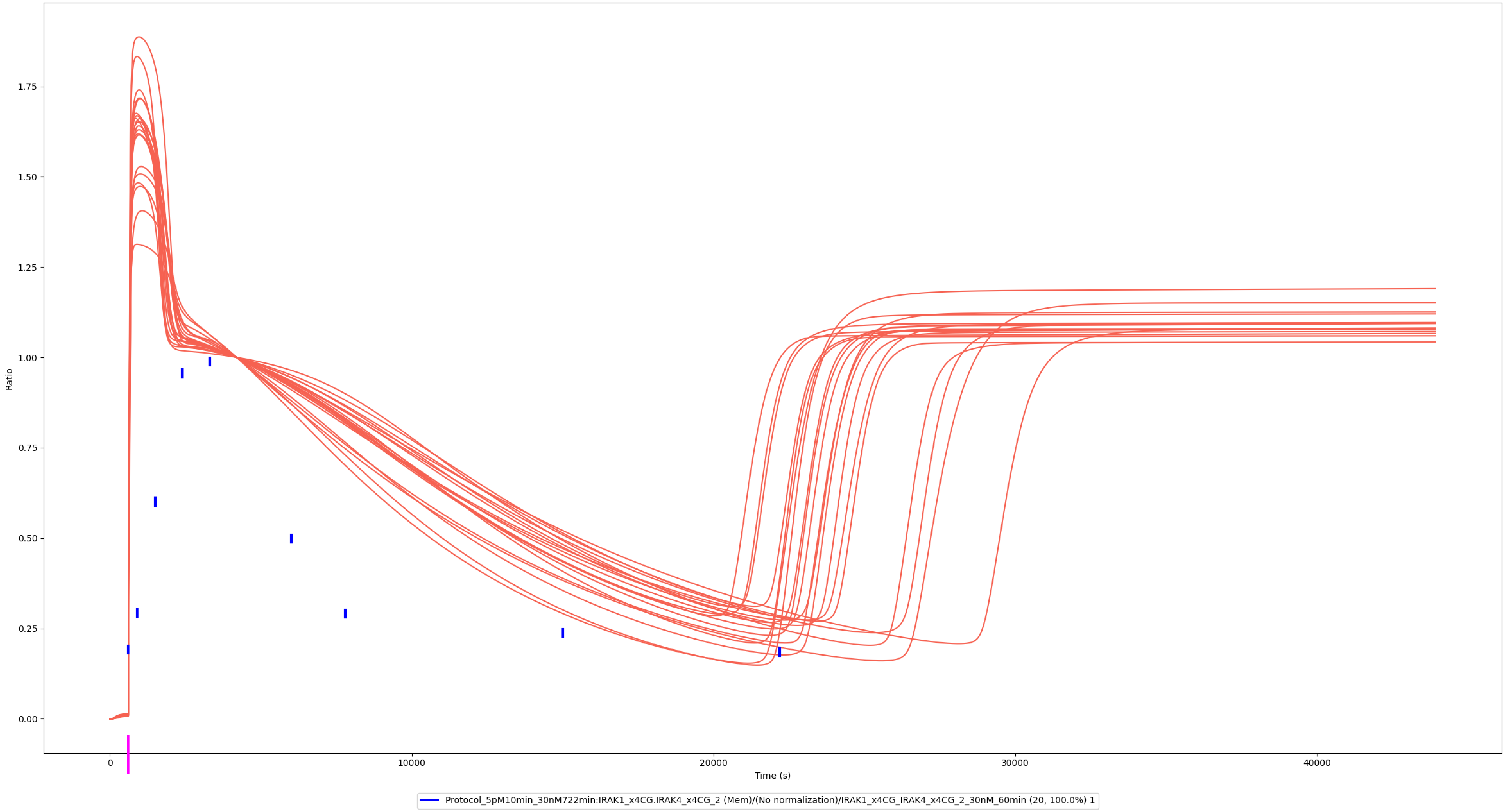

### IRAK2

### Experiment vs. Simulation: 30 nM LPS, IRAK4:IRAK2

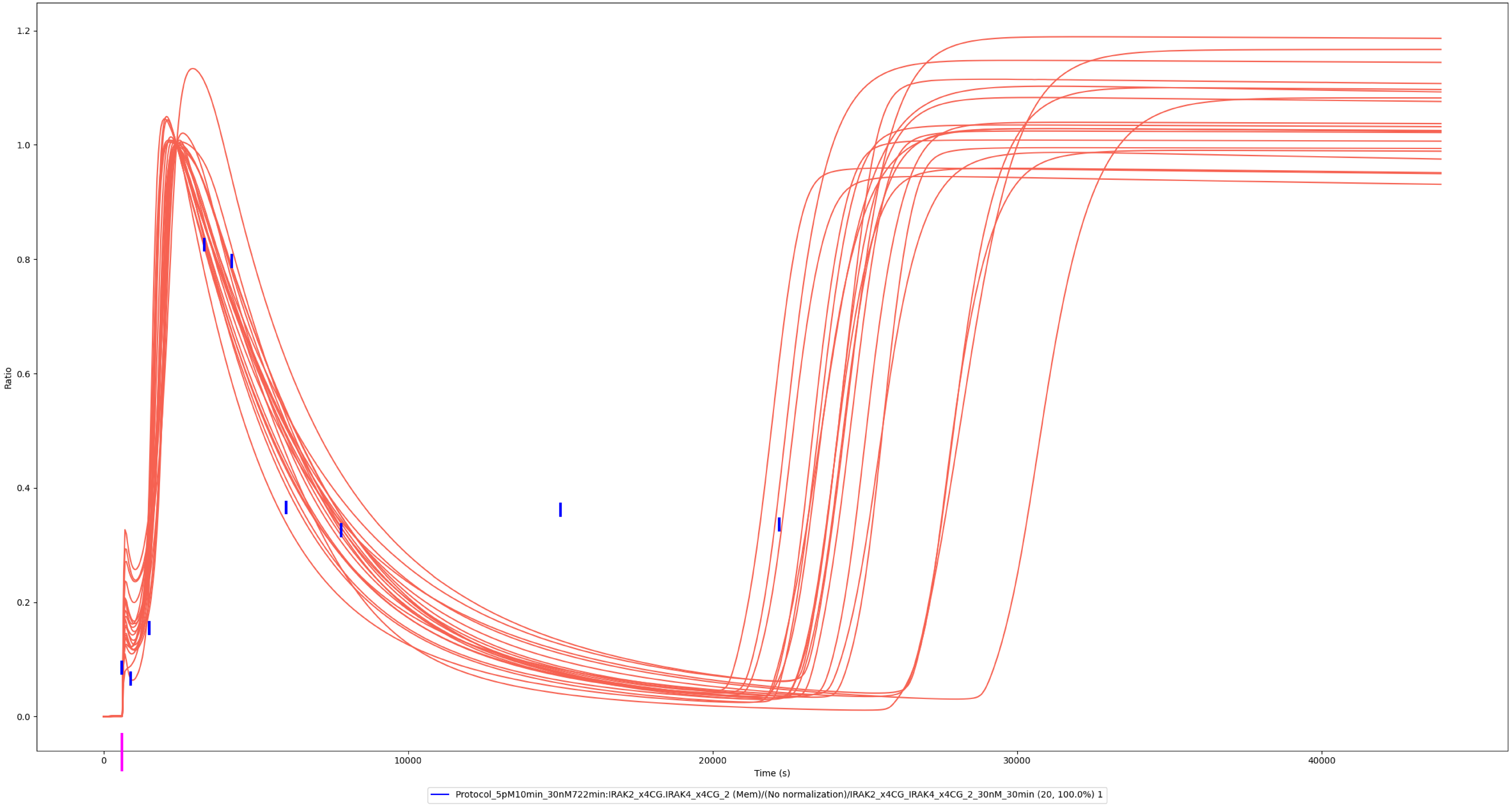

### Experiment vs. Simulation: 30 nM LPS, IRAK4:IRAK2

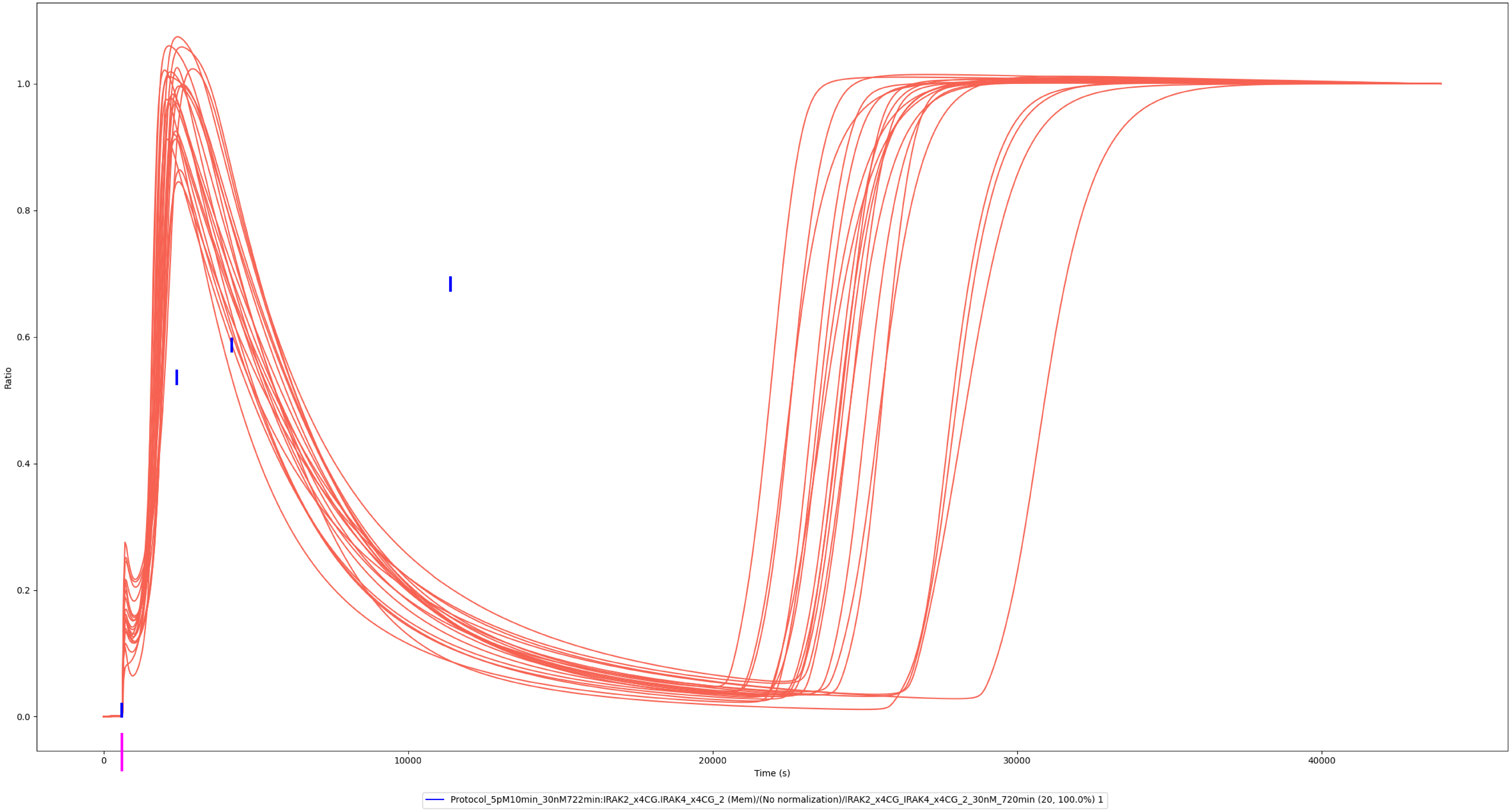

### Experiment vs. Simulation: 30 nM LPS, IRAK4:pp-IRAK2

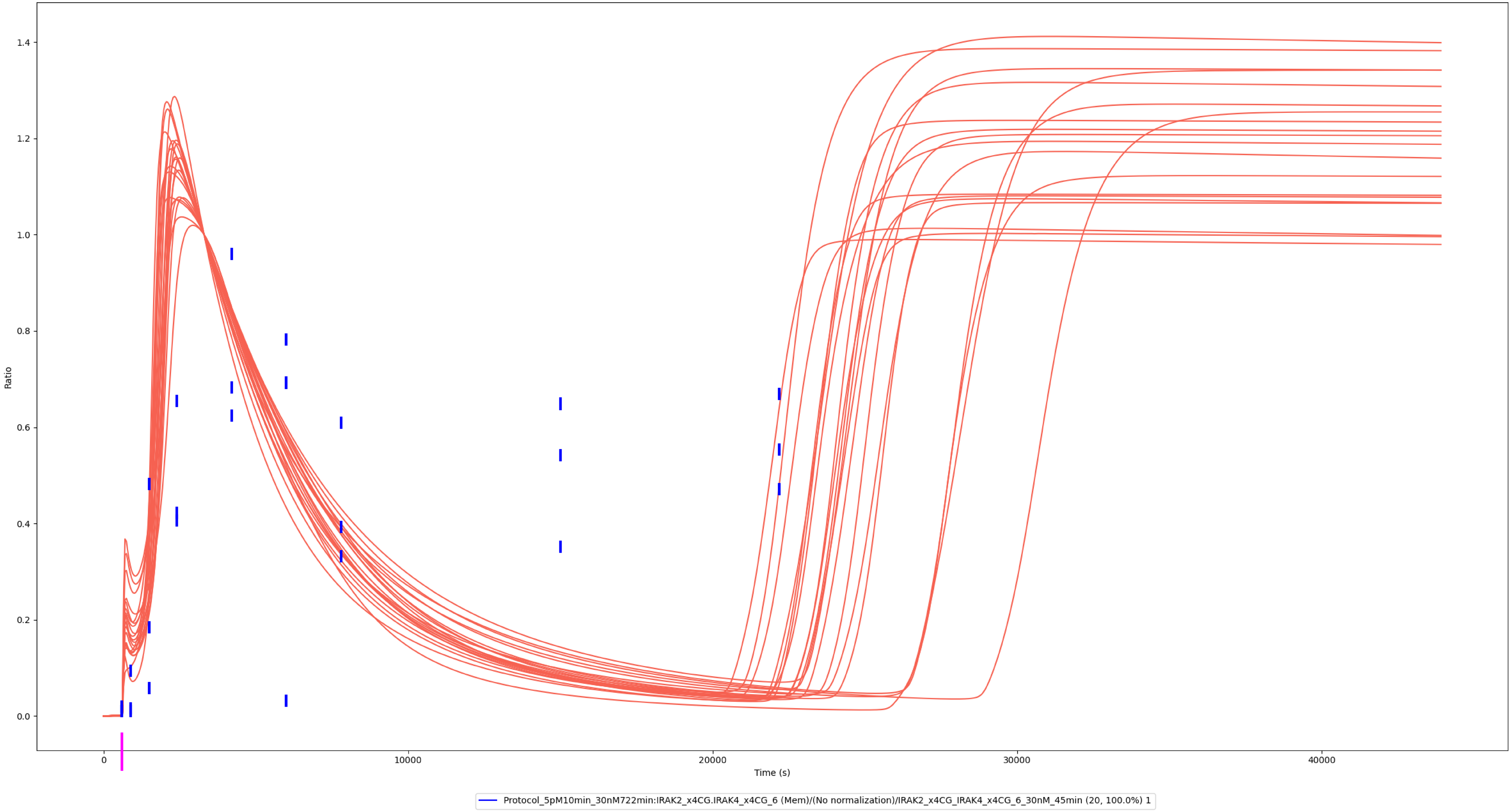

### Experiment vs. Simulation: 30 nM LPS, pp-IRAK2

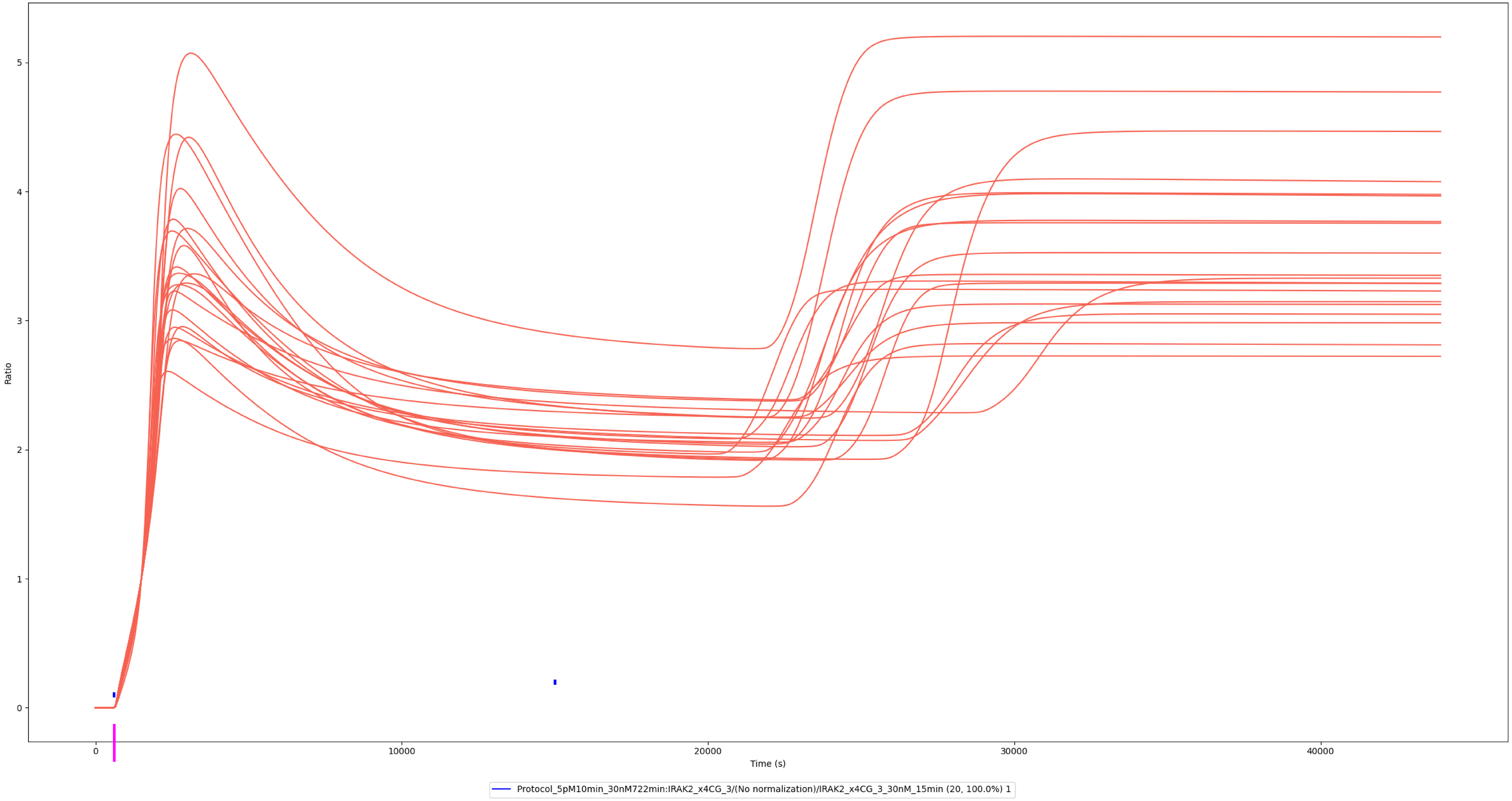

### Experiment vs. Simulation: 30 nM LPS, pp-IRAK2

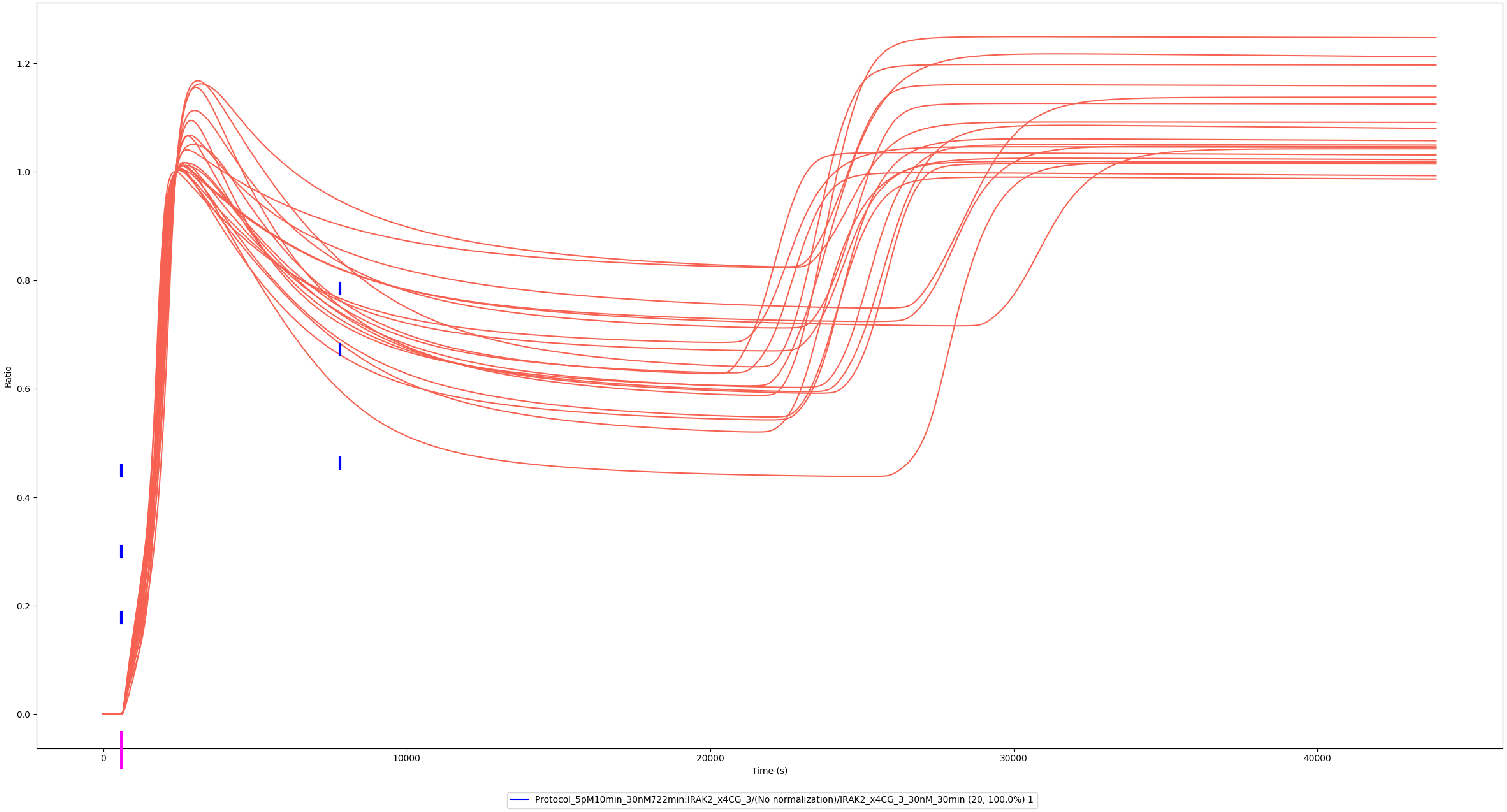

### TRAF6

### Experiment vs. Simulation: 30 nM LPS, IRAK1:TRAF6

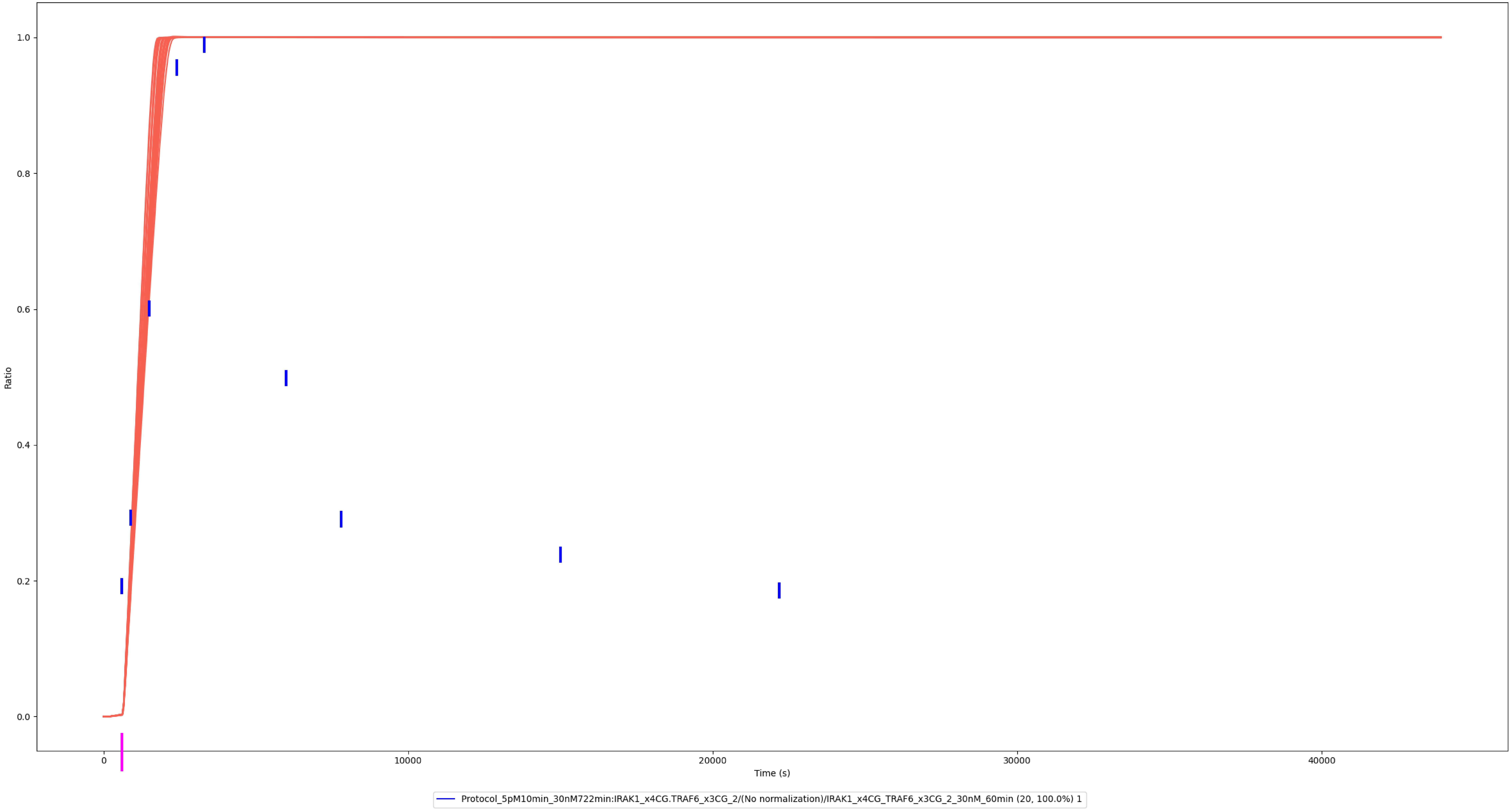

### Experiment vs. Simulation: 30 nM LPS, IRAK2:TRAF6

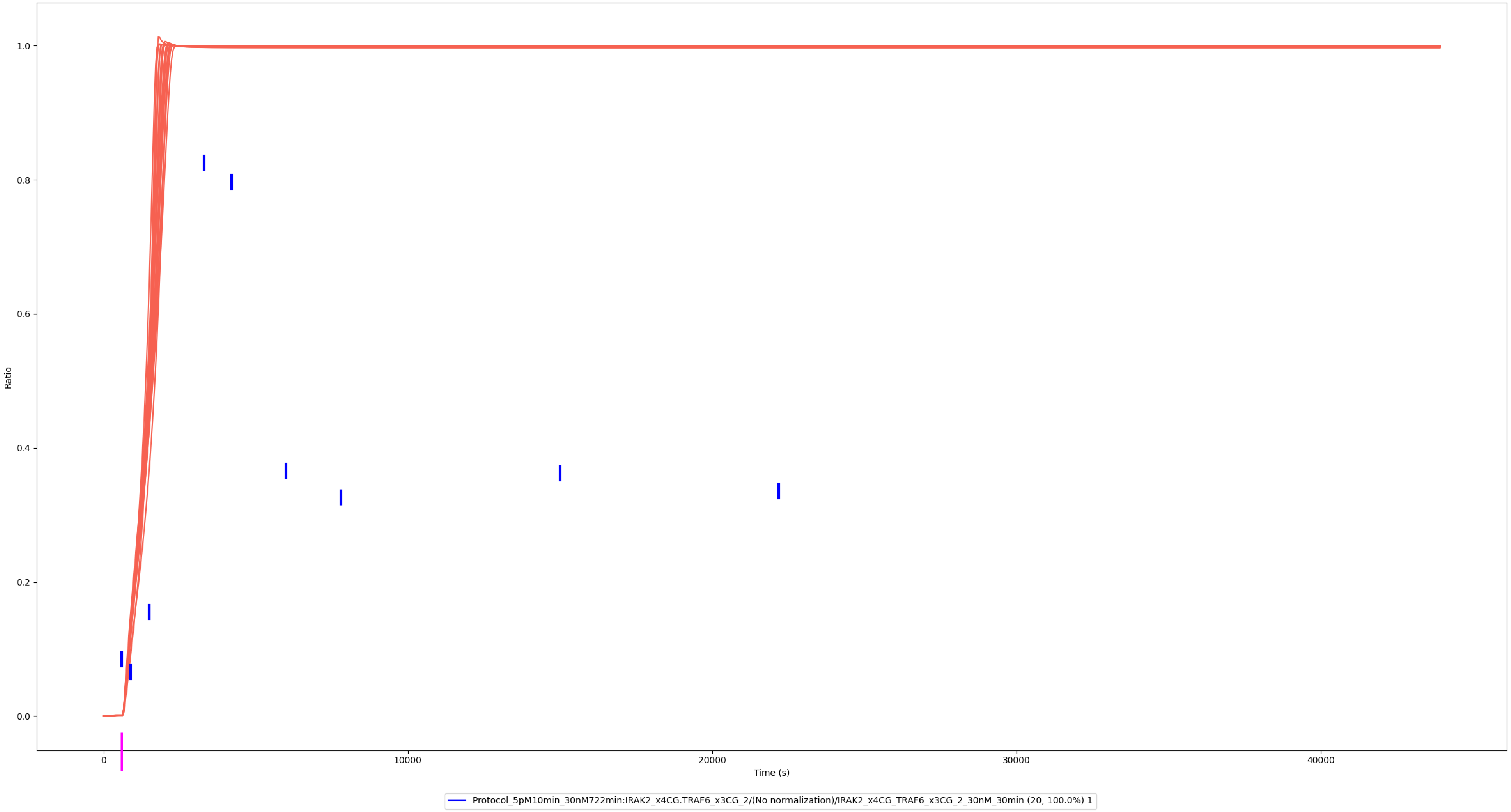

### Experiment vs. Simulation: 30 nM LPS, pp-IRAK2:TRAF6

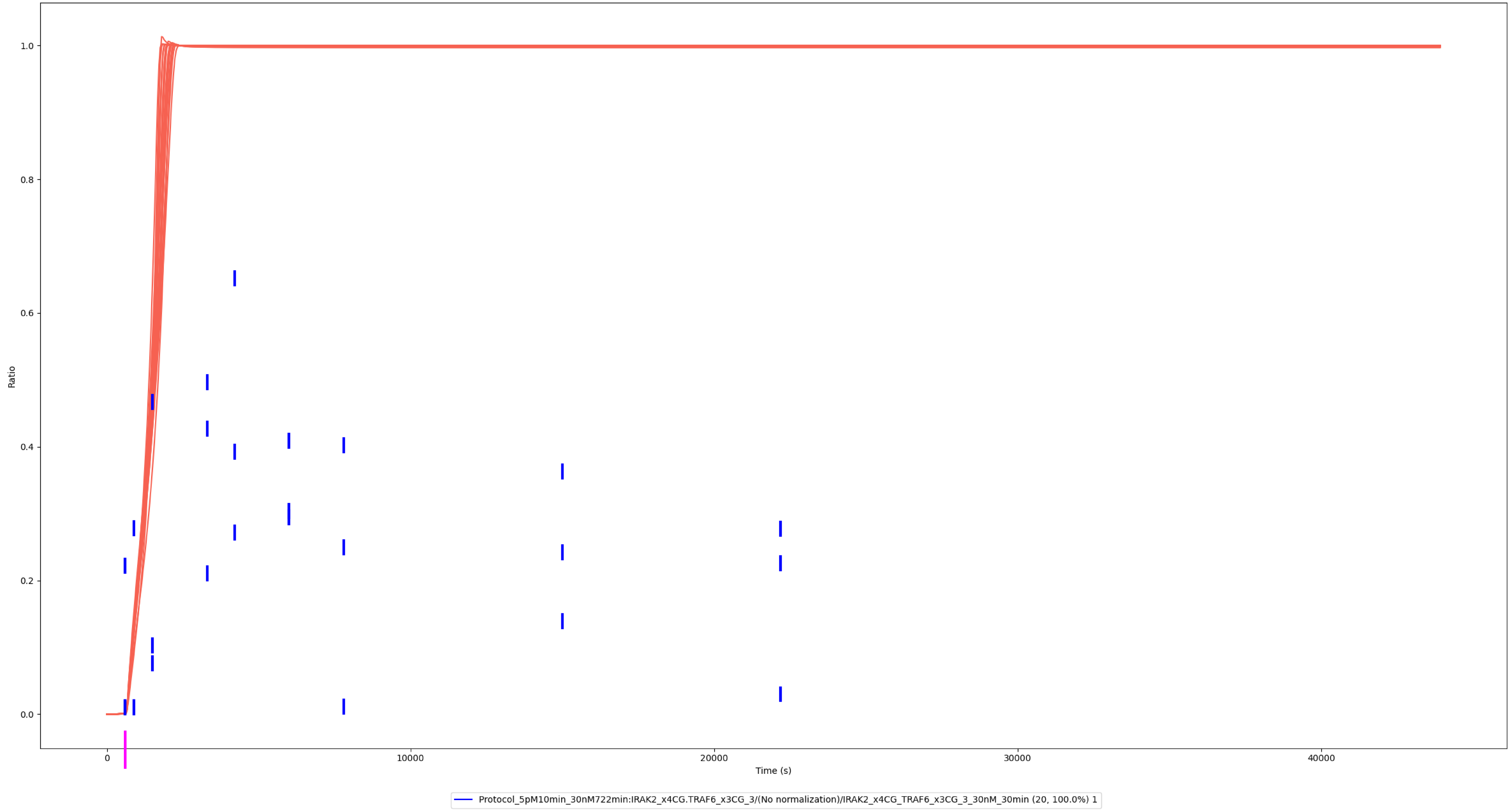

### NEMO

### Experiment vs. Simulation: 30 nM LPS, IRAK1:TRAF6:NEMO:IKKα/β

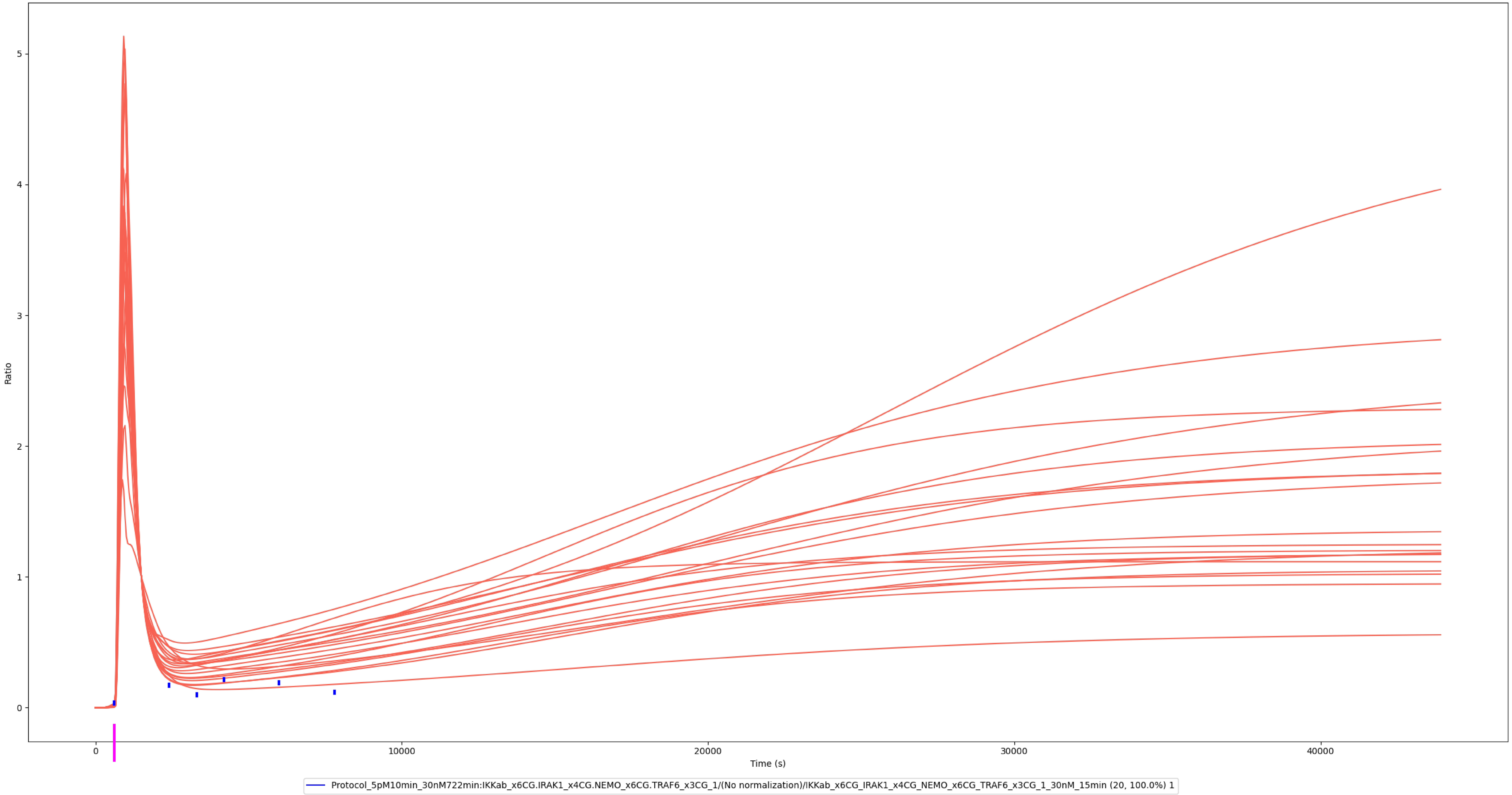

### Experiment vs. Simulation: 30 nM LPS, IRAK2:TRAF6:NEMO:IKKα/β

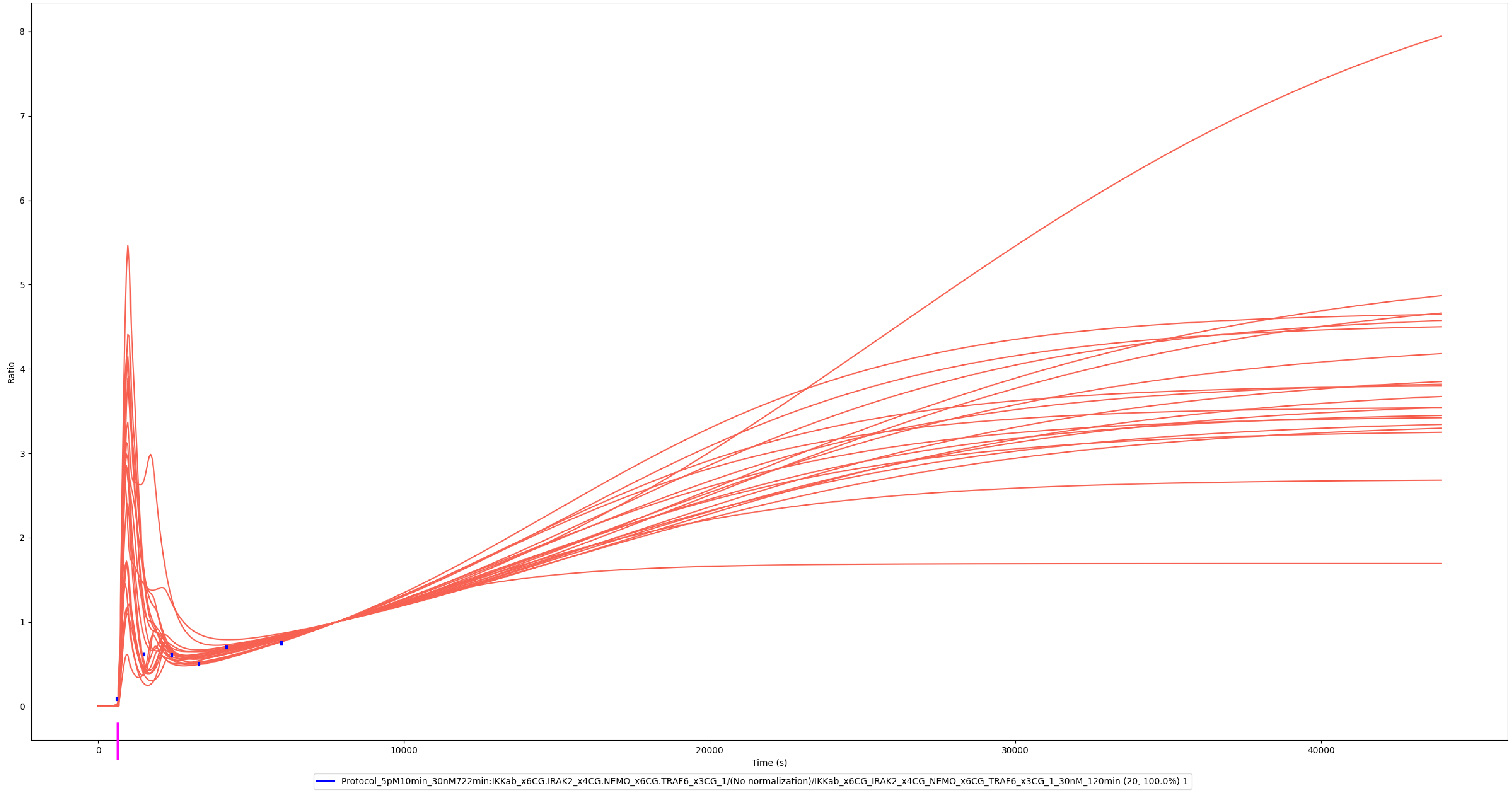

### IKKα/β

### Experiment vs. Simulation: 1 nM LPS, p-IKKα/β

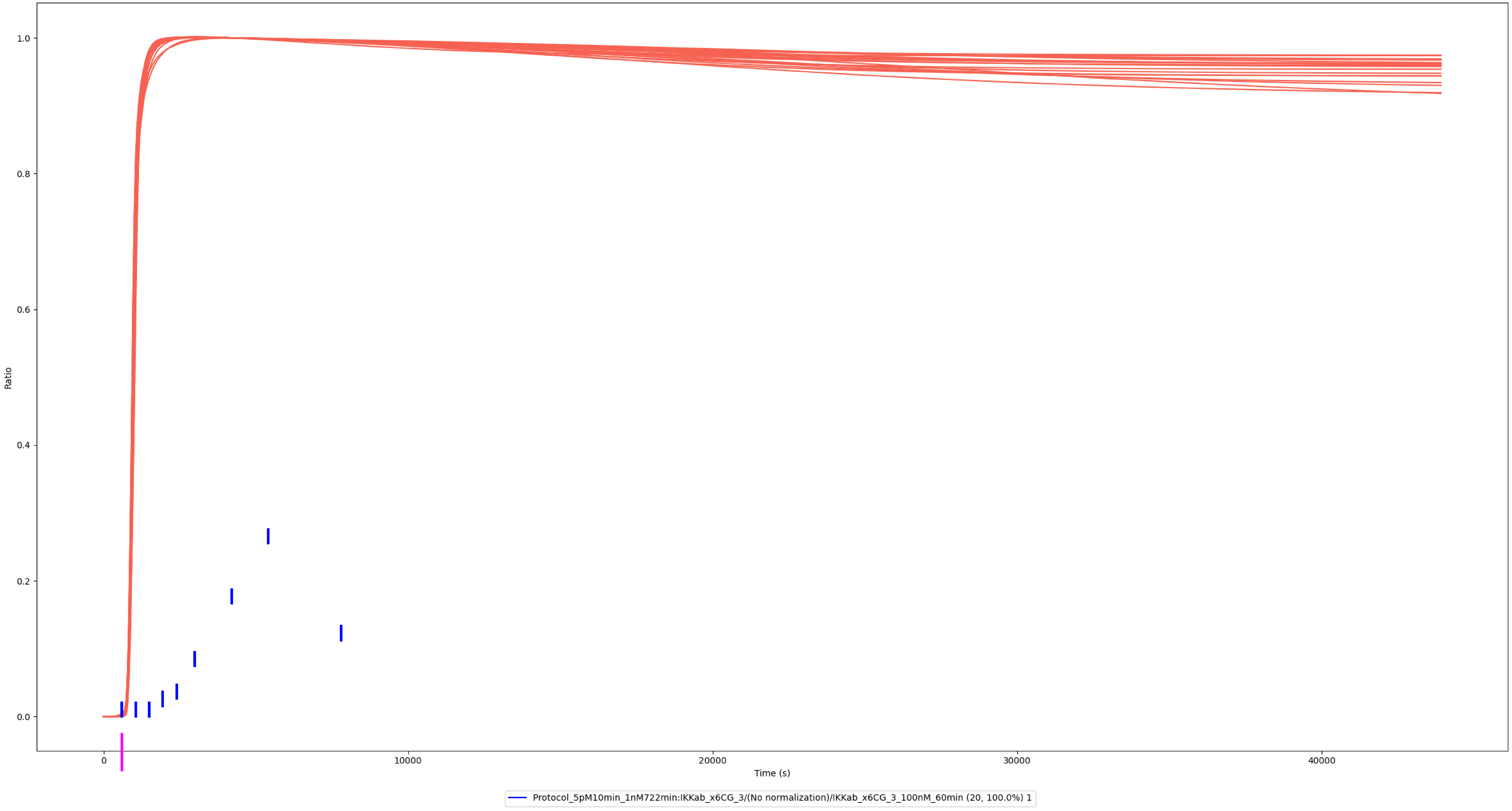

### Experiment vs. Simulation: 10 nM LPS, p-IKKα/β

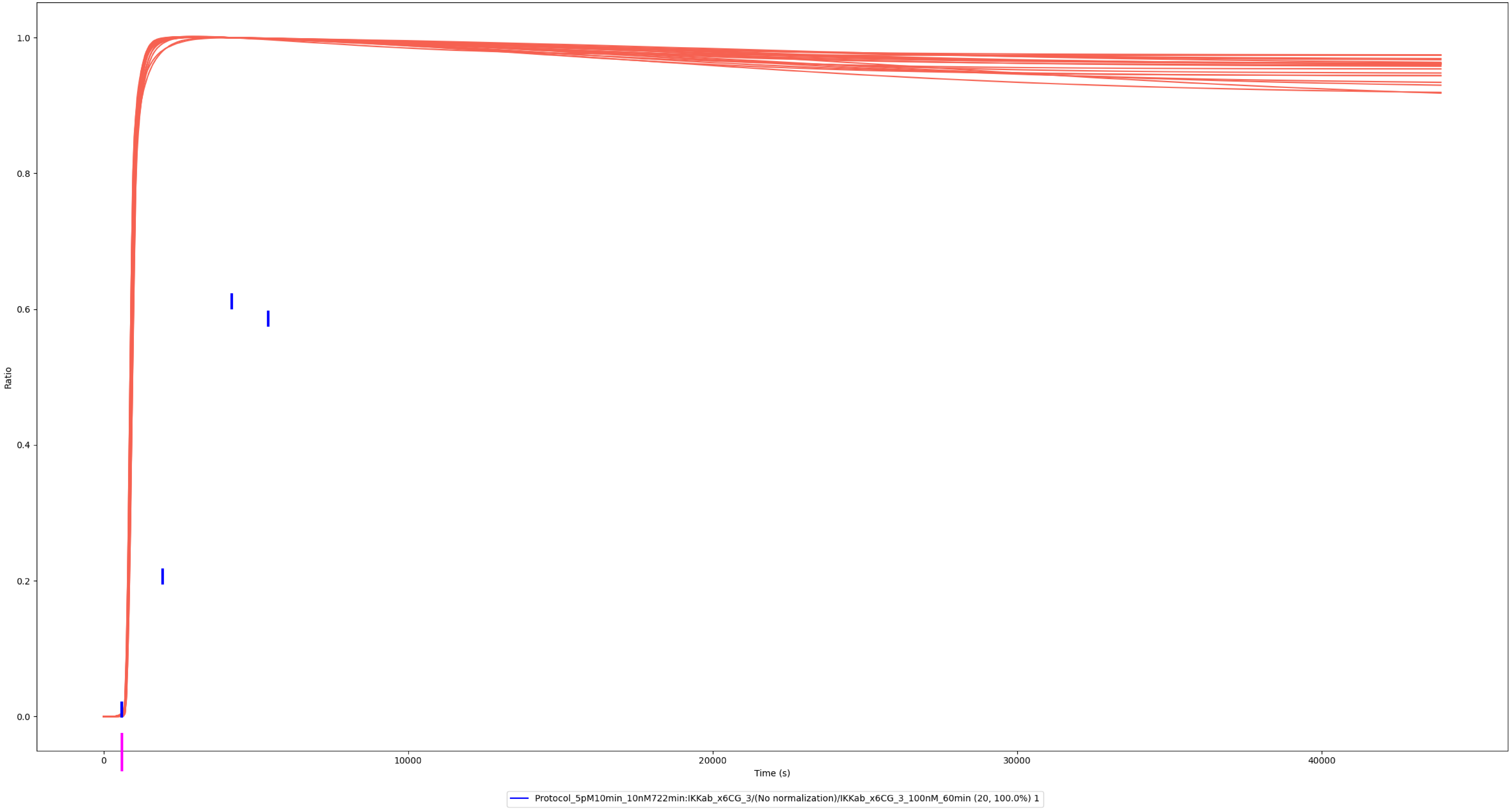

### Experiment vs. Simulation: 100 nM LPS, p-IKKα/β

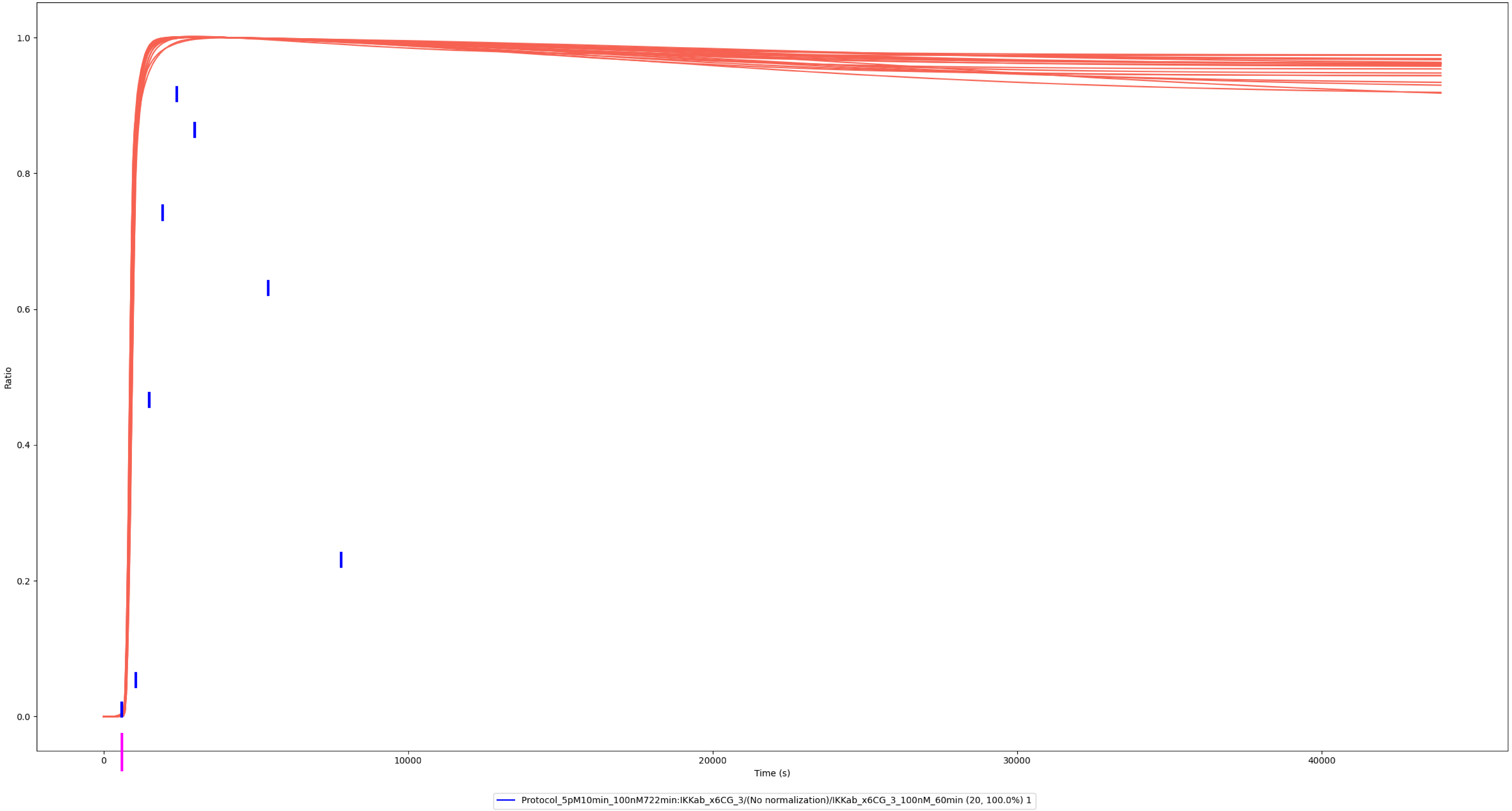

### Experiment vs. Simulation: 1.5 nM LPS, pp-IKKα/β

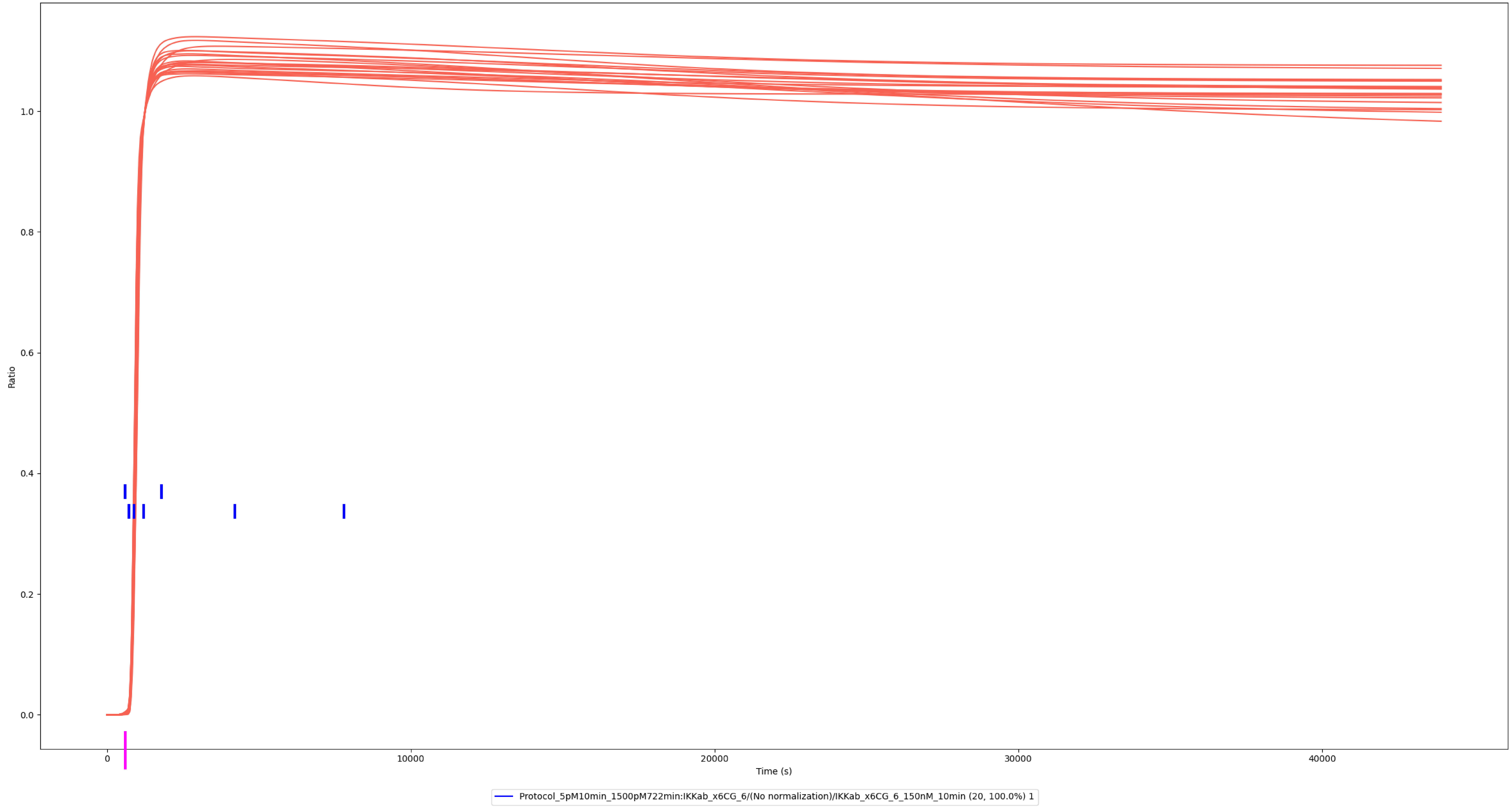

### Experiment vs. Simulation: 15 nM LPS, pp-IKKα/β

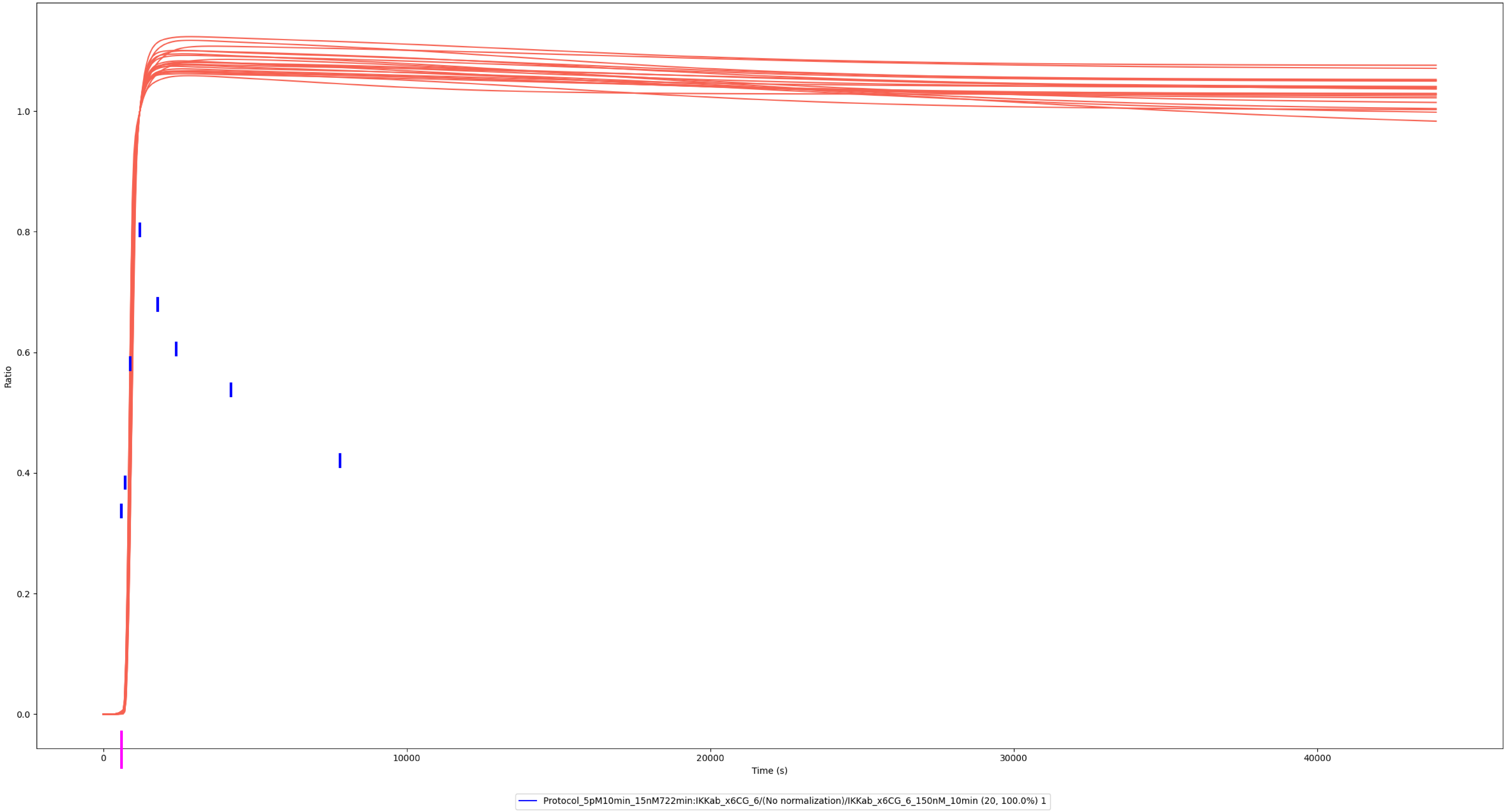

### Experiment vs. Simulation: 150 nM LPS, pp-IKKα/β

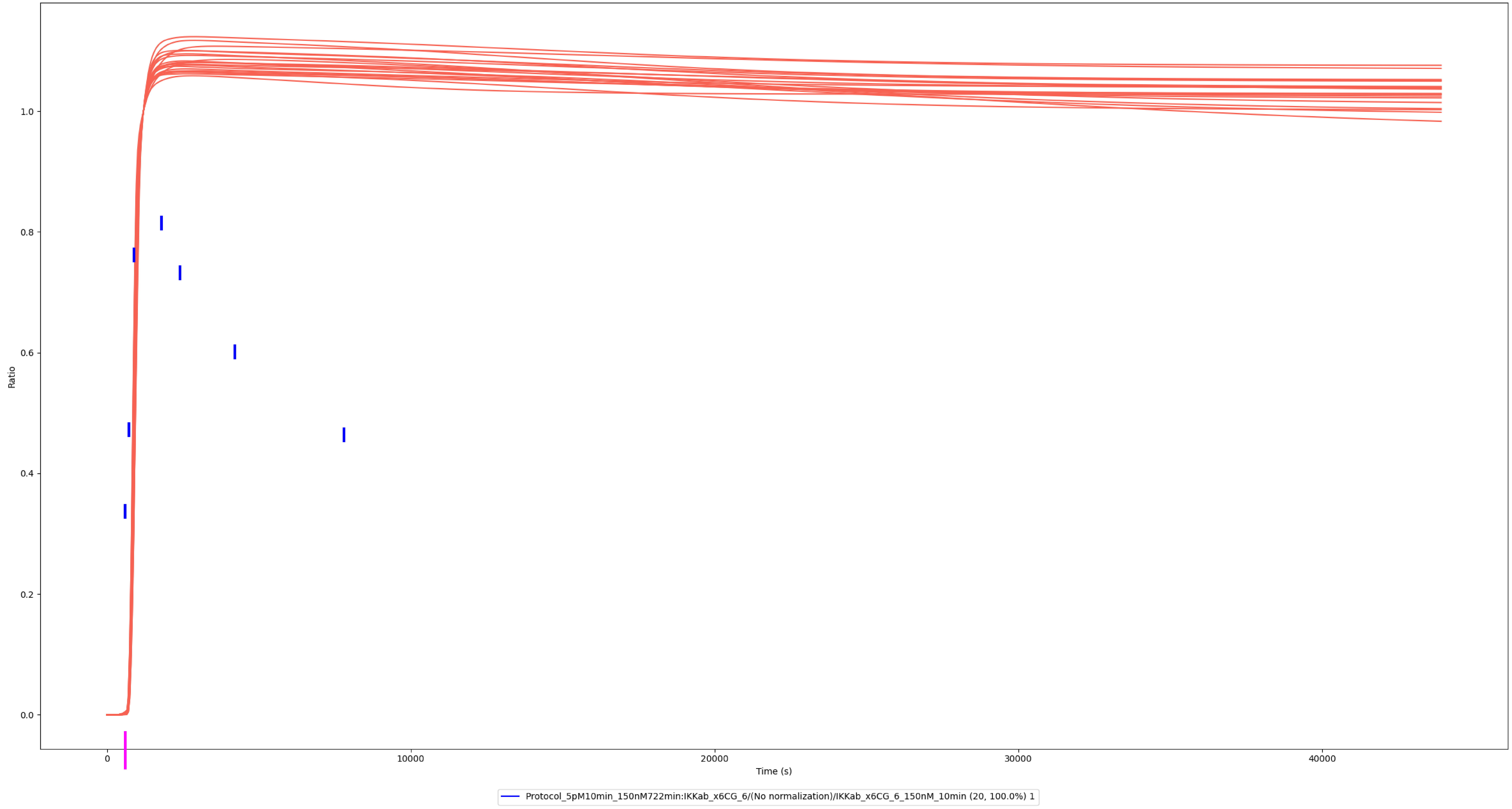

### IκBα

### Experiment vs. Simulation: 0.01 nM LPS, IκBα

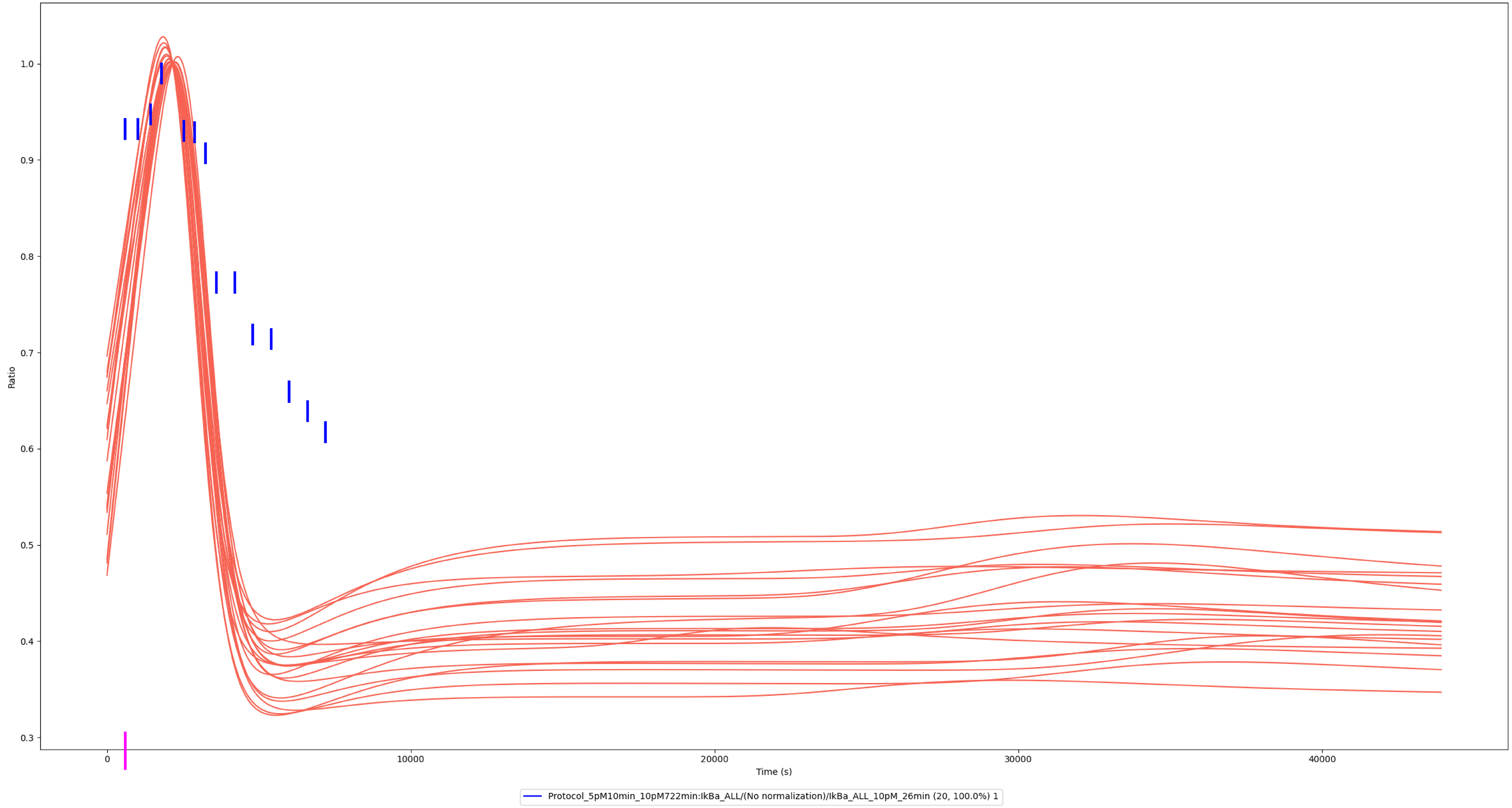

### Experiment vs. Simulation: 0.1 nM LPS, IκBα

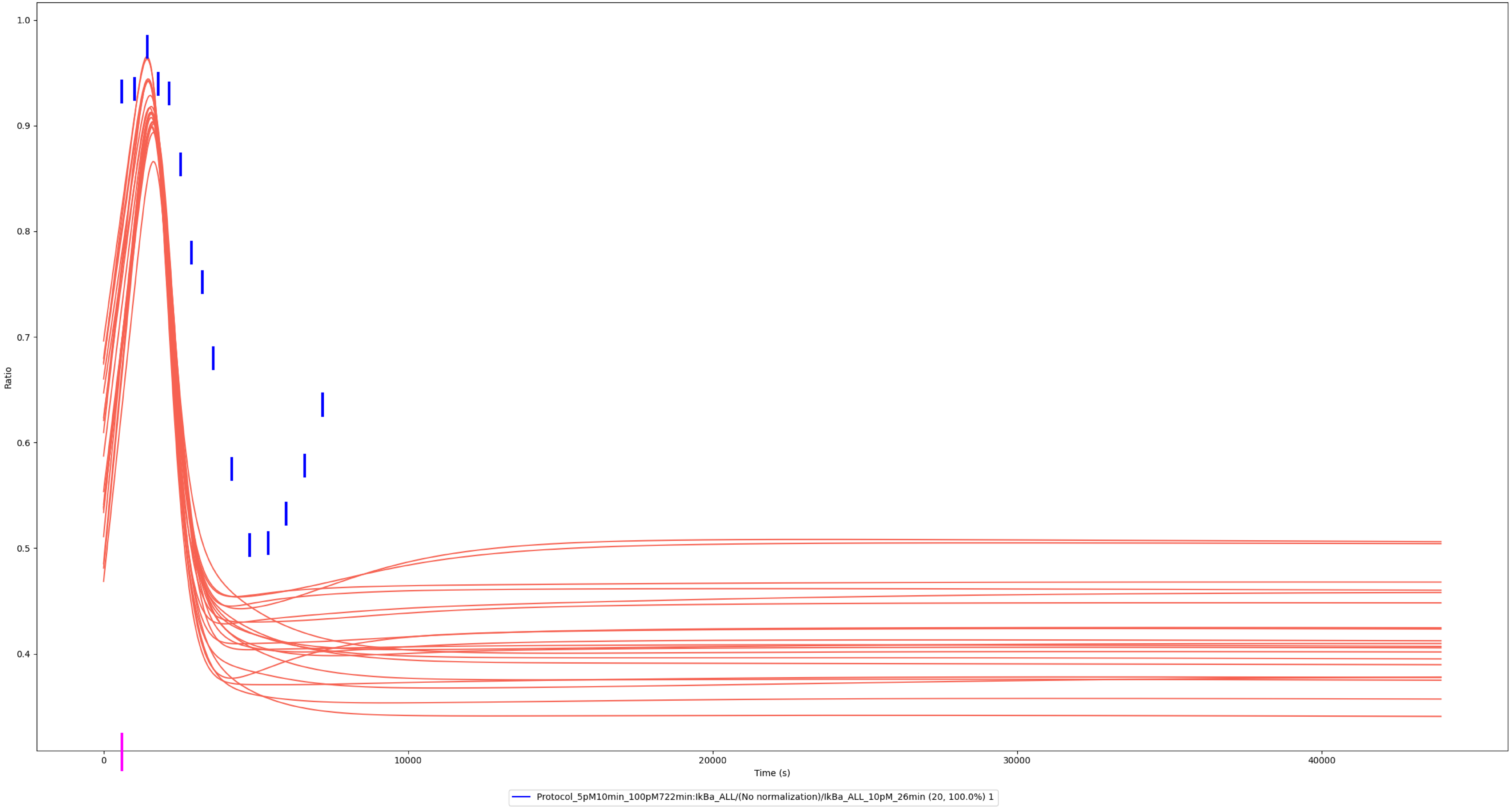

### Experiment vs. Simulation: 1 nM LPS, IκBα

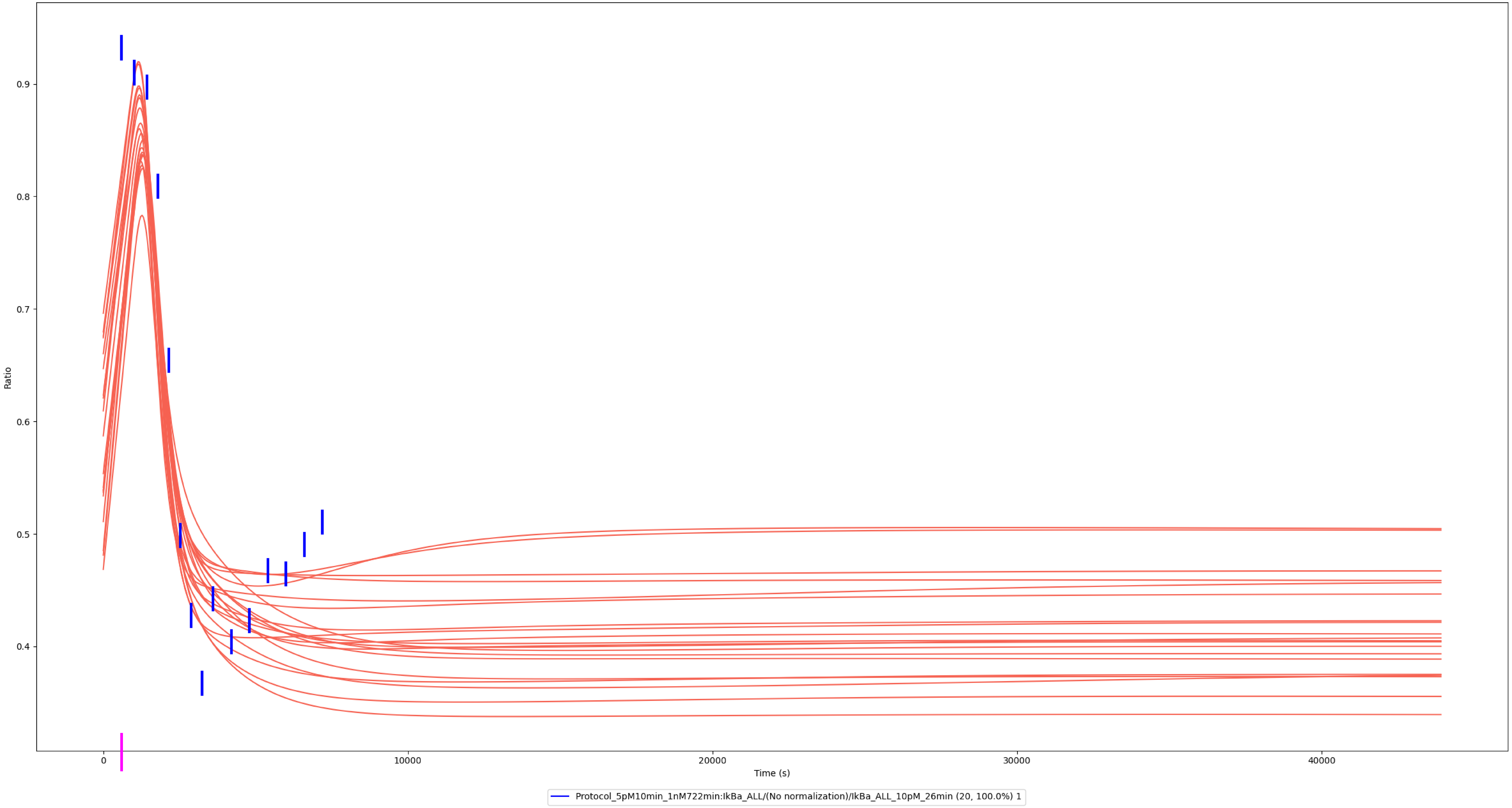

### Experiment vs. Simulation: 10 nM LPS, IκBα

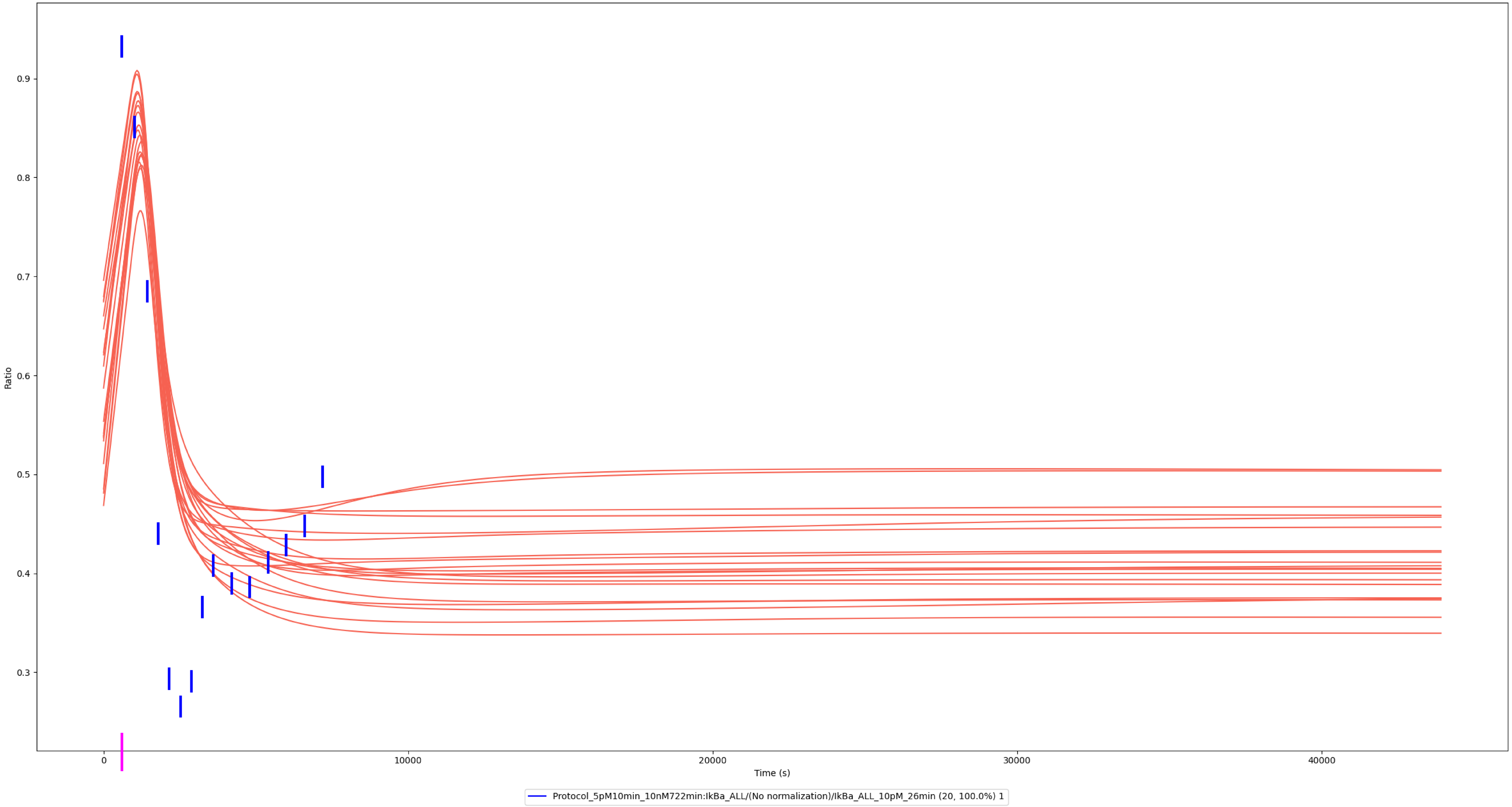

### Experiment vs. Simulation: 100 nM LPS, IκBα

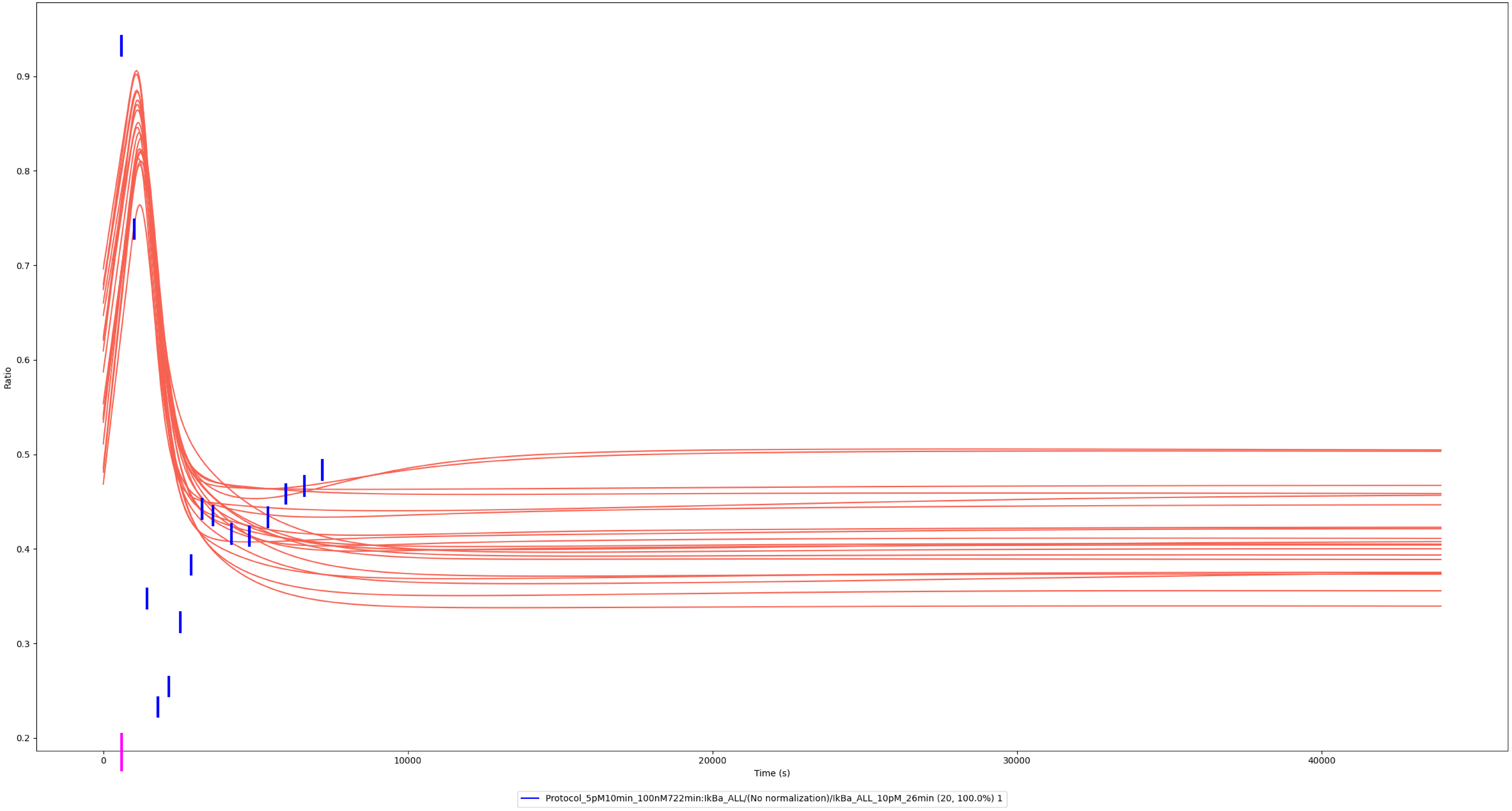

### Experiment vs. Simulation: 1,000 nM LPS, IκBα

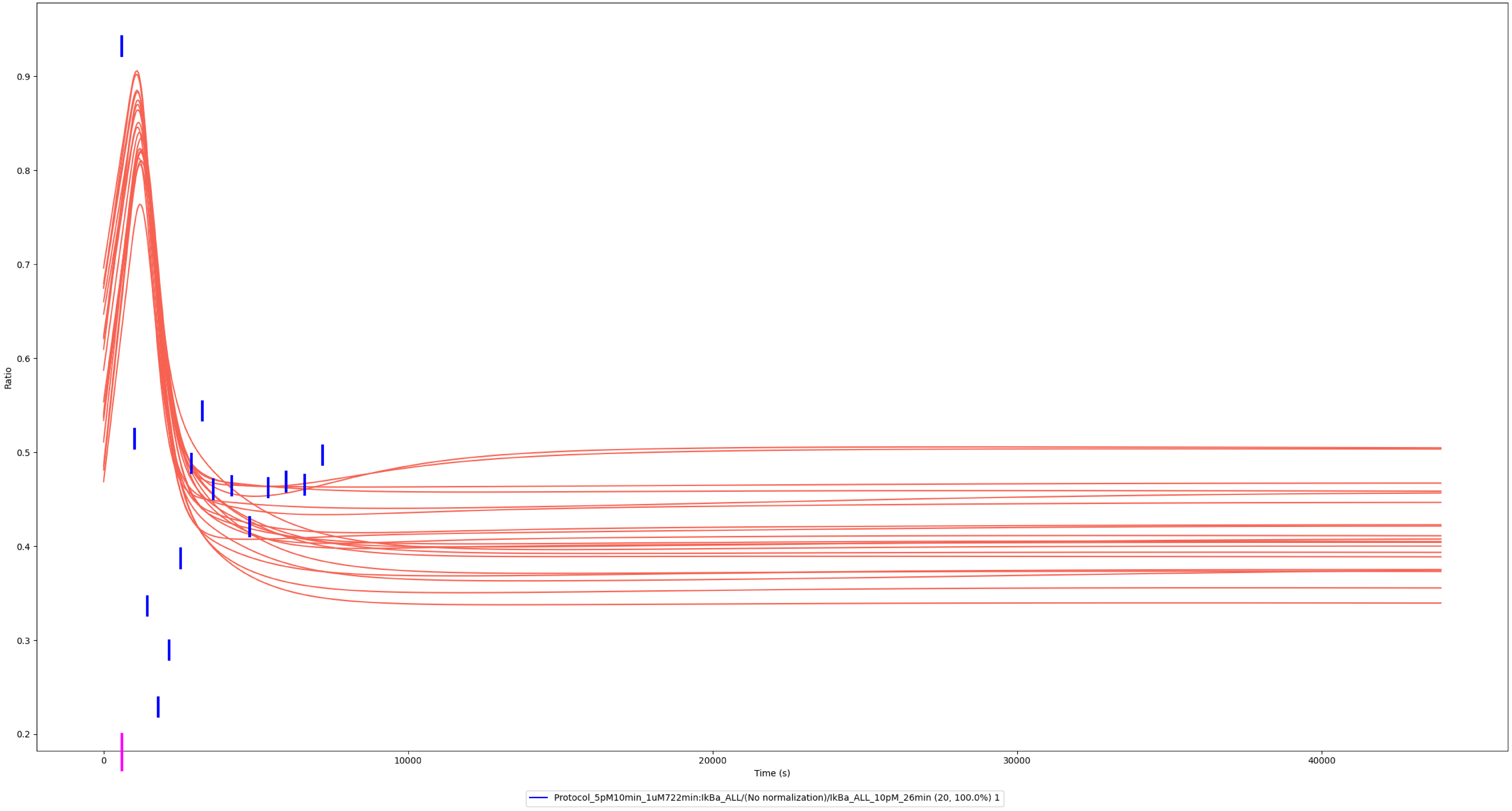

### Experiment vs. Simulation: 1 nM LPS, IκBα

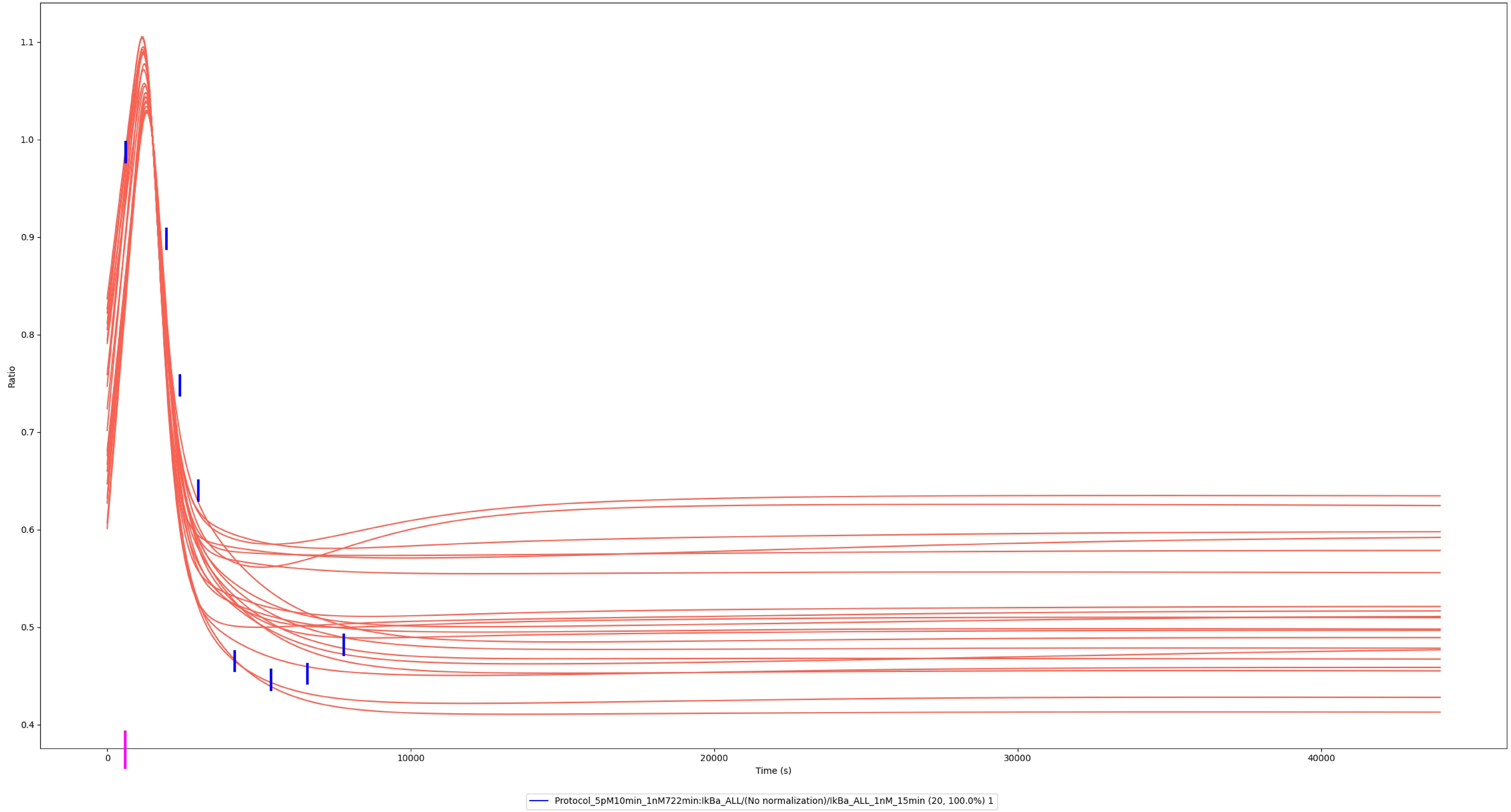

### Experiment vs. Simulation: 100 nM LPS, IκBα

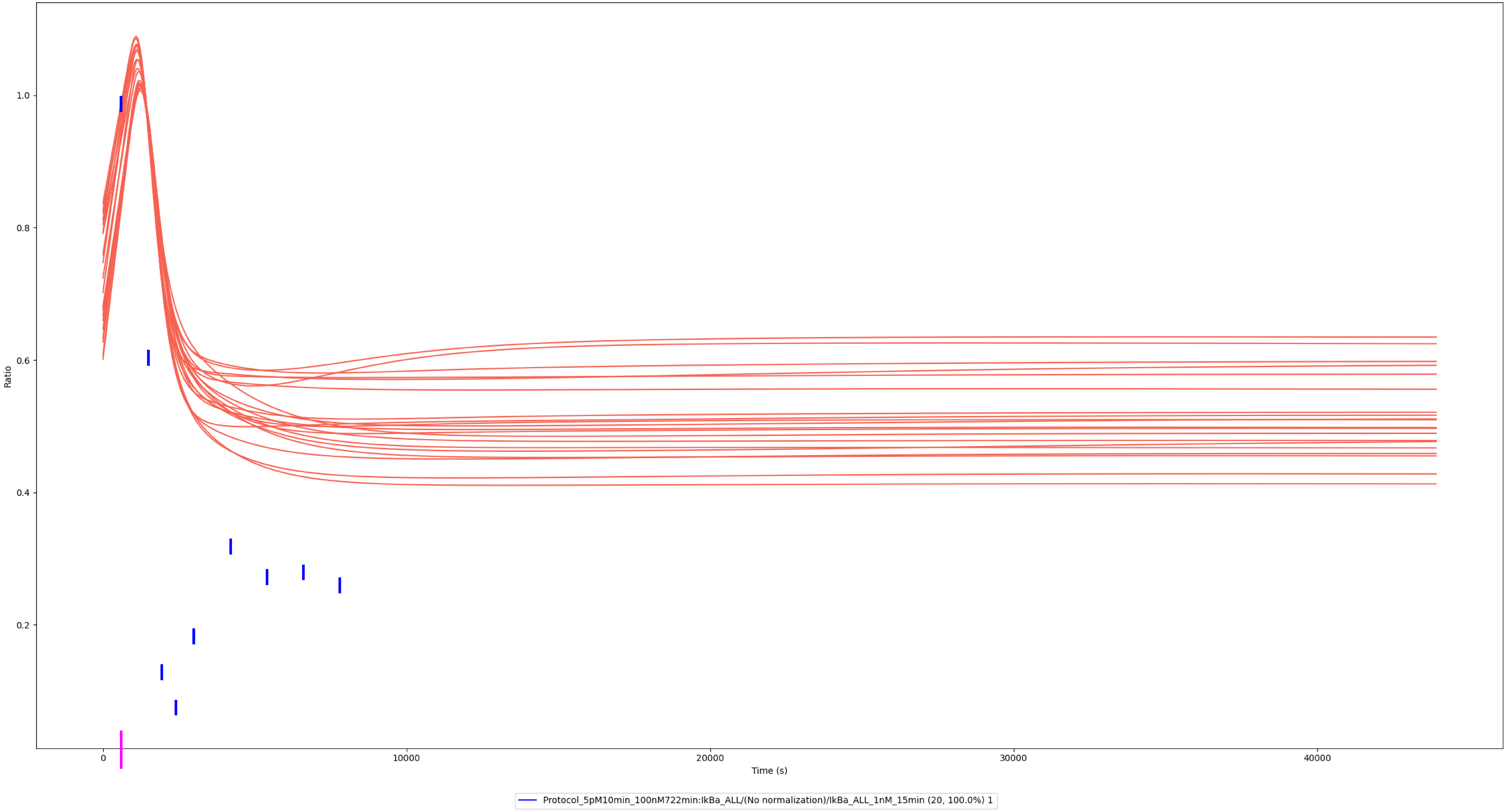

### RelA

### Experiment vs. Simulation: 0.3 nM LPS, Nuclear / Total RelA

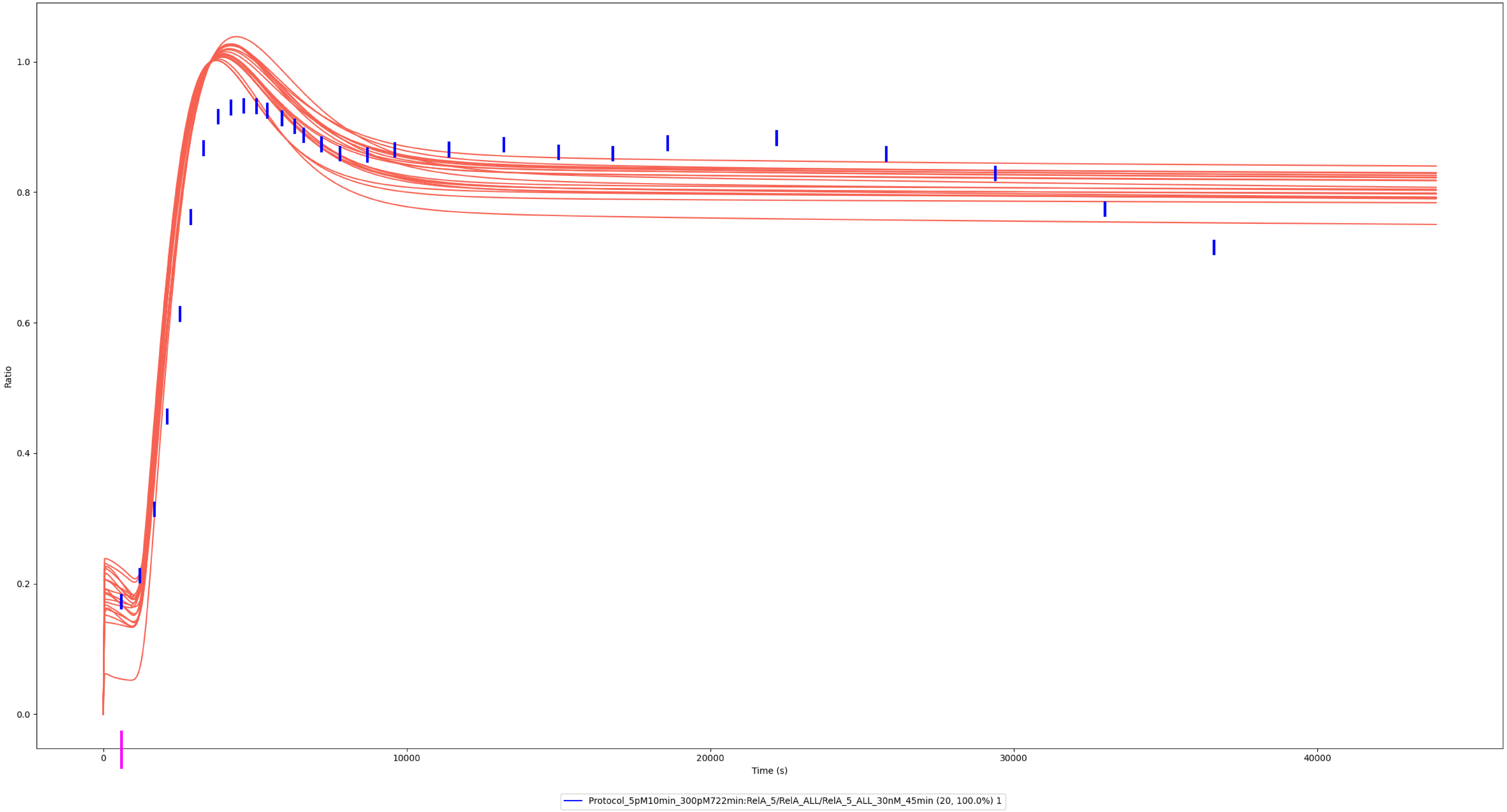

### Experiment vs. Simulation: 3 nM LPS, Nuclear / Total RelA

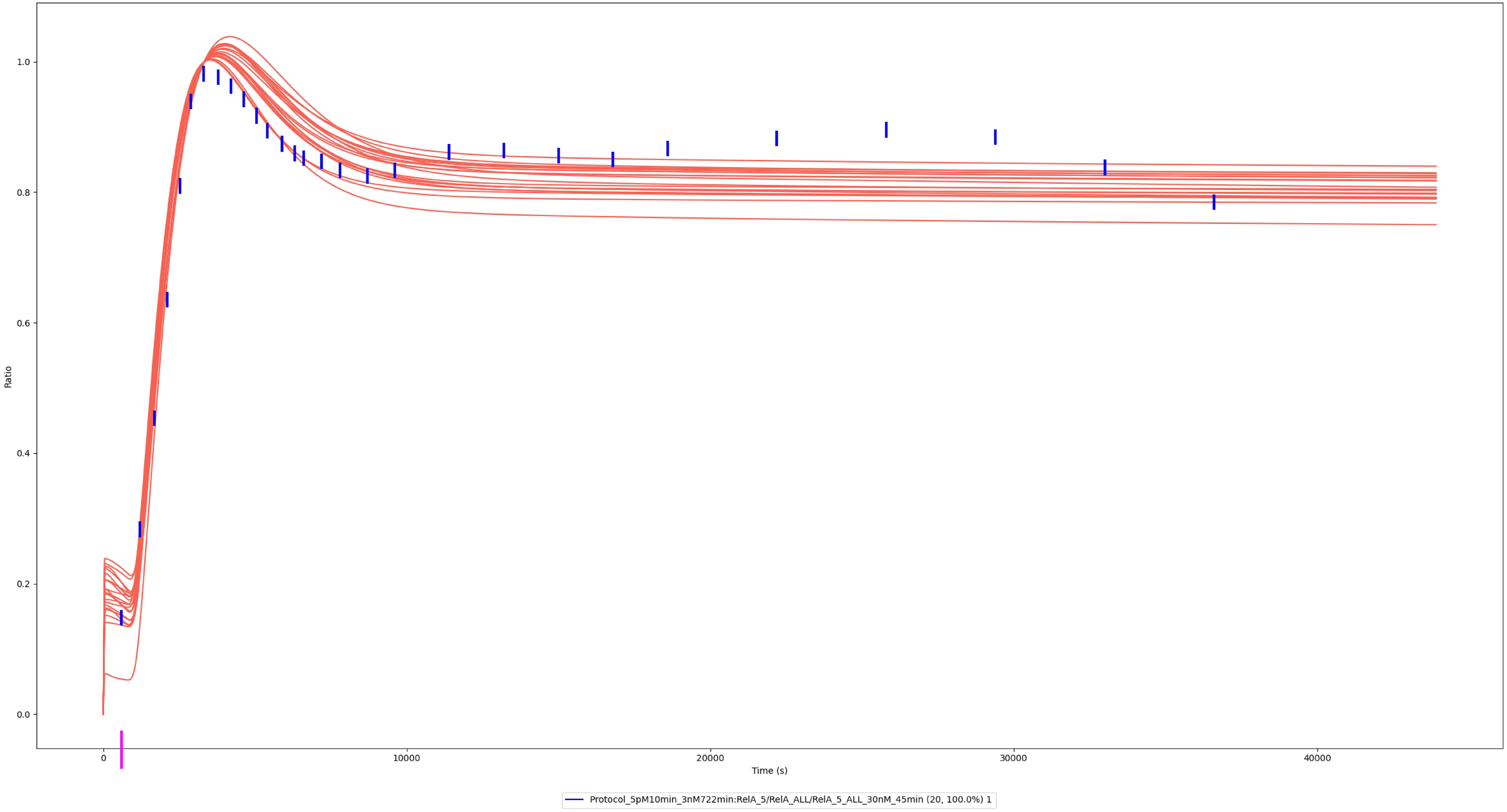

### Experiment vs. Simulation: 30 nM LPS, Nuclear / Total RelA

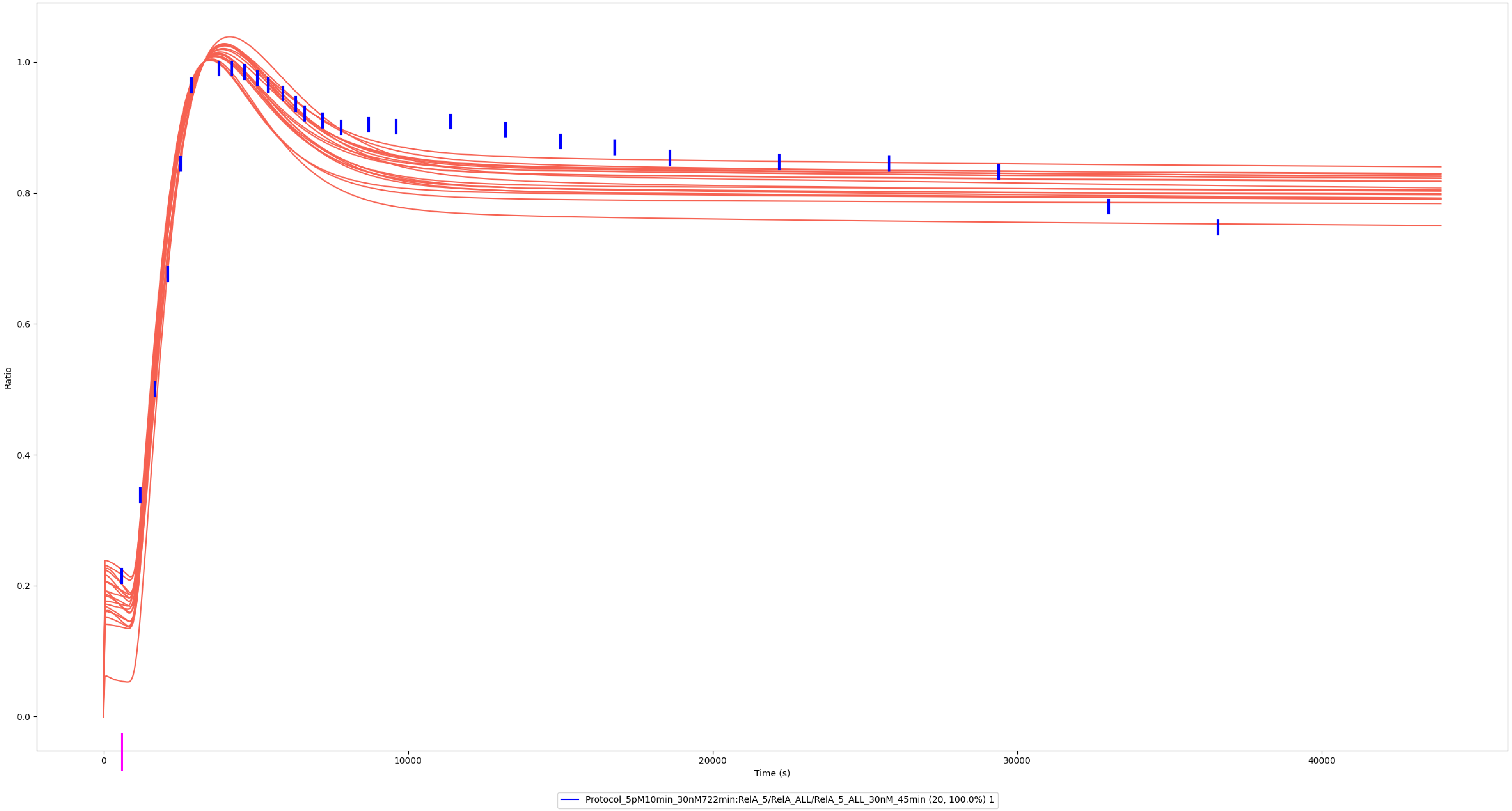

### Experiment vs. Simulation: 150 nM LPS, Nuclear / Total RelA

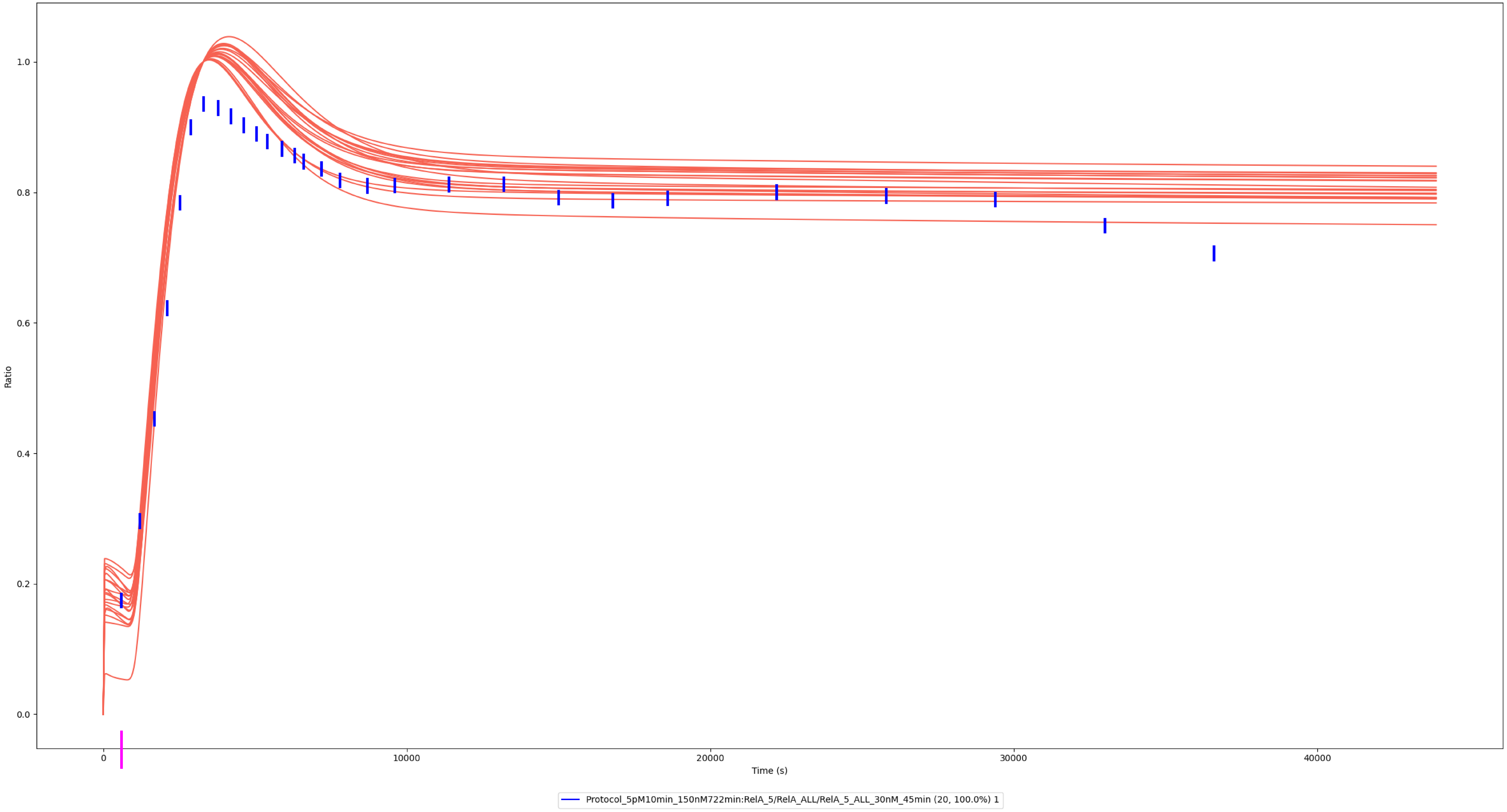

### cRel

### Experiment vs. Simulation: 0.3 nM LPS, Nuclear / Total cRel

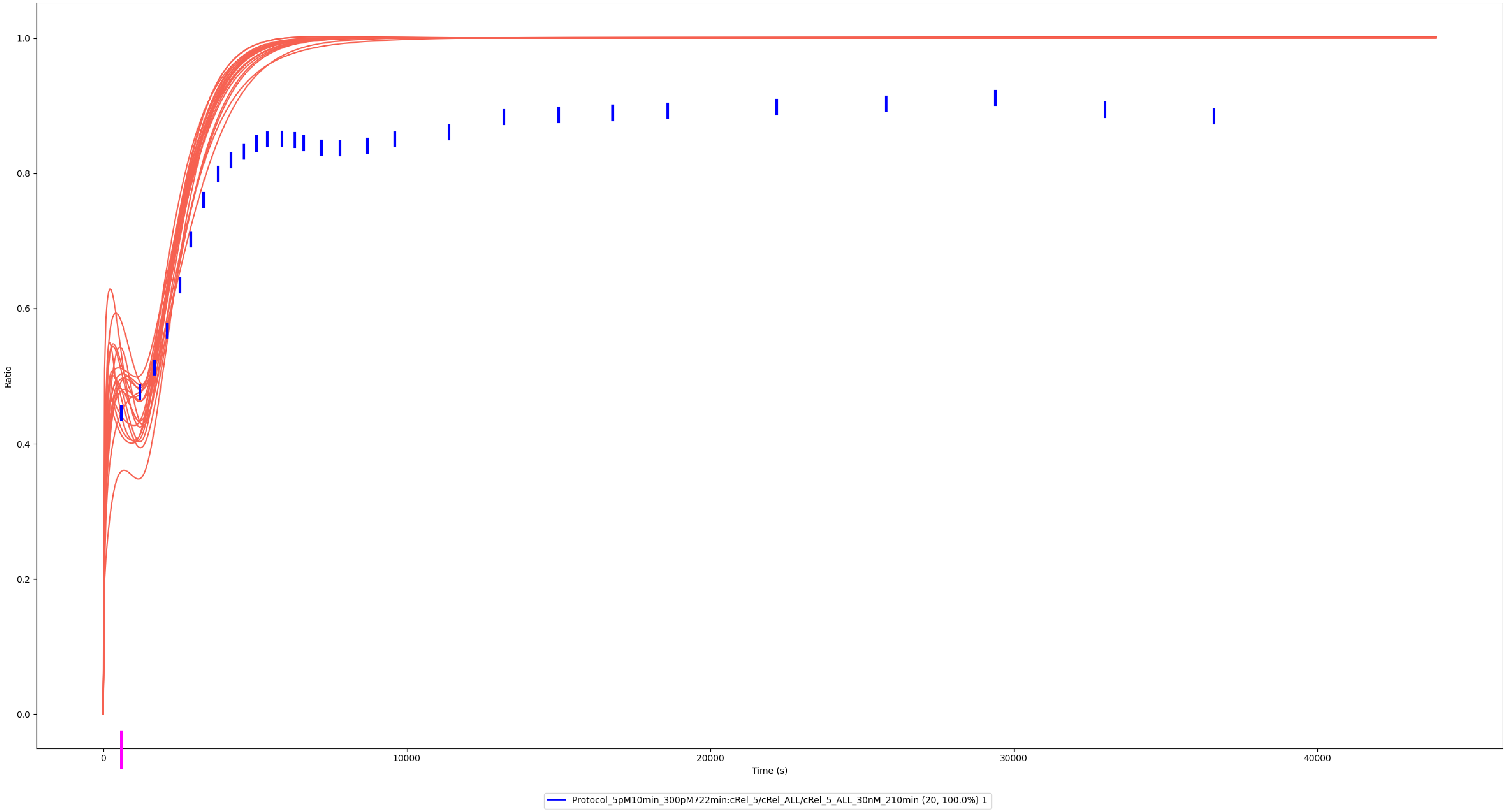

### Experiment vs. Simulation: 3 nM LPS, Nuclear / Total cRel

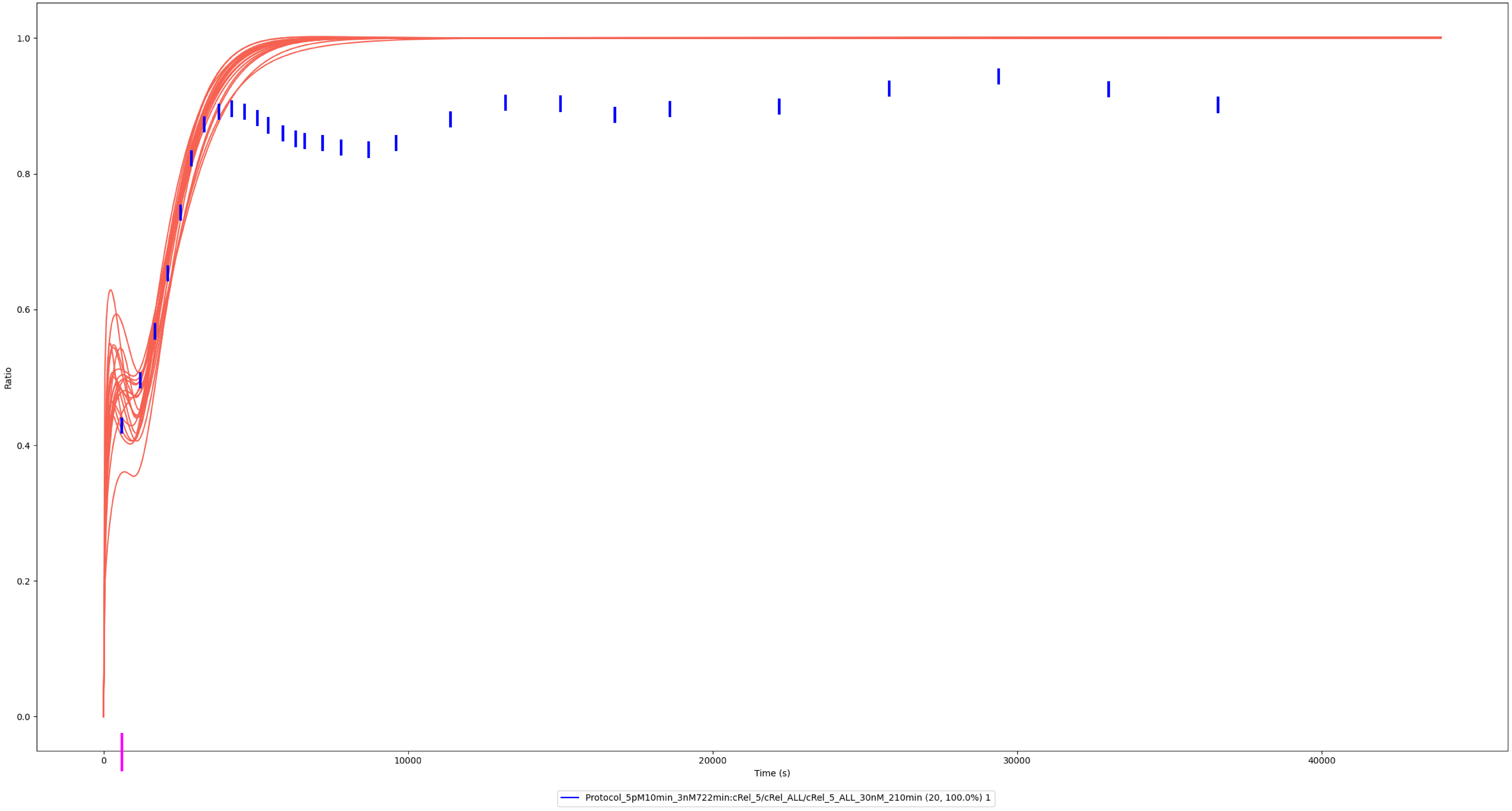

### Experiment vs. Simulation: 30 nM LPS, Nuclear / Total cRel

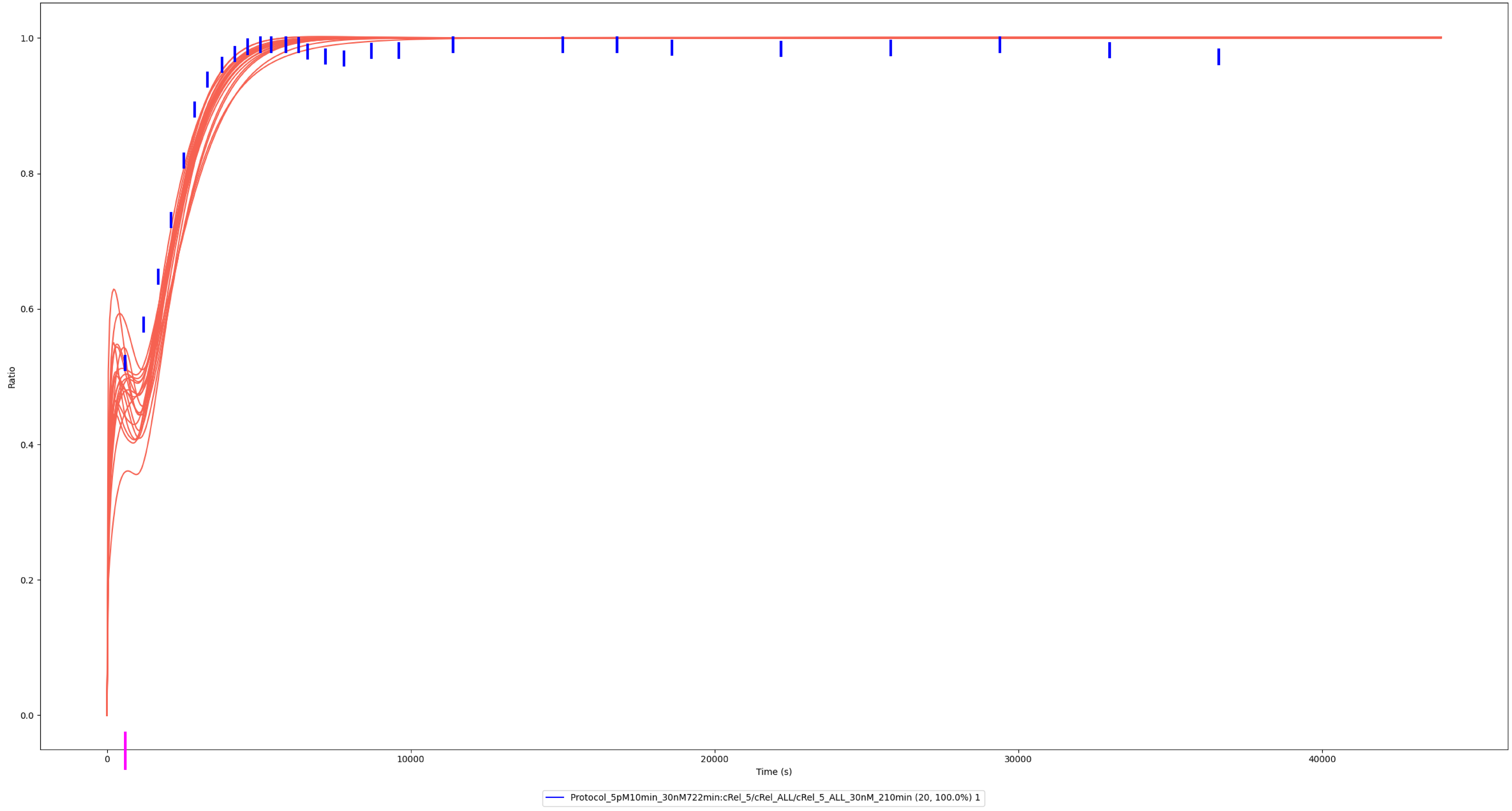

### Experiment vs. Simulation: 150 nM LPS, Nuclear / Total cRel

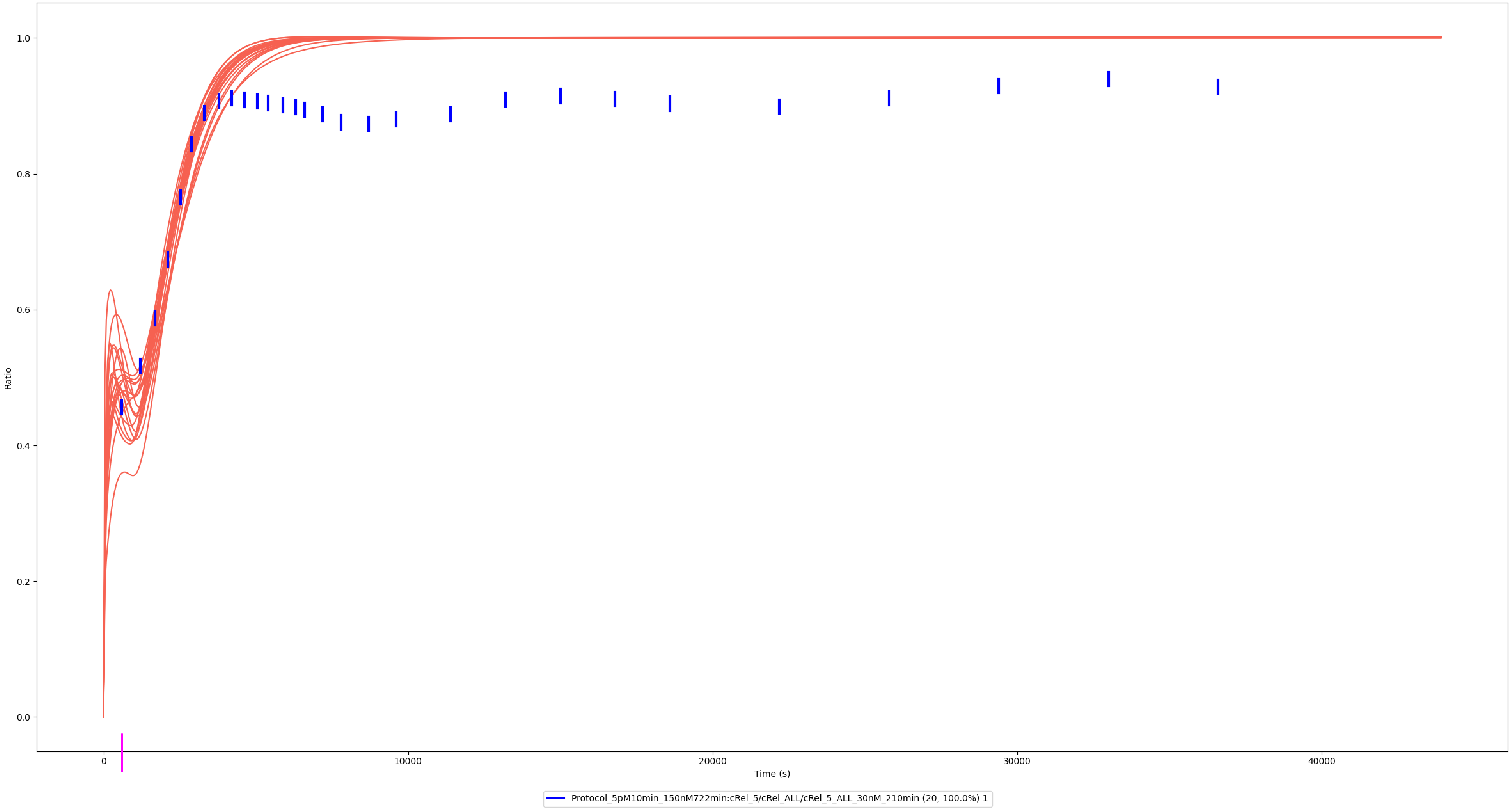

### MKK1/2

### Experiment vs. Simulation: 1 nM LPS, pp-MKK1/2

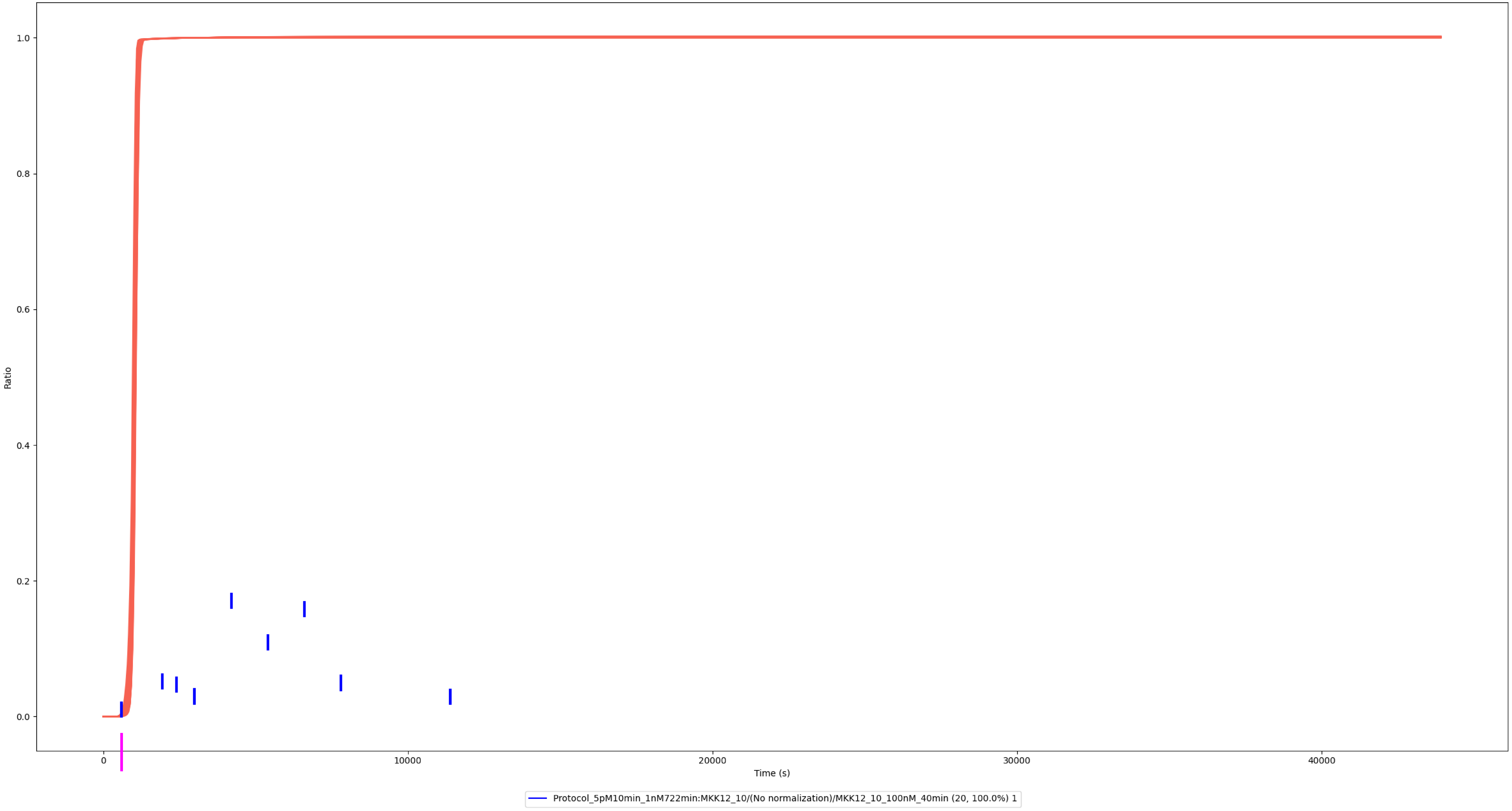

### Experiment vs. Simulation: 10 nM LPS, pp-MKK1/2

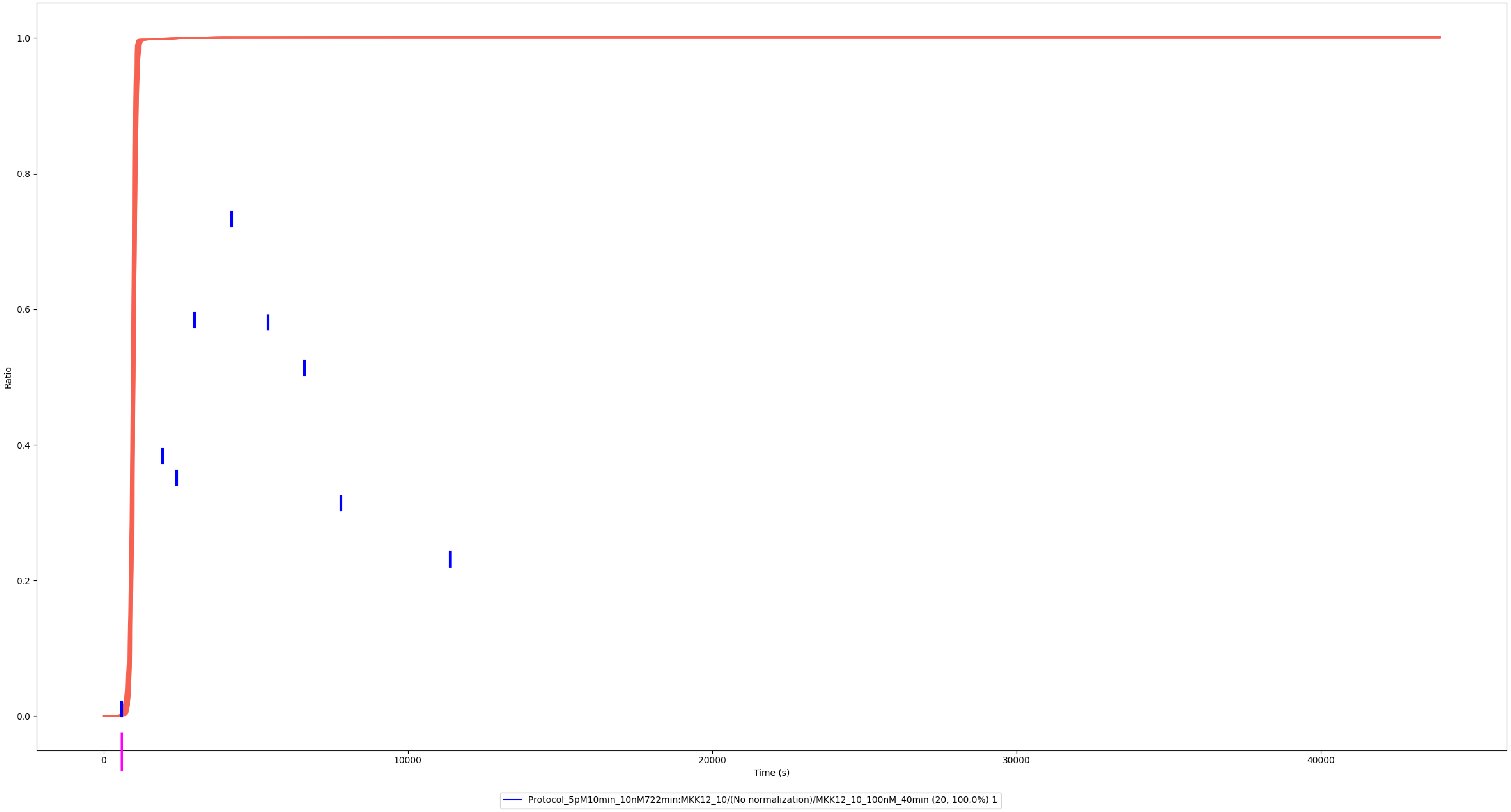

### Experiment vs. Simulation: 100 nM LPS, pp-MKK1/2

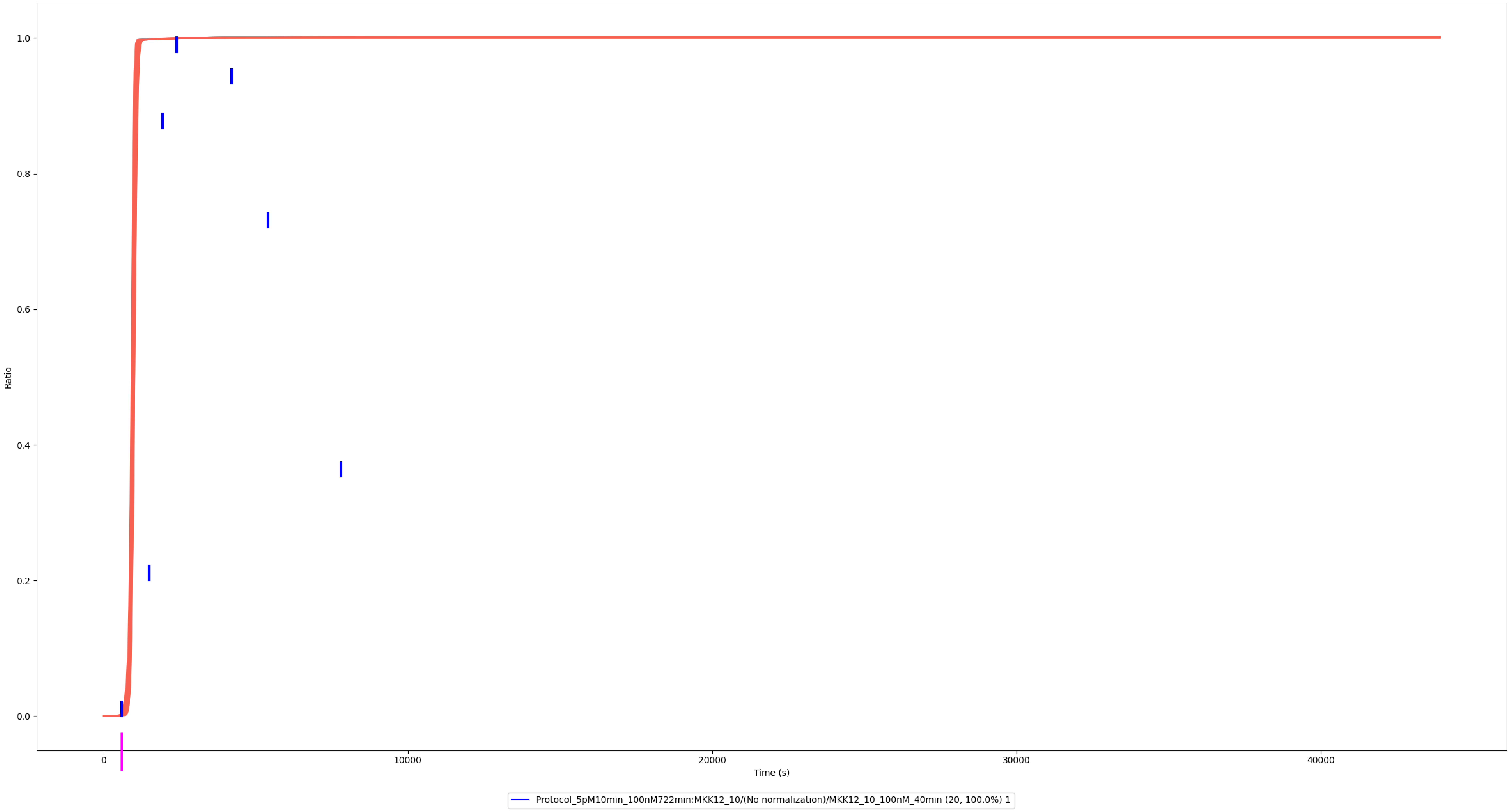

### MKK4

### Experiment vs. Simulation: 30 nM LPS, pp-MKK4

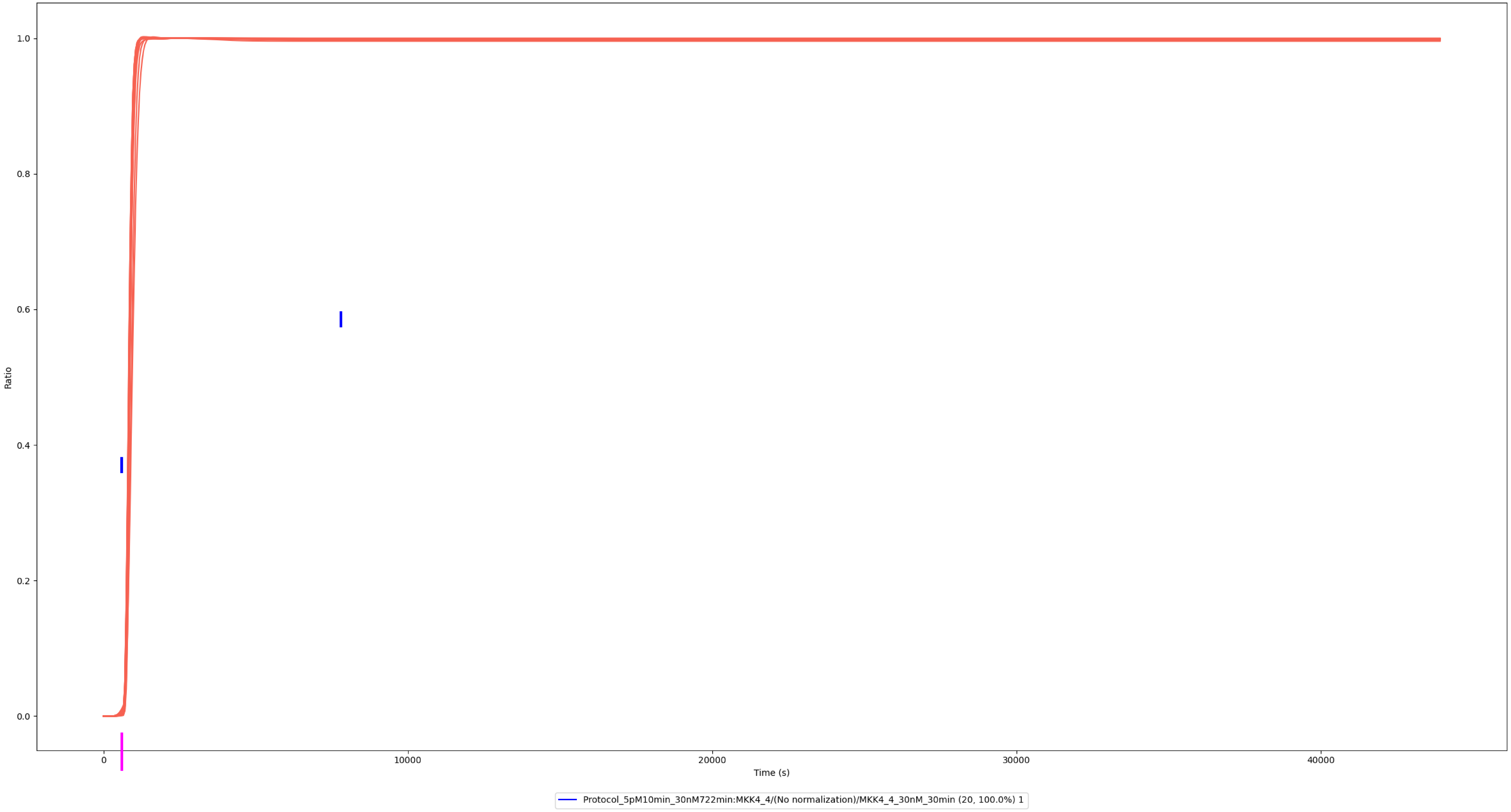

### ERK1/2

### Experiment vs. Simulation: 0.01 nM LPS, pp-ERK1/2

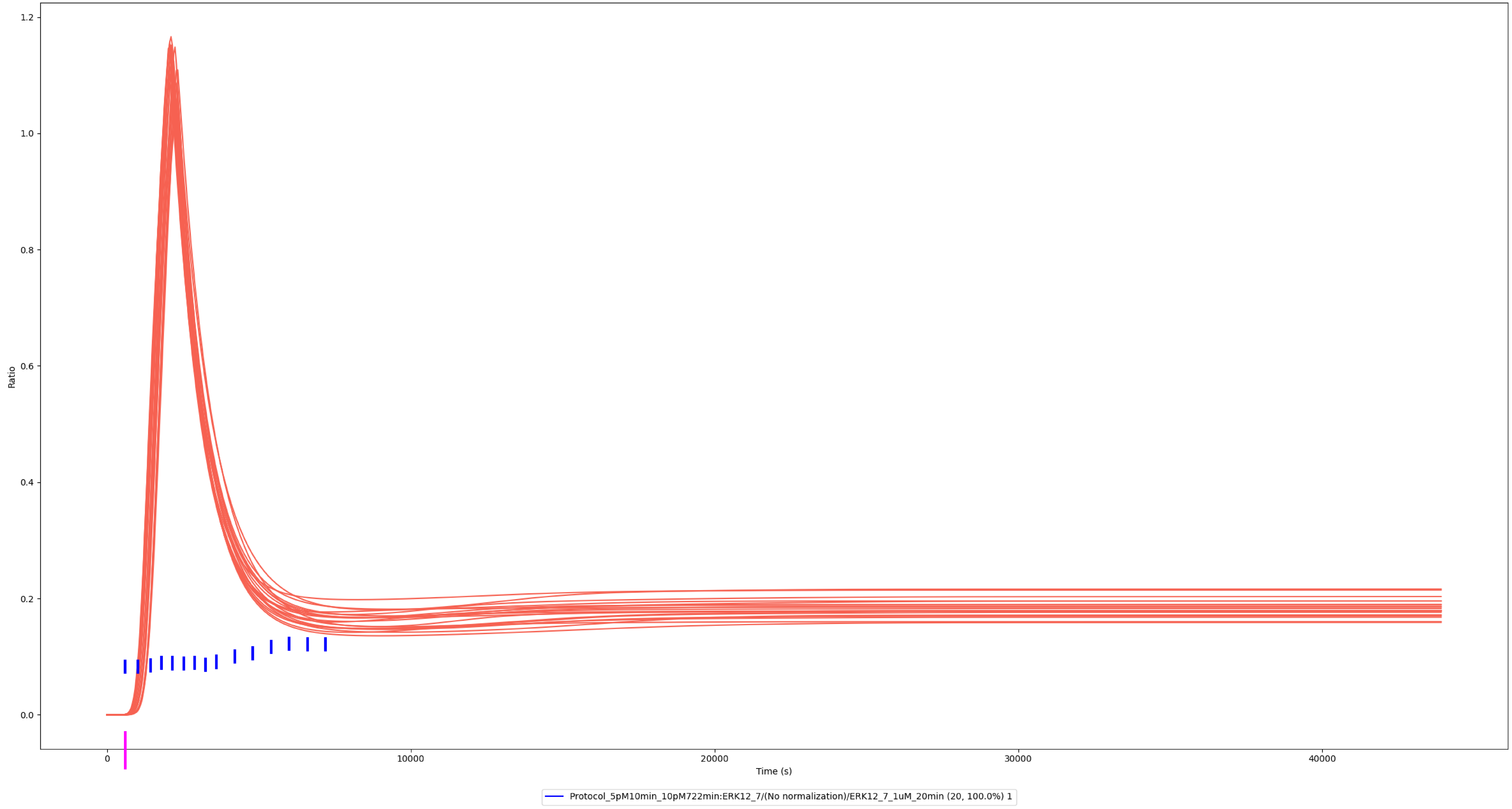

### Experiment vs. Simulation: 0.1 nM LPS, pp-ERK1/2

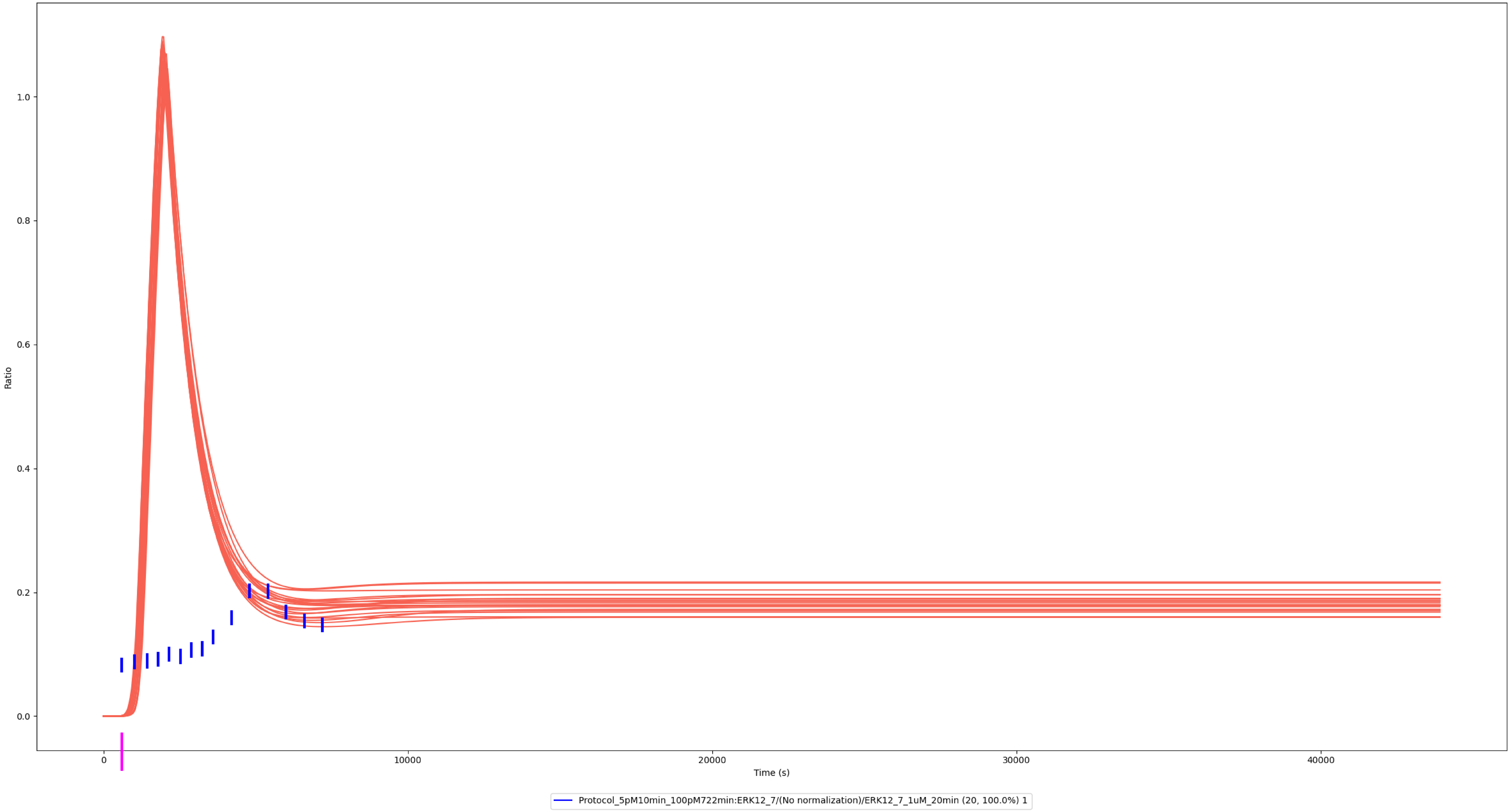

### Experiment vs. Simulation: 1 nM LPS, pp-ERK1/2

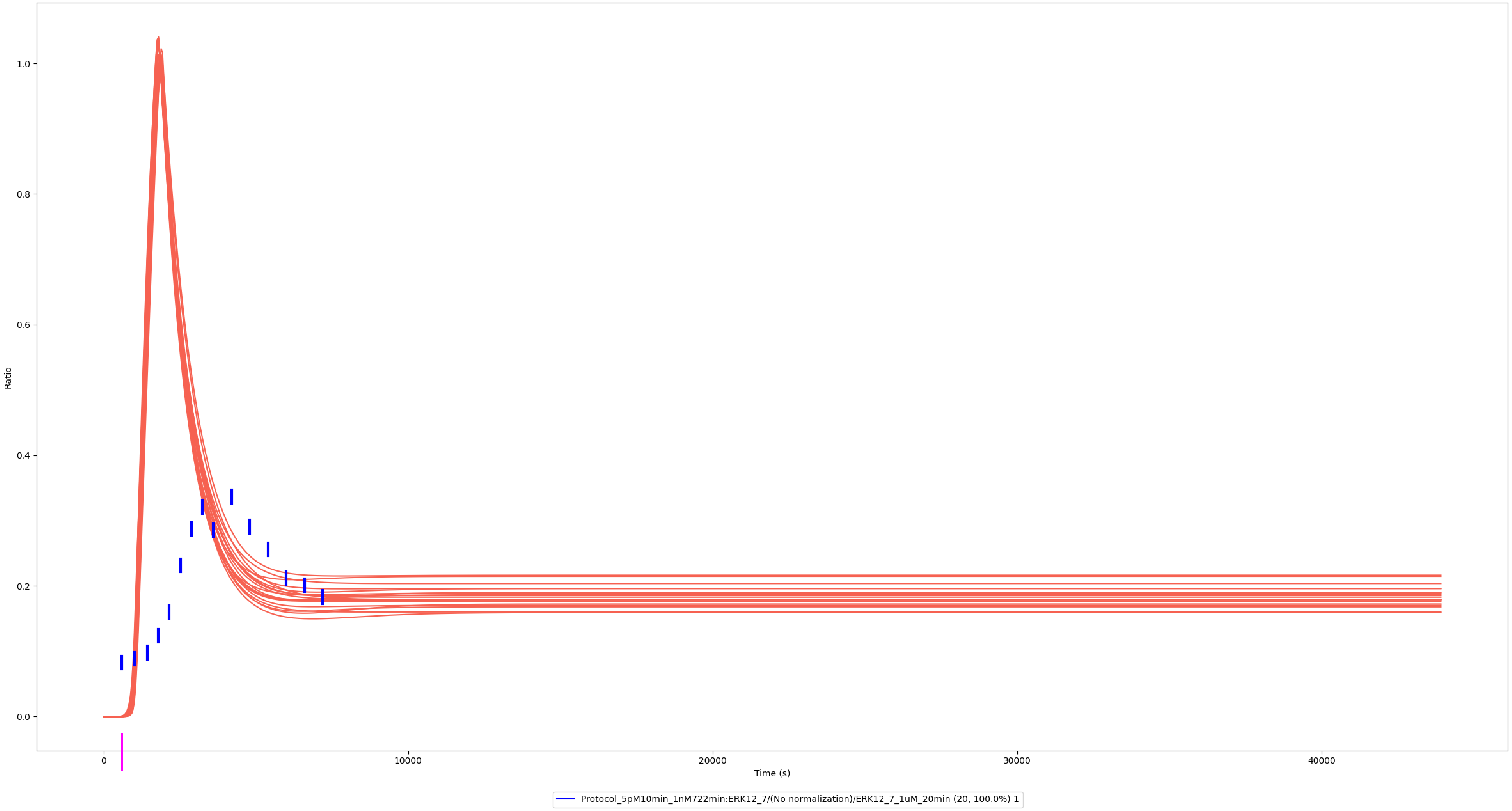

### Experiment vs. Simulation: 10 nM LPS, pp-ERK1/2

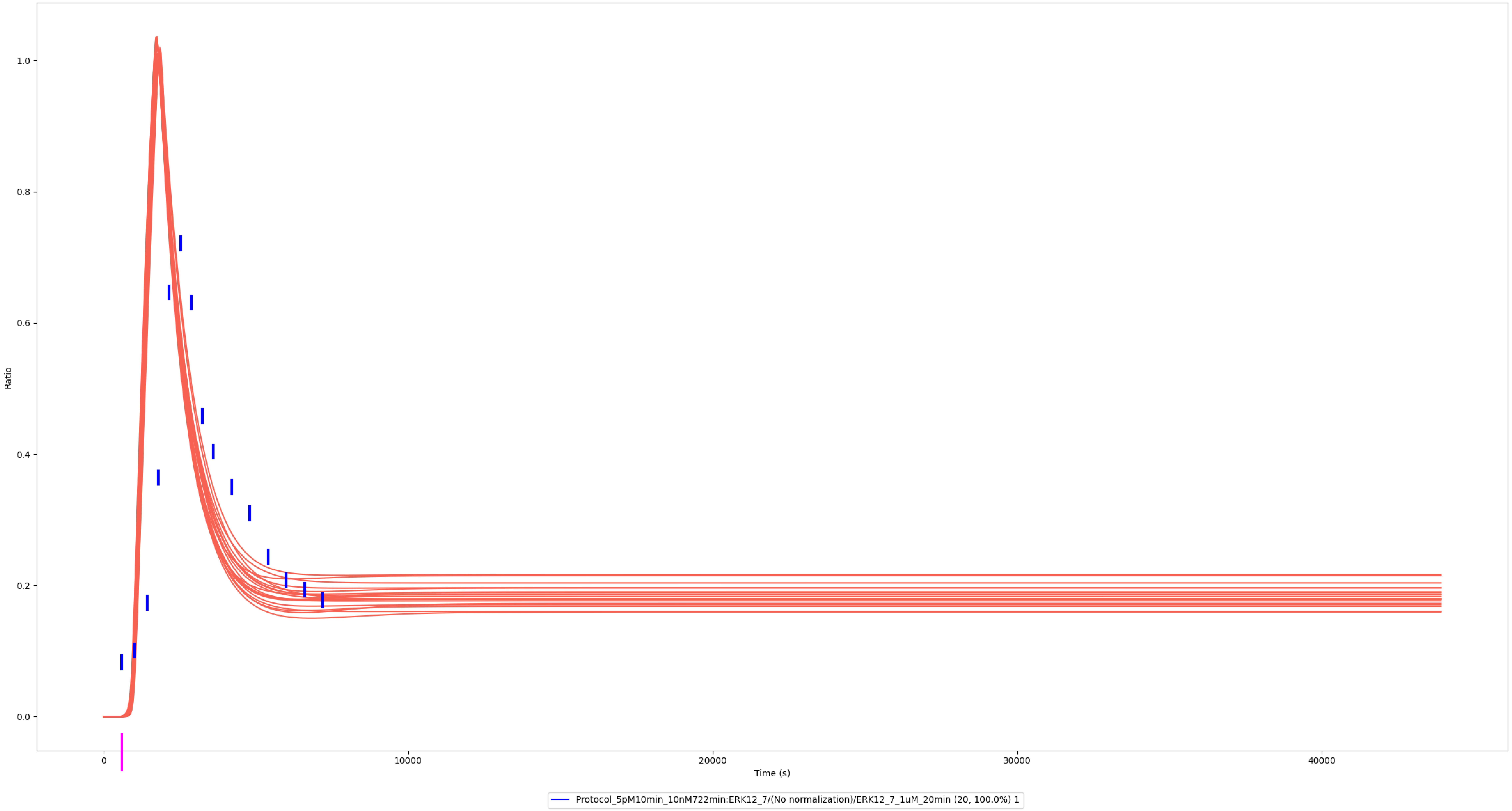

### Experiment vs. Simulation: 100 nM LPS, pp-ERK1/2

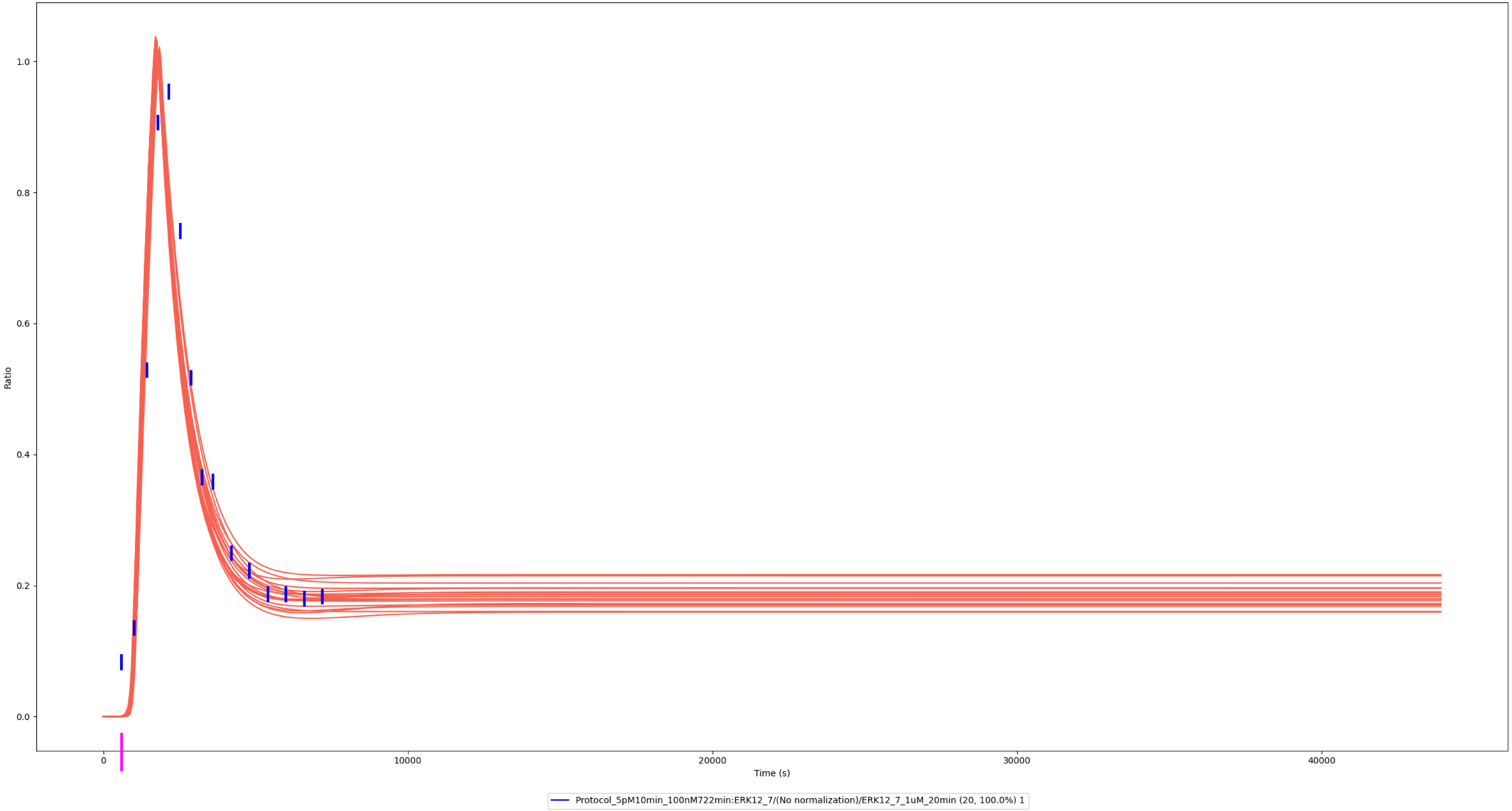

### Experiment vs. Simulation: 1,000 nM LPS, pp-ERK1/2

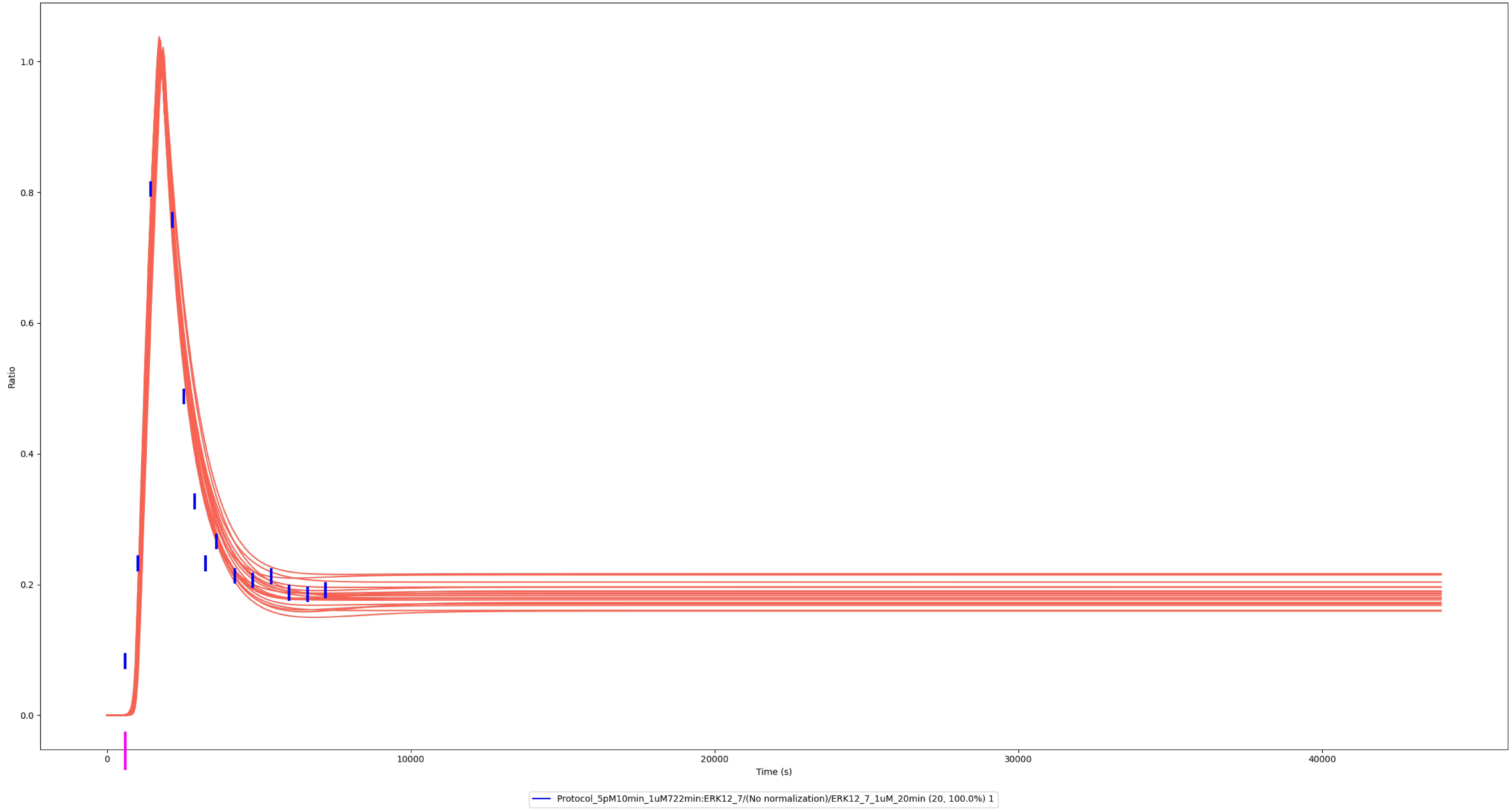

### Experiment vs. Simulation: 1 nM LPS, pp-ERK1/2

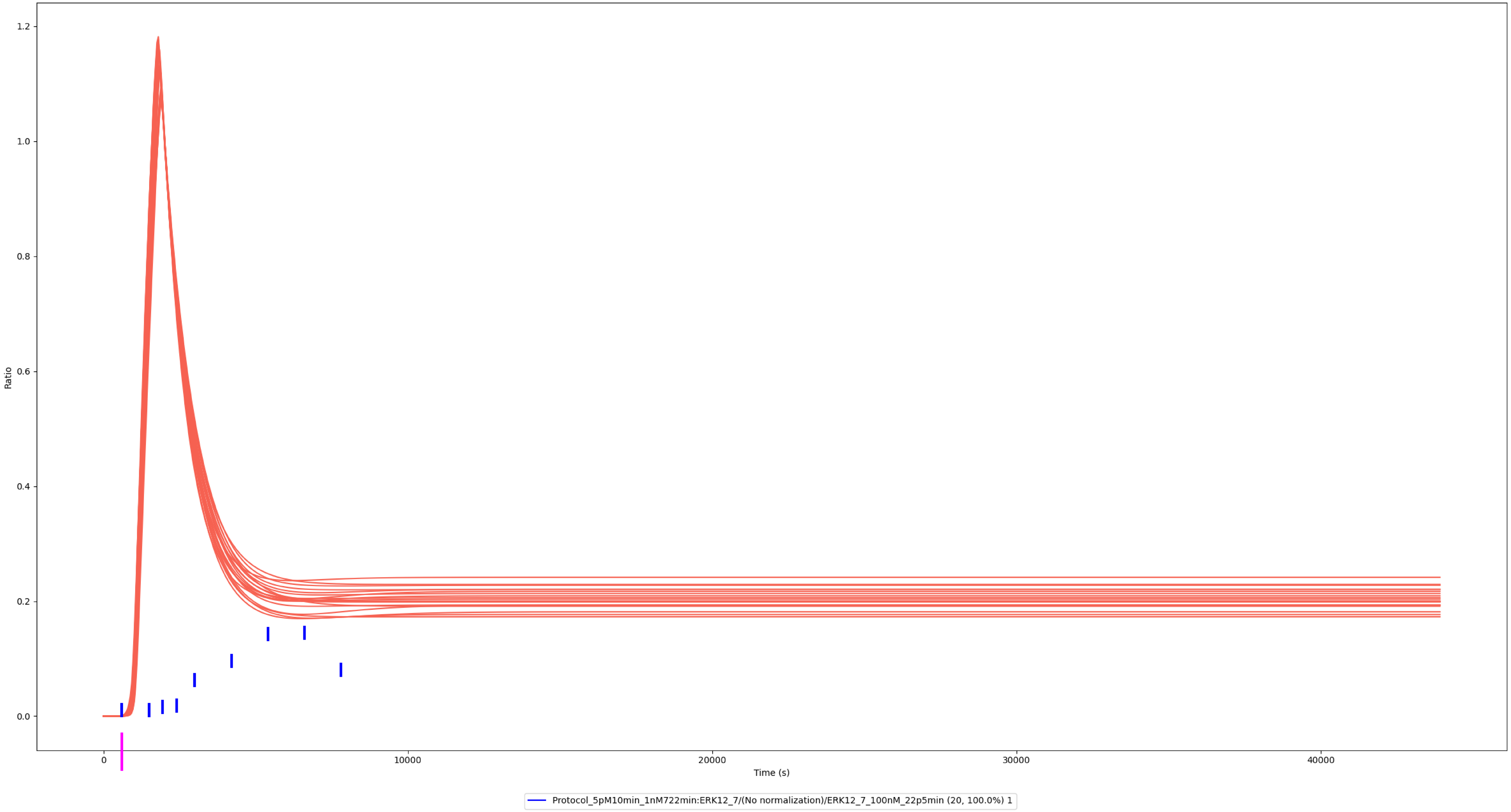

### Experiment vs. Simulation: 100 nM LPS, pp-ERK1/2

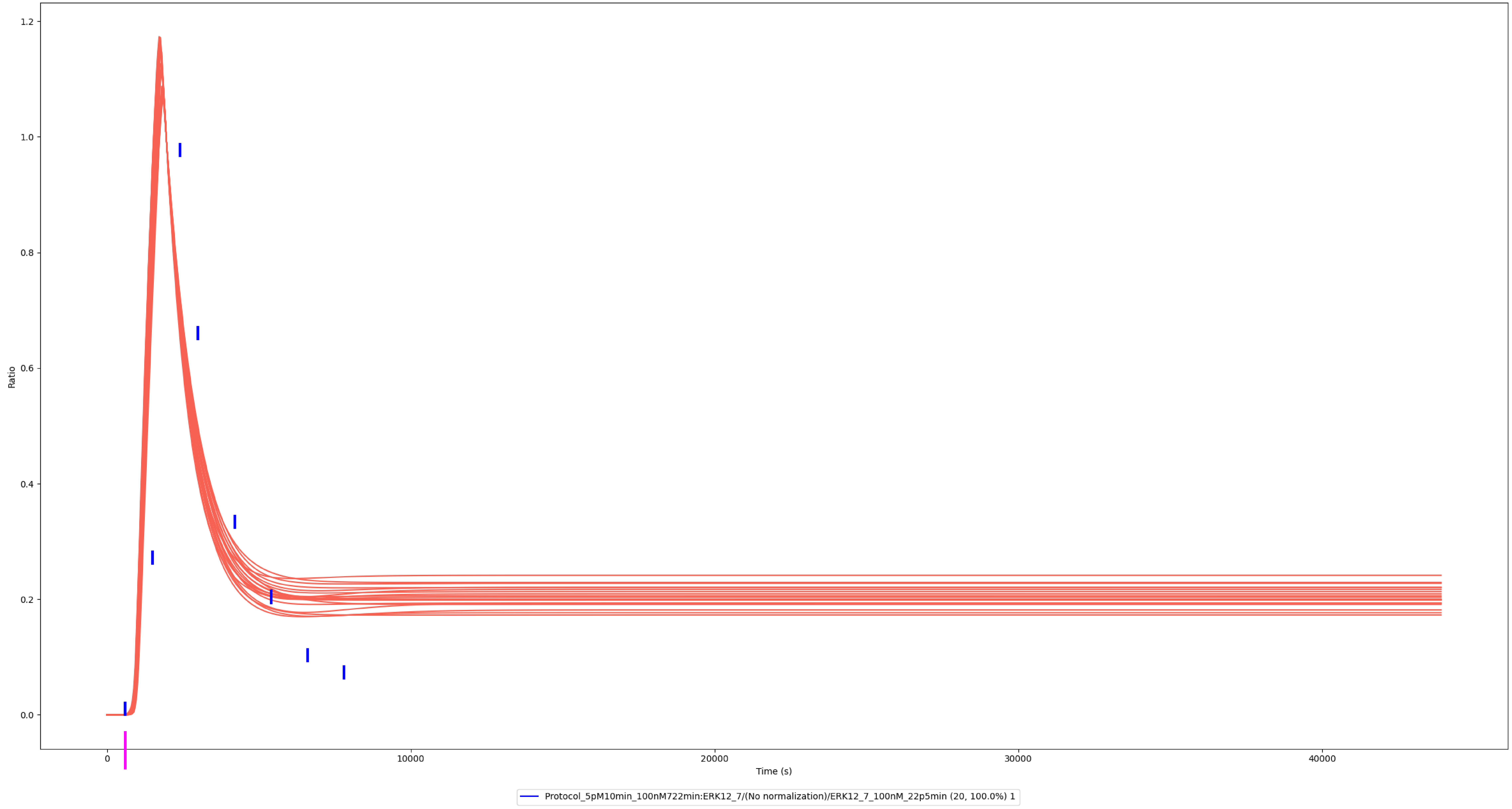

### Experiment vs. Simulation: 1 nM LPS, pp-ERK1/2

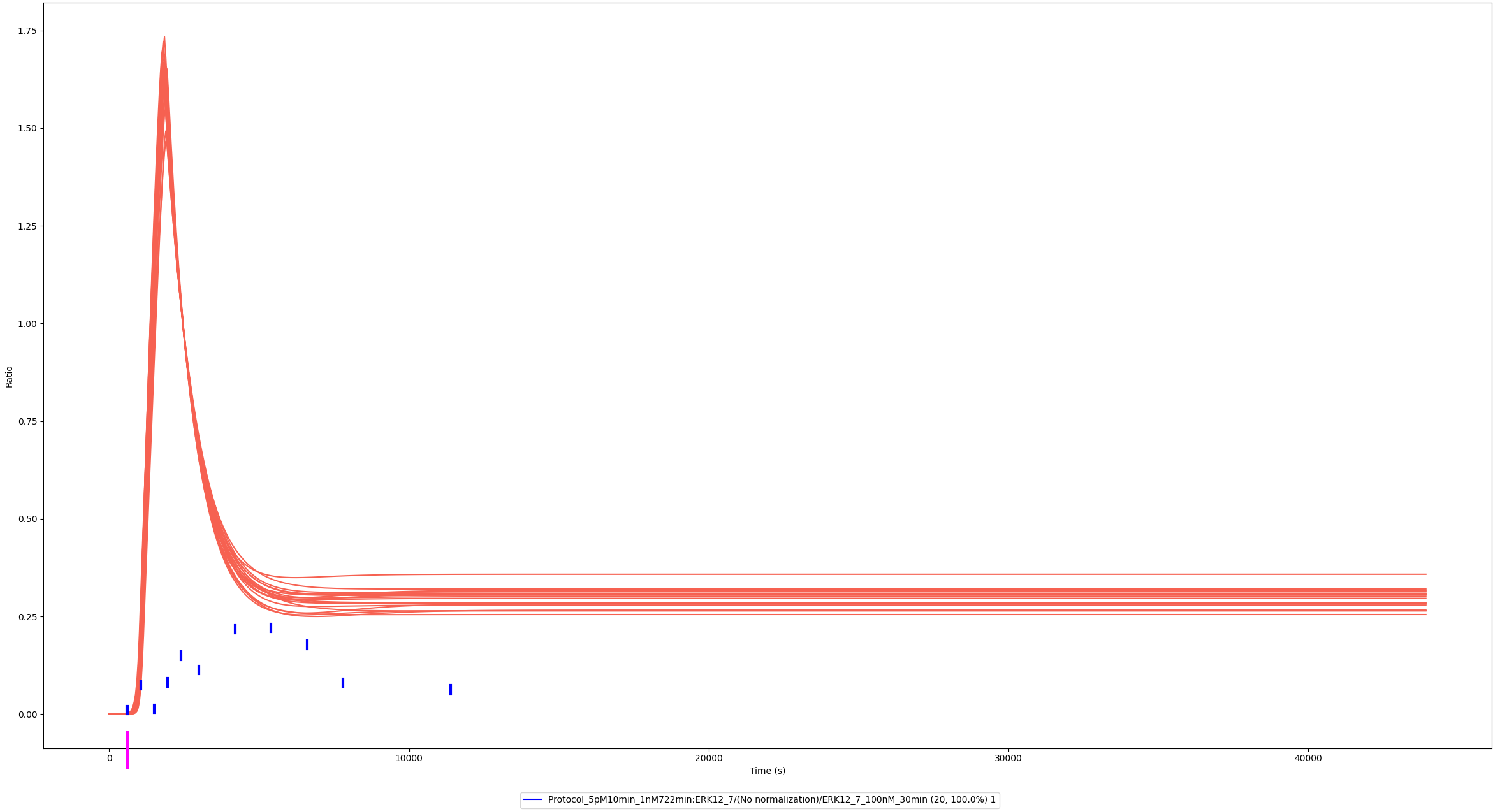

### Experiment vs. Simulation: 10 nM LPS, pp-ERK1/2

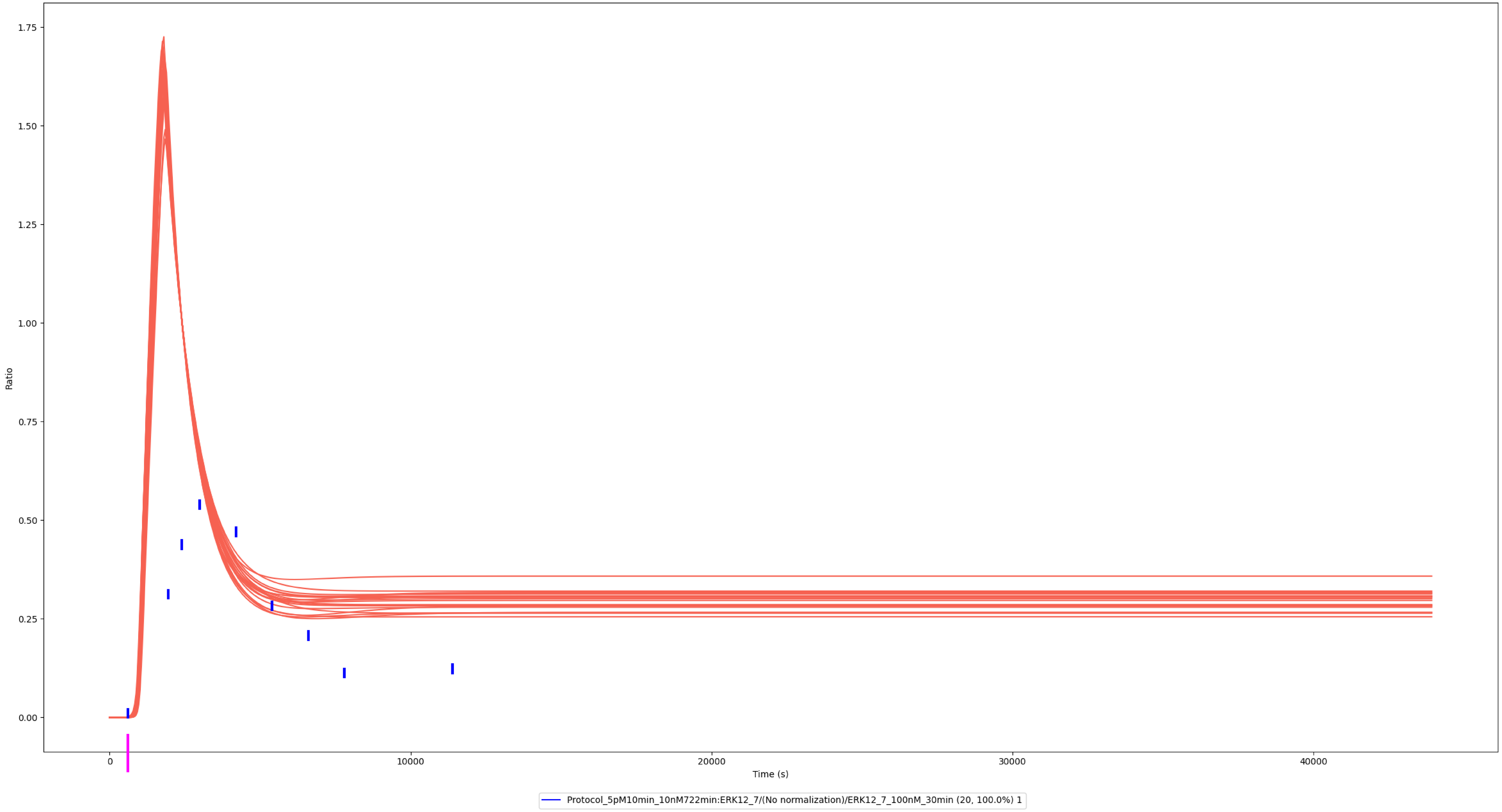

### Experiment vs. Simulation: 100 nM LPS, pp-ERK1/2

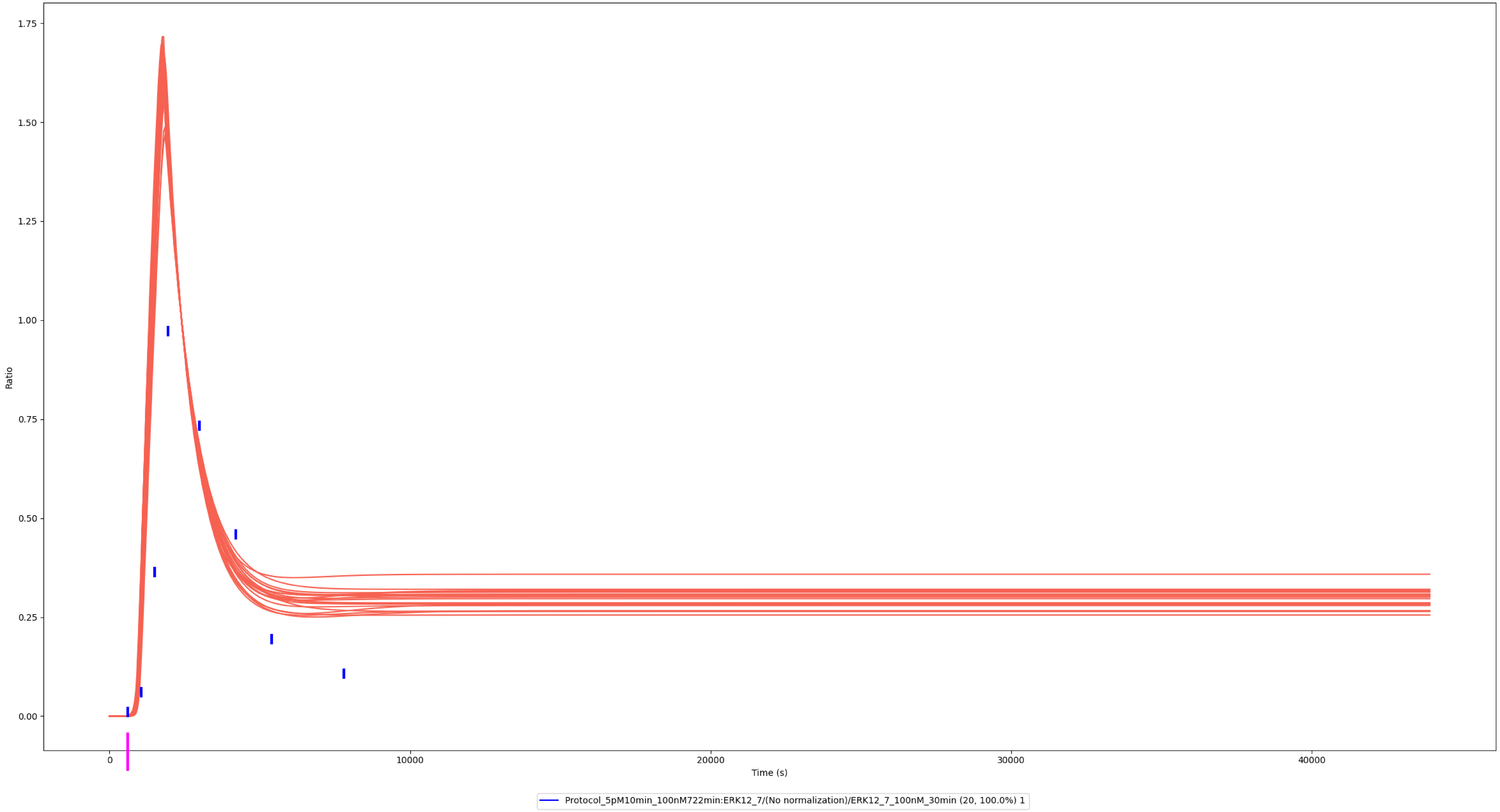

### Experiment vs. Simulation: 3 nM LPS, pp-ERK1

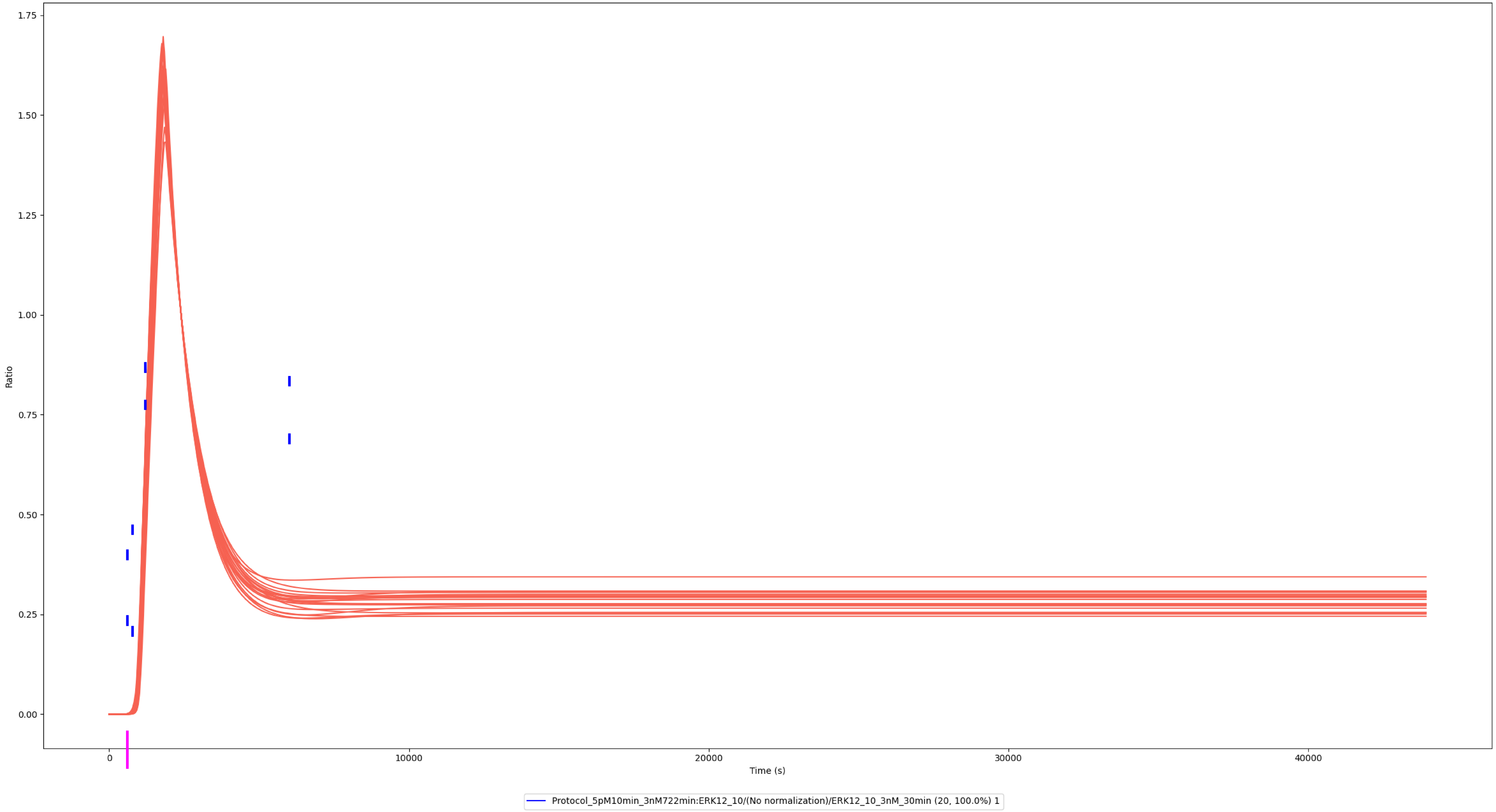

### Experiment vs. Simulation: 30 nM LPS, pp-ERK1

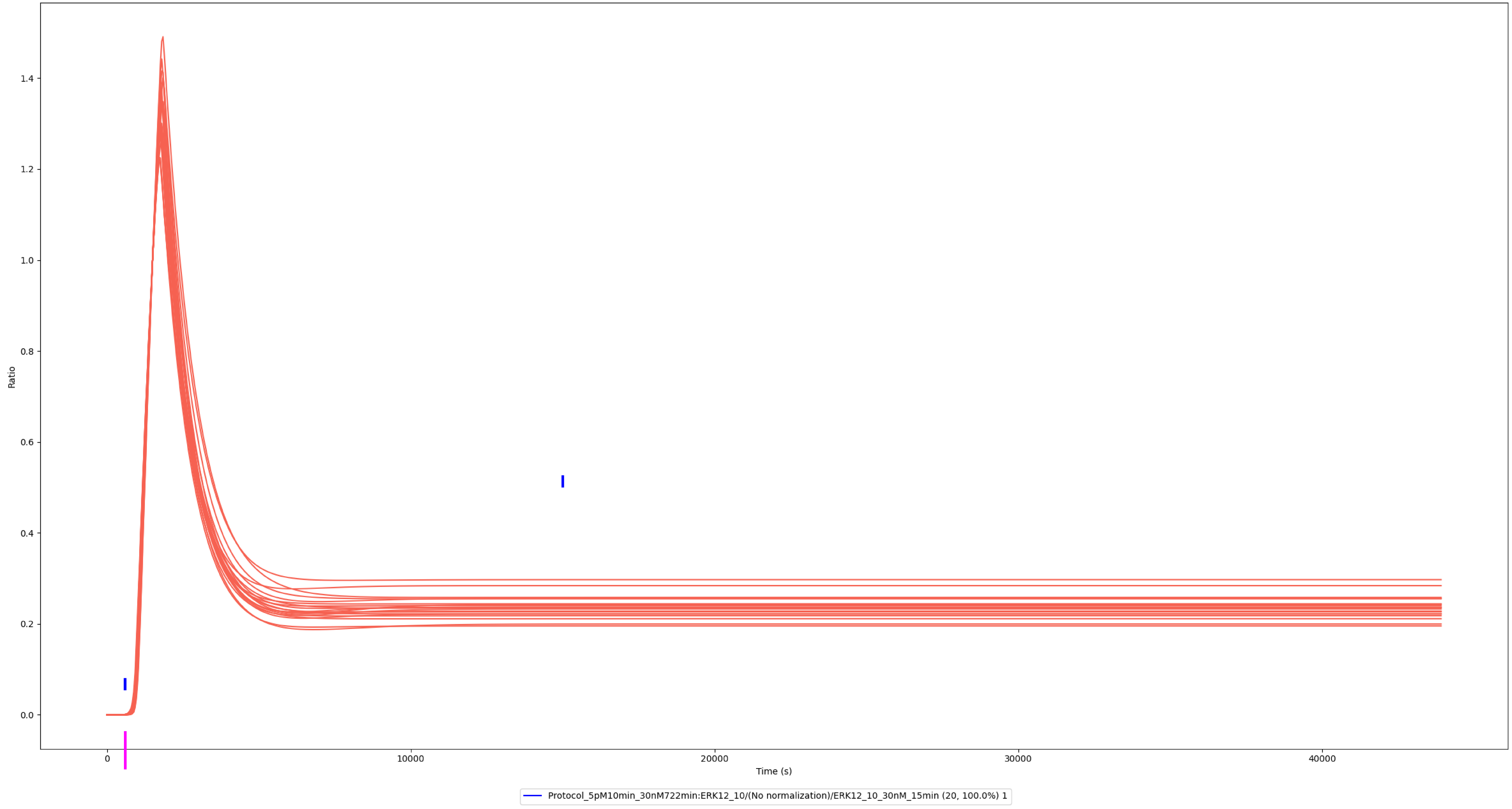

### Experiment vs. Simulation: 30 nM LPS, pp-ERK1

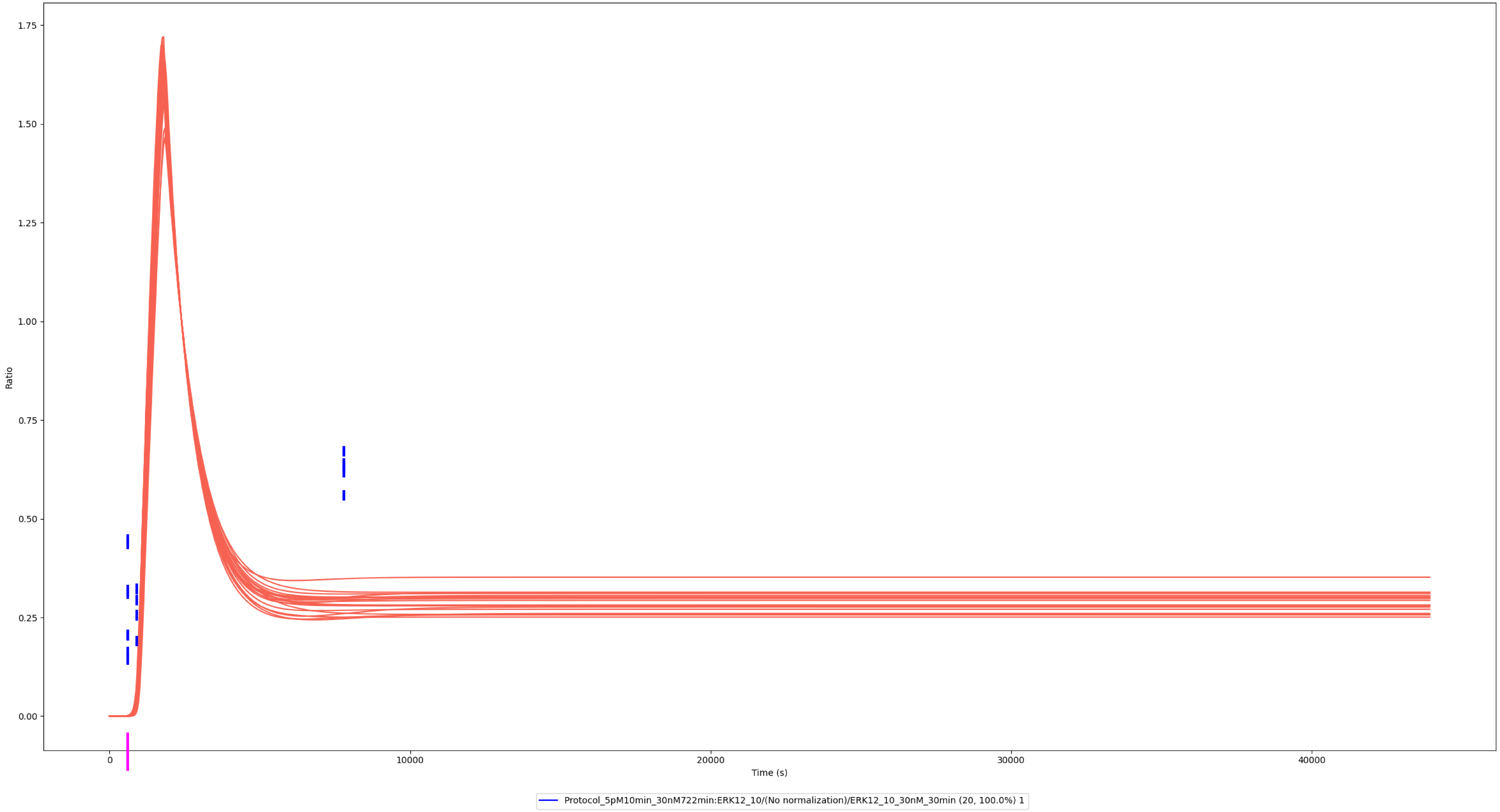

### Experiment vs. Simulation: 300 nM LPS, pp-ERK1

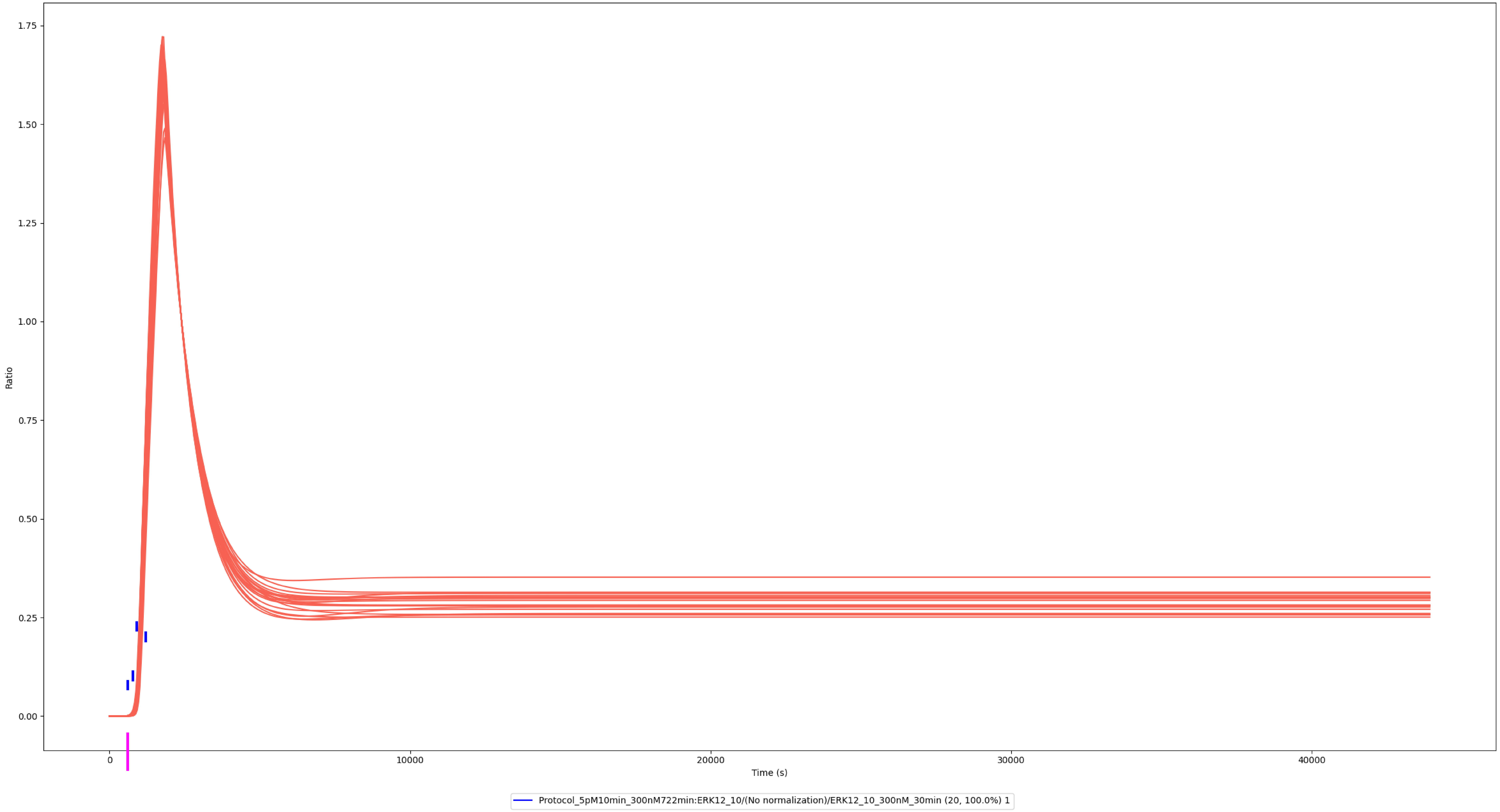

### Experiment vs. Simulation: 3 nM LPS, pp-ERK2

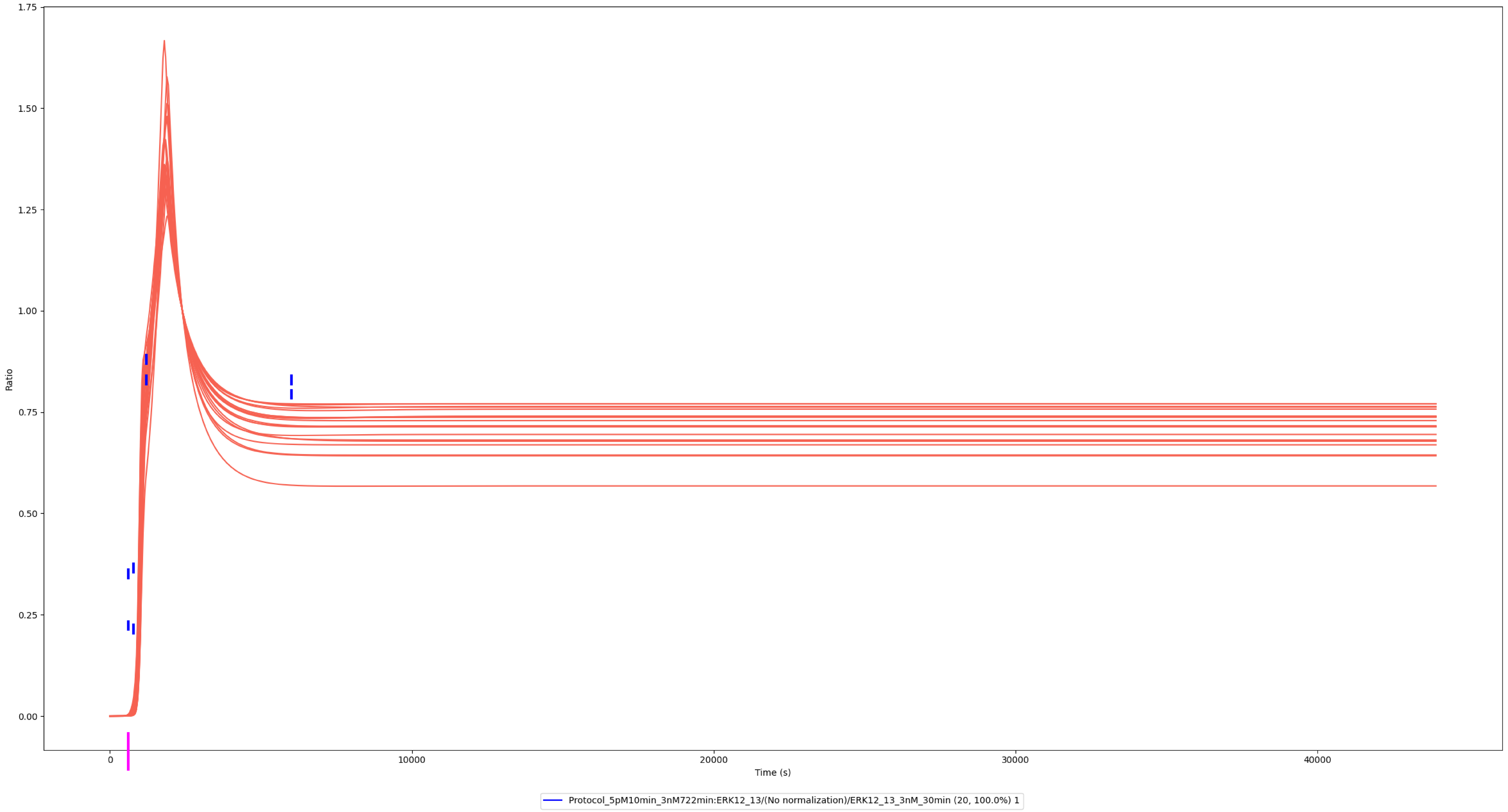

### Experiment vs. Simulation: 30 nM LPS, pp-ERK2

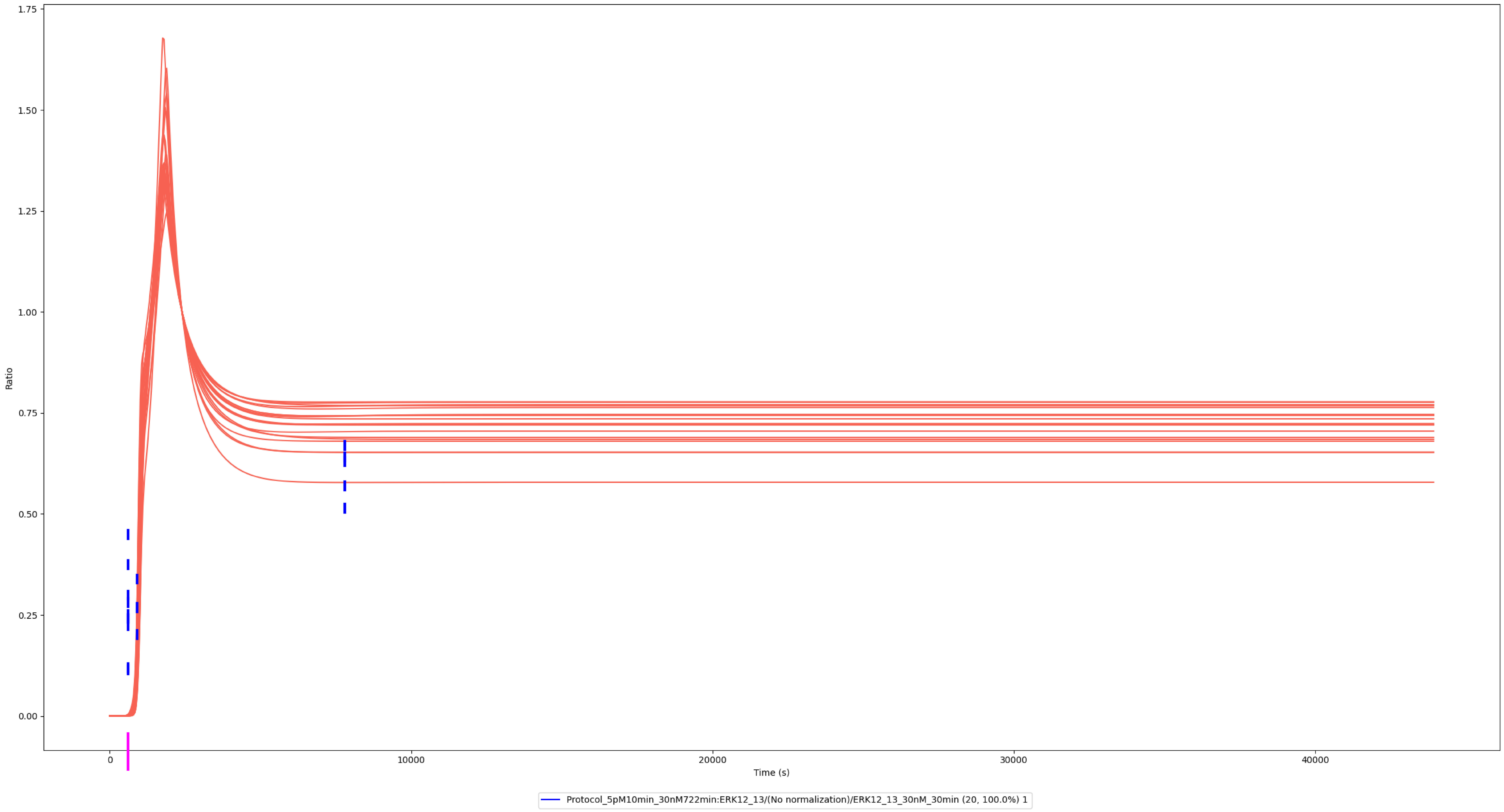

### JNK1/2/3

### Experiment vs. Simulation: 3 nM LPS, pp-JNK1/3

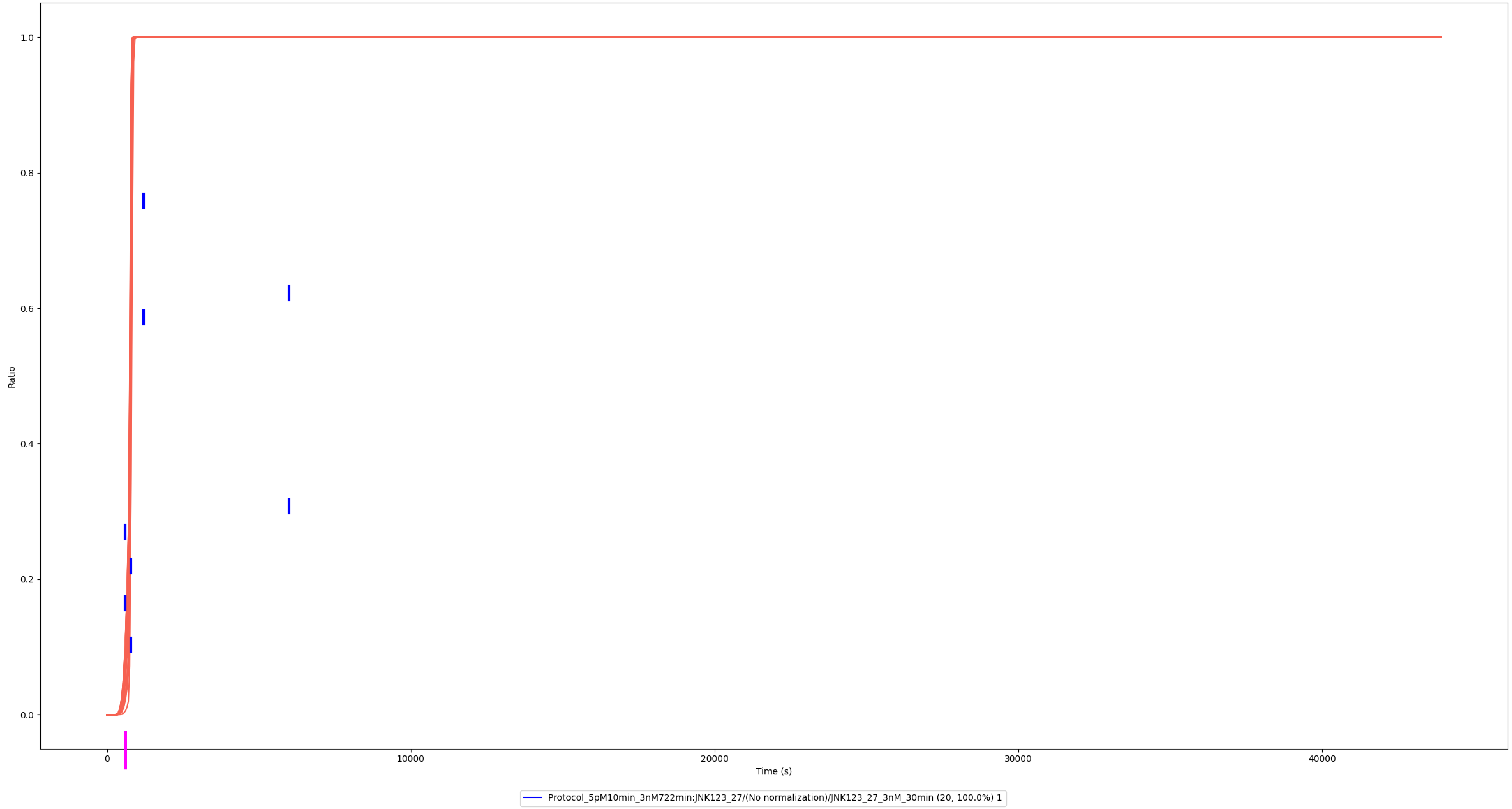

### Experiment vs. Simulation: 30 nM LPS, pp-JNK1/3

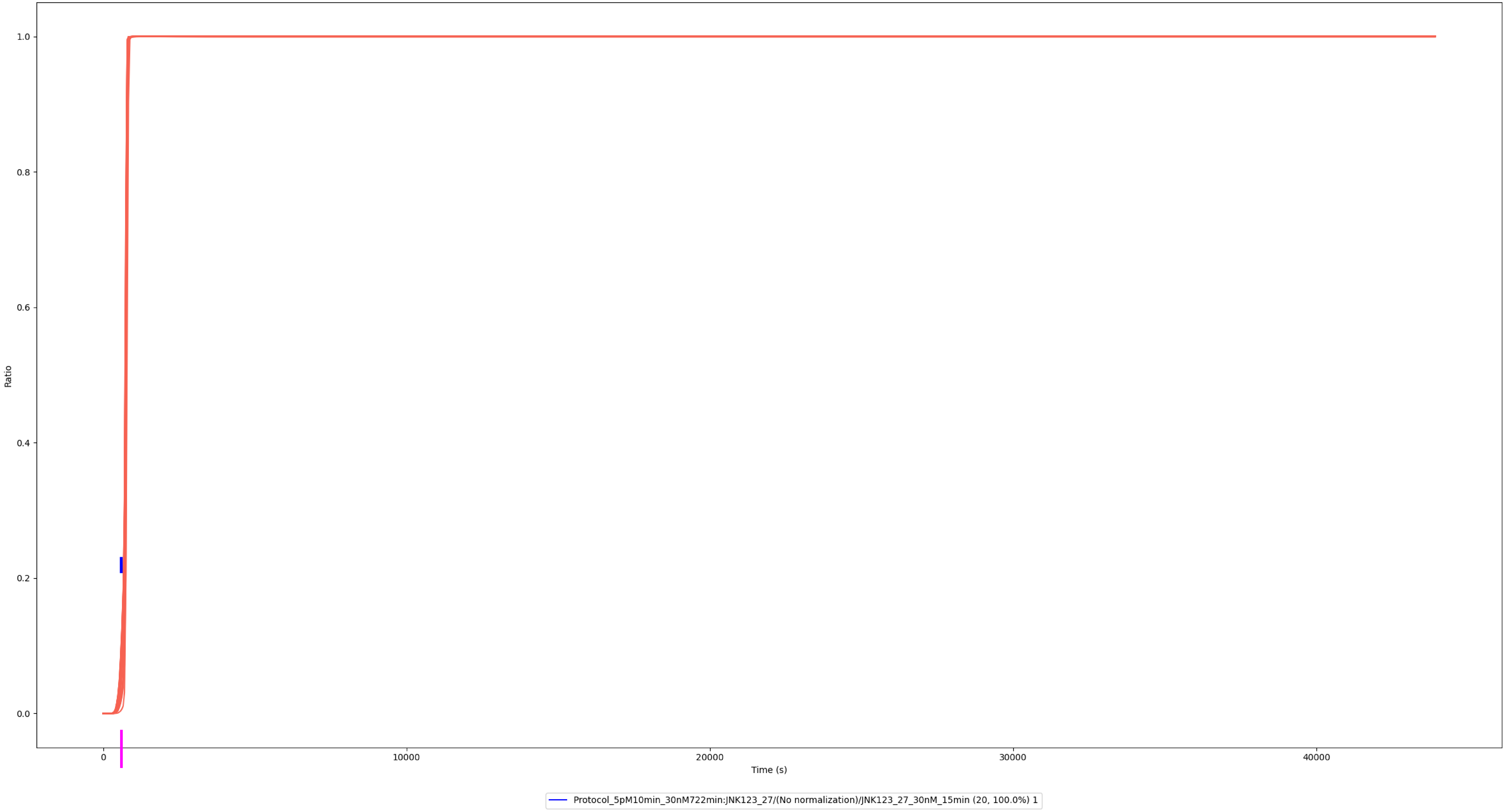

### Experiment vs. Simulation: 30 nM LPS, pp-JNK1/3

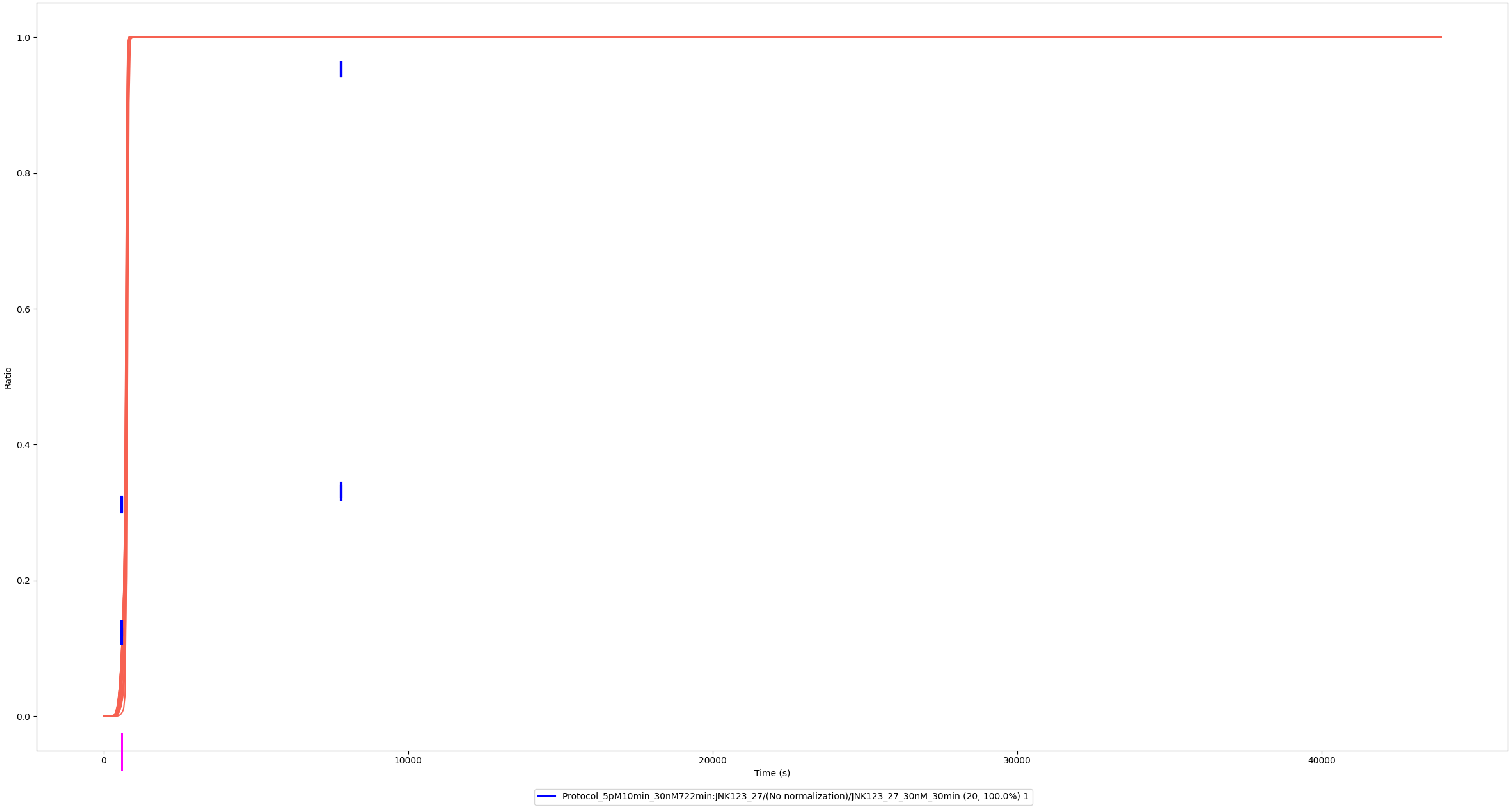

### Experiment vs. Simulation: 30 nM LPS, pp-JNK2

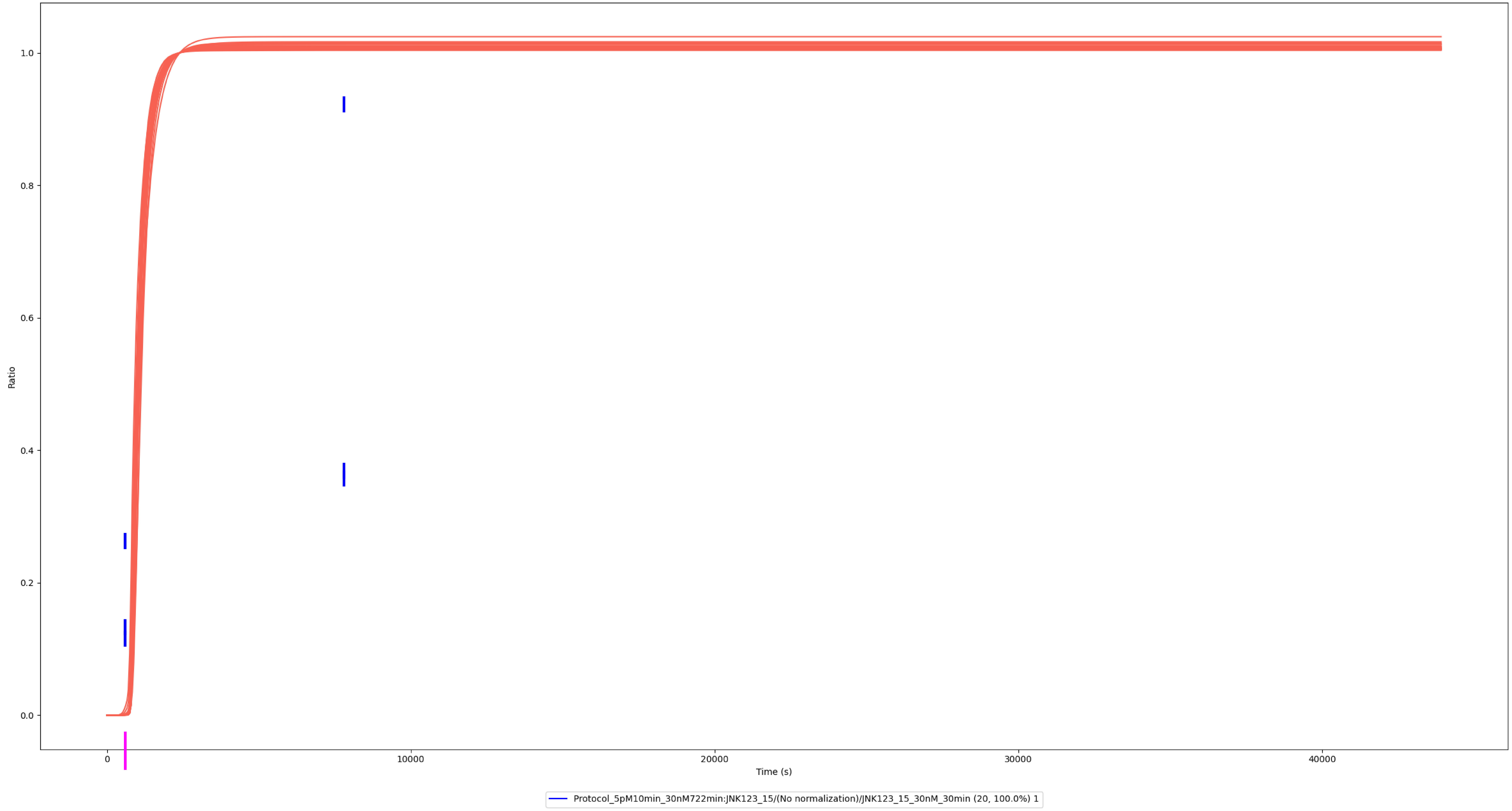

### Experiment vs. Simulation: 30 nM LPS, pp-JNK2

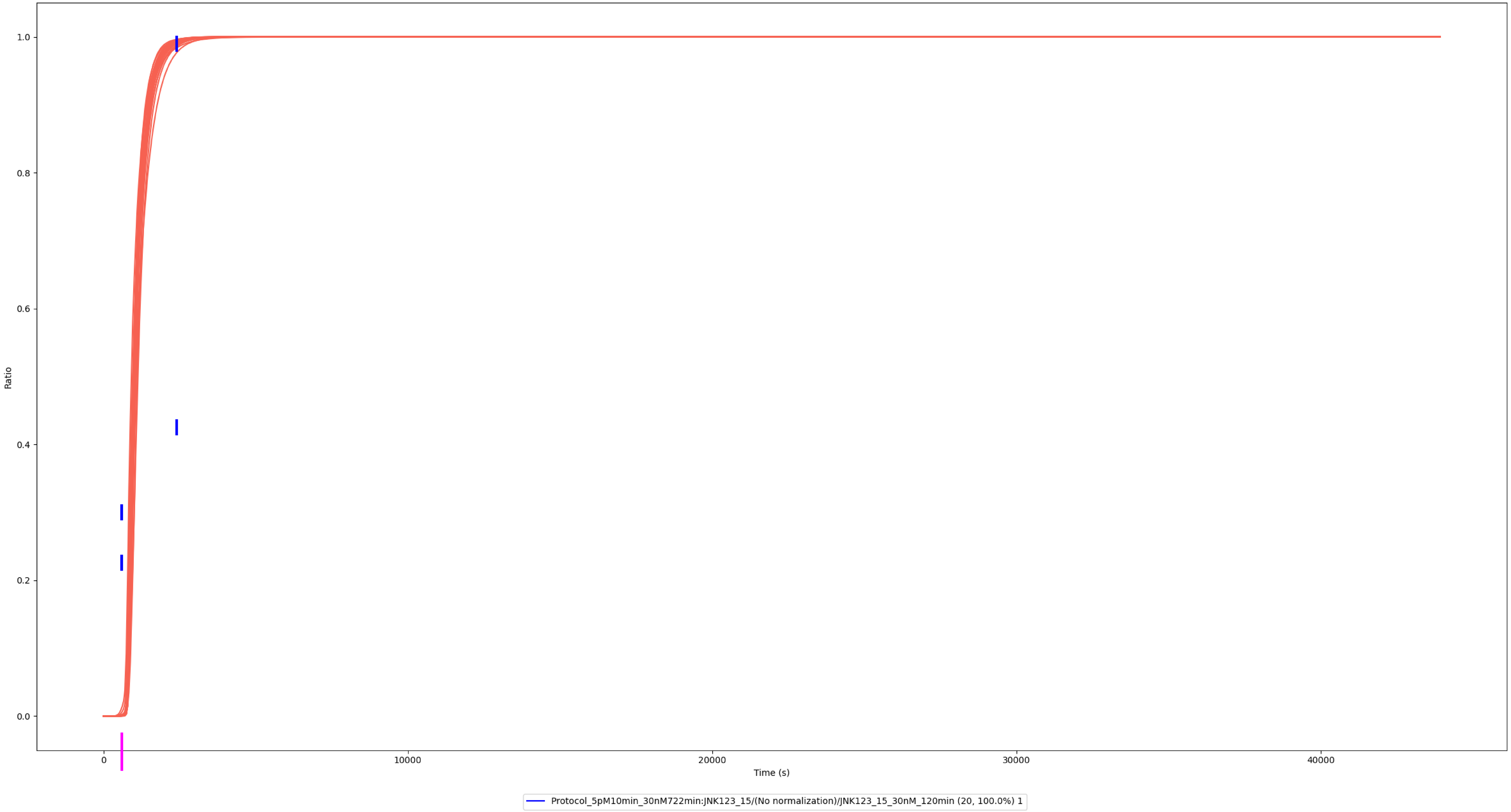

# p38α/β/γ/δ

### Experiment vs. Simulation: 0.01 nM LPS, pp-p38α/β/γ/δ

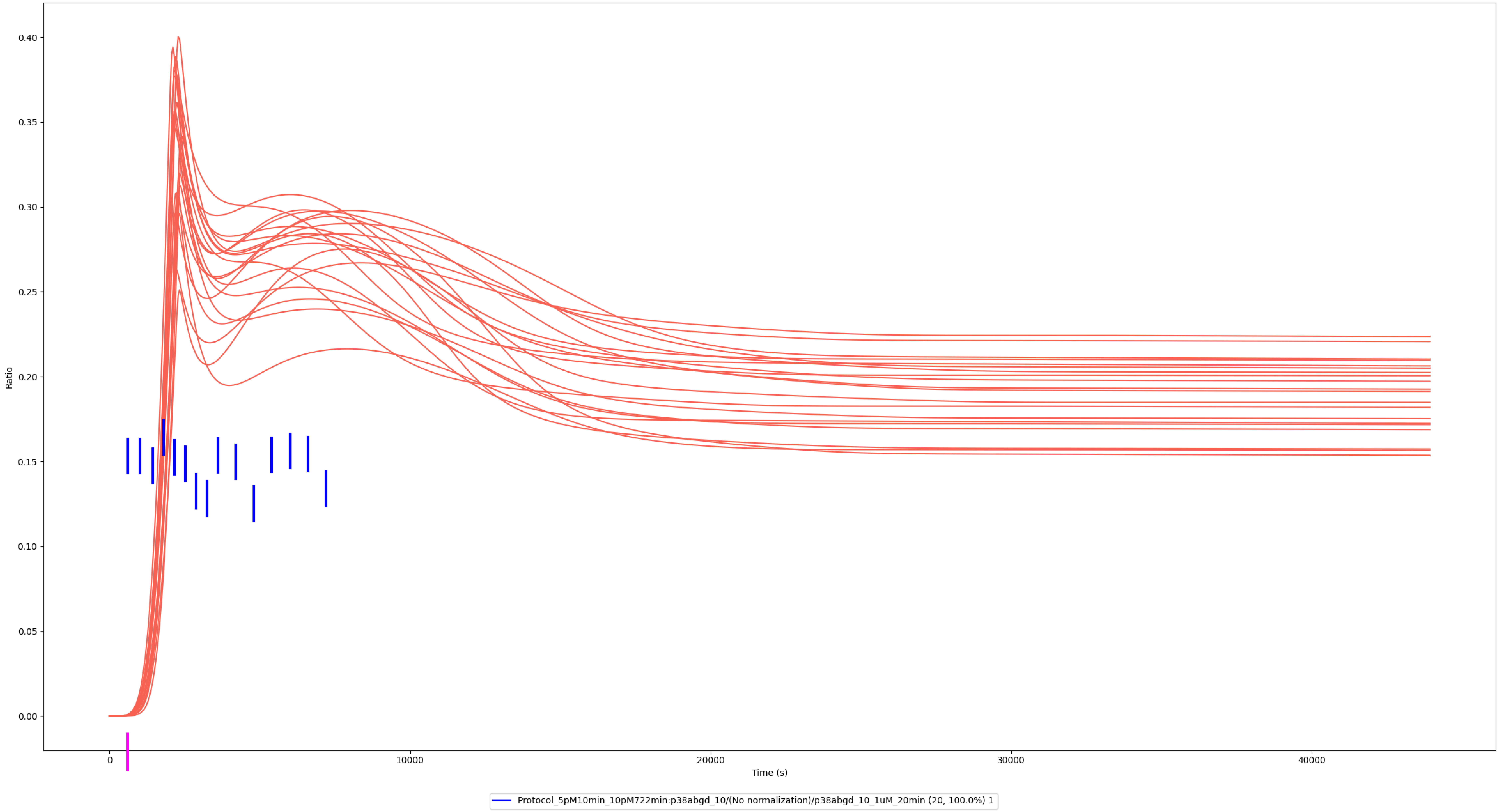

### Experiment vs. Simulation: 0.1 nM LPS, pp-p38α/β/γ/δ

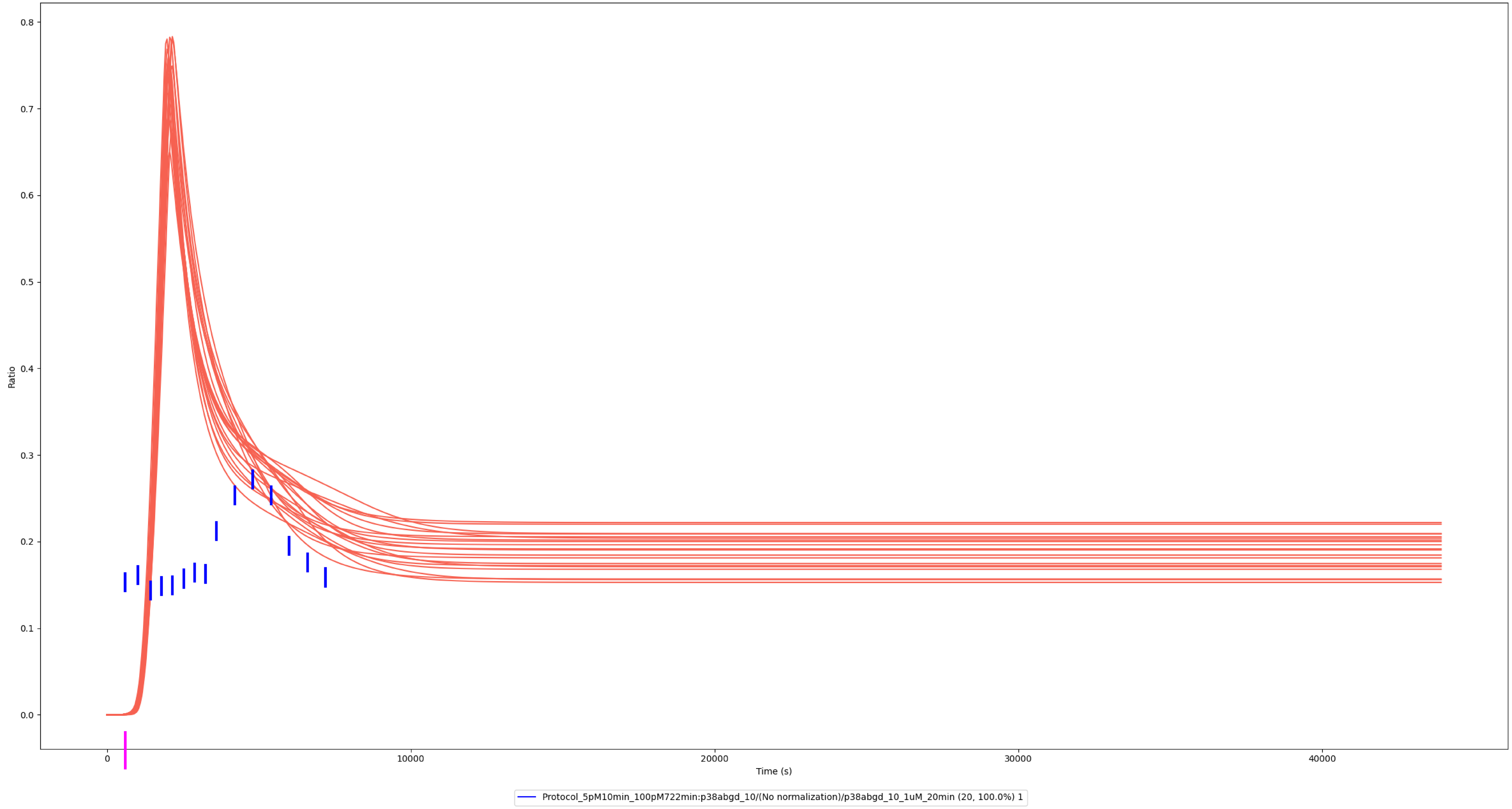

### Experiment vs. Simulation: 1 nM LPS, pp-p38α/β/γ/δ

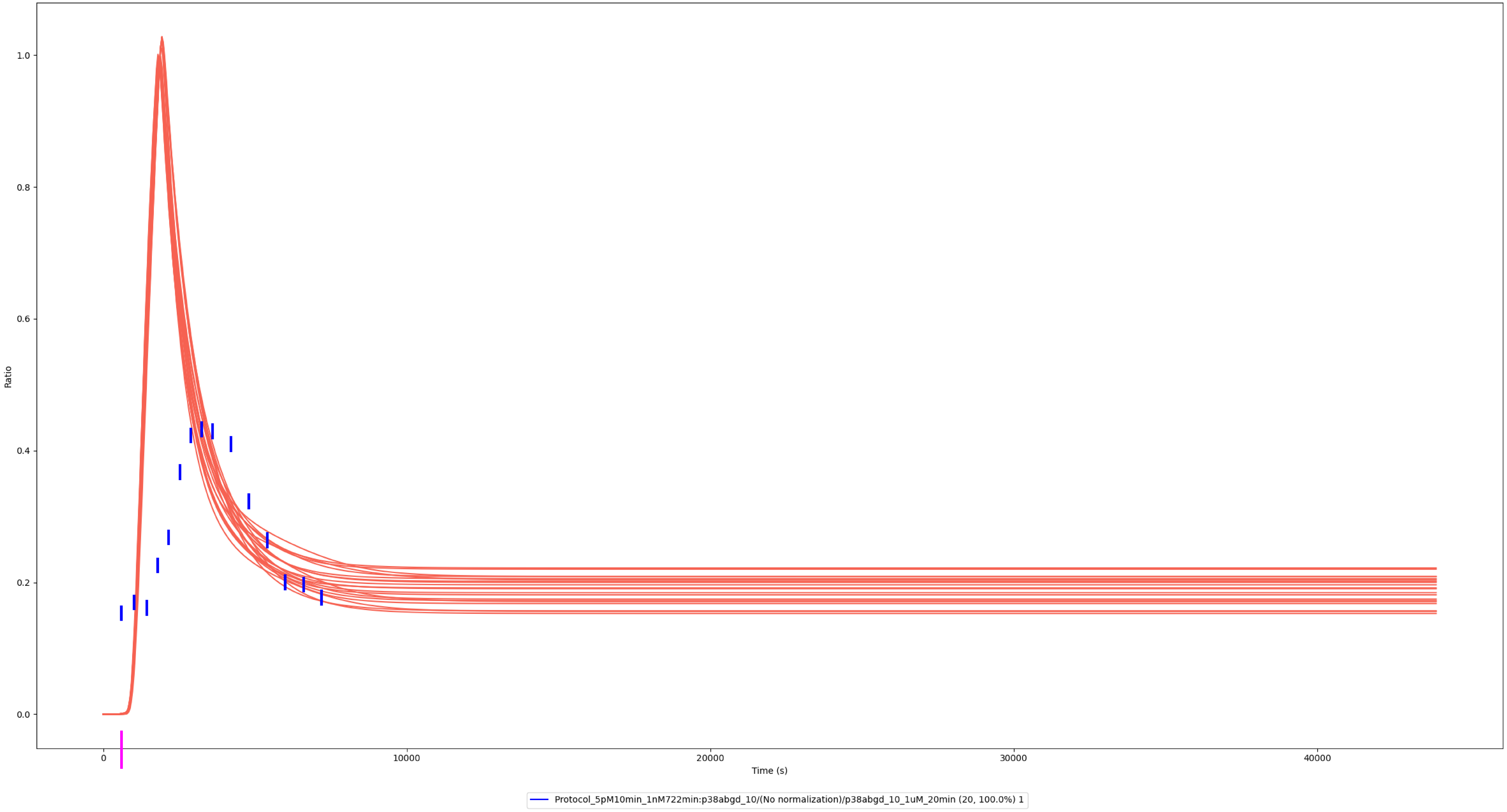

### Experiment vs. Simulation: 10 nM LPS, pp-p38α/β/γ/δ

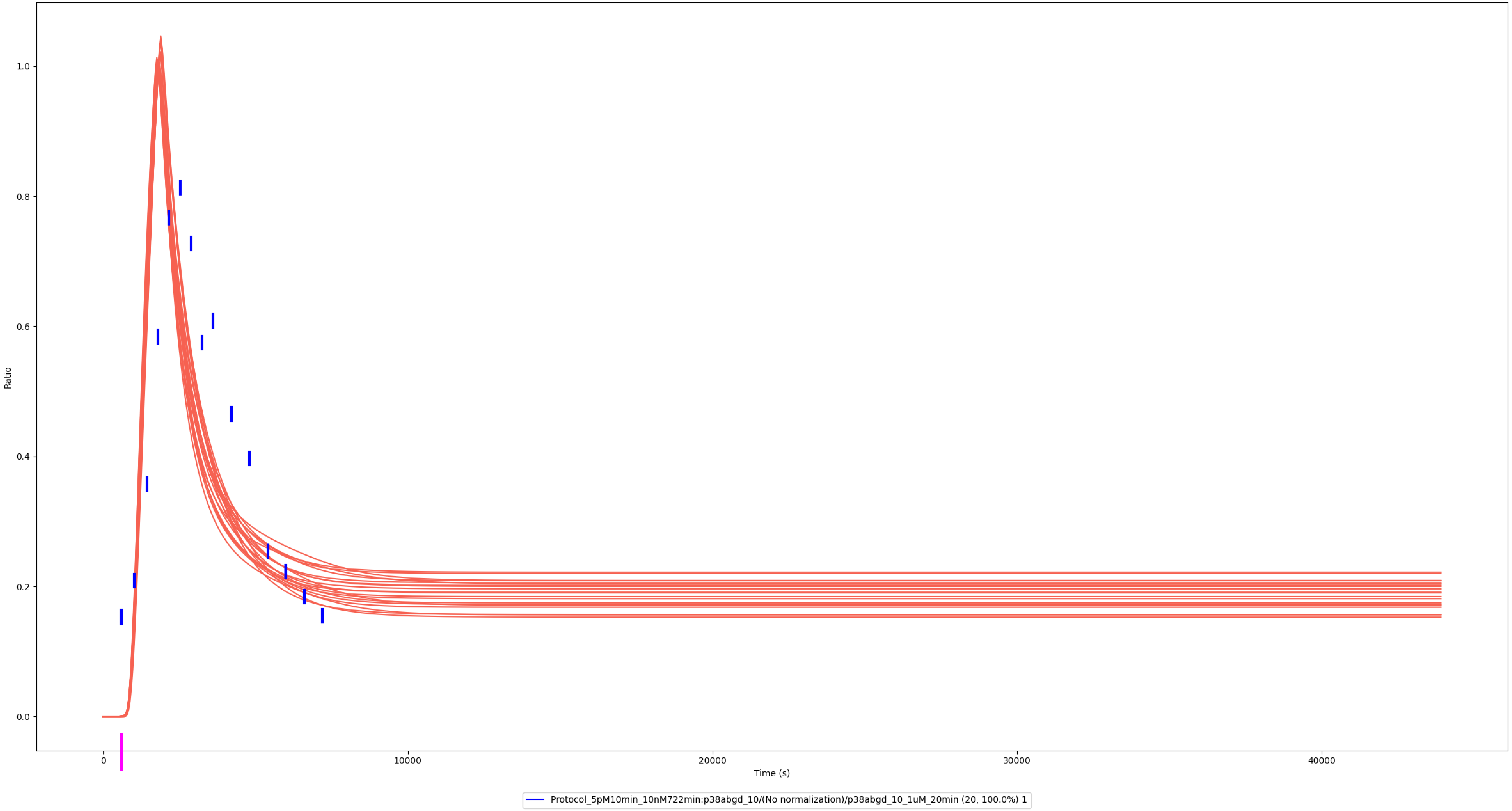

### Experiment vs. Simulation: 100 nM LPS, pp-p38α/β/γ/δ

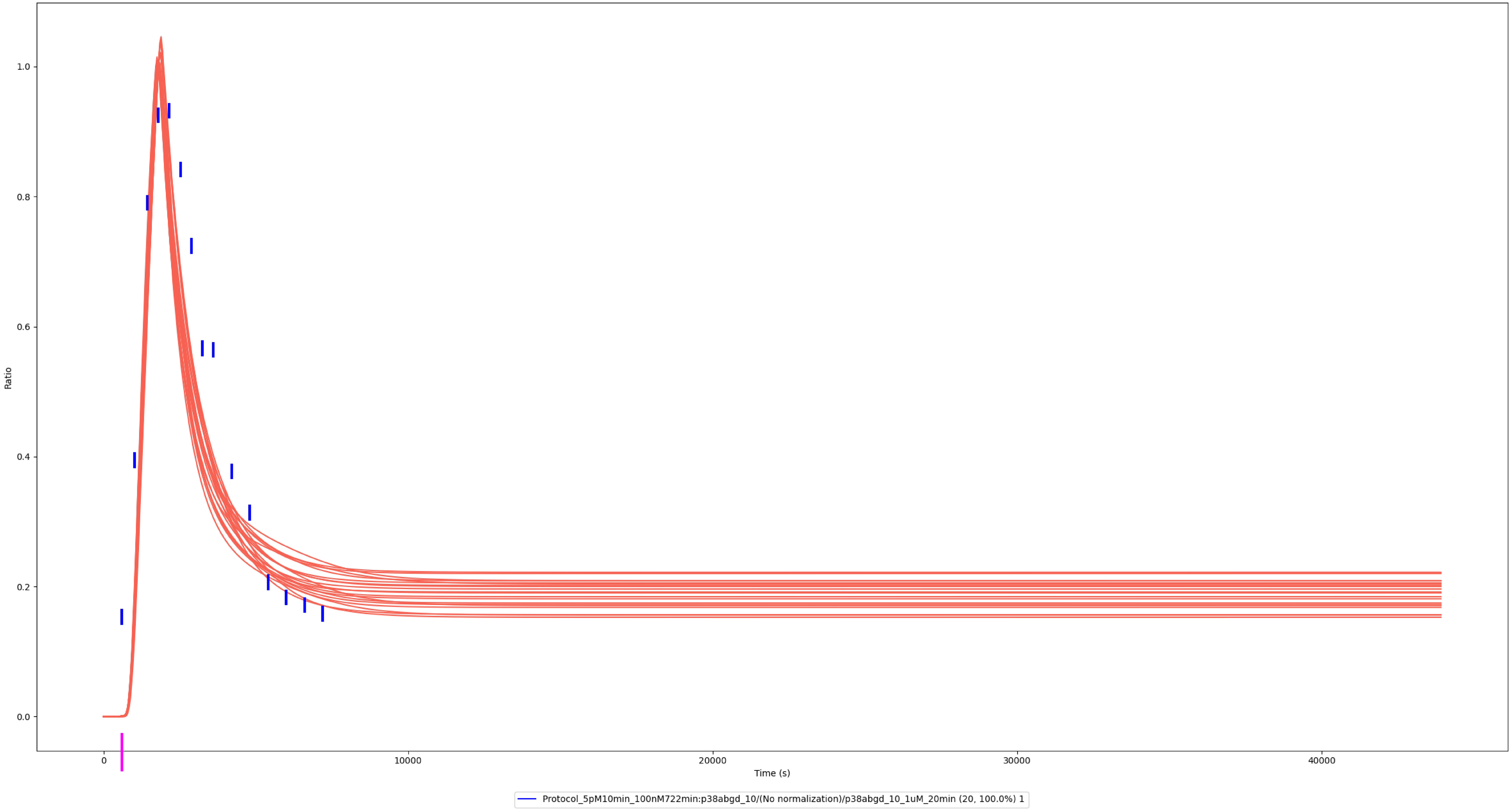

### Experiment vs. Simulation: 1,000 nM LPS, pp-p38α/β/γ/δ

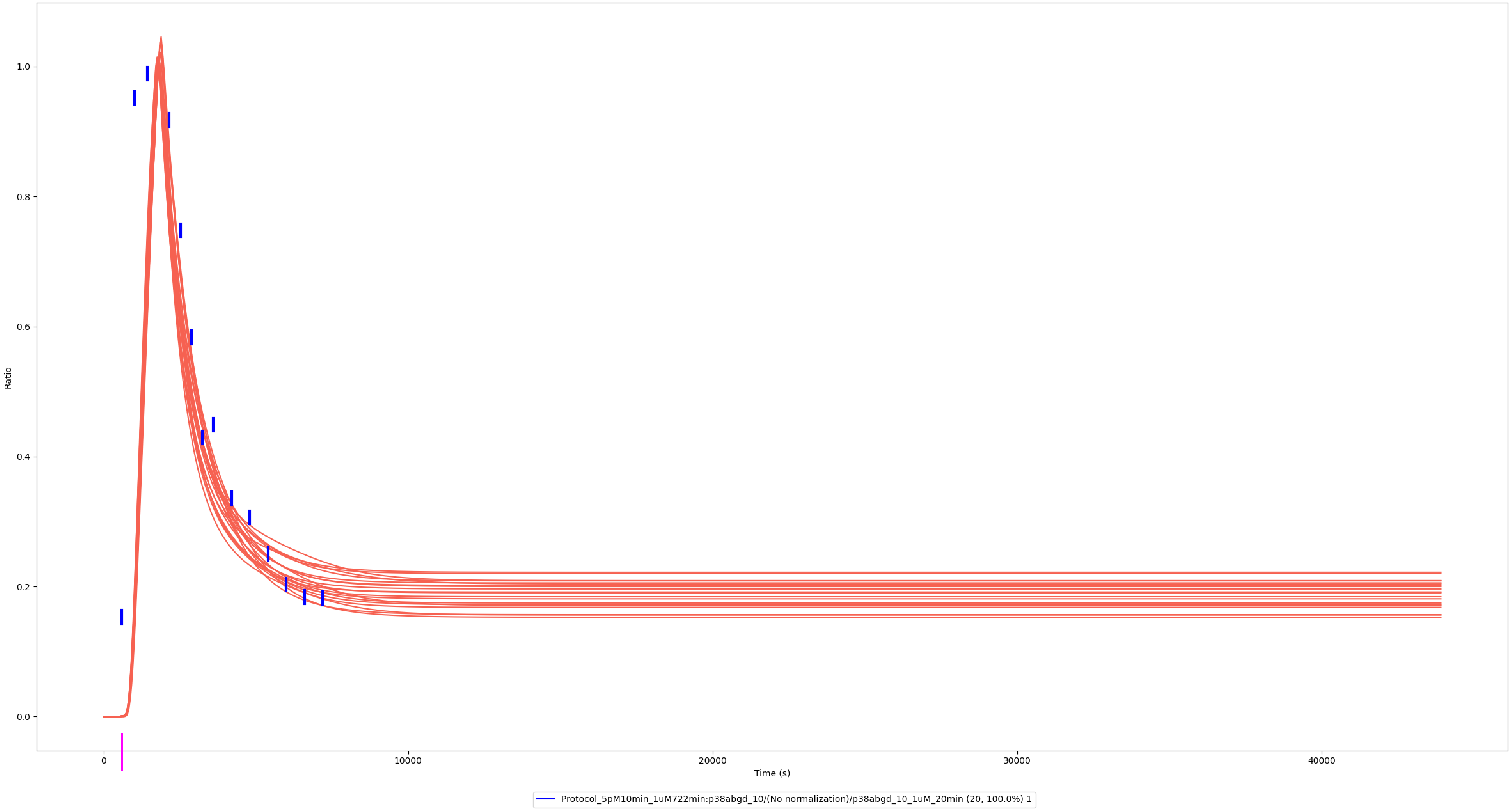

### Experiment vs. Simulation: 1 nM LPS, pp-p38α/β/γ/δ

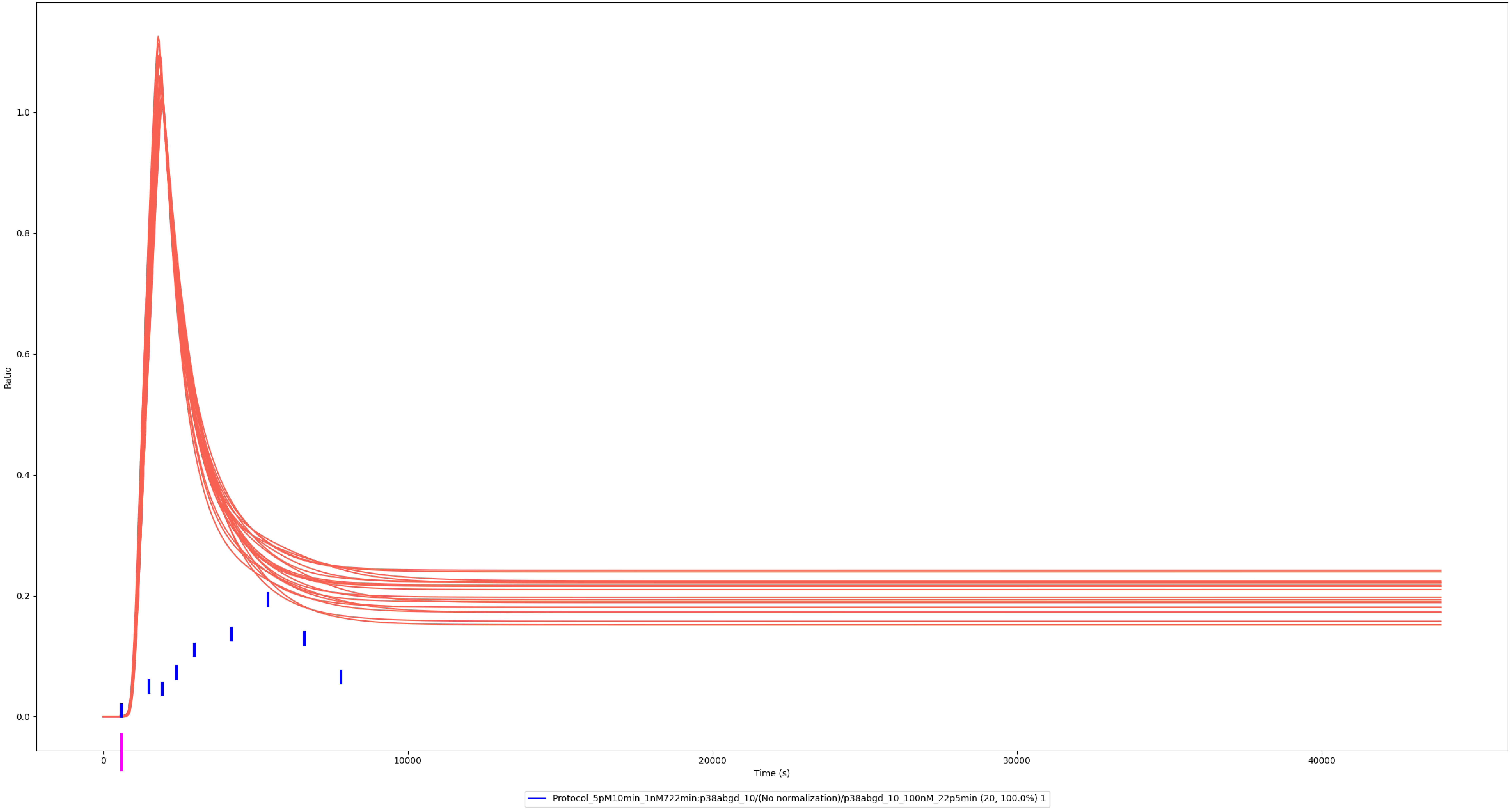

### Experiment vs. Simulation: 100 nM LPS, pp-p38α/β/γ/δ

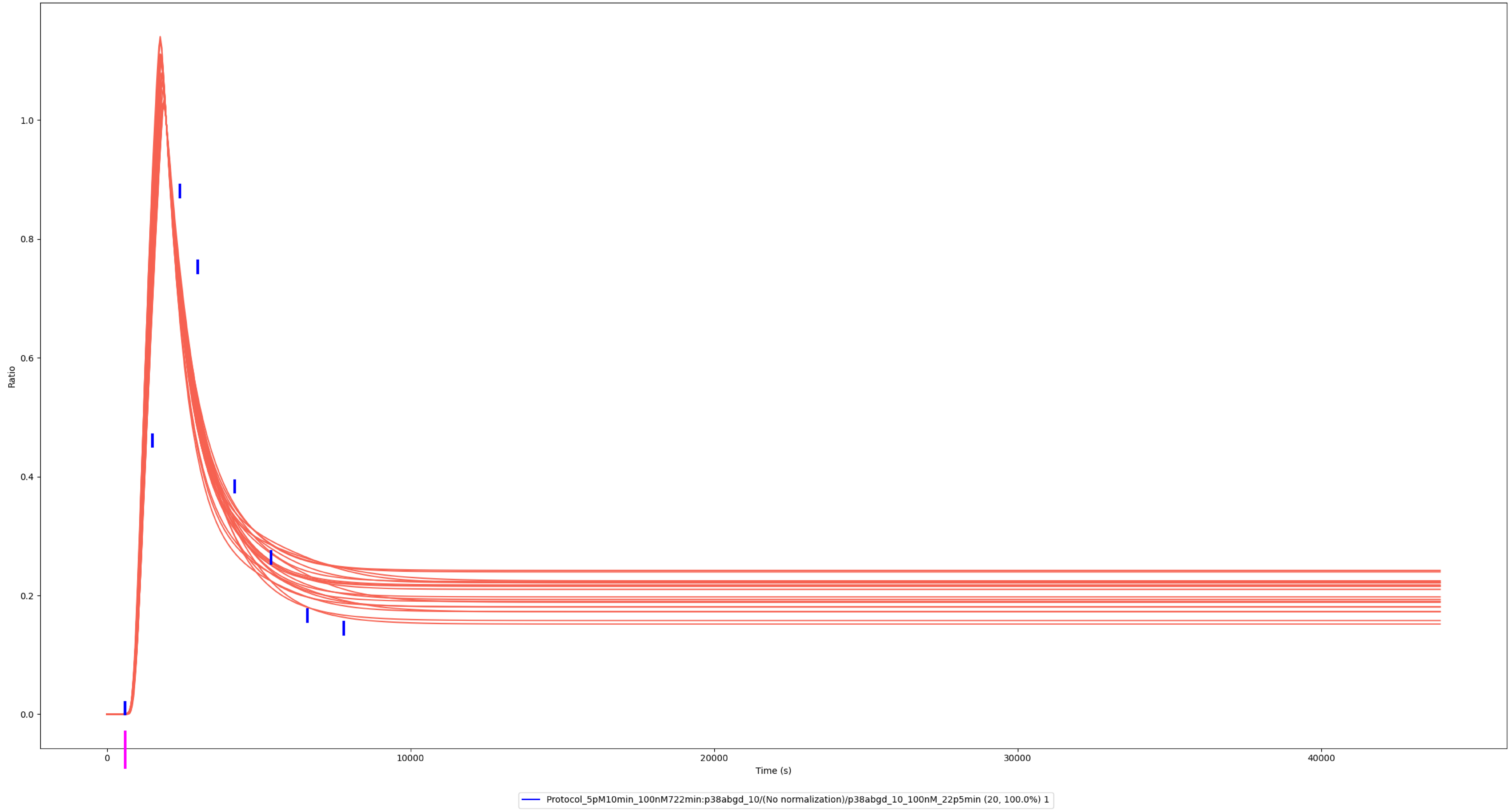

### Experiment vs. Simulation: 3 nM LPS, pp-p38α

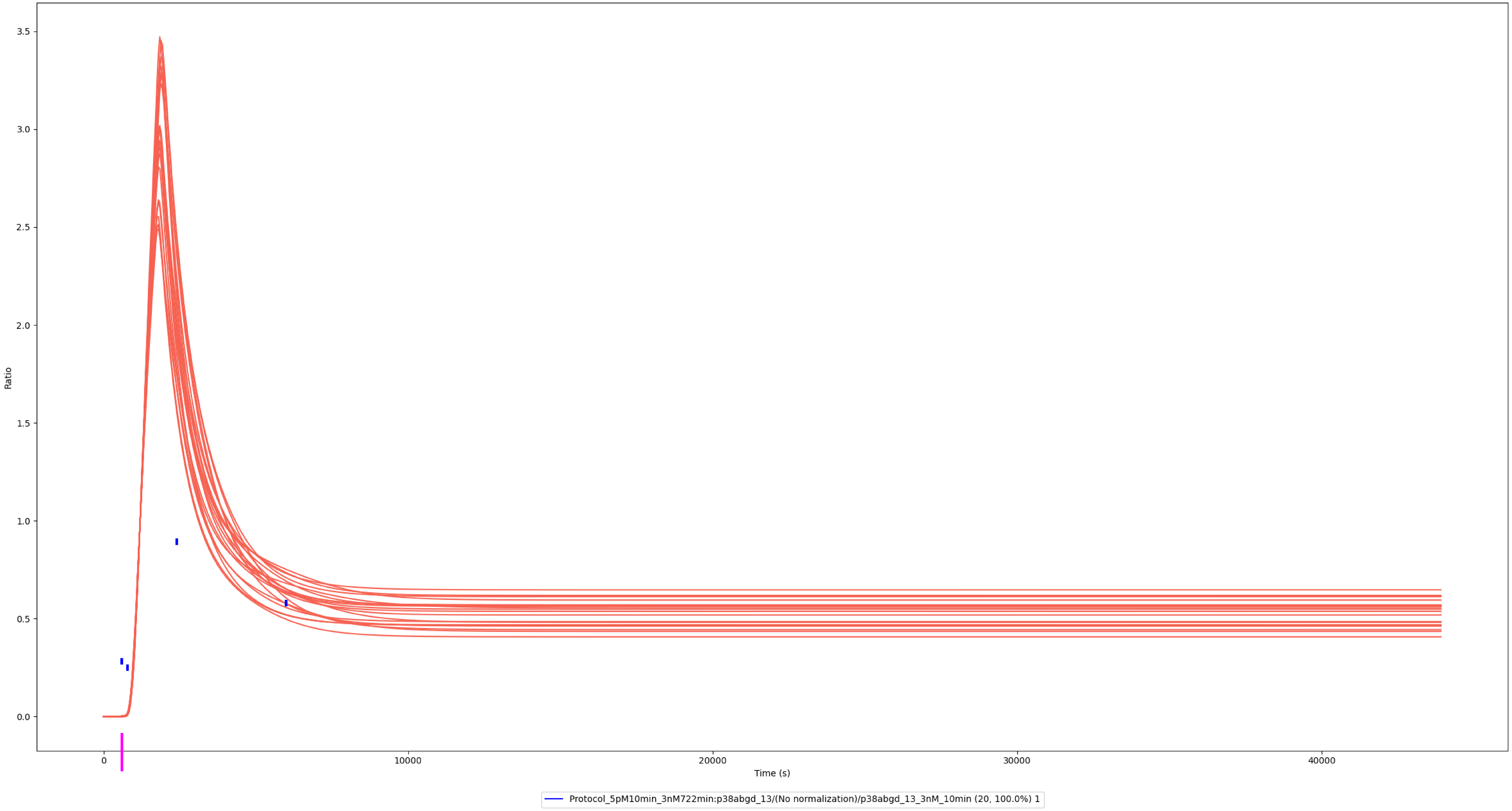

### Experiment vs. Simulation: 30 nM LPS, pp-p38α

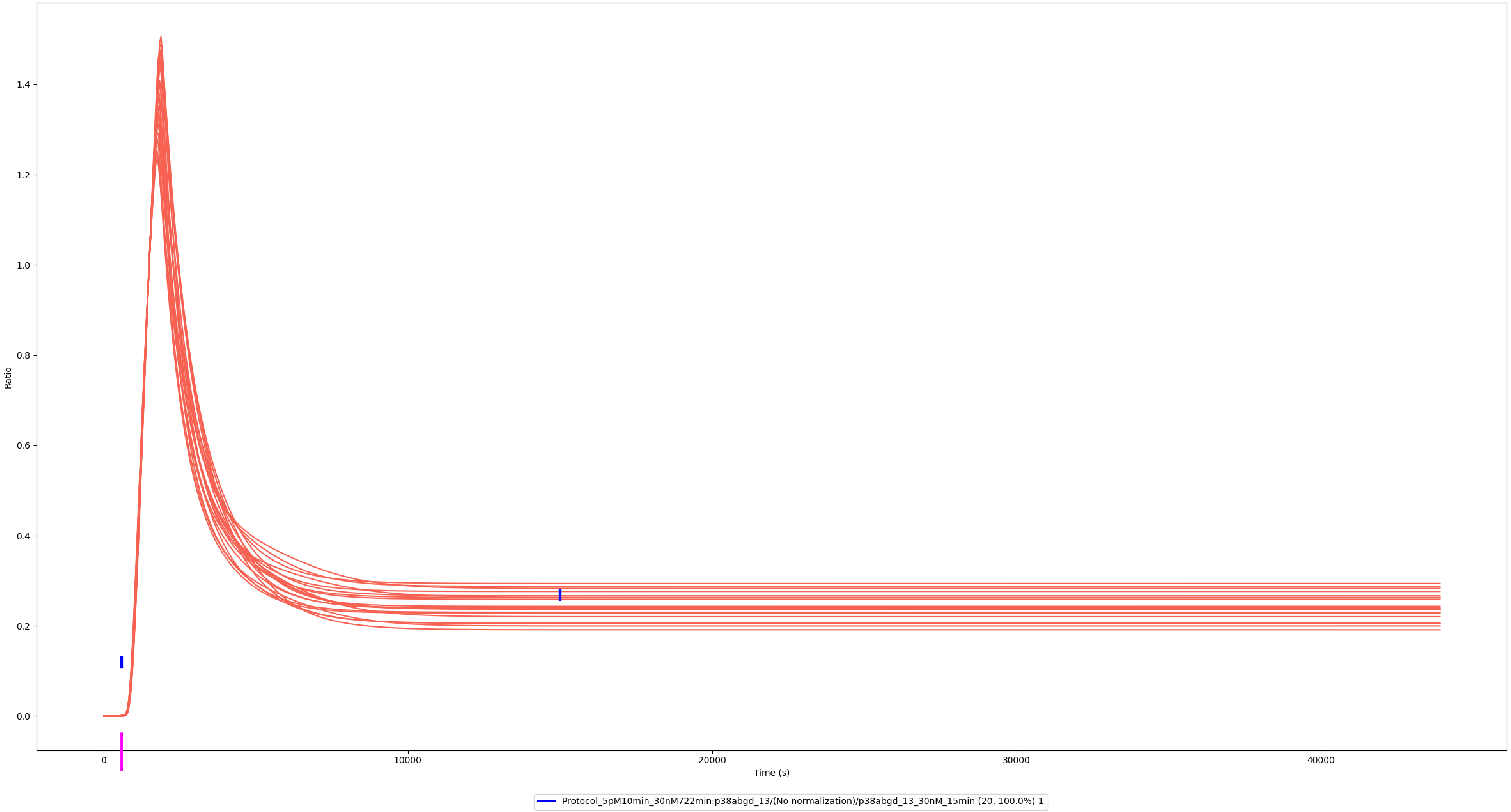

### Experiment vs. Simulation: 30 nM LPS, pp-p38α

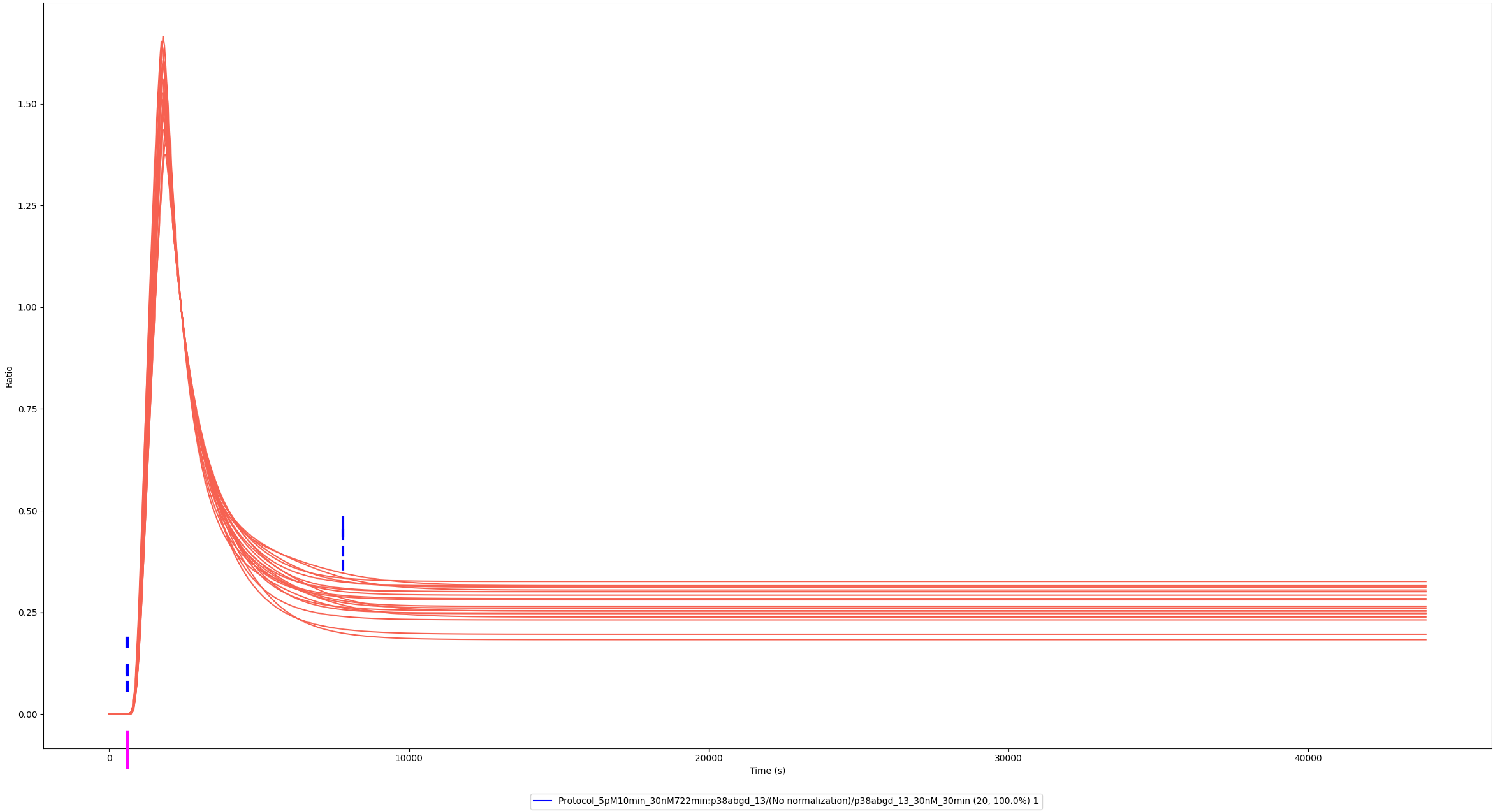

### Experiment vs. Simulation: 30 nM LPS, pp-p38β

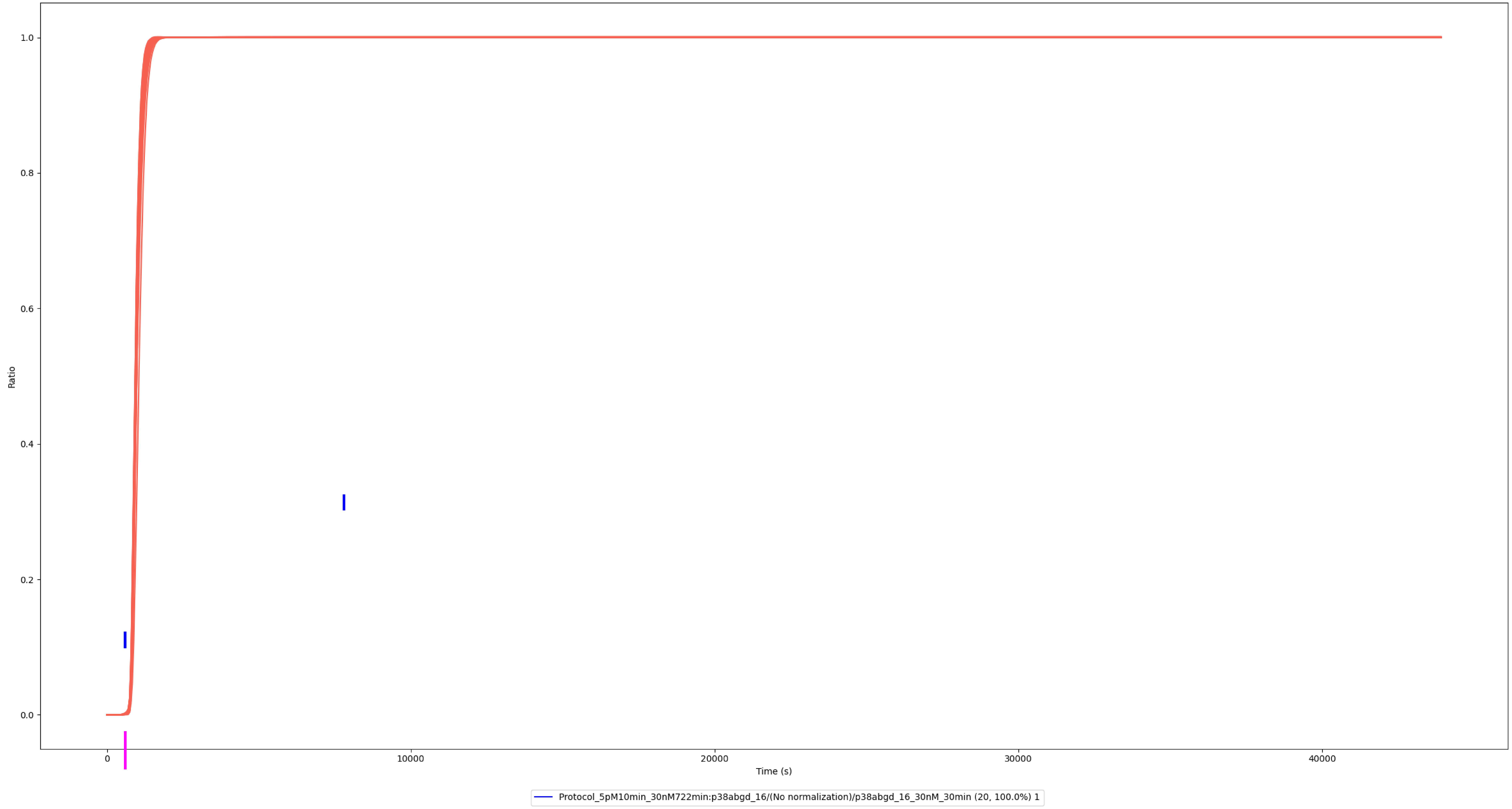

### Experiment vs. Simulation: 30 nM LPS, pp-p38β

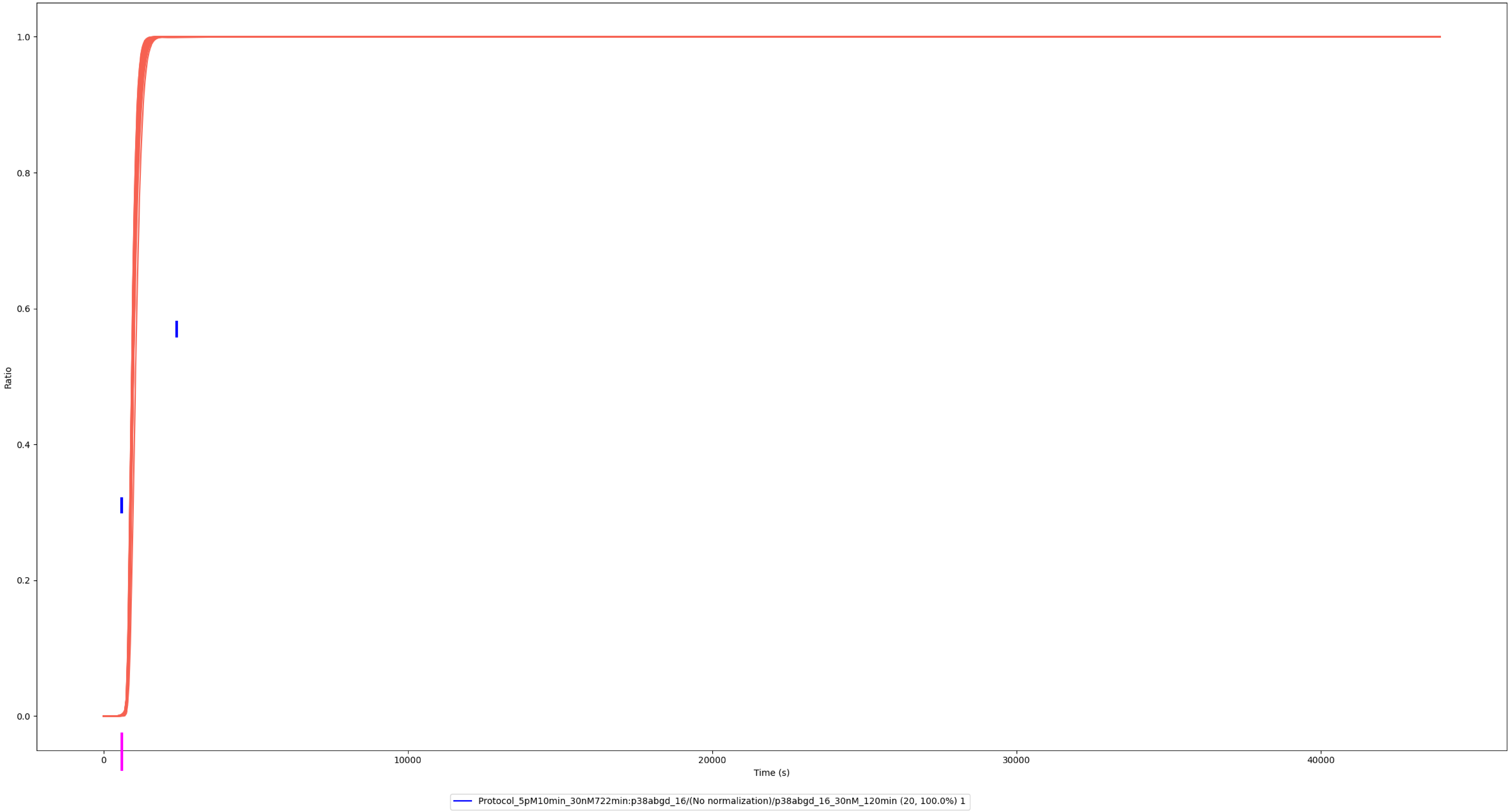

### Experiment vs. Simulation: 30 nM LPS, pp-p38γ

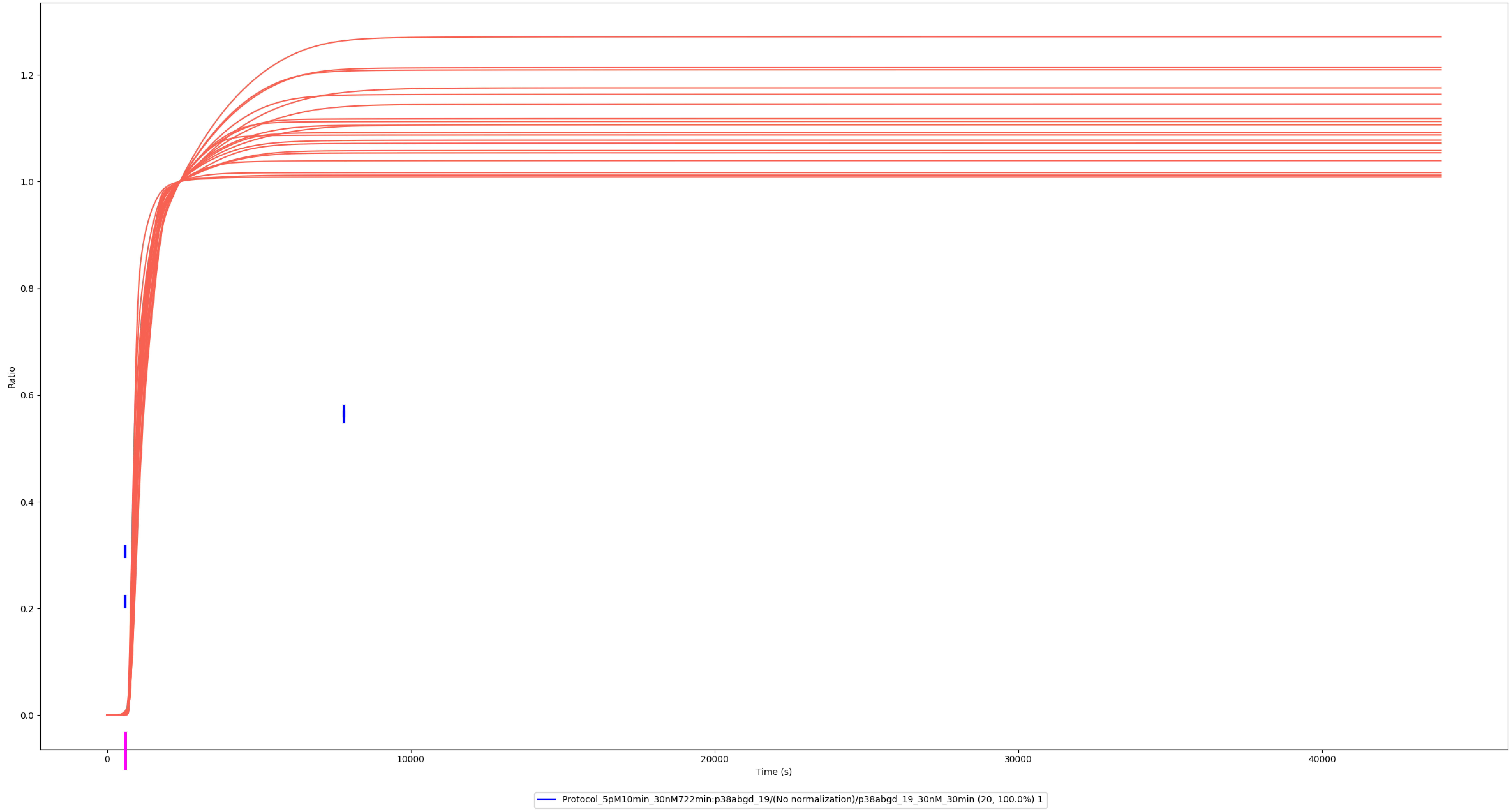

### Experiment vs. Simulation: 30 nM LPS, pp-p38γ

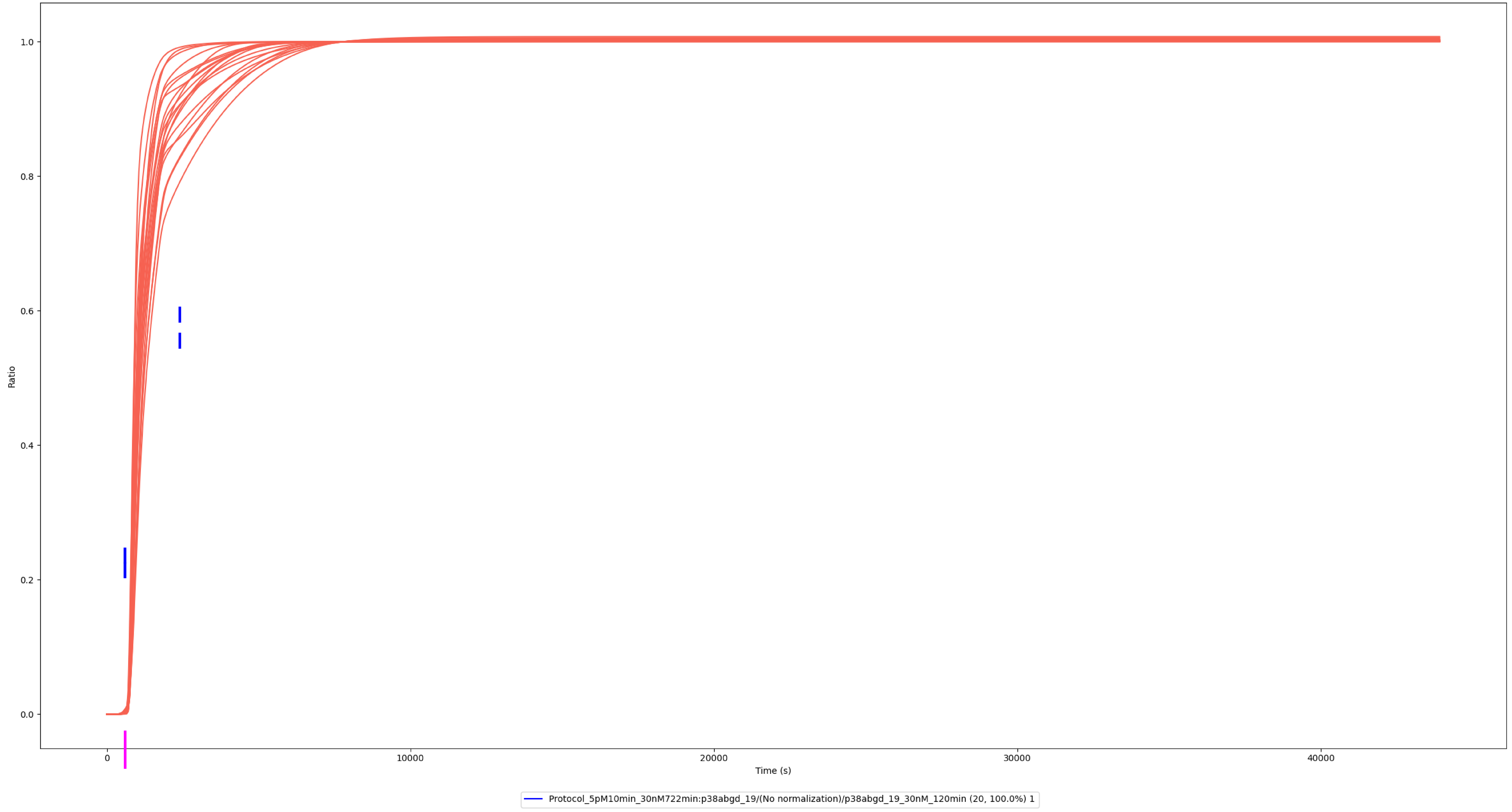

### MSK1

### Experiment vs. Simulation: 3 nM LPS, pppp-MSK1

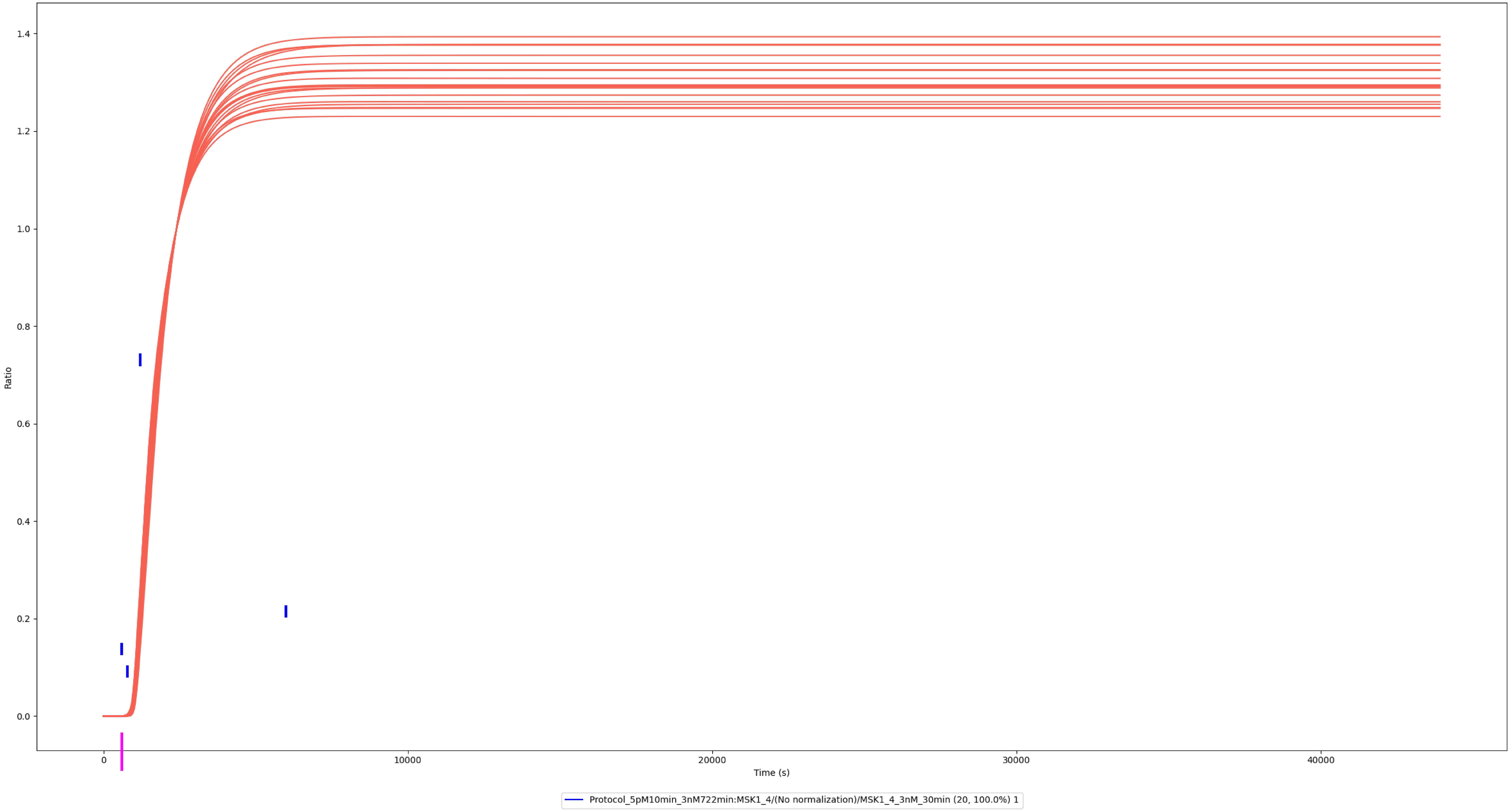

### Experiment vs. Simulation: 30 nM LPS, pppp-MSK1

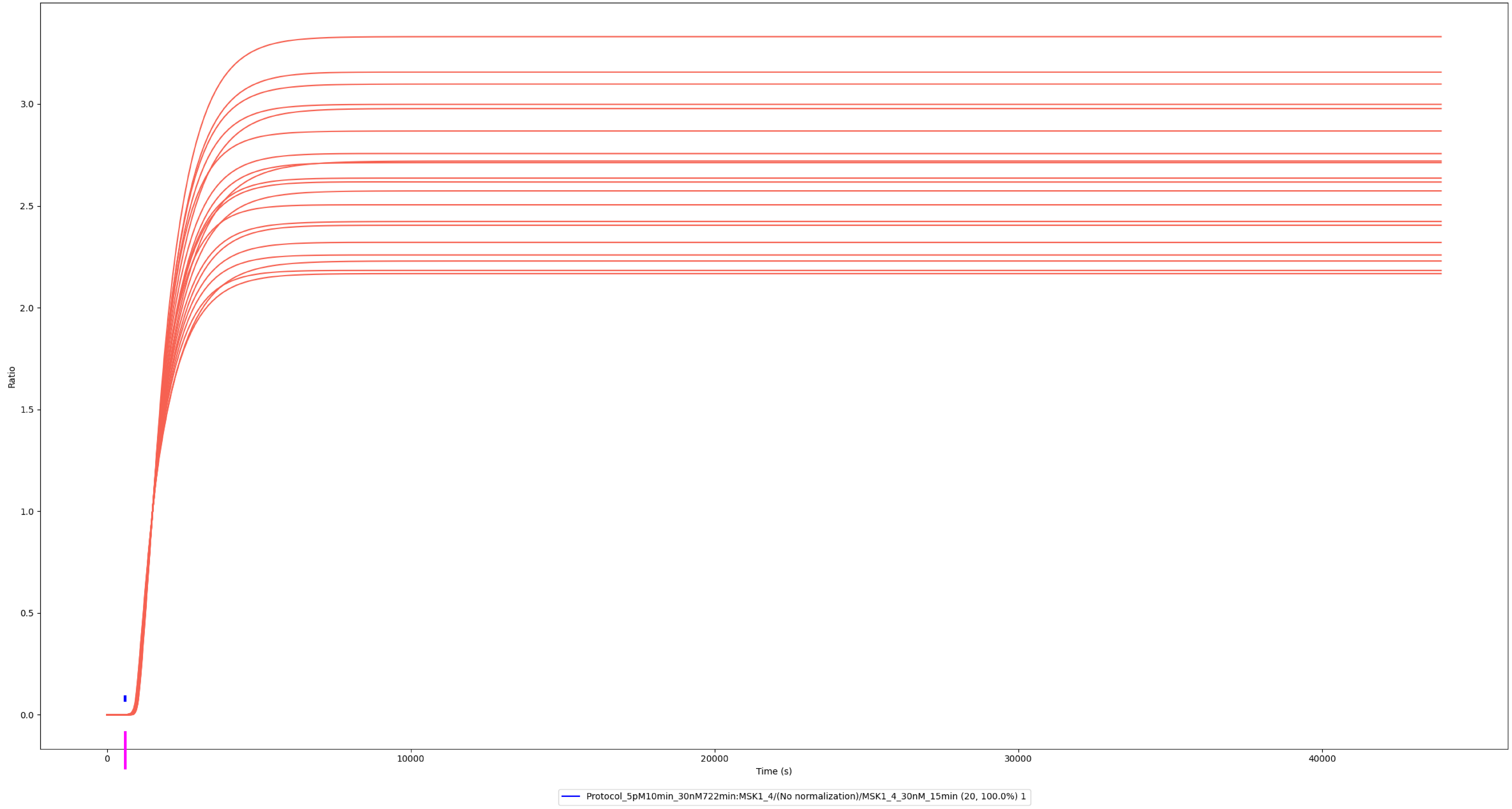

### Experiment vs. Simulation: 30 nM LPS, pppp-MSK1

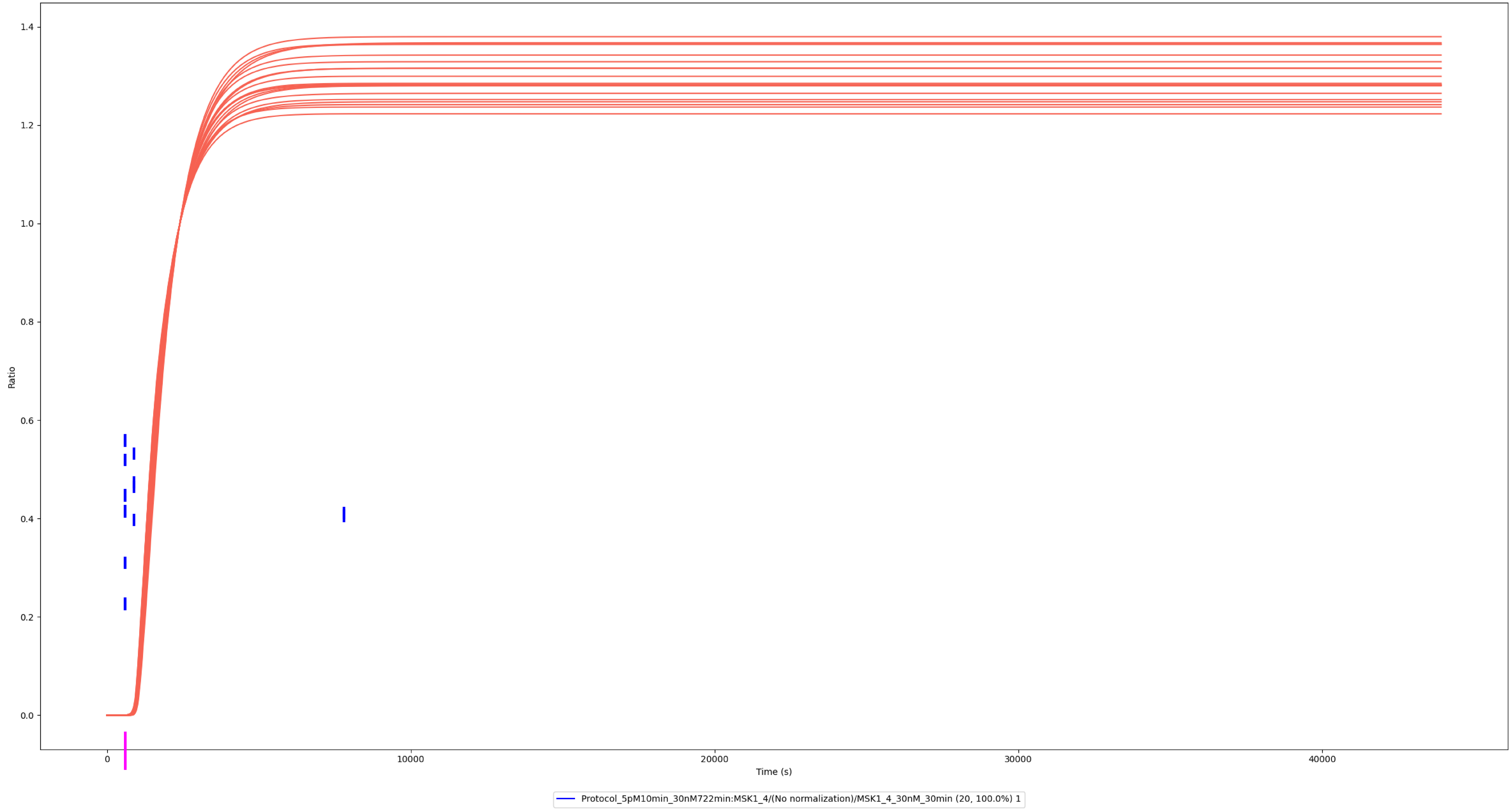

### Jun, JunB, JunD

### Experiment vs. Simulation: 30 nM LPS, p-Jun

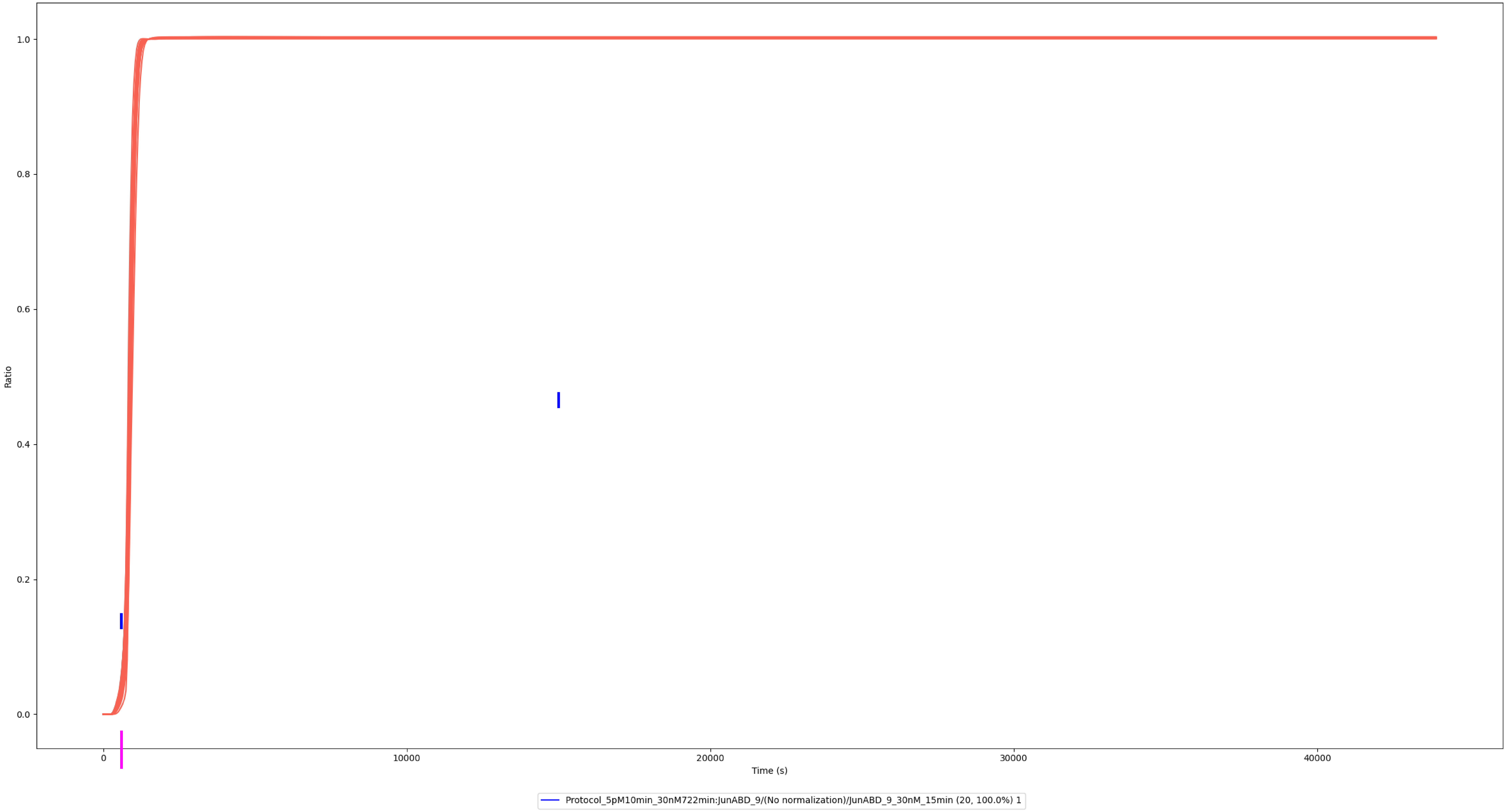

### Experiment vs. Simulation: 30 nM LPS, p-Jun

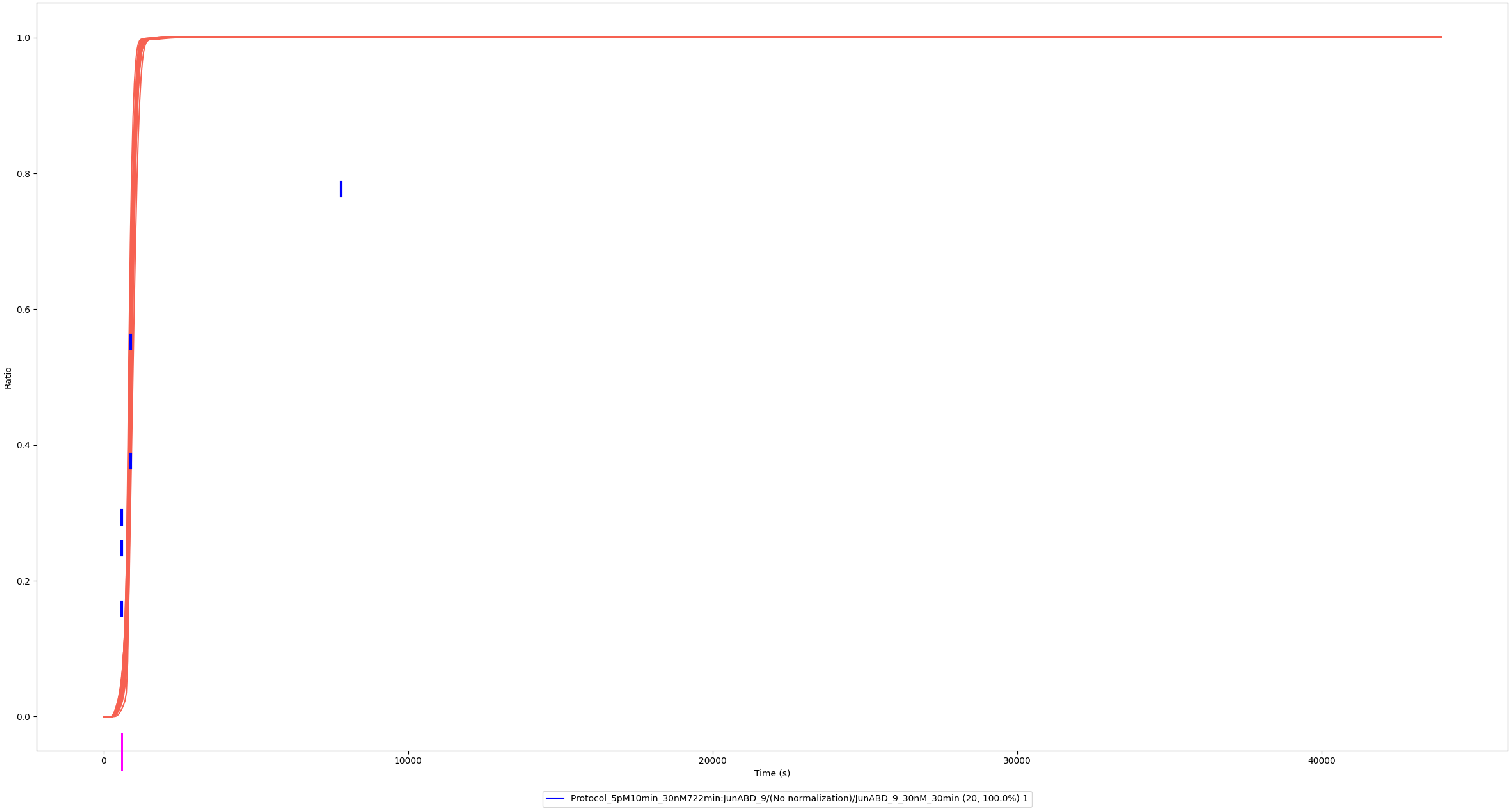

### Experiment vs. Simulation: 30 nM LPS, p-Jun

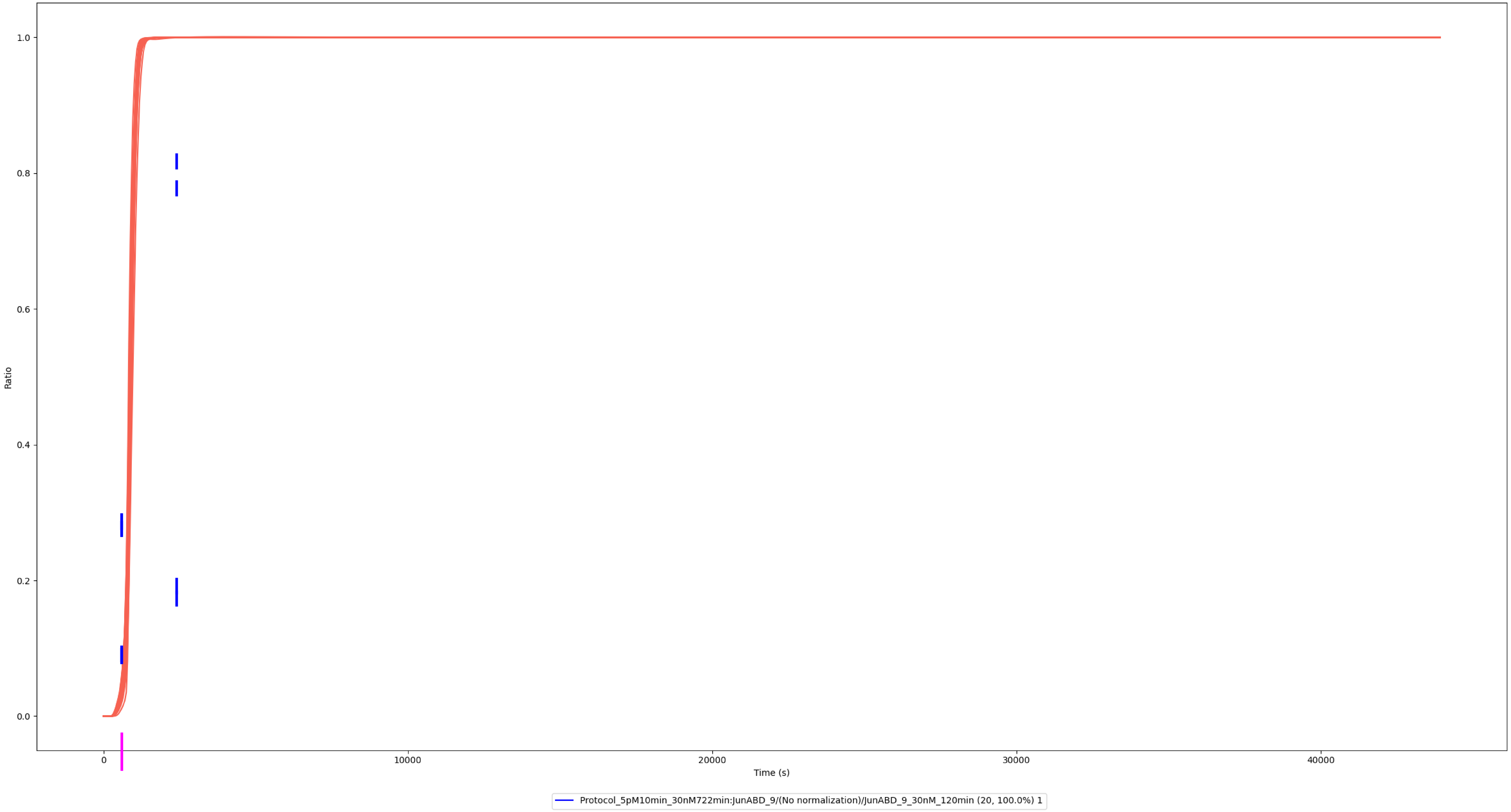

### Experiment vs. Simulation: 300 nM LPS, p-Jun

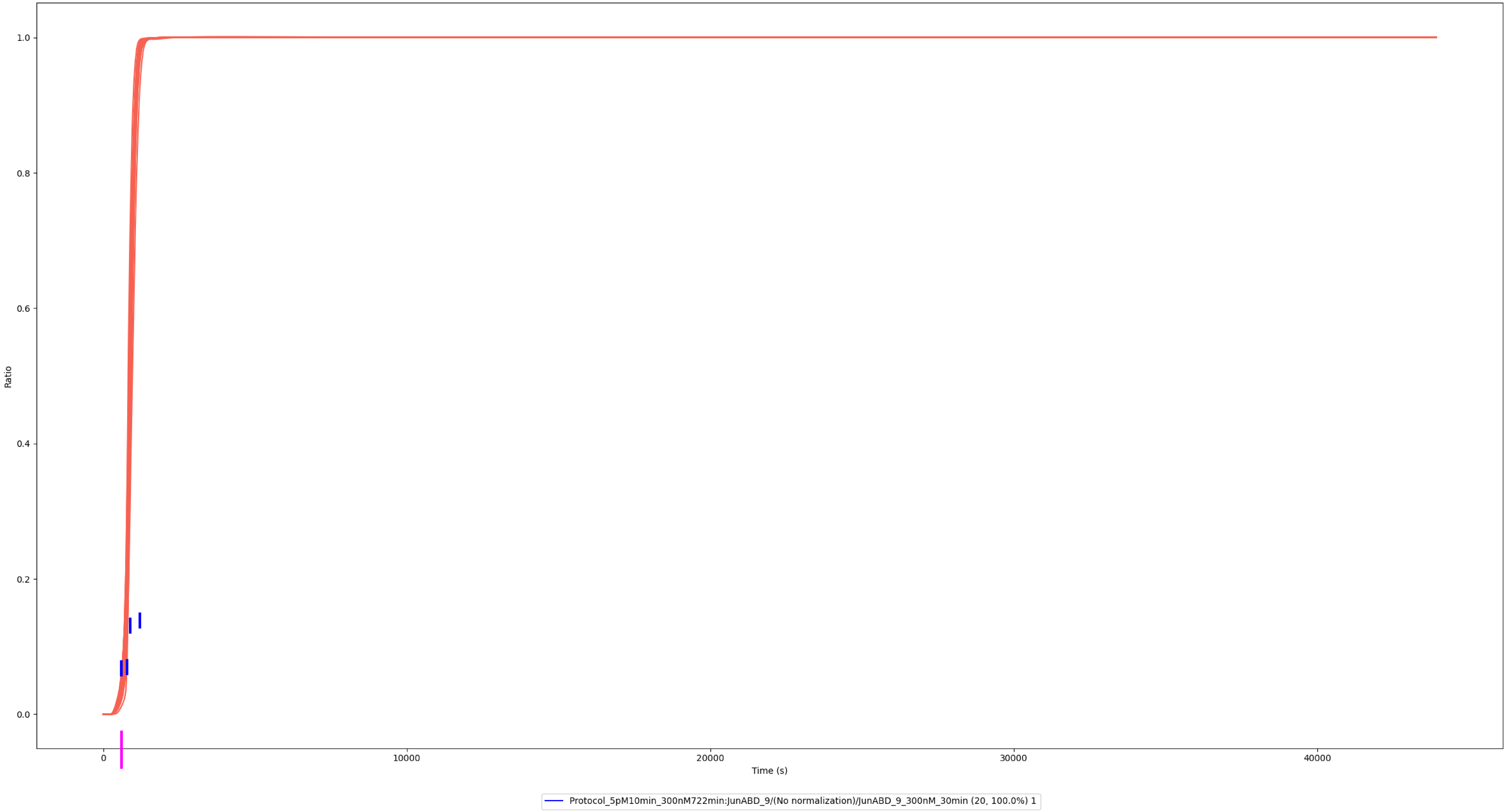

### Experiment vs. Simulation: 30 nM LPS, p-JunB

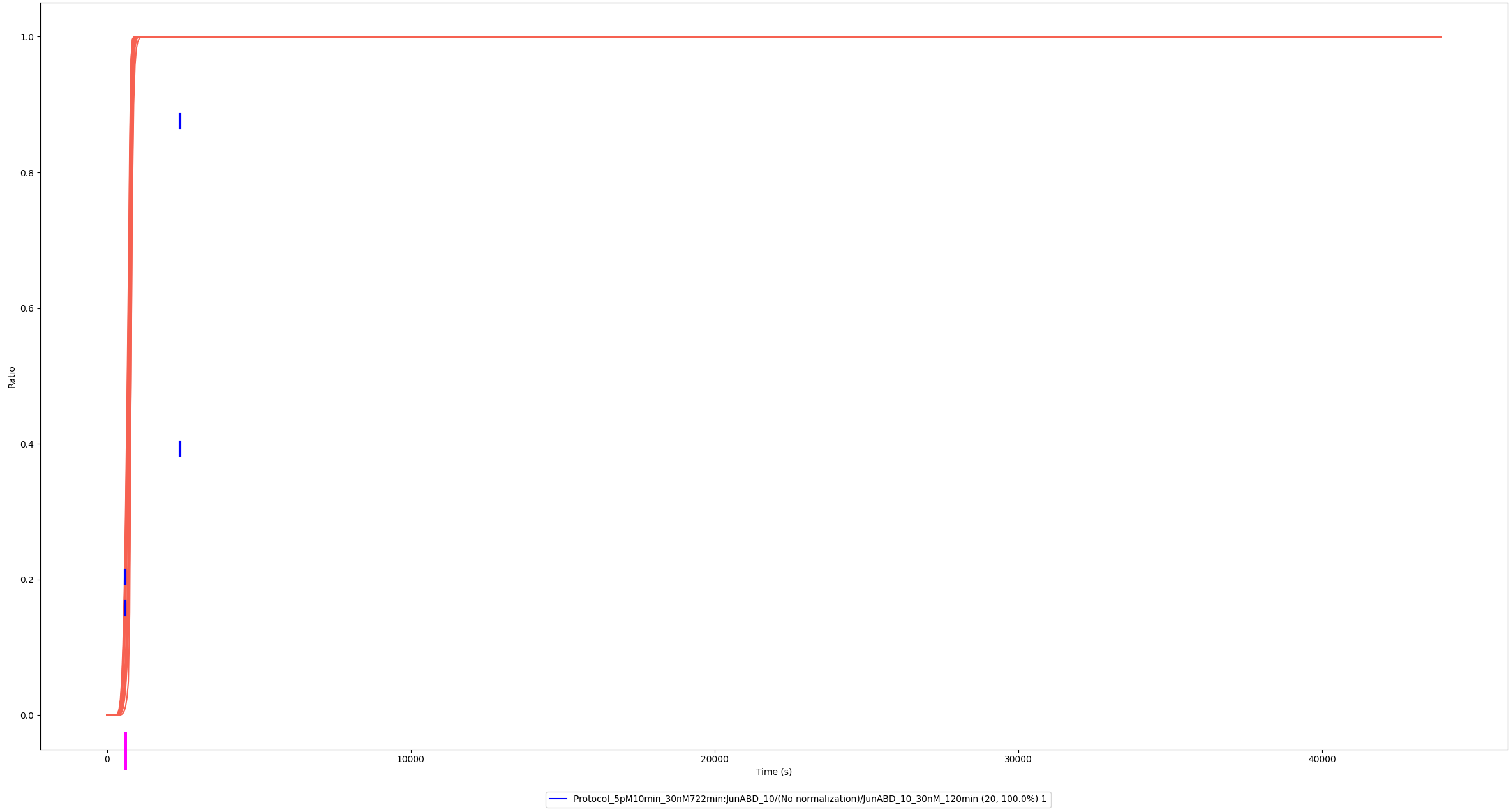

### Experiment vs. Simulation: 30 nM LPS, p-JunD

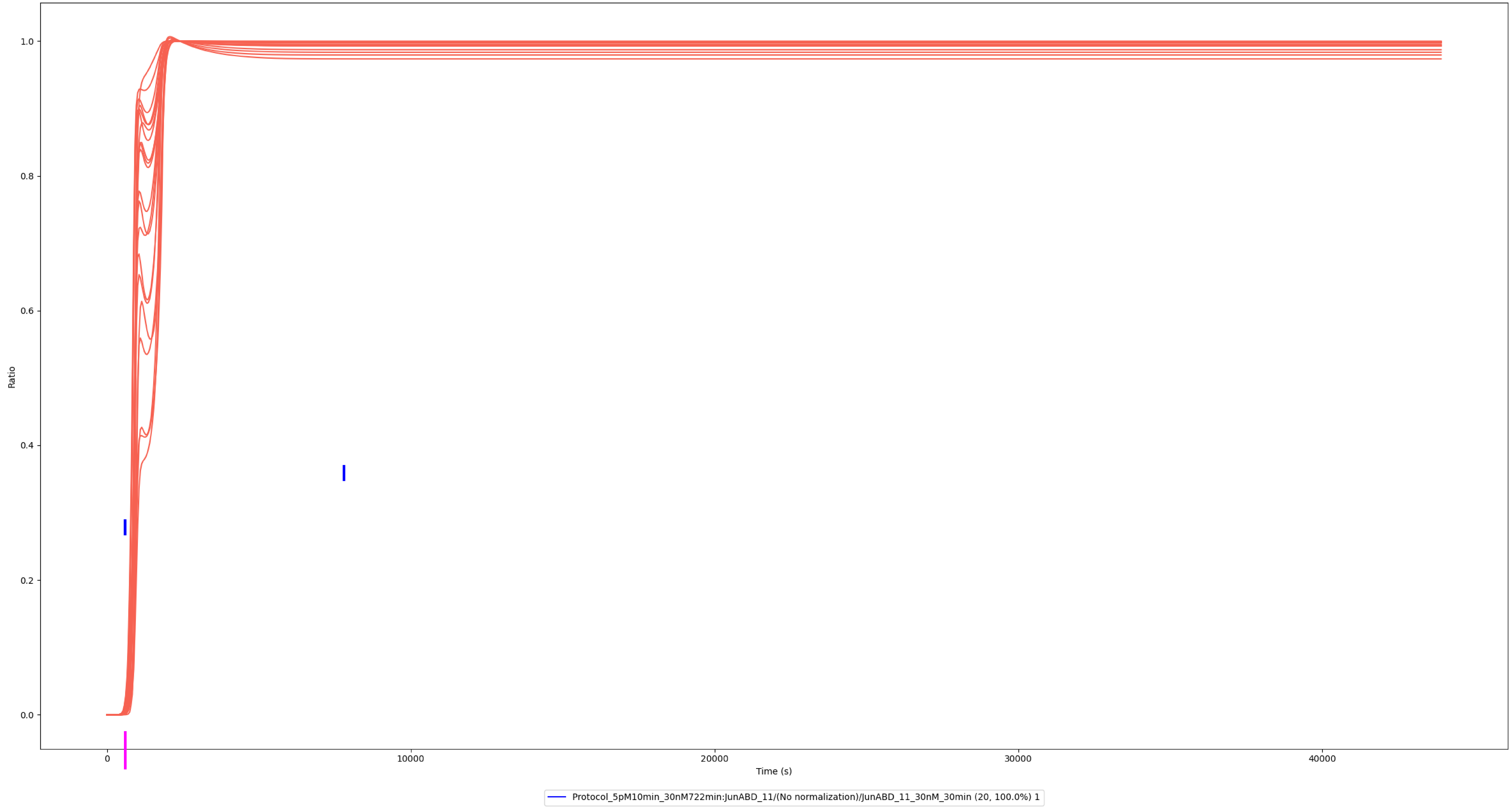

### Experiment vs. Simulation: 30 nM LPS, p-Jun/p-JunD

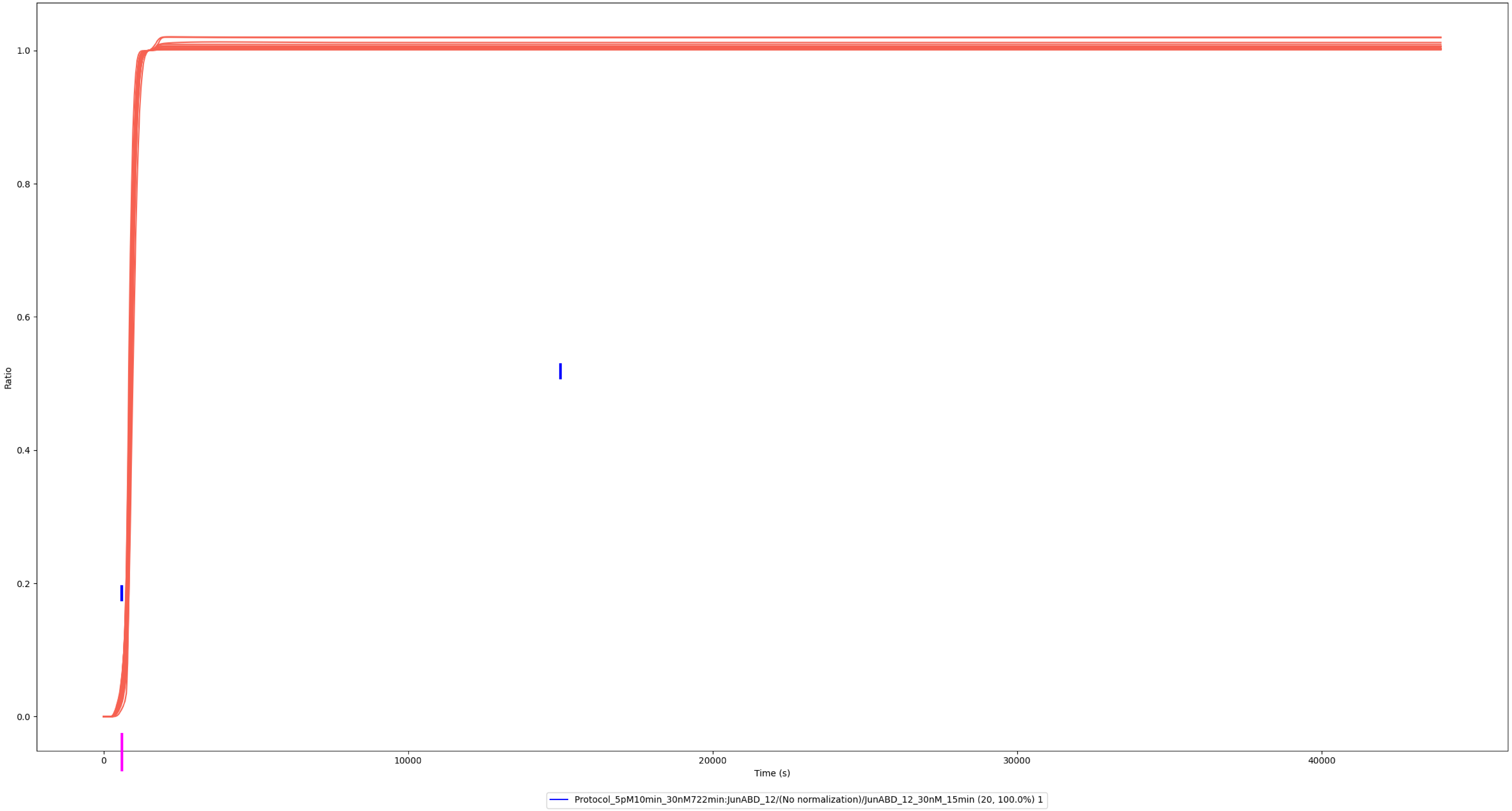

### Experiment vs. Simulation: 30 nM LPS, p-Jun/p-JunD

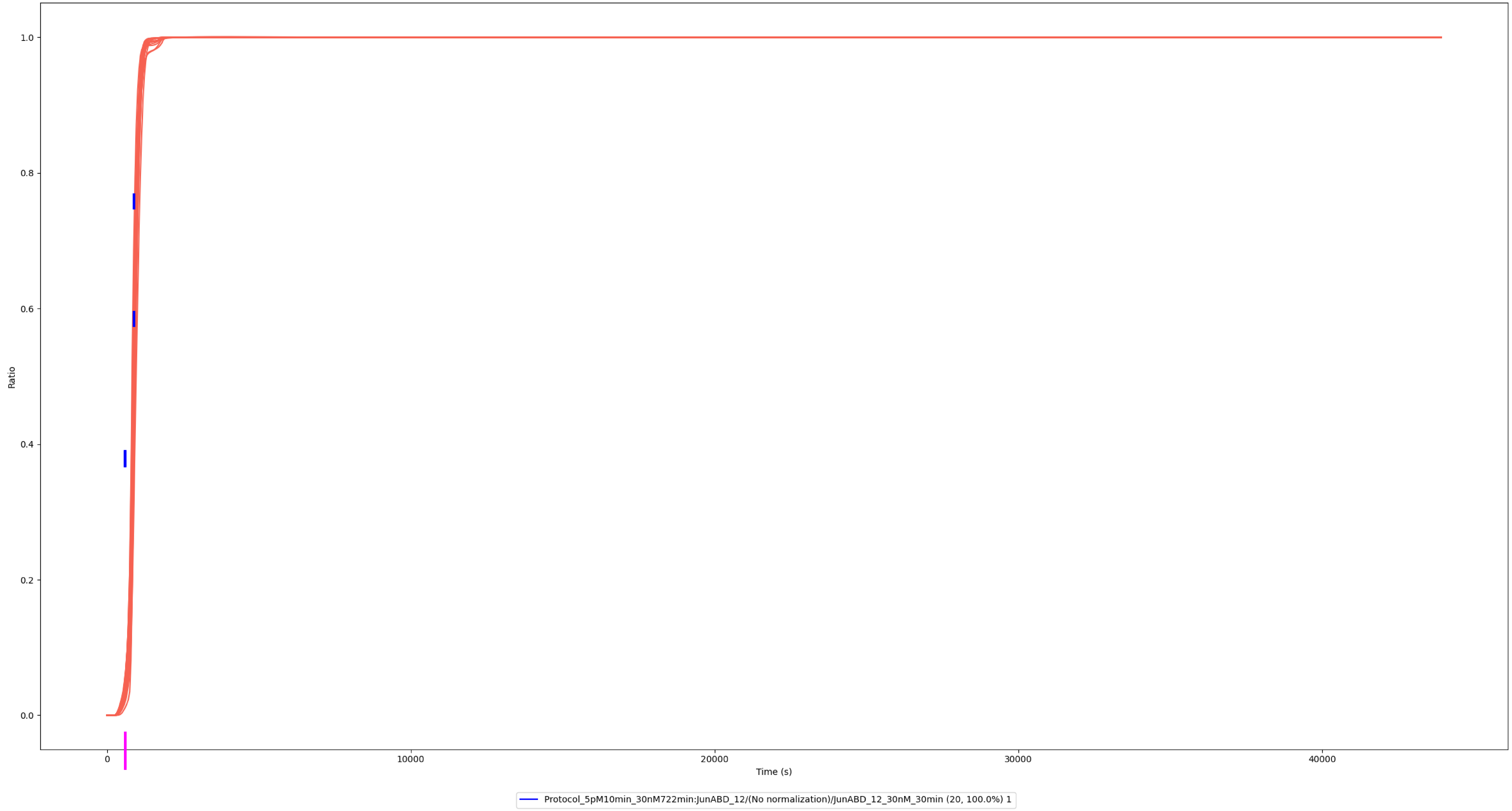

### Experiment vs. Simulation: 30 nM LPS, p-Jun/p-JunD

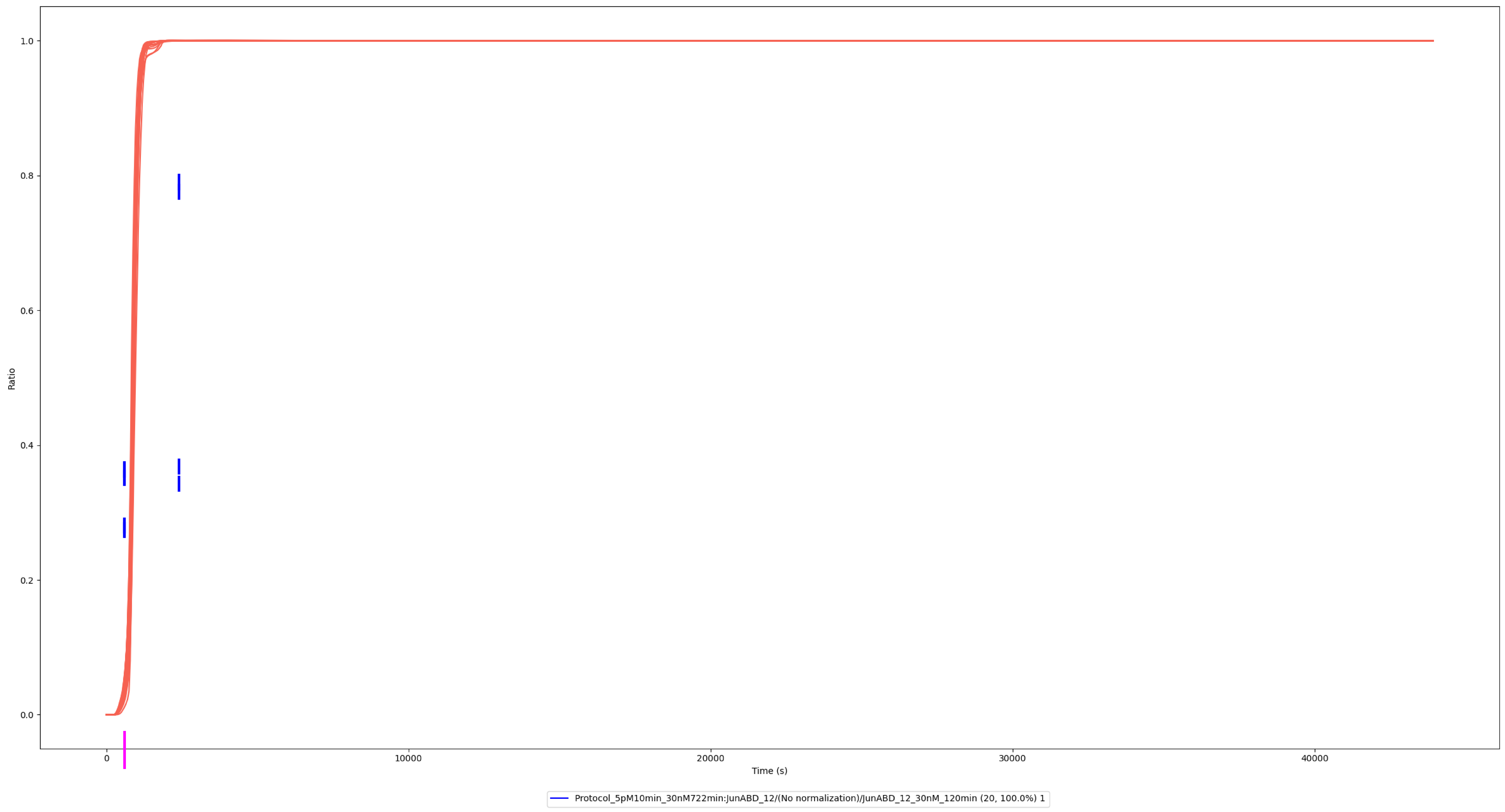
