## Supplemental Information for "Quantitative Modeling of TLR Signaling Reveals Missing Negative Feedback Guiding Identification of TANK-IKKε Checkpoint"

**Supplemental Methods:**

**Model Parameter Variation**

During model training, millions of pathway simulations were performed using SimAnalyzer. An *Analysis* is a set of simulations that used the same set of model parameters (the same set of intracellular and cell-surface reactant concentration values and the reaction rate values), but each simulation used a different *Protocol* (i.e., extracellular *LPS* concentration; the 14 *Protocols* are described below).

All the *Analyses* used the same set of model parameter initial values. For each *Analysis* and varied model parameter, a *Factor* value was generated using random sampling. SimAnalyzer can use advanced algorithms for sampling (e.g., Leaped Halton Sequence), but simple random sampling was used for this project. Each model parameter value was equal to its initial value multiplied by its *Factor*. The random sampling used log-normal distributions, and the median of each log-normal distribution was set to equal one (the *Mean* (equal to the mean of the natural logarithm of the distribution) was set to equal zero). These log-normal distributions had a Probability Density Function (PDF) equal to:

$$PDF\left( x \right)=\frac{1}{x\sigma\sqrt{2\pi}}exp\left[ {{-ln\left( x \right)}^{2}}/{2\sigma^{2}} \right]$$

where σ is the *Scale*, which equals the standard deviation of the natural logarithm of the distribution. These log-normal distributions had a Cumulative Distribution Function (CDF) equal to:

$$CDF\left( x \right)=\Phi\left( \frac{ln\left( x \right)}{\sigma} \right)$$

where Φ is the CDF of the standard normal distribution. The position x of the third quartile Q3 (i.e., the 75^th^ percentile) is related to σ by:

$$\sigma=\frac{ln\left( Q3 \right)}{\Phi^{-1}\left( 0.75 \right)}$$

Thus, if the value of Q3 is known, then σ, PDF(x), and CDF(x) are fully determined.

For the parameters having no measurement or estimate, an initial parameter value was set using the default value (described above), and the Q3 of the log-normal distribution was selected ad hoc such that the log-normal distribution CDF appropriately overlapped the corresponding eCDF (Supplemental Figure S2). If an initial parameter value was of higher confidence (with a calculated estimated accuracy), a narrower log-normal distribution was selected ad hoc to reduce the size of the parameter search space. Q3 values were selected based on the estimated confidence of each initial model parameter initial value (Supplemental Table S7, Supplemental Figure S3). The highest confidence parameters used a relatively narrow log-normal distribution with Q3 = 2.5, resulting in ~90% of the *Factor* values being within the range of 0.1 to 10. The lowest confidence values used a relatively wide log-normal distribution with Q3 = 15, resulting in ~90% of the *Factor* values being within the range of 0.001 to 1000.

As mentioned above, the concentration values of the extracellular reactants were not subjected to model parameter variation (unlike the intracellular and cell-surface reactants). Also, endo-deubiquitination reactions sometimes required a rapid subsequent *Dissociation Reaction* (k_off_ = 1000/s), and this k_off_ parameter was not varied. Finally, the IκBα/β k_prod_ parameters were not varied. The IκBα/β mRNA concentration and k_deg_ parameters were varied using Q3 set to 3.

Minimum and maximum constraints were used for each type of model parameter. For intracellular and cell-surface reactant concentrations, the range was 1 – 10^9^ copies/cell (939.67 fM – 939.67 μM), which should cover the full theoretical range. For comparison, we measured that there were 2.27×10^8^ copies of actin per mouse BMDM cell [1]. For k_on_, the range was 1 – 10^10^ /(M×s), which should cover the full theoretical range [2-6] For k_off_, k_trans_, k_cat_, and k_deg_, the range was 10^−5^ to 10^5^ /s, which should cover the full theoretical range [2, 3, 5, 7] If a sampled *Factor* value would result in a non-compliant parameter value, the *Factor* value was discarded, a new *Factor* value was randomly sampled, and this was repeated until the parameter value was compliant.

**Pathway Model Training**

During model training, model parameters were varied, simulations were performed (described below), and the results were compared to the corresponding experimental time-course data. In SimAnalyzer, each datum used for model training is termed a *Filter Constraint* (*FC*). In Simmune, *FCs* can be used as filters, but this feature was not used for this project. Each *FC* was a ratio defined by:

$$FC Ratio=\frac{Constraint Value}{Reference Value}$$

A *Constraint* and its corresponding *Reference* can be defined using identical or differing *Complexes*, *Protocols*, and/or time-points. A *Constraint Value* and its corresponding *Reference Value* can both simply be *Complex* concentration values. If so, the *FC Ratio* is defined as:

$$FC Ratio=\frac{Constraint Observable Complex Concentration}{Reference Observable Complex Concentration}$$

For example, in an earlier publication [8] pp-ERK1/2 (pp-ERK1 + pp-ERK2) were quantitated according to the equation:

$$FC Ratio=\frac{[pp\text{-}ERK1/2] at 1 nM LPS for 44 min}{[pp\text{-}ERK1/2] at 1000 nM LPS for 20 min}=0.3213$$

Thus, the *FC Ratios* were defined using the above normalized fold change values from the time-course experiments.

Alternatively, a *Constraint Value* and its corresponding *Reference Value* can both be concentration ratios. If so, the *FC Ratio* is defined as:

$$FC Ratio=\frac{\frac{Constraint Observable Complex Concentration}{Constraint Normalization Complex Concentration}}{\frac{Reference Observable Complex Concentration}{Reference Normalization Complex Concentration}}$$

For example, from [9], we used:

$$FC Ratio=\frac{\frac{[Nuclear RelA] at 0.3 nM LPS for 32 min}{[Total RelA] at 0.3 nM LPS for 32 min}}{\frac{[Nuclear RelA] at 30 nM LPS for 45 min}{[Total RelA] at 30 nM LPS for 45 min}}=0.6134$$

Note that freely diffusing and membrane-associated reactants are treated independently by *FCs* (*FC Constraints* and *References* can be defined using either a freely diffusing form or a membrane-associated form of a complex but not using the total quantity). Also, the *FC*s used concentration values of the *Complex*, not of the corresponding *Molecule(s)*.

For each experiment (i.e., time-course and/or LPS dose response curve), the time and dose of the maximum resulting abundance value was used as the *Reference*, and the other conditions were used as *Constraints*. After an *Analysis* was complete, a *Distance Score* was calculated for each *FC*:

$$Distance Score=w\times\left| {FC}_{exp}-{FC}_{sim} \right|$$

where *w* is a weight factor (for this project, the weight factor values were set to equal one), *FC_exp_* is the experimentally measured *FC Ratio*, and *FC_sim_* is the corresponding *FC Ratio* produced by the pathway simulations (the bar symbols indicate the absolute value). Note that in SimAnalyzer, *Ranking Weight* = *w* × *FC_exp_*.

During model training pathway simulations, some of the model parameter sets resulted in some of the reactants being essentially inactive (~0 copies/cell) over much of the time-course. This could result in an *FC Ratio* of approximately zero divided by zero. This error was avoided in SimAnalyzer by using a *Limit of Minimal Concentration* of 10 pM (10.642 copies/cell), and a *Penalty Score if Lower than Limit* of 10,000 (unless noted otherwise). For each *FC*, if the *Constraint Normalization Complex Concentration*, *Reference Observable Complex Concentration*, or *Reference Normalization Complex Concentration* was less than the *Limit of Minimal Concentration* (10 pM), then concentration ratios were not used to calculate a Distance Score. Instead, the *Penalty Score if Lower than Limit* (10,000) was used as the *Distance Score*.

In addition to the experimental *FCs*, a second type of *FCs* (“activation *FCs*”) was used. To ensure that model training would result in at least a minimal level of activation of the entire TLR4 pathway, each stably activated form of each *Complex* (i.e., not a transient form such as a typical enzyme:substrate complex), was used to define an activation *FC*. The same conditions (1 nM LPS for 62 min (*Constraint*) and 60 min (*Reference*)) and ratio value (*FC Ratio* = 1) were used for all 101 of the activation *FCs*. For example, for ppERK1 it was:

$$FC Ratio=\frac{[pp\text{-}ERK1] at 1 nM LPS for 62 min}{[pp\text{-}ERK1] at 1 nM LPS for 60 min}=1$$

Because of the short time difference (2 min) used to define all the activation *FCs*, the corresponding *Distance Scores* were insignificant unless the denominator was less than the *Limit of Minimal Concentration* (10 pM; if so, the *Penalty Score if Lower than Limit* (10,000) was incurred). Note that we previously reported that IκBα, pp-ERK1/2, and pp-p38α/β/γ/δ were all significantly activated (>20% of their maximal activation) in mouse BMDMs by stimulation with 1 nM LPS for 60 min [8].Three proteins (JNK3, p38γ, and p38δ) were excluded because their initial concentration was <200 copies/cell.

For each *Analysis* (i.e., set of simulations identical except for the *Protocol*), the *Discordance* was defined as the arithmetic mean of the *Distance Scores* calculated across all the *FCs*. The *Discordance* was used to rank the parameter sets. A lower *Discordance* indicated a better match to the experimental time-course data. The trained model was the model with the lowest *Discordance* value.

**Pathway Simulation**

SimAnalyzer was used to perform pathway simulations for model training in two steps. First, initial *Analyses* were performed using model parameters determined using random sampling (described above). Subsequently, *Analyses* were performed using parameters determined using an application of the GA [10].

All of the simulations (initial and GA) used the Rowmap ODE solver (Integration Method = GRK4T of Kaps-Rentrop, Maximum Dimension of the Krylov Space = 70, Minimum Step Size Reduction = 0.25, Maximum Step Size Increase = 2, Damping Factor = 0.8, Timesteps = 1000, Absolute Tolerance = 1.0E-15 M, Relative Tolerance = 1.0E-06, Maximum Complex Size = 12 *Molecules* (this is not a restriction for this model, as the *Association Reactions* cannot produce a complex larger than 12 *Molecules*)[11, 12]. Rowmap is not inherently restricted to output solutions having only positive values, and it occasionally produced erroneous negative reactant concentration values. To address this, if any reactant concentration value at any *in silico* time-point was less than -10 pM (-10.642 copies/cell), the simulation was halted and classified as failed, and the results were discarded.

The number of initial *Analyses* was 200,000 (200,000 *Analyses* × 14 *Protocols* = 2,800,000 pathway simulations). The model parameters for the initial *Analyses* were determined using random sampling (described above). After the initial *Analyses* were complete, GA *Analyses* were performed. Eighty-five GA iterations were performed with 2,000 *Analyses* per iteration (85 iterations × 2,000 *Analyses* × 14 *Protocols* = 2,380,000 simulations). During each GA iteration, the top four ranked *Analyses* were simply passed on to the next iteration (85 iterations × 4 redundant *Analyses* × 14 *Protocols* = 4,760 redundant simulations; these were not actually resimulated), and 1,996 novel parameter sets were generated and used to perform new simulations. After each GA iteration, a *Homogeneity* test was performed. The arithmetic mean of the smallest 10% of the *Discordance* scores was calculated. This value was used as the central value, and the (arithmetic) mean absolute deviation was calculated. If this value was less than 0.1, *Homogeneity* was set to be true. Otherwise, *Homogeneity* was set to be false. If *Homogeneity* was true, 100 *Analyses* (5%) simply used randomly sampled model parameters using the same algorithm as the initial *Analyses*, and the remaining 1,896 *Analyses* were performed using GA pairing, crossing-over, and mutation. If *Homogeneity* was false, no random sampling was performed, and the 1,996 *Analyses* were performed using GA pairing, crossing-over, and mutation.

During the GA analyses, *Analyses* first underwent pairing to determine parent *Analysis* pairs to be used to produce offspring model parameter sets for the subsequent GA iteration. The *Analyses* were ranked by their *Discordance* from 0 to NA − 1, where NA is the number of *Analyses* per GA iteration (Rank 0 was the top ranked *Analysis*). Each parent *Analysis* was determined by calculating a Rank value using random sampling: Rank = RoundDown(NA × RndExp). The RoundDown function rounds a number downward to the nearest integer. RndExp is a value randomly sampled from an exponential probability distribution with Rate Parameter equal to 0.95 (this is called the GA *Selection Strength* in SimAnalyzer). Further, the values of RndExp were restricted to being ≥0 and <1 (noncompliant values were discarded, RndExp was resampled, and this was repeated until RndExp was compliant). If the same Rank value was determined for both parents, a new Rank value was calculated until the two parents were different.

For each GA iteration, pairs of parents were determined. Each pair of parents produced two offspring. All the varied model parameters were ordered in a one-dimensional array. The two offspring were produced by performing a single crossing-over operation at a random location in the array, and then by performing mutation.

The GA *Mutation Probability* was set to 25%, and the GA *Mutation Width* (*MW*) was set to 0.1. Mutation of a *Factor* was performed by multiplying the current *Factor* value by a *Mutation Factor* *(MF)* value where MF = (1 − MW) + (2 × MW × R) where R was randomly sampled from a uniform distribution ranging from 0 to 1. If the resulting model parameter was outside of the minimum to maximum range (described above), the MF value was discarded, R was resampled, and this was repeated until the model parameter was compliant. The GA was implemented such that it ignored the *Rate Constraints* (described above).

There was a total of 5,175,240 non-redundant simulations (2,800,000 + 2,380,000 − 4,760). The maximum simulation duration was set to 60 min (wall-clock time), and 2,222 (0.6%) of the 369,660 non-redundant *Analyses* failed because a simulation timed out. A total of 138,135 (37.4%) *Analyses* failed for any reason. A total of 231,525 non-redundant *Analyses* (3,241,350 simulations) finished successfully. Model training took three weeks to perform using our workstation powered by two AMD EPYC Rome 7702 CPUs for a total of 128 cores and 256 threads.

**Homomer Modeling**

Coarse-grained modeling was used to model homomeric complexes. Each homomeric complex was modeled using a single *Molecule* (rather than using multiple individual copies; the *Molecule* concentrations were adjusted accordingly). For example, the myddosome model contained one copy of *Molecule MyD88_x6CG* to represent six copies of MyD88. The molecules modeled in this manner were IKKα (hexamer), IKKβ (hexamer), IRAK1 (tetramer), IRAK2 (tetramer), IRAK4 (tetramer), MAL (tetramer), MyD88 (hexamer), NEMO (hexamer), PI(4,5)P_2_ (tetramer, it forms a 4:4 complex with MAL), and TRAF6 (trimer). Some TLR4 signaling pathway molecules are statically homomeric irrespective of TLR4 signaling. For example, ELK1 is a stable homodimer. Statically homomeric molecules were modeled as homomers (e.g., *Molecule ELK1_x2*).

**Homolog Modeling**

Model training (described below) used experimental time-course data that included relative abundance measurements of homologs. For example, we performed western blots (described above) to measure the relative abundance of pp-ERK1/2 (pp-ERK1 + pp-ERK2) in LPS stimulated mouse BMDMs. Simmune does not include an ability to sum abundance values of multiple *Complexes* during model training. Consequently, a *Molecule* named *ERK12* was defined with *Features* *ERK1* and *ERK2*. ERK1 was modeled as *Molecule ERK12* with *Feature ERK1* set to *On* and *Feature ERK2* set to *Off*, and likewise for ERK2. Using *ERK12*, Simmune was able to use the relative abundance values of pp-ERK1/2 from the western blots. Using this strategy, molecule-nonspecific experimental values were used for model training, while each homolog was nevertheless individually modeled (Supplemental Table S1).

**Polyubiquitination Modeling**

Three types of polyubiquitination (K48, K63, and K63-M1) were modeled using both a *Feature* and a *Binding Site*. For example, TRAF6 can be K63 polyubiquitinated with M1 branches, and NEMO can bind to TRAF6 at its M1 polyubiquitin (*Feature* *K63M1Ub* = *On*, *Binding Site* *M1Ub_BS7*). The reaction network included endo-deubiquitination reactions. Consequently, this could require rapid subsequent dissociation. For example, for the NEMO-TRAF6 complex, If TRAF6 underwent K63 endo-deubiquitination, NEMO was set to dissociate quickly from TRAF6 (k_off_ = 1000/s). This k_off_ parameter was not varied in SimAnalyzer during model training (model training is described below).

**Macrophage Modeling**

Spatially resolved modeling of the intracellular space was not performed in this study. Nevertheless, SimAnalyzer required the volume and surface area of the *in silico* mouse macrophage cell. We found that suspended mouse BMDMs examined by light microscopy were spherical with a diameter of 15.46 ± 4.0 μm (median ± interquartile range). This morphology is consistent with mouse BMDM measurements done by others [13-16]. Thus, the *in silico* mouse macrophage was modeled as a sphere with a diameter of 15 μm, (cell surface area = 706.86 μm^2^, and cell volume = 1767.15 μm^3^). Consequently, 1 copy/cell is equivalent to 0.93967 pM. Because the simulations were not spatially resolved, localization was modeled using a *Feature* and translocalization (e.g., RelA nuclear import) was modeled using a *Transformation Reaction*.

**Model Simulation Protocols**

Fourteen SimAnalyzer *Protocols* were created. They were identical except for the extracellular concentration of unbound LPS at t = 600 sec. The *in silico* timespan was from 0 to 43920 sec = 732 min = 12.2 h. The extracellular unbound LBP concentration at t = 0 sec was set to 183 nM, which is the basal plasma concentration [17, 18]. The extracellular unbound MD2 concentration at t = 0 sec was set to 195 pM, which is the basal plasma concentration [19, 20]. The extracellular unbound LPS concentration at t = 0 sec was set to 5 pM, which is approximately the basal plasma concentration (the LPS concentration range is undetectable to 200 pg/ml in healthy human plasma, and it is 17 pg/ml in basal mouse plasma) [21]. The extracellular unbound LPS concentration at t = 600 sec was set to 0.01, 0.03, 0.1, 0.3, 1, 1.5, 3, 10, 15, 30, 100, 150, 300, or 1000 nM. Unlike the intracellular and cell surface *Molecules*, the concentration values of the extracellular *Molecules* were not subjected to variation (SimAnalyzer does not have this ability; parameter variation during model training is described below).

**Western Blots**

Cells were stimulated with 1, 10, or 100 nM lipopolysaccharide (LPS; Kdo2-Lipid A from *E. coli*, Avanti Polar Lipids, Birmingham, AL) for the indicated time. SDS-PAGE and western blots were performed for p-JNK1 (pTyr185 and pThr183/pTyr185; 81E11), p-ERK1/2 (ERK1 + ERK2, pThr202 and pThr202/pTyr204; D13.14.4E), p-IKKα/β (IKKα pSer176 + IKKβ pSer177; C84E11), pp-MKK1/2 (MKK1 + MKK2, pSer217/pSer221; 41G9), and pp-ERK1/2 (ERK1 + ERK2, pThr202/pTyr204; E10) (Cell Signaling Technology Inc., Danvers, MA).

**Supplemental Protocol and Scheme:**

**Protocol for Protein Complex Structure Model Refinement (see Scheme 1)**

**1 pLDDT criterion:**

Go over all residues *I* in the complex and classify them according to pLDDT.

if I has a pLDDT ≤ 60.0 set I as I_u_

if I has a pLDDT > 60.0 set I as I_c_

where subscripts u and c stand for *unconfident* and *confident* predictions. This criterion defines the set {c/u}_1_ of high (s_c_) and low (s_u_) quality fragments, s = 1, 2, ..., N_0_, for the raw AF model.

**2 Add length criterion:**

Go over the N_0_ fragments s in {c/u}_1_ and determine their length (s), defined as the number of residues in s.

if s = s_u_ and (s) ≤ 8 switch all residues I in s to I_c_ (possibly a flexible/disordered loop or linker)

if s = s_u_ and (s) > 8	keep all residues I in s as I_u_

if s = s_c_	keep all residues I in s as I_c_

This criterion redefines {c/u}_1_ as the new set {c/u}_2_ of fragments s = 1, 2, ..., N ≤ N_0_.

**3 Add interfacial criterion:**

Go over the N fragments in {c/u}_2_ and find all the s_u_ fragments that interface with at least one s_c_.

For each s_c_ and s_u_ pair, measure the distances r_IJ_ between all (N_u_) C_a_ atoms of I in s_u_ and all (N_c_) C_a_ atoms J in s_c_

if r_IJ_ ≤ 7.0 Å I is interfacial; switch I from I_u_ to I_c_ and issue warning

if r_IJ_ > 7.0 Å I is not interfacial; keep I as I_u_

Once the u/c reassignment of residues in s_u_ is completed, calculate the fraction *f* = N_u2c_/N_u_, where N_u2c_ is the number of residues in s_u_ that switched from u to c.

if *f* ≥ 0.6 switch all remaining I_u_ to I_c_

if *f* < 0.6 do nothing

This criterion defines a new set {c/u}_3_ of fragments s = 1, 2, …, Μ ≤ N

**4 Add terminal-domains criterion:**

Go over the M fragments in {c/u}_3_ and identify the first (s_u,NTD_) and last (s_u,CTD_) s_u_ fragments of each chain *P*.

if s_u,NTD_ = 2 and s = 1 is a s_c_ fragment and not interfacial (criterion 3) with any other s_c_ switch all I in s = 1 from I_c_ to I_u_

if s_u,CTD_ = n*_P_* – 1 (where n*_P_* is the number of segments in chain *P*) and fragment s = n*_P_* is a s_c_ fragment and not interfacial switch all I in s = n*_P_* from I_c_ to I_u_

This criterion eliminates loose NTD and CTD domains not involved in interfacial events.

**5 Final structures:**

Remove all remaining s_u_ fragments from the system.

**Workflow Scheme:**

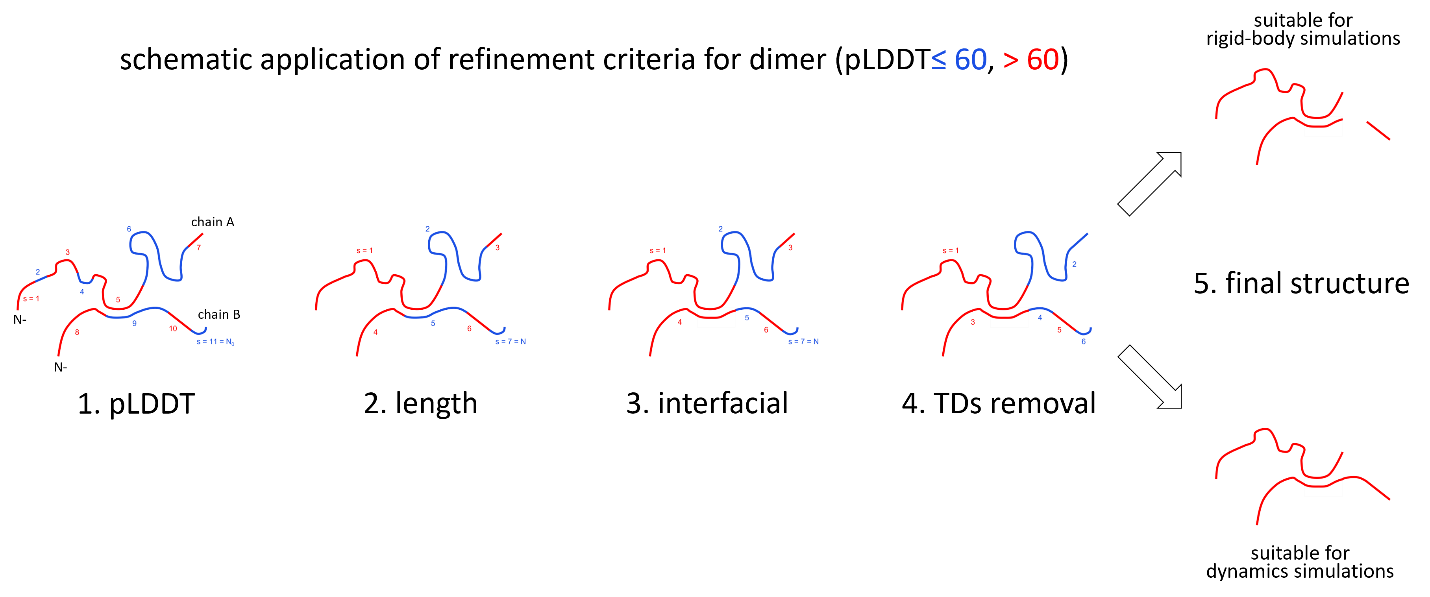
