## Supplemental Figures for "Quantitative Modeling of TLR Signaling Reveals Missing Negative Feedback Guiding Identification of TANK-IKKε Checkpoint"

### Slide 1
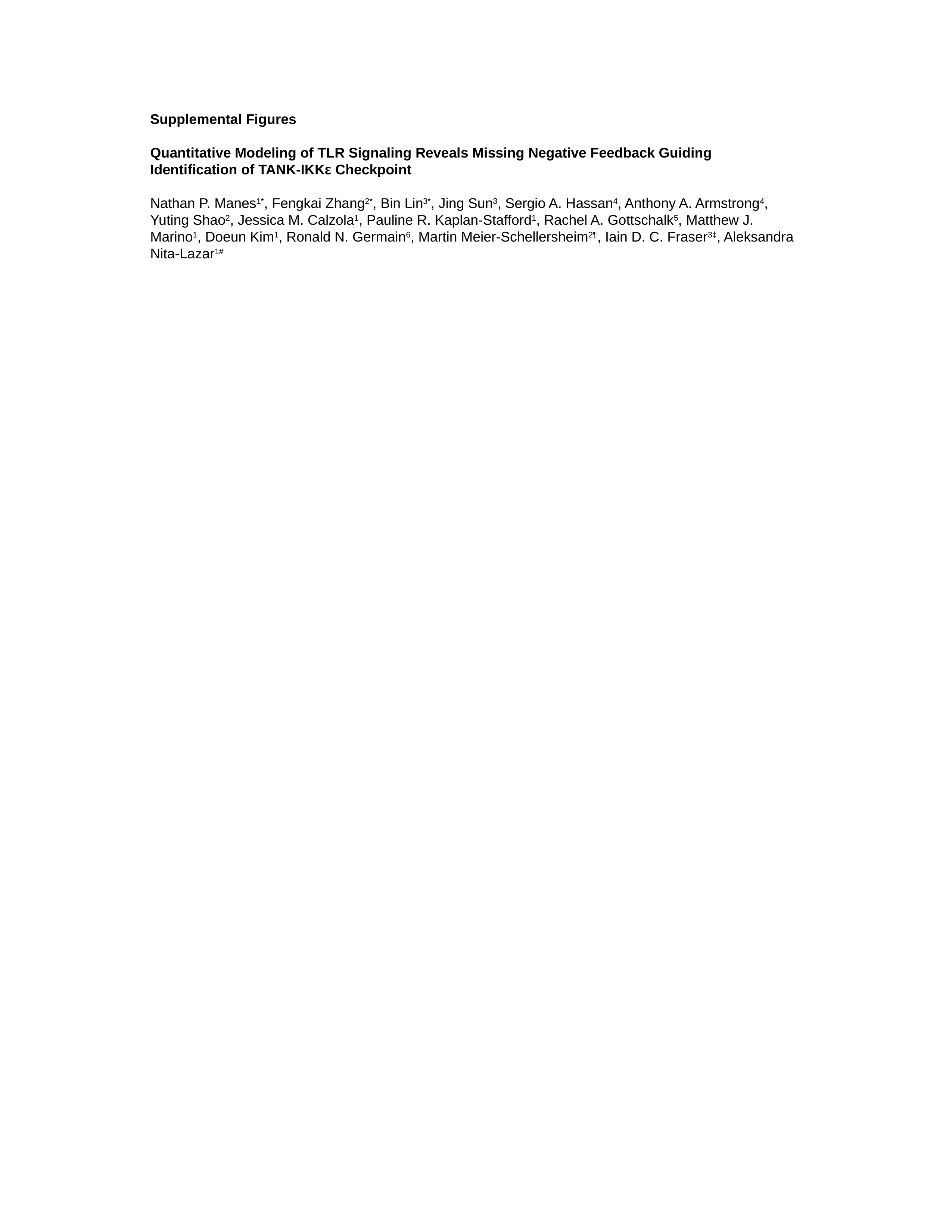

Supplemental Figures
Quantitative Modeling of TLR Signaling Reveals Missing Negative Feedback Guiding Identification of TANK-IKKε Checkpoint
Nathan P. Manes^1*^, Fengkai Zhang^2*^, Bin Lin^3*^, Jing Sun^3^, Sergio A. Hassan^4^, Anthony A. Armstrong^4^, Yuting Shao^2^, Jessica M. Calzola^1^, Pauline R. Kaplan-Stafford^1^, Rachel A. Gottschalk^5^, Matthew J. Marino^1^, Doeun Kim^1^, Ronald N. Germain^6^, Martin Meier-Schellersheim^2¶^, Iain D. C. Fraser^3‡^, Aleksandra Nita-Lazar^1#^

### Slide 2
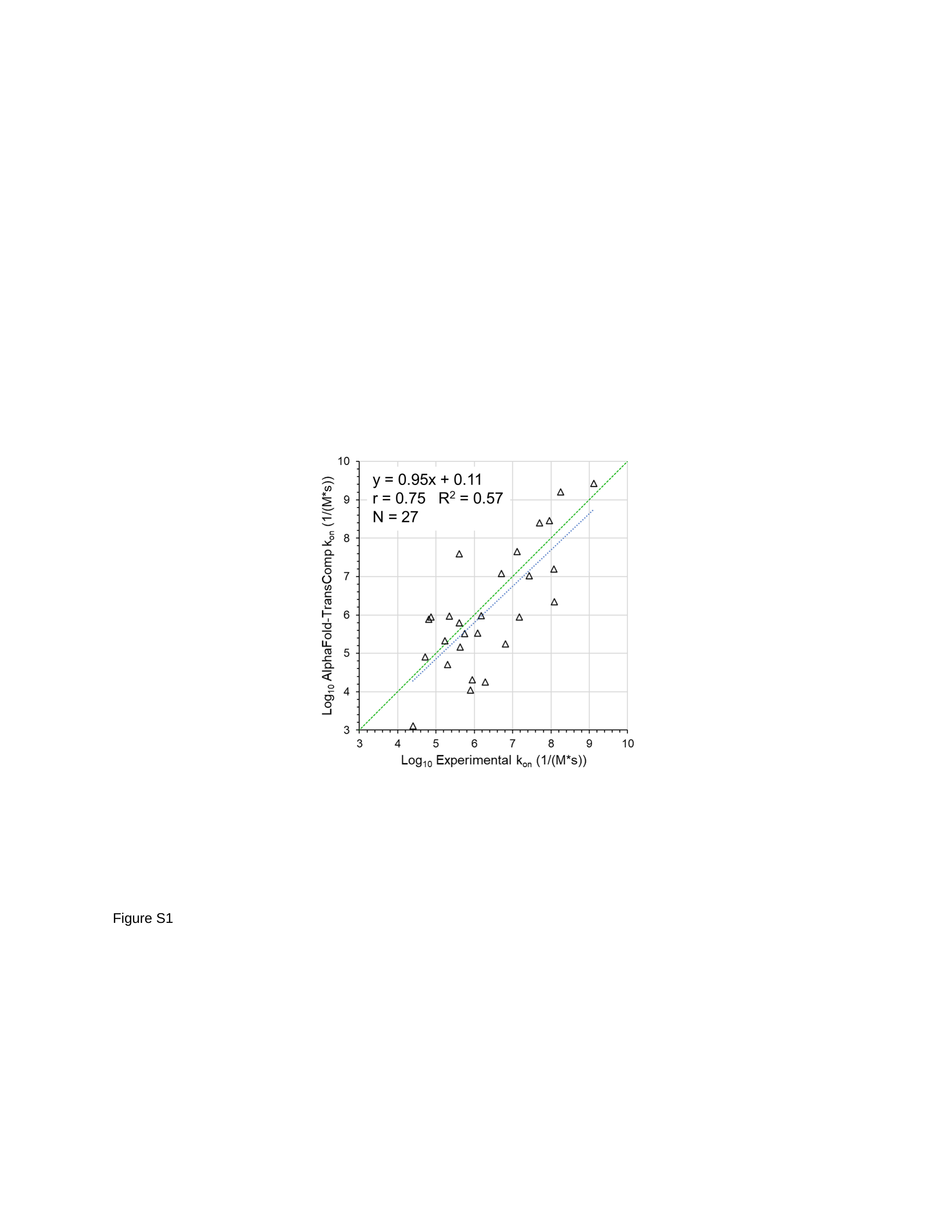

Figure S1

### Slide 3
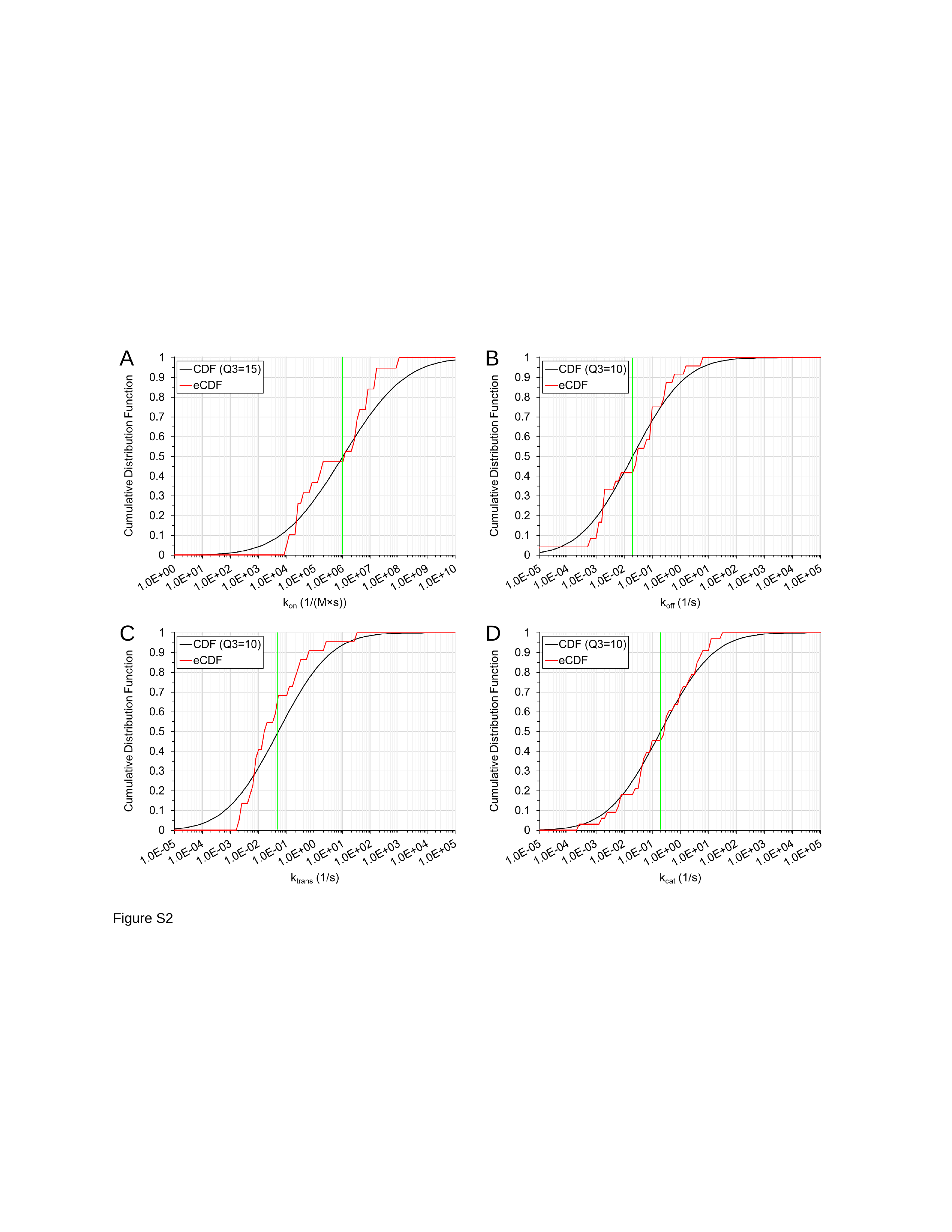

A
B
C
D
Figure S2

### Slide 4
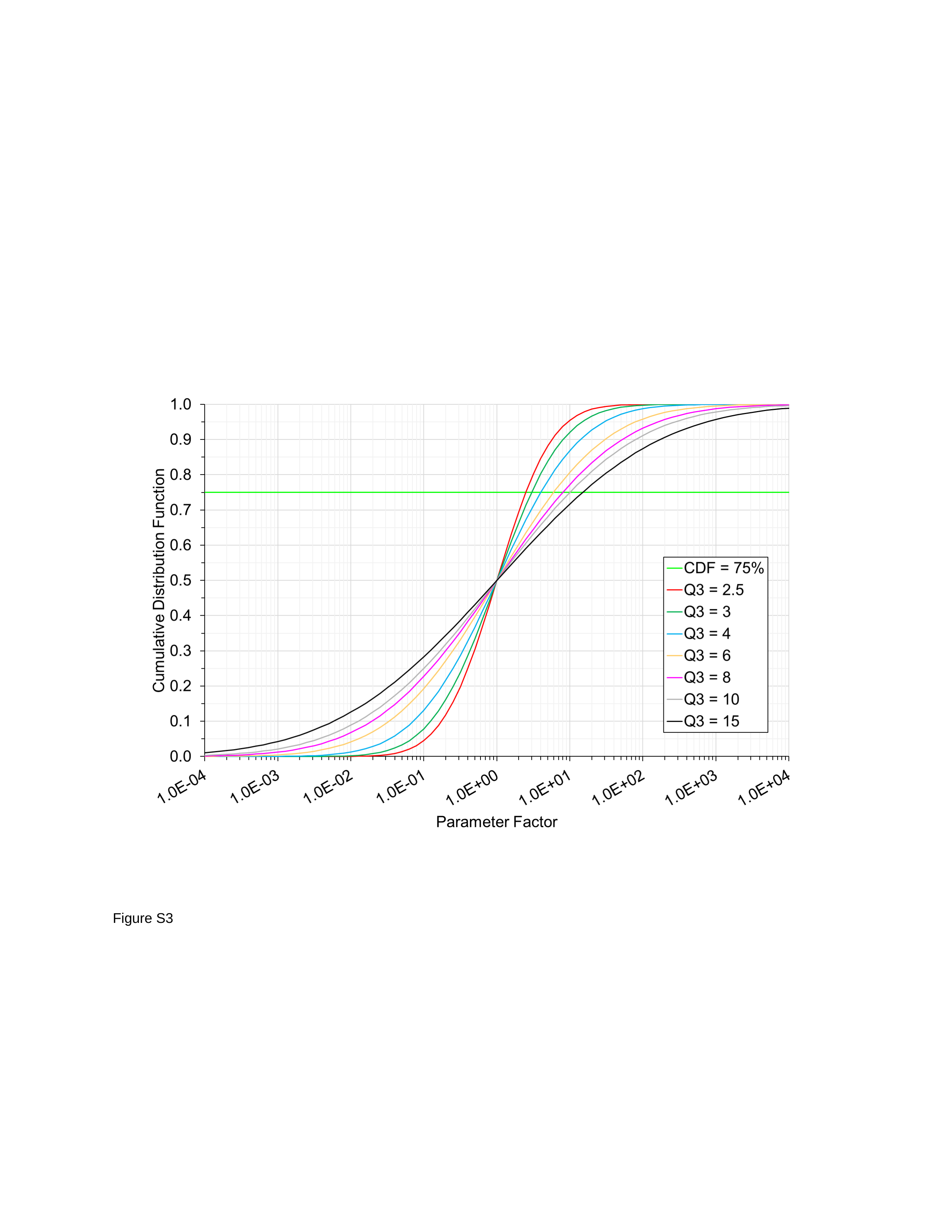

Figure S3

### Slide 5
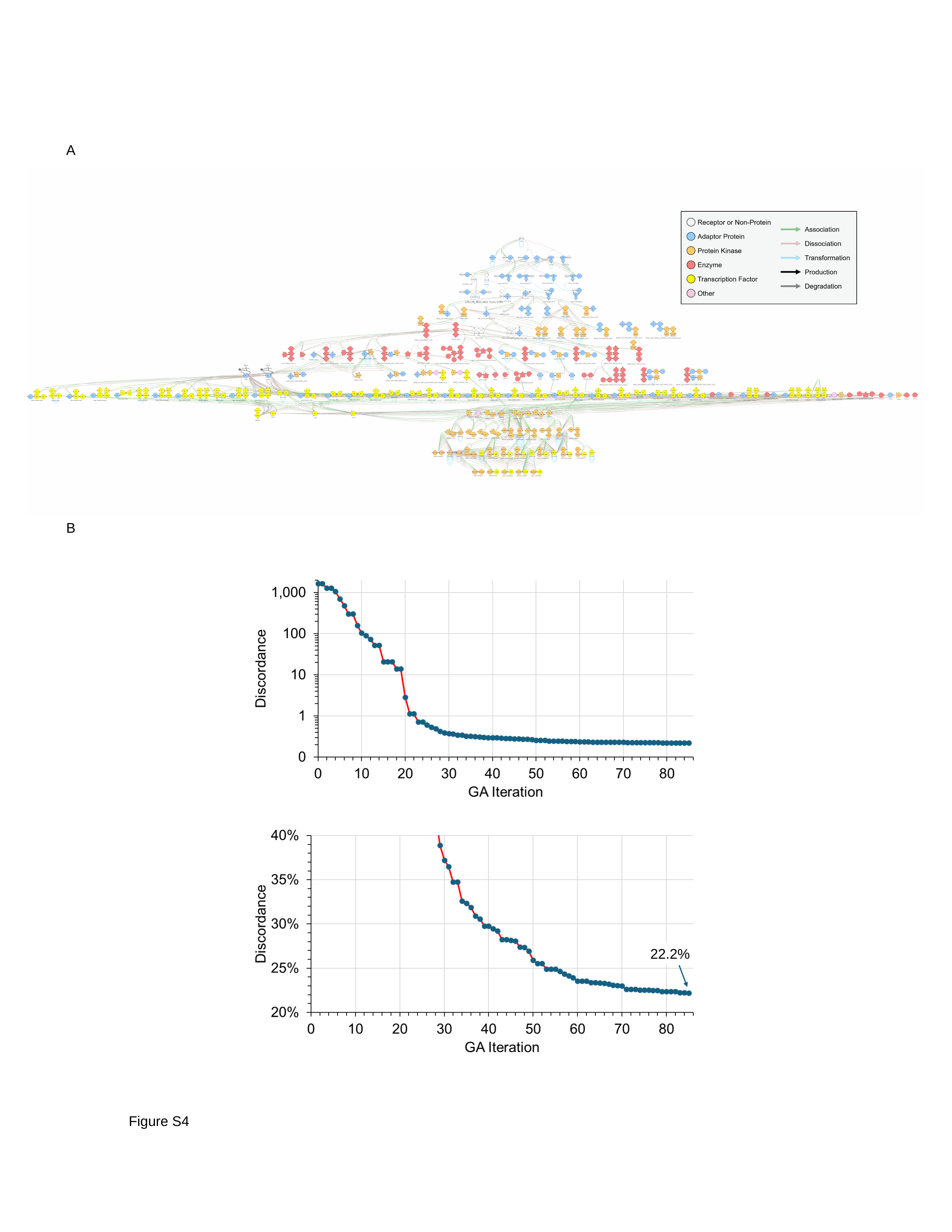

A
B
22.2%
Figure S4

### Slide 6
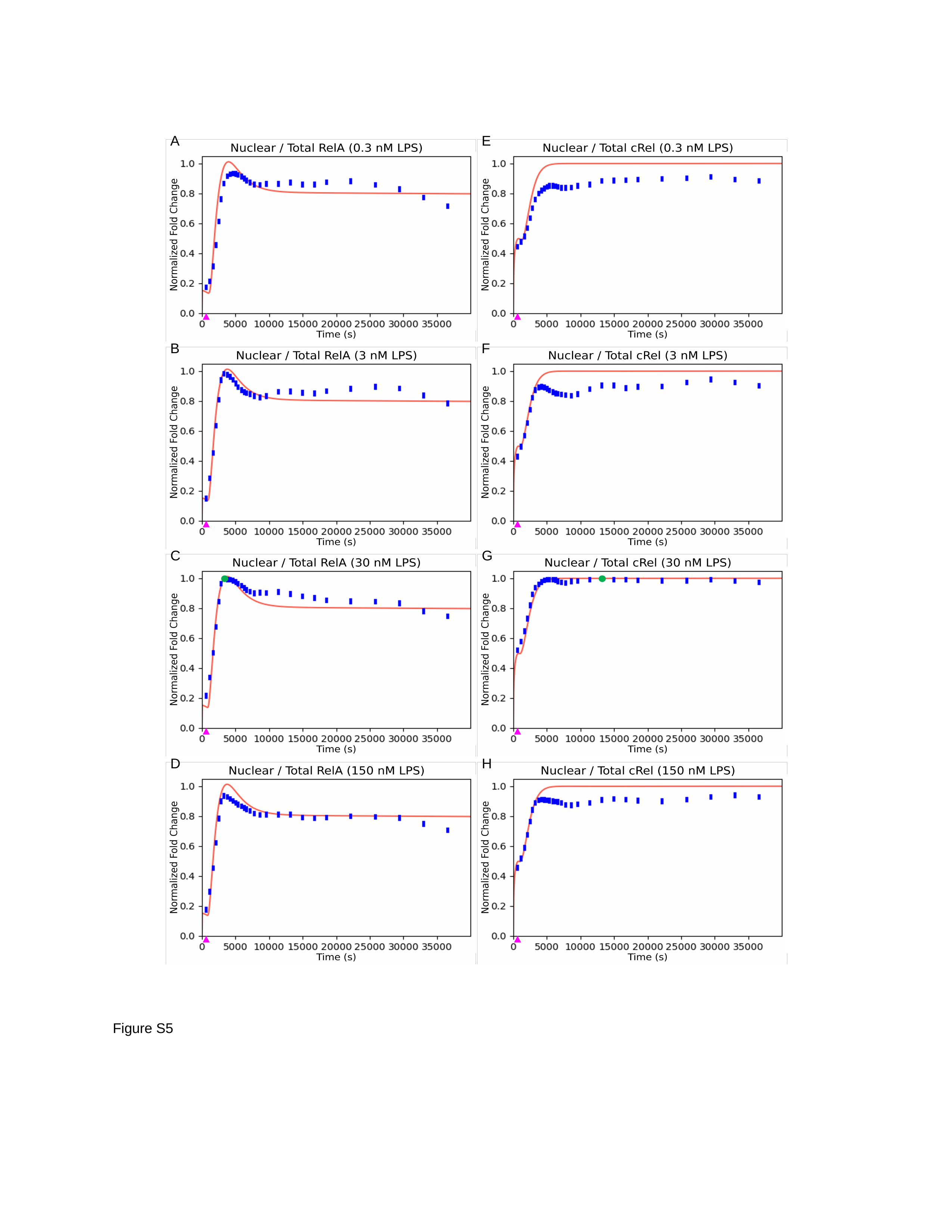

A
E
B
F
C
G
D
H
Figure S5

### Slide 7
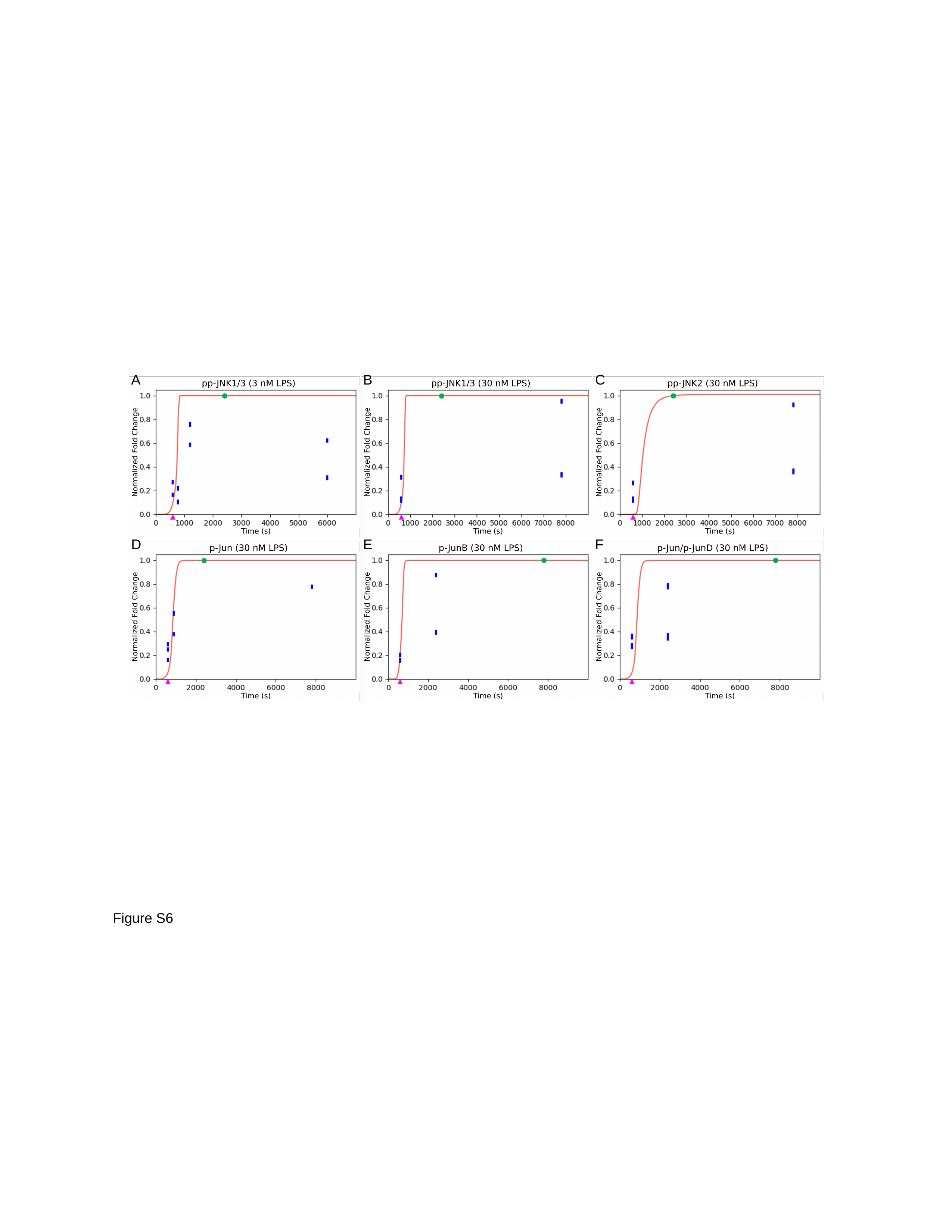

A
B
C
D
E
F
Figure S6

### Slide 8
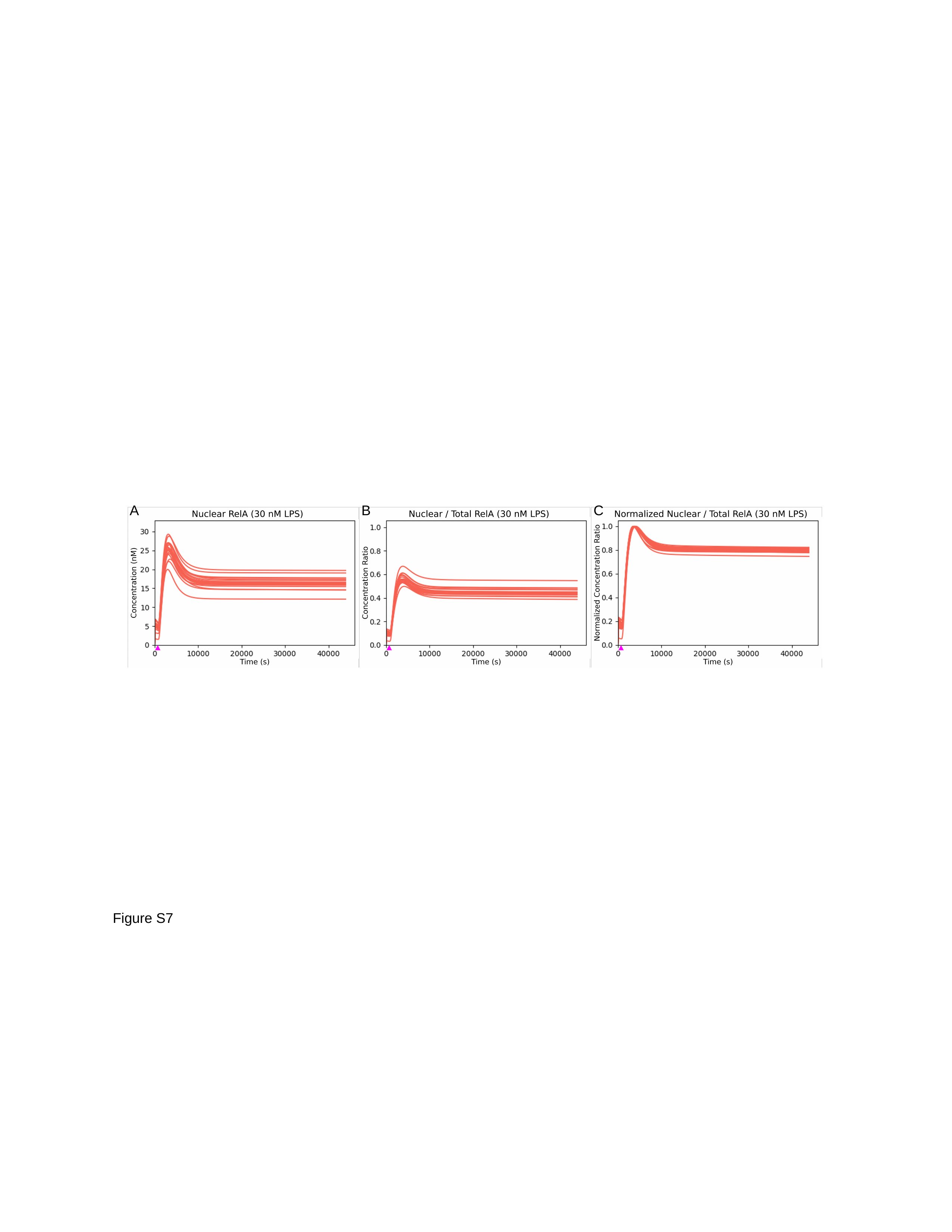

A
B
C
Figure S7

### Slide 9
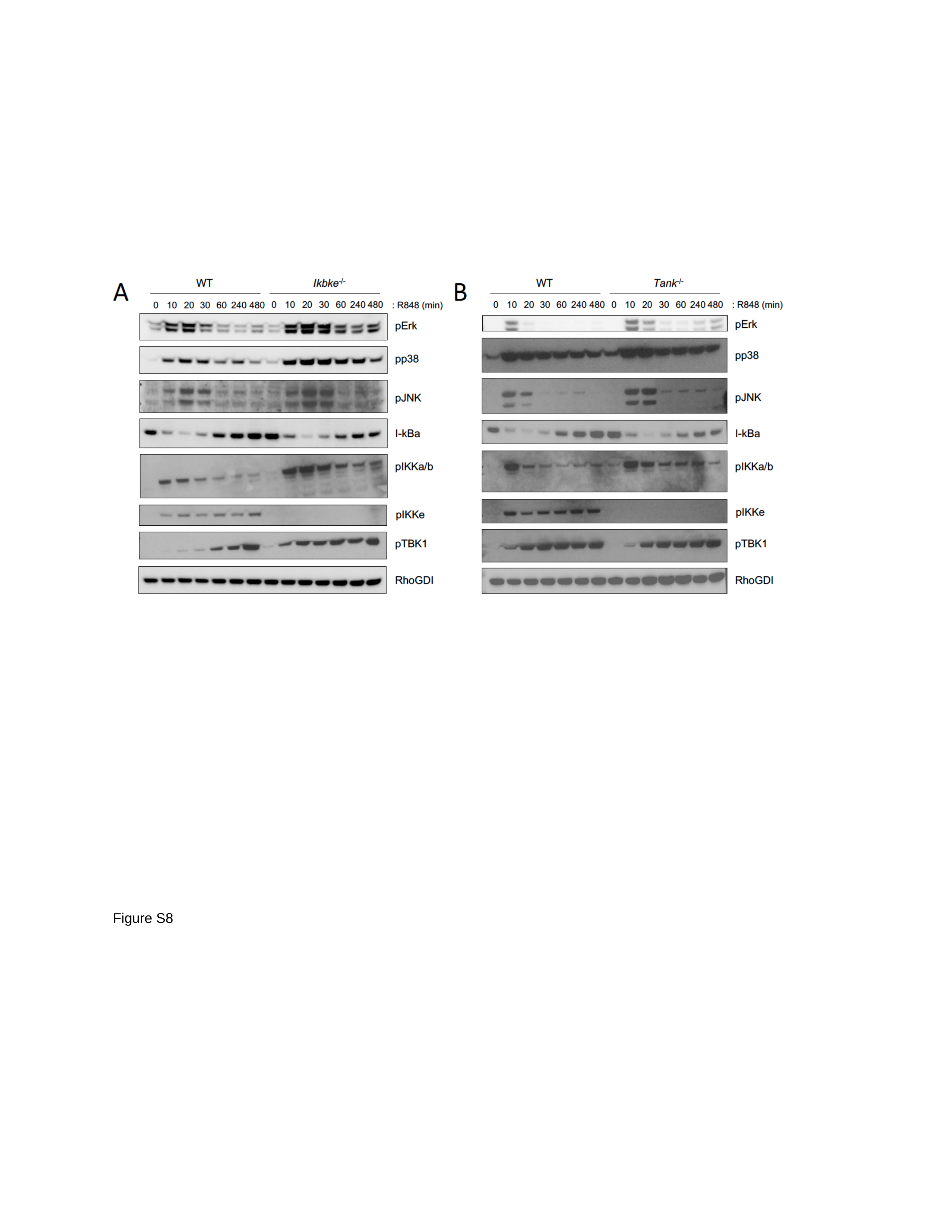

Figure S8
